# A lineage-resolved multimodal single-cell atlas reveals the genomic dynamics of early *C. elegans* development

**DOI:** 10.1101/2024.12.02.626321

**Authors:** Servaas N. van der Burght, Francesco N. Carelli, Alex Appert, Yan Dong, Matthew Hill, Toby Buttress, Richard Butler, Julie Ahringer

**Author notes:** Authors contributed equally to this work (Servaas N. van der Burght, Francesco N. Carelli, Alex Appert, Yan Dong).

## Abstract

Multimodal single-cell profiling provides a powerful approach for unravelling the gene regulatory mechanisms that drive development, by simultaneously capturing cell-type-specific transcriptional and chromatin states. However, its inherently destructive nature hampers the ability to trace regulatory dynamics between mother and daughter cells. Taking advantage of the invariant cell lineage of Caenorhabditis elegans, we constructed a lineage-resolved single-cell multimodal map of pre-gastrulation development, which allows the tracing of chromatin and gene expression changes across cell divisions and regulatory cascades. We characterise the early dynamics of genome regulation, revealing that zygotic genome activation occurs on an accessible chromatin landscape pre-patterned both maternally and zygotically, and we identify a redundant family of transcriptional regulators that drive a transient pre-gastrulation program. Our findings demonstrate the power of a lineage-resolved atlas for dissecting the genome regulatory events of development.

## Introduction

Starting from a single cell, a whole animal is generated by differential interpretation of the genome sequence. While we are beginning to understand the regulatory networks underlying differentiated cell types, we still know little about the genomic controls of development. Unlocking the dynamics of genome regulation is central to understanding how organisms are built.

Single-cell profiling provides the means to address this. Profiling gene expression in tissues and organisms has transformed our understanding of cell-type diversity (Rood et al. 2024), and the ability to profile chromatin accessibility (an indicator of regulatory element activity) in parallel has enabled investigation of the genome regulatory controls of the observed cellular gene expression patterns (Flynn et al. 2023). However, profiling is inherently destructive, hampering study of the dynamic regulation that controls development. Computational approaches are used to infer cellular progression, and the impressive recent application to co-profiling data from Drosophila and zebrafish embryogenesis has made key progress towards tracking genomic changes across developmental trajectories (Liu et al. 2024; Kim et al. 2024; Calderon et al. 2022). However, the methods are unable to establish direct transitions through cell divisions. The ability to co-track gene expression and chromatin accessibility changes between mother and daughter cells would illuminate how the genome is read.

Due to its invariant lineage (Sulston et al. 1983), C. elegans is an organism where this can be done. Profiles of cells from embryos of different ages can be captured and then mapped to their position on the lineage tree, thereby linking mother and daughter cell profiles. A pioneering study profiling gene expression across C. elegans embryogenesis showed the feasibility and power of this approach (Packer et al. 2019). However, the map lacked genome state information so the genome regulatory processes that drive the observed cellular differences could not be determined.

To investigate the genomic and transcriptomic unfolding that underlies the beginning of development, we have generated a map of gene expression and chromatin accessibility in single nuclei up to the gastrulation stage. We show that zygotic genome activation occurs on a pre-patterned accessible genome, with some accessible sites maternally inherited and some newly induced in the zygote. We uncover a transient gene expression program shared by pre-gastrulation cells that starts at ZGA, and a family of redundant transcriptional regulators that control the program. The map also reveals stepwise genomic regulation across the different early lineages. Our results illuminate the initial events of genome regulation and demonstrate the power of lineage-resolved multimodal analyses.

## Results

### A lineage resolved single-cell multimodal atlas of *C. elegans* pre-gastrulation development

We generated a single-cell multimodal atlas of chromatin accessibility and gene expression in early development using the 10X Genomics multiome platform (Figure 1A, B; Supplementary Figure 1). Using embryo collections containing predominantly pre-gastrulation embryos, we co-profiled nuclear transcription and chromatin accessibility in single nuclei (see methods). Following quality controls, we obtained 48775 high-quality nuclei across five biological replicates, with a median of 1091 detected genes and 1809 ATAC fragments per profiled nucleus.

**Figure 1.**
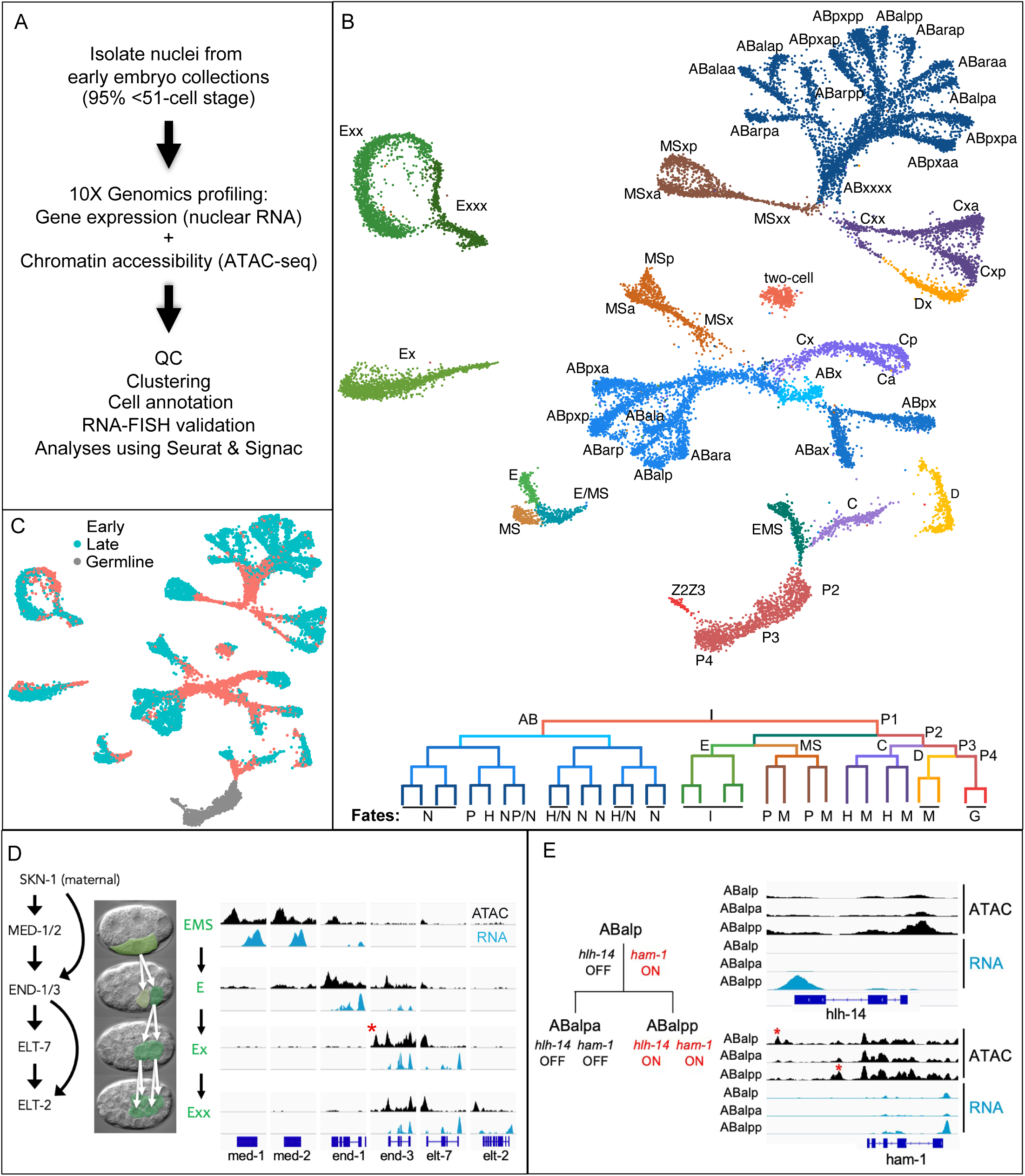
A multimodal atlas of *C. elegans* embryogenesis. A) Experimental pipeline. B) Top: UMAP visualization and cell type annotation of pre-gastrulation cell types and selected post-gastrulation cell types. Bottom: Cell lineage tree representing cells in the top panel, and the the types of cells they will produce (N: neuron; P: pharynx; H: hypodermis; I: intestine; M: muscle; G: germline). C) Early and late-cell cycle gene annotation. D) Mother to daughter cell changes in chromatin accessibility and gene expression across the the transcription factor cascade specifying intestinal cell fate in the E- lineage visualised by ATAC-seq (black) and RNA-seq (blue) signals. Red asterisk indicates element open in *end-3* for only one cell cycle. E) Chromatin and gene expression differences between ABala and its two daughters, ABalpa and ABalpp at the *hlh-14* and *ham-1* loci. ATAC-seq (black) and RNA-seq (blue)

After clustering analyses, we annotated pre-gastrulation cells and a small number of cells at the beginning of gastrulation using cell-type markers (Supplementary Table 1 and 2) from the literature (Tintori et al. 2016; Packer et al. 2019; Cole et al. 2024) and RNA FISH validation experiments (Supplementary Figure 1). The remaining post-gastrulation cells were assigned to a lineage group (Supplementary Figure 2 and Supplementary Table 3). As previously observed (Packer et al. 2019; Cole et al. 2024), some small groups of cells in the pre-gastrulation map (e.g., bilaterally symmetric cells) had indistinguishable expression profiles. The earliest cluster consists of nuclei from 2-cell embryos (AB and P1 are unresolved); almost no nuclei from one-celled zygotes were profiled as most had divided by the time nuclei were isolated. In all, the early embryo multimodal map contains 58 different clusters, which contain 192 to 1431 nuclei per cluster (Figure 1B, Supplementary Figure 3, Supplementary Table 4).

We observed that some clusters exhibited profiles similar to two annotated sister cells. For example, we labeled one cluster as MSx because it shared characteristics with both MSa and MSp, and another as ABxxxx, representing all 16 AB cells at that stage. These clusters showed high expression of replication-coupled histone genes (Supplementary Figure 4), suggesting that they may represent cells recently born and in an early cell cycle stage. To test this, we performed RNA FISH for pes-10 across the AB16 cell cycle, as it is more highly expressed in the shared ABxxxx cluster than in individually annotated AB16 cells.

Supporting our hypothesis, early AB16 cells exhibited higher pes-10 levels compared to late AB16 cells (Supplementary Figure 4).

After training a random forest classifier on a limited, high-confidence set of early cell cycle clusters, we developed a model to annotate early cell cycle nuclei (Fig. 1C). For simplicity we labelled classified cells as “early” and others as “late”, as the specific cell-cycle period captured by the model remains unknown. Using these annotations, we observed gene-specific differences in transcription and chromatin accessibility between early and late cells (Supplementary Figure 5). Further refining the model should illuminate the interplay between genome regulation and the cell cycle.

To assess the temporal resolution of our map, we analyzed chromatin and gene expression dynamics during the well-studied early intestinal transcription factor cascade in C. elegans. The early part of the cascade begins with the activation of med-1 and med-2 in the EMS cell at the 4-cell stage by the maternal transcription factor skn-1 and culminates in the expression of elt-2 when there are four intestinal cells (McGhee 2013) (Figure 1D).

Our results align with known expression timing (McGhee 2013) and reveal striking temporal dynamics across the cascade (Figure 1D). As expected, the promoters of med-1 and med-2 are open and the genes are transcribed in EMS but promoters are closed and transcription halted in its daughter E. In E, transcription of the targets end-1 and end-3 begins. end-3 is transiently expressed, turning off in Ex, while end-1 continues to be expressed in Ex, where a new regulatory element opens briefly before closing in the Exx daughters. elt-7 begins transcription in the Ex though its promoter opens earlier, in E. Finally, the promoter of elt-2 opens, and its transcription begins, when there are four E daughters (Exx). Such stepwise changes are seen in all lineages (e.g. Figure 1E), underscoring the map’s fine temporal resolution.

### Zygotic genome activation starts on a pre-patterned genome

In animals, the single-celled totipotent zygote is initially transcriptionally quiescent, regulated by maternally inherited proteins and RNAs (Kojima et al. 2024). A critical event in the transition from maternal to zygotic control is the transcriptional activation of the zygotic genome, termed zygotic genome activation (ZGA). While the mechanisms of ZGA vary across species, they often involve the opening of regulatory elements by pioneer factors during early cleavages (Zhou & Heald 2023). In C. elegans, ZGA begins in the three somatic blastomeres at the 4-cell stage. The fourth blastomere, germline precursor P2 remains quiescent. This is triggered by the nuclear entry of TAF-4, an essential component of the RNA polymerase machinery held in the cytoplasm by OMA-1/2 proteins until their degradation in late 2-cell embryos (Guven-Ozkan et al. 2008). However, the genomic regulation underlying ZGA is not known.

To explore early chromatin dynamics, we first compared patterns of accessibility in somatic cells in early embryonic stages through analysing the Fraction of Reads in Peaks (FRiP).

Although a similar number of ATAC fragments were recovered at each stage (Supplementary Figure 3), the fraction of reads in peaks (FRiP) varied by approximately two-fold (Figure 2A). FRiP was highest at the 2-cell stage, dropped sharply at the 4-cell stage (coinciding with ZGA), and then gradually increased as embryogenesis progressed. The differences in relative accessibility in peak regions indicates that chromatin organisation changes during early cleavages, with a major change at the 4-cell stage, when ZGA begins.

**Figure 2.**
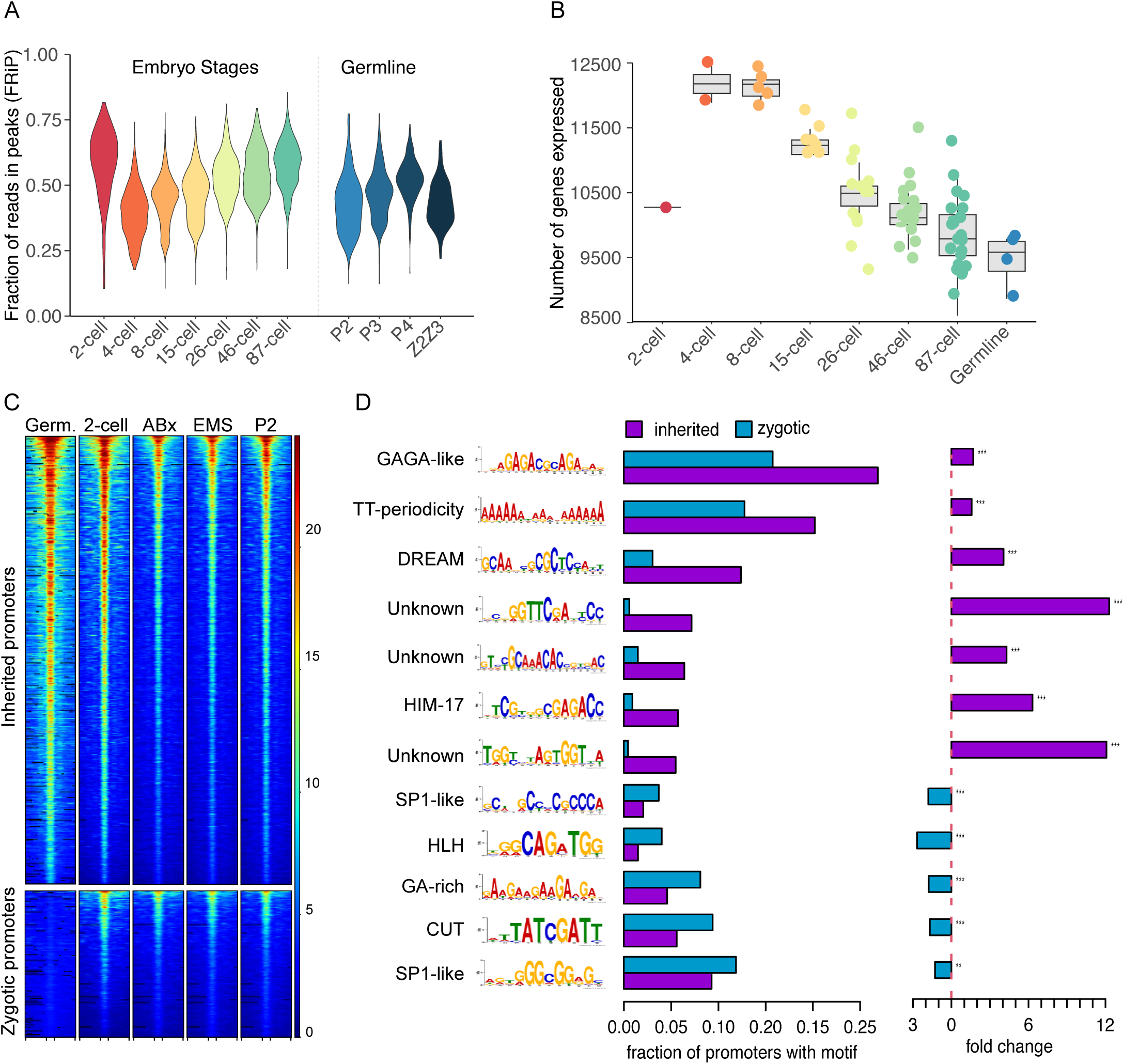
Chromatin accessibility landscape at ZGA. A) Fraction of reads in peaks (FRiP) measured from somatic cells of different embryo stages (left) and from germline precursors (right). B) Number of genes expressed in somatic cells from each embryo stage calculated from sampling 200,000 UMIs from each cell type. Each dot represents one cell type. C) Heatmap showing ATAC-seq signals on promoters accessible in the somatic cells at the 4-cell stage (+/- 500bp). Clustering was driven by germline signal. Germ. is adult germline ATAC-seq from (Serizay et al. 2020). D) Left: motifs enriched in inherited and/or zygotic promoters; predicted binding factors were defined using Tomtom. Center: fraction of inherited and zygotic promoters with one or more occurrences of the motif. Right: fold change of fraction of inherited/zygotic (purple) or zygotic/inherited (blue) promoters. Benjamini-Hochberg-adjusted P-values from chi-squared test: *** = P < 0.001, ** = P < 0.01.

We next investigated the relationship between the chromatin accessibility changes and gene expression. In line with ZGA, we detected an increase in the number of expressed genes at the 4-cell stage, after which the number gradually dropped (Figure 2B). We found that the high number of detected genes at the 4-cell stage was primarily due to expression of a larger number of lowly expressed genes (Supplementary Figure 6). The finding that 4-cell embryos undergoing ZGA have the lowest FRiP and the highest number expressed genes suggests that they may have a less organised and more permissive genome that temporarily permits widespread, potentially spurious gene expression. The change in chromatin accessibility patterns may be due to the sudden onset of transcription at the 4-cell stage, which is likely to disrupt nucleosome organisation. We note that H3.3 levels reduce during early cleavages (Gleason et al. 2023), which could potentially also play a role in these patterns.

To study ZGA in more detail, we identified the promoters accessible at the 4-cell stage and investigated the origin of their accessibility (Supplementary Table 5). Surprisingly, most of the 5,801 promoters we detected to be open in any cell at the 4-cell stage were accessible not only in the transcriptionally active somatic blastomeres but also in the transcriptionally quiescent germline precursor P2 (Figure 2C). Moreover, the accessibility patterns were largely shared with the quiescent 2-cell embryos, indicating that promoter accessibility is established prior to ZGA.

We hypothesized that this early accessibility may be maternally inherited. Indeed, 74% of the accessible sites at ZGA were also accessible in the adult maternal germline (Fig. 2C), consistent with maternal inheritance; mRNA from the associated genes was also detected as maternal RNA in zygotes (Supplementary Figure 7A). The remaining 26% of sites accessible at ZGA were closed in the maternal germline and newly opened in the embryo (Fig. 2C); consistent with zygotic specificity, maternal mRNA was rarely detected for these genes (Supplementary Figure 7A).

We observed that expression of genes with inherited promoters was largely uniform in our map (Supplementary Figure 7B) and indeed across stages as well as tissues in adults (Supplementary Figure 8). Consistent with ubiquitous expression, gene ontology analyses showed enrichment for genes with diverse housekeeping functions (Supplementary Figure 9). Expression of genes with promoters that open in the zygote gradually increases in early embryogenesis (Supplementary Figure 7B) and the majority have tissue restricted expression in adults (Supplementary Figure 8). This set is enriched for proteolysis functions, transcription factors, and other types of nucleic acid binding (Supplementary Figure 9).

We also observed different patterns of motif enrichments in the inherited and zygotic promoter sets, which provide leads for future investigations (Figure 2D). Notably, the inherited promoters were more highly enriched for two motifs previously shown to be associated with ubiquitous gene expression (GAGA and TT-periodicity (Serizay et al. 2020)) whereas the zygotic motifs had more frequent presence of HLH and CUT homeodomain transcription factor motifs.

To summarise, our findings indicate that ZGA occurs on a pre-patterned accessible genome. This regulatory architecture includes maternally inherited accessible elements and zygotically established sites. We propose that the release of TAF-4 from cytoplasmic sequestration enables transcription within this pre-existing chromatin landscape for the transition to zygotic genome control.

### A transient gene expression program regulated by CUT homeodomain transcription factors

We observed that pes-10, one of the first strictly zygotic genes identified (Seydoux & Fire 1994), displayed zygotically established promoter accessibility and contained a CUT motif in its promoter. Notably, pes-10 exhibited a striking expression pattern, characterized by transient high expression in pre-gastrulation cells (Fig. 3A). This distinctive pattern suggested the involvement of a specific regulatory program.

**Figure 3.**
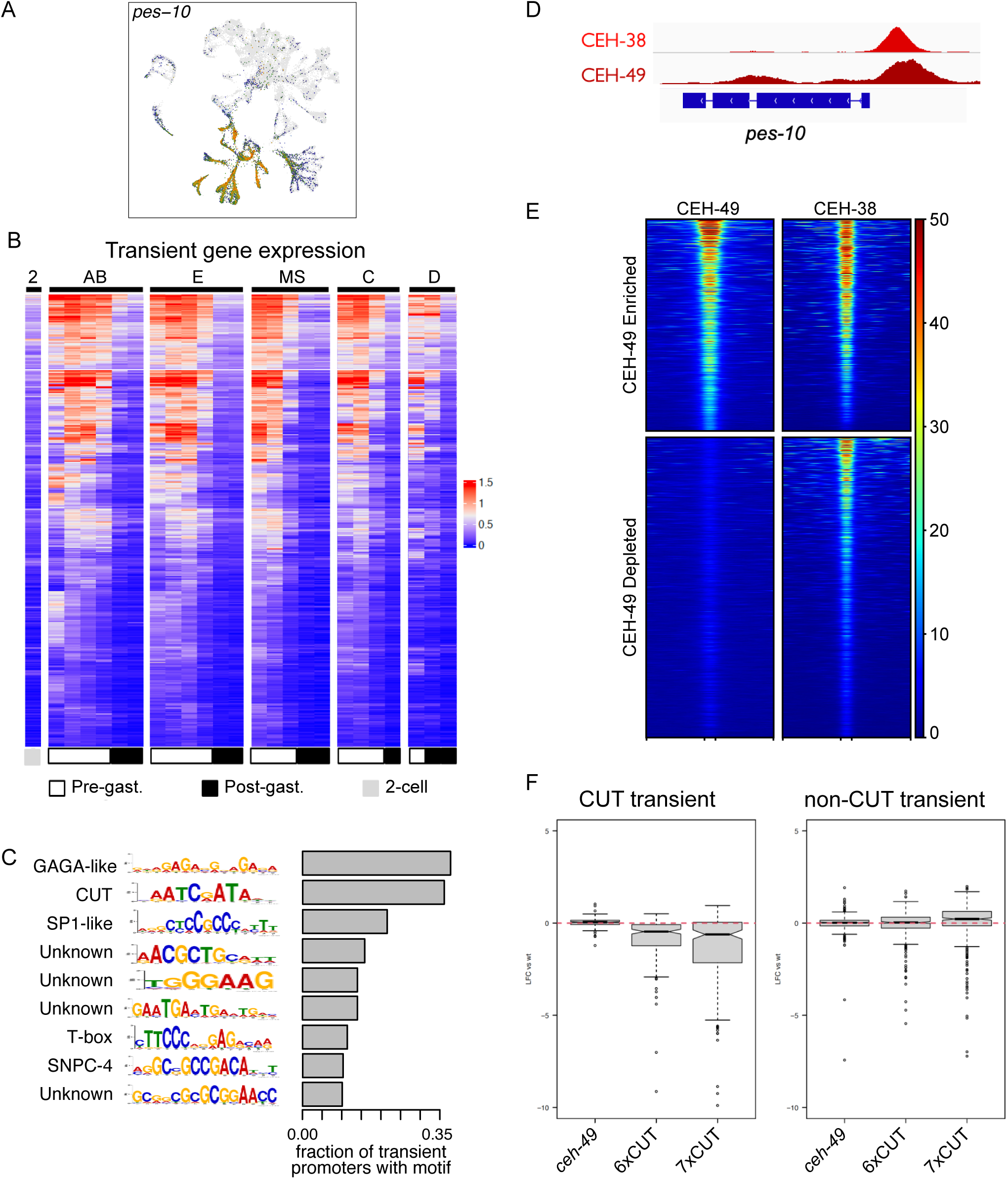
A transient gene expression program in pre-gastrulation embryos. A) Feature plot showing *pes-10* expression profile in pre-gastrulation cells across the full UMAP. B) Heatmap showing average expression of transient genes (log10 corrected counts + 1) in each somatic lineage. Cells from pre- vs post-gastrulation embryos are distinguished in white and black at the bottom of the heatmap. Columns are ordered temporally: AB = ABx, ABxx, ABxxx, ABxxxx, ABxxxxx, ABxxxxxx; E = EMS, E, Ex, Exx, Exxx; MS = MS, MSx, MSxx, MSxxx, MSxxxx; C: C, Cx, Cxx, Cxxx; D = D, Dx, DxxCxxx. C) Left: motifs enriched in promoters from transient genes; predicted binding factors were defined using Tomtom. Right: fraction of transient gene promoters with one or more occurrences of the motif. D) ChIP-seq binding profile of CEH-38 and CEH-49 at the *pes-10* locus. E) CEH-38 and CEH-49 ChIP seq signal across all promoters bound by CEH-38 and/or CEH-49. Clustering was driven by CEH-49 signal. F) The log2FC of expression in CUT mutant strains relative to wild-type animals in transient genes with or without a CUT protein at one or more of their promoters.

To investigate this further, we searched for other genes with similar transient expression patterns. We performed differential expression analysis comparing gene expression between pre-gastrulation and post-gastrulation cells (see Methods). This analysis identified 551 genes with a pattern of high expression across pre-gastrulation stage cells followed by a drop at gastrulation (Fig. 3B; Supplementary Table 6).

We next used MEME-chip to find motifs enriched in the promoters of the transient genes. The CUT motif was highly enriched, second only to a GA-biased motif, and was present in 38% of promoters (Figure 3C). The *C. elegans* genome encodes seven CUT homeodomain family genes: five are ubiquitously expressed, ceh-48 is neuron-specific, and ceh-49 is specific to the embryo and the late adult germline (Cao et al. 2017; Ghaddar et al. 2023). A recent study demonstrated that CUT family members act through the CUT motif to redundantly regulate pan-neuronal gene expression (Leyva-Díaz & Hobert 2022). By analyzing strains with increasing numbers of CUT gene knockouts, the study found progressively stronger gene expression defects, with the most severe defects observed when all six neuronally expressed CUT genes were mutated (“6X CUT”). The remaining unmutated embryo-specific gene, *ceh-49*, is highly expressed in early embryos and is itself a transient gene (Supplementary Figure 10). Except for *ceh-48*, the CUT genes are expressed in the pre-gastrulation embryos at varying levels (Supplementary Figure 10).

We used ChIP-seq to investigate CUT protein binding patterns in early embryos and ask whether they associate with promoters of transient genes, selecting CEH-38, which has broad expression, and CEH-49, which has transient expression in pre-gastrulation cells. We found that the CUT proteins were present at the *pes-10* promoter (Figure 3D). Genome-wide there was a high degree of overlap (Supplementary Table 7 and 8). We identified 4403 genes with a promoter bound by a CUT protein, with 1257 promoters showing high ChIP-seq signal for both CEH-38 and CEH-49, and 1783 promoters showing CEH-38 signal and low or no CEH-49 (Figure 3E)(Supplementary Table 9). Among the transient genes, 48% (N=265) had a promoter bound by at least one of the two CUT proteins, and the majority were bound by both (201 in the CEH-49-enriched vs 78 in the CEH-49-depleted set).

To assess potential roles of the CUT family in regulating a transient gene expression program, we created a knockout of the pre-gastrulation CUT gene *ceh-49* as well as knocking out *ceh-49* in the 6X CUT strain to make a 7X CUT strain. We then conducted gene expression profiling in early embryos from these three strains together with wild-type and assessed the effects (Supplementary Figure 11A-C, Supplementary Tables 10-13).

Hundreds of genes were differentially expressed in 6X CUT and 7X CUT mutants (n=366 and n=547, respectively), but essentially no differences were observed in the single *ceh-49* mutant. Consistent with CUT proteins promoting transcription, the majority of genes were downregulated (78% in 6X CUT and 67% in 7X CUT), with 235 shared between the two mutants. Of these, 63 have transient expression. As a group, the transient genes with CUT promoter binding were significantly downregulated in both the 6X CUT and the 7X CUT backgrounds (Figure 3F), supporting a role for the CUT proteins in promoting the transient program.

In line with the reported functional redundancy in the CUT family in the nervous system (Leyva-Díaz & Hobert 2022), we observed that further removal of *ceh-49* from the 6X CUT in the 7X CUT strain phenotype caused significant downregulation of 30 transient genes, most of which were downregulated in the 6X CUT strain (Supplementary Tables 11 and 13). The transient genes without CUT binding were not downregulated in the CUT mutants (Figure 3F), indicating these genes are under distinct regulatory control.

To further characterize this regulatory program, we annotated the CUT associated transient genes using WormCat. These genes were strongly enriched for proteolysis-related functions, including many components of the SCF ubiquitin ligase complex and a large number of F-box proteins (Supplementary Figure 12). The class is also enriched for genes involved in mRNA regulation and nucleic acid binding. Notably, this included six orthologs of the ZFP36L1/Tis11B family, which are RNA-destabilizing proteins (Baou et al. 2009).

These functions align with processes required during the maternal to zygotic transition, when the proteome is remodeled and maternal RNAs are regulated or degraded (Kojima et al. 2024). It has also been observed that SCF complexes are highly abundant and critical for maternal protein degradation during early embryogenesis in diverse species (Xie et al. 2019). We propose that the CUT-regulated transient program ensures sufficient levels of key regulators that facilitate this key transition, enabling the zygote to take control of development from the mother.

## Discussion

Our lineage-resolved multimodal single-cell atlas is a powerful resource for elucidating the regulatory events of early C. elegans development. The integration of chromatin accessibility and gene expression across the early lineage reveals the step-by-step changes in genome regulation as development unfolds.

Through analysing the map, we found that zygotic genome activation (ZGA) occurs on a pre-patterned chromatin landscape that is generated by inheritance of maternally accessible elements together with the pre-ZGA establishment of new sites in the zygote. This provides a template ready for initiation of transcription at the 4-cell stage when TAF-4 is able to enter the nucleus. Many of the maternally inherited accessible sites are promoters for ubiquitously expressed housekeeping genes, suggesting these promoters may always be open. The mechanism is unknown but could be via an active process, and/or by sequence features that destabilise nucleosomes. In contrast to inherited accessibility, the opening of promoter chromatin in the zygote is likely driven by maternally provided transcription factors, e.g., translation of maternal RNA for the SKN-1 transcription factor leads to its presence in EMS and P2 (Bowerman et al. 1993) where it likely opens the promoters of its targets such as MED-1/2. In support of this, the MED-2 promoter also opens in the P2 germline precursor (Supplementary Figure 13), though med-1 is not transcribed in P2 due to germline transcriptional silencing (Mello et al. 1996; Seydoux et al. 1996).

We also uncovered a transient gene expression program starting at ZGA driven by redundant CUT homeodomain transcription factors. We suggest that this program, enriched for genes involved in protein degradation and RNA regulation, contributes to the cellular remodelling required during the maternal to zygotic transition. It is striking that the CUT family is also redundantly required in C. elegans for the unrelated process of pan-neuronal gene expression (Leyva-Díaz & Hobert 2022). CUT homeodomain transcription factors are found across animals and act in diverse processes (Leyva-Díaz 2023), supporting the view that they do not have specialised developmental roles. In the C. elegans nervous system, the CUT proteins were shown to function together with terminal selector transcription factors (Leyva-Díaz & Hobert 2022), suggesting that this family may provide transcription in conjunction with cell-type specific regulators.

The mechanism of ZGA in C. elegans contrasts with mechanisms observed in other systems, such as zebrafish and Drosophila, where pioneer factors gradually remodel chromatin over the course of many pre-ZGA cleavages but maternal patterning of accessibility has not been reported (Kojima et al. 2024). Interestingly, a recent study in mouse provided evidence that some zygotic accessibility patterns are similar to those in early oocytes, suggesting inheritance of information, but the mechanism of this potential process is unclear (Liu et al. 2020; Jung et al. 2019).

Importantly, the lineage-resolved nature of our dataset circumvents challenges associated with reconstructing temporal dynamics computationally, providing fine temporal resolution of stepwise changes in gene expression and chromatin accessibility across cell divisions.

Extending this approach to later stages will enable detailed dissection of the regulatory processes in play across development, and advances in lineage reconstruction should make this possible in organisms without a defined lineage.

## Methods

### Strains

The following strains were used in this study: N2 Bristol as WT, JA1928 *ceh-49(we67)* V, OH16377 *ceh-38(tm321)* II; *ceh-44(ot1028)* III; *ceh-48(tm6112)* IV; *otIs356* V; *otDf1* X, JA1941 *ceh-38(tm321)* II; *ceh-44(ot1028)* III; *ceh-48(tm6112)* IV; *ceh-49(we67)* V; *otIs356* V; *otDf1* X, OH16224 *ceh-49(ot1016[ceh-49::gfp])* V, PHX4799 *ceh-38(syb4799[ceh-38::GFP])* II, OH17051 *ceh-48(ot1125[ceh-48::GFP])* IV.

### *C. elegans* maintenance and cultures

C. elegans strains were cultured using standard methods at 20℃ on Nematode Growth Medium (NGM) plates with OP50 (Brenner 1974). To obtain large quantities of early embryos, liquid cultures were grown. Briefly, gravid animals were washed from NGM plates using M9 buffer (3 g KH2PO4, 6 g Na2HPO4, 5 g NaCl, 1 ml 1 M MgSO4, H2O to 1 litre) and treated with hypochlorite solution (1.2% hypochlorite, 0.5M NaOH) for approximately 5 min. Embryos were quickly washed three times using M9 buffer and hatched at 18℃ in M9 buffer without food to obtain synchronized L1 larvae. L1s were grown at 18℃ to gravid adult worms in S-Basal (0.1 M NaCl and 0.05 M potassium phosphate (pH 6.0), 5 µg/ml cholesterol) with *E. coli* HB101 at a concentration of 1000-3000 worms/mL. An additional round of culturing was performed to obtain larger populations if needed.

### CRISPR/Cas9 genome editing

We used the CRISPR/Cas9 method as described by Ghanta *et al*. (Ghanta et al. 2021). Microinjections were performed as described by Mello & Fire (Mello & Fire 1995). To generate JA1928 *ceh-49(we67)* V, we injected guide g19 (AUAUUCUUUGAAAAUCUGAA) and g20 (UUUGGAUGUAAAUAUGUCAA), and repair template O099 (CAAAACTTCAAAATTATATATTAATTAAACCCTTCCAATGGAAAACATTGAAAAGAAATGTTTAAAGATT) in N2. To generate JA1941, in which the entire CUT family is mutated, we injected guides g19, g20, and repair template O099 in OH16377.

### Single cell multiomics

To collect early embryos, a culture containing ∼1 million animals was treated with hypochlorite solution (1.2% hypochlorite, 0.5M NaOH) when approximately ⅓ of the adults in a culture, grown for ∼62 hours at 18℃ with E. coli HB101 as a food source, had one to 6 embryos. Embryos were washed 2x with M9 and 1x with Egg salts (118 mM NaCI, 40 mM KCI, 3.4 mM CaCI2, 3.4 mM MgCl2, 5 mM Hepes, pH 7.2). To remove the egg shell, embryos were treated with chitinase (Sigma-Aldrich C6137) in Egg salts (0.2 U/ml) for 4 to 4.5 min. Embryo morphology was monitored under a dissecting microscope until they appeared more rounded, indicating egg shell degradation. Chitinased embryos were quickly washed 3x in ice cold Egg salts and were kept on ice from now on. Digested but intact embryos were washed 1x in Nuclei Extraction Buffer (NEB) (Quarato et al. 2021) with 2% BSA and resuspended in NEB+ (NEB with 2% BSA, 1U/ml proteinase inhibitor Roche #3335402001, and 1U/ml Protector RNAse inhibitor Roche #3335402001). Nuclei were extracted by douncing ∼800,000 to 1 million digested embryos in a steel tissue grinder (Wheaton Dounce DURA-GRIND 357572). Debris was pelleted at 150xg in a benchtop centrifuge (Eppendorf 5804) at 4℃. Supernatant was transferred to a new tube and nuclei were pelleted at 700xg and washed 2x in NEB+. Finally, nuclei were resuspended at 10,000 nuclei/ml in diluted nuclei buffer (10x Genomics) before performing ATACseq and RNAseq multiomics using the 10x Genomics Single Cell Multiome kit (10xGenomics, Chromium Single Cell Multiome ATAC + Gene Expression) according to the manufacturer’s instructions. We performed seven profiling experiments (named exp024, exp025, exp027, exp031, exp032, exp042, and exp043; ;exp042 and exp043 are technical replicates)

### HCR RNA FISH

HCR RNA FISH v3 (Choi et al. 2018) probes and amplifiers were ordered from Molecular Instruments. Embryos were collected by hypochlorite treatment and washed in PBS, resuspended in 4% paraformaldehyde in PBS, and flash frozen in liquid nitrogen, and stored at -80C before use. Synchronized young adult worms from a liquid culture were washed 2x in PBS and fixed as described above. Before in situ hybridization, samples were fixed by thawing for 30 min in room temperature water. Samples were washed 2x in PBS and permeabilized in 70% ethanol at 4°C for 24 hrs to one week. Samples were washed 2x in PBST (0.1% Tween 20 in PBS) and in situ hybridization was performed according to the manufacturer’s instructions. Briefly, samples were incubated in 2 mg/ml glycine in PBST for 15 min on ice. Samples were then washed 2x in PBS-T and incubated for 5 min in a 1:1 PBST to probe hybridization buffer at room temperature. Samples were pre-hybridized at 37°C in 300 μl pre-warmed probe hybridization buffer for one hour. A probe solution was prepared with 2 pmol per probe for qHCR or 8 pmol per probe for dHCR in 200 μl pre-warmed probe hybridization buffer. The probe solution was added to the sample and incubated at 37℃ overnight. Excess probes were removed by washing 4x for 15 min in probe wash buffer at 37℃. Samples were washed 2x in SSCT (0.1% Tween20, 150 mM NaCl, 15 mM Na3C6H5O7, pH 7) and pre-amplified in 300 μl amplification buffer for 30 min at room temperature. 30 pmol of hairpin h1 and hairpin h2 were snap cooled separately by melting at 95°C for 90 seconds in a thermocycler and cooled to room temperature in the dark for 30 min. A hairpin solution was prepared by adding 30 pmol of each hairpin to 200 μl of amplification buffer before adding it to the sample. Samples were incubated in the dark at room temperature for 3-16 hrs for qHCR or 45 min for dHCR. Excess hairpins were removed by washing with 5x SSCT at room temperature for 2x 5 min, 2x 30 min, 1x 5 min. Samples were mounted in Vectashield Plus (Vector Laboratories, H-1900), stored at 4 °C, and imaged within two weeks.

### Micrscopcopy and image analysis

HCR RNA FISH slides were imaged using a Nikon AX R confocal on a Nikon Ti2-E inverted microscope equipped with a Nikon resonant scanner and NIS Elements NIS AI acquisition software, version 5.42.06. A Nikon Plan Apo 60x/1.4 NA objective was used for qHCR and a Nikon PLan Apo 100x /1.45 NA objective was used for dHCR. Laser intensities were controlled across samples. Images were analyzed using FIJI (version 2.14.0) (Schindelin et al. 2012). qHCR expression levels were analyzed by hand and dHCR nuclear dots were quantified using a custom script.

### RNA-seq

RNA-seq experiments (two biological replicates) on early embryos of N2, ceh-49(we67) (JA1928), 6xCUT (OH16377) and 7xCUT(JA1941) were performed as described in (Carelli et al. 2022). .

### ChIP-seq

Chip-seq experiments (two biological replicates) on early embryos of CEH-49::GFP (OH16224) and CEH-38::GFP (PHX4799) were performed as described by (Carelli et al. 2022).

### Computational methods Genome annotation

Genome, gene and protein annotations were downloaded from Wormbase (Sternberg et al. 2024) (release WS285). The C. elegans repetitive elements annotation for the genome release ce10 (corresponding to Wormbase WS220) was downloaded from Dfam (Hubley et al. 2016)(release 3.5) and converted to ce11 (corresponding to WS285) using LiftOver (Hinrichs et al. 2006).

### Processing of sequencing data

#### Demultiplexing

Single cell multiomics data were demultiplexed using the mkfastq function from cellranger- arc (version 2.0.1) to generate FASTQ files.

#### scATAC-seq reads mapping and initial peak calling

scATAC-seq reads were mapped on the C. elegans genome using the cellranger-arc count function. The fragment files (“atac_fragments.tsv.gz”) from all samples were then used in bulk as input to call an initial set of ATAC-seq peaks. Each fragment longer than 150bp was split in two and both ends were each extended to 150bp around the fragment’s ends (Tn5 cut sites); fragments shorter than 150bp were used without splitting them. All sequences were then used to call peaks with MACS2 (Zhang et al. 2008)(version 2.2.7.1) with the following settings: --call-summits --extsize 150 --shift 0 --gsize ce --keep-dup all. All peaks were then re-sized to 200bp sound their MACS2 summits using bedtools (Quinlan & Hall 2010)(version 2.31.0) to obtain an all-samples bulk peakset.

#### scRNA-seq reads mapping and transcriptome extension

scRNA-seq reads were initially mapped on the C. elegans genome with STARsolo (Kaminow et al. 2021)(version 2.7.10a) using the WS285 gene annotation as a guide. We then used a similar approach as in (Packer et al. 2019) to extend the 3’ gene annotation (Supplementary Table 16). Briefly, RNA-seq reads from all samples were deduplicated using umi_tools dedup (Smith et al. 2017)(version 1.0.0). We then calculated expression levels from each transcript from an annotated protein-coding gene, lincRNA, or pseudogene after extending their 3’ end by up to 500bp in 100bp steps. Any 100bp extension was cut short if it: a) overlapped another transcript; b) overlapped an accessible site (all samples bulk peakset, see “scATAC-seq reads mapping and initial peak calling” section); or a blacklisted region. We then allowed a transcript extension if: a) the original transcript had a coverage > 100 reads; b) if the tentative extension exceed by at least 10% the coverage of the original transcript; c) if the counts per extension bin do not exceed the count in the previous bin by more than 10%. If the three requirements were met, then we kept the shortest extension retaining more than 90% of the additional reads.

After the transcriptome extension step, we re-run STARsolo for each sample using the extended transcriptome as a guide and with the following settings: --soloType CB_UMI_Simple --soloCBwhitelist whitelist/737K-arc-v1.txt --soloUMIlen 12 --soloCellFilter None --soloFeatures GeneFull --soloMultiMappers EM --alignIntronMax 20000.

### Single cell multiomics data analysis

#### Identification of empty droplets

Empty droplets in each sample were identified based on their RNA-seq profile using the EmptyDrops (Lun et al. 2019) R function from the dropletutils package (version 1.14.2). Per-cell UMI counts were manually investigated to remove low count barcodes - likely caused by barcode hopping - and to set the lower bound of the UMI counts likely to represent empty droplets. An FDR < 0.001 was used to distinguish putative empty droples from real cells (cell barcodes).

#### Initial cell clustering - pre-filtering and doublets removal

Cell barcodes from each sample were initially pre-processed to remove: a) all barcodes with >5% of all UMIs derived from genes encoded by the mitochondrial genome; b) all barcodes with > 10’000 UMIs; c) all barcodes with a scATAC-seq fragment count (measured on accessible sites detected by cellranger-arc) lower than 200 or higher than 5 MADs (median absolute deviations). The gene expression matrix of the remaining barcodes was then filtered to retain only protein-coding genes, lincRNAs, or pseudogenes and used to generate an initial barcode clustering using the Seurat package (Hao et al. 2021)(version 4.3.0). Data were normalised using SCTransform v.2 (Choudhary & Satija 2022), the dimensionality of the normalised count matrix reduced used PCA, and barcodes clustered based on the first 50 principal components following the standard Seurat pipeline. Based on this initial clustering, we then identified and removed doublets using the scDblFinder (Germain et al. 2021)(version 1.8.0) R package. After doublets removal, the RNA count matrix was again normalised using SCTransform and cells clustered as previously described.

#### Ambient RNA correction

We used SoupX (Young & Behjati 2020)(version 1.6.2) to minimize the contribution of cell-free RNA (ambient RNA) to the RNA profile of each barcode. Since marker genes automatically detected by SoupX were heavily enriched for genes with high RNA counts in P-cells, we decided to exclude potential P-cell clusters from the contamination estimate step, and then run SoupX (with settings: soupQuantile = 0.80) on the remaining clusters for each sample. The contamination estimate was then used to adjust the full RNA count matrix (including putative P-cell clusters).

#### Multiomics samples merging

The ambient RNA-corrected count matrix and the ATAC-seq count matrix - quantified over the scATAC-seq all samples bulk peakset (see “scATAC-seq reads mapping and initial peak calling” section) - of each sample were again normalised using SCTransform v.2 and the TF-IDF method implemented in Signac (Stuart et al. 2021)(version 1.6.0), respectively. The resulting normalised gene count and ATAC-seq fragment count matrices from each sample were then merged into a single Seurat object using the Seurat merge function, and the whole set of barcodes then processed as previously described. The relatively homogeneous contribution of barcodes from each sample to each cell cluster suggested minimal batch effects among all samples. The major differences in barcode-to-cluster contribution could be largely explained by differences in staging between samples, with exp024, exp025 and exp027 contributing in proportion more post-gastrulation cells than the other samples.

#### Calling ATAC-seq peaks for each cell cluster

scATAC-seq fragments from each cell cluster were obtained using the SplitFragments function from Signac. All ATAC-seq fragments were split and re-centred around the cut sites (as described in the “scATAC-seq reads mapping and initial peak calling” subsection), and peaks called for each cluster using MACS2 with settings “--llocal 2000 --call-summits --bdg -- SPMR --extsize 150 --shift 0 --gsize ce --keep-dup all --nomodel” and then re-scaled to 100bp around the summit. Peaks from each clusters called at a MACS2 q-value < 0.00001 were then removed if they overlapped for >50 of their length an exon (except for those overlapping a 5’UTR or the first 100bp of a noncoding transcript, in which case any overlap was allowed). All peaks from each cluster were then merged into a unique set of peaks and extended to 200bp around the midpoint to obtain an initial cluster-based peakset (round-1 peakset).

#### Multimodal clustering of 10x data and final dataset cleanup

The gene expression matrix and the ATAC-seq fragment matrix computed on the round-1 peakset (and normalised using the TF-IDF method) for all cells were used to run a multimodal cell clustering using the Weighted Nearest Neighbor (Hao et al. 2021) workflow implemented in Seurat. We used the FindMultiModalNeighbors function with the first 50 dimensions for both the RNA and the ATAC modality (but excluding the first LSI component for the ATAC-seq modality).

We noticed that some cell clusters showed a heavy skew in the contribution of cells from the 7 multiomics samples. Since we did not expect the cellular composition to be markedly different among our samples, we flagged clusters made up of:

- at least 4-fold more cells from samples exp031 and exp032 combined, than from exp042 and exp043 combined (and vice-versa);

- at least 80% of cells from clusters exp024, exp025 and exp027 combined.

The 10730 barcodes from these 16 flagged clusters showed overall higher levels of cell-free (ambient) RNA, RNA from mitochondrially-encoded genes and ribosomal RNA, and low levels of lincRNA expression and ATAC-seq fragments per cell. These characteristics, coupled with the strong bias in sample of origin, suggested that those barcodes might represent damaged cells, and we thus decided to remove them from our dataset, leaving us with a final number of 48775 cells. The final cell set was again processed to normalise the RNA-seq and ATAC-seq count matrices and recluster cells using the WNN method.

#### Cell type annotation

We used a set of marker genes derived from literature searches and in-house RNA-FISH experiments to manually assign cell clusters to specific cell types and cell lineages, i.e. cells derived from the same founder cell after the same number of cell divisions (for example, the ABxxx cell lineage includes the 8 cell types obtained after 3 rounds of cell divisions of the AB precursor). Ambiguities in the precise assignment of a cell cluster to a given cell type were indicated with an “x”. For example:

- the MSxa cluster includes both the MSaa and the MSpa cells;

- the ABxxx cell cluster contains cells from multiple cell types from the ABxxx lineage.

No cell type assignment was attempted for clusters ABxxxxx, ABxxxxxx, MSxxx, MSxxxx, Cxxx, and Dxx since few specific cells would be present in our dataset, hampering a confident assignment.

*Two-cell embryo*: one cell cluster (“twocell”) displayed an enrichment of RNA signal from *cpg-3* and *mesp-1*, two genes with high expression in the oocytes based on data from Ghaddar *et al*. (Ghaddar et al. 2023). RNA-FISH confirmed that *cpg-3* and *mesp-1* levels are highest in 2-cell embryos.

*Germline precursors*: four clusters showed very high levels of RNA derived from P-granule-associated genes (Lee et al. 2020), suggesting they represented germline precursors. The three largest clusters had variable enrichment of different P-granule genes RNA, and displayed a gradual increase in the ATAC-seq signal in a set of germline-specific promoters described in (Carelli et al. 2022), and were thus annotated as P2, P3 and P4. Further supporting this annotation, we anecdotally observed higher ATAC-seq levels in promoters of genes specifically expressed in their somatic sister cells (e.g. high accessibility on *med-1* and *med-2* promoters in EMS and P2, and on the C44B9.3 promoter in D and P4).

The smallest cluster showed a lower enrichment of P-granule genes RNA, and the specific expression of *prg-2* and other genes enriched in the Z2 and Z3 cells in the scRNA-seq dataset from (Packer et al. 2019), and was thus annotated as “Z2Z3”.

*Unassigned cells*: 5783 cells from 9 clusters could not be confidently assigned to any cell type or cell lineage.

#### Final ATAC-seq peakset

We extracted ATAC-seq fragments from each cell type and defined a new set of peaks per cell type following a similar approach as described in section “Calling ATAC-seq peaks for each cell cluster”. Genome-wide, CPM-normalised ATAC-seq profiles for each cell type were generated using the bamCoverage function from deepTools2 (Ramírez et al. 2016)(version 3.5.2). After merging ATAC-seq peaks of each cell type into a global set of accessible sites (round-2 pre-IDR peakset), we further refined the set of regions accessible in each cell type using an IDR-based approach (Li et al. 2011). Specifically, we aggregated ATAC-seq fragment counts measured on the round-2 pre-IDR peakset from the following combinations of samples:

- Group 1: exp024, exp031 and exp042;

- Group 2: exp025, exp027, exp032 and exp043.

The two groups of samples were chosen to balance the total number of cells for each cluster. The aggregate counts of each accessible region from the two groups were used as input to run IDR (version 2.0.4.2) on each cell type. Peaks passing an IDR threshold of 0.05 were kept to define sets of accessible regions in each cell type, for a combined set of 30634 ATAC-seq peaks called in any cell of our dataset (final peakset, Supplementary Table 14). These regions were used to generate the final ATAC-seq count matrix of our multiomics dataset.

#### Early cell cycle detection

A few clusters were automatically assigned to either individual cell types (e.g. EMS, C…) or to specific lineages (e.g. ABxxx, MSx…), yet showed the following common features:

- High expression of most histone genes;

- Low ATAC-seq FRiP;

- Low expression of marker genes of their mother cell.

These features, combined with experimental evidence from RNA-FISH, suggested that these clusters represented cells at an earlier stage of their cell cycle, and were thus annotated with the “early” suffix.

To distinguish early- from late-cell cycle cells in our datasets, we used the expression profile of a subset of early (“earlyABxxx”, “earlyEMS”, “earlyCx”, “earlyMSxx”) and late (“ABala”, “ABalp”, “ABpxa”, “EMS”, “Ca”, “Cp”, “MSxa”, “MSxp”) cell types to train a random forest model using the caret R package (Kuhn 2007)(version 6.0_91) with settings “tuneGrid = data.frame(mtry = 50), ntree=100”. Accuracy measured on early (“earlyABxxxx”, “earlyEandMS”, “earlyCxx”, “earlyMSx”) and late (“ABaraa”, “ABarpa”, “ABpxap”, “E”, “MS”, “Cxa”, “Cxp”, “MSa”, “MSp”) cell cycle cells was ∼0.87%, and allowed us to confidently annotate 17520 early- and 31255 late-cell cycle cells.

scRNA-seq coverage tracks

STARsolo alignment outputs from each sample were combined into distinct sets of cell-type-specific BAM files based on the barcodes assigned to each cell type using the filterbarcodes function from sinto (https://timoast.github.io/sinto/index.html, version 0.8.1). Each BAM file was then de-duplicated using the umi_tools dedup function (Smith et al. 2017), and strand specific coverage tracks generated using bamCoverage from deepTools2.

#### Bulk RNA-seq analysis

Paired-end reads from bulk RNA-seq experiments (whole embryo RNA or nuclear RNA) were trimmed and pre-processed using trim_galore (https://github.com/FelixKrueger/TrimGalore, version 0.6.7), and then aligned on the WS285 genome using STAR (Dobin et al. 2013) with settings “--outWigType wiggle --twopassMode None --alignIntronMax 20000” to generate bigWig tracks. Gene expression was quantified on the extended WS285 transcriptome using Kallisto (Bray et al. 2016)(version 0.48.0).

For nuclear bulk RNA-seq data, the STAR alignment was further filtered to retain only uniquely mapped read pairs using SAMtools (Li et al. 2009)(version 1.18) which were then converted into a BED file of RNA fragments (connecting the read1 and read2 ends) using BEDTools.

#### Bulk ATAC-seq and ChIP-seq analysis

Paired-end reads from bulk ATAC-seq (from (Serizay et al. 2020)) and ChIP-seq experiments (this work) were trimmed and pre-processed using trim_galore, and then aligned on the WS285 genome using bwa mem (Li 2013), retaining only reads with MAPQ > 10. CPM-normalised coverage tracks were generated with bamCoverage. ChIP-seq peaks were called using the YAPC method (Jänes et al. 2018) with an IDR < 0.001.

#### Promoter annotation

Promoters were annotated using a similar approach as described in (Jänes et al. 2018). Briefly, accessible sites (“final peakset”) were annotated as promoters if:

- Strand-specific, nuclear RNA-seq coverage was significantly downstream of an accessible site. The analysis was performed using two replicates of early-embryo (this paper) or mixed-stage embryo (Jänes et al. 2018) nuclear RNA-seq data. The increase in signal was measured in the 200bp downstream of each accessible site (oriented based on the RNA-seq strandedness) and compared either the 200bp upstream, or to the signal in the accessible region. Significant differences (LFC > 1.5; adjusted P-value < 0.1 when comparing the 200bp downstream vs the 200bp upstream regions; adjusted P-value < 0.05 when comparing the 200bp downstream vs the accessible site OR if the accessible site was located within a protein coding gene, lincRNA, or pseudogene) were determined using DESeq2 (Love et al. 2014)(version 1.34.0).
- The summit of the accessible site was located upstream of the coding sequence start site of a transcript, or at most 250bp downstream of the transcription start site of a noncoding transcript. Accessible site were not considered if the 5’ end of a non-first exon of another gene (in the same orientation) was located in the intervening region.
- The intervening distance between the accessible site and the downstream gene was shorter than 200bp OR if gaps longer than 50bp in the nuclear RNA-seq coverage (measured combining the two early embryo or mixed embryo replicates) were located in a longer intervening region.

Moreover, when two adjacent (closer tha 300bp) accessible sites met all the aforementioned criteria, and one site had an ATAC-seq coverage signal (combined fragment counts from all cells from the multiomics dataset, measured using BEDtools) lower than 90% of the adjacent peak, only the site with the highest ATAC-seq signal was retained as a promoter. In total, this approach allowed to annotate 6888 accessible sites as promoters.

We further complemented our promoter annotation by transferring the promoter-to-gene annotation for all our accessible sites overlapping a promoter annotated in (Serizay et al. 2020). With this approach, 3963 additional promoters were added to our set.

Finally, a promoter regulating the first or an internal gene in an operon (WS285 annotation) was automatically assigned to all downstream genes. In total, our promoter set thus included 10851 promoters regulating 11083 genes (Supplementary Table 15).

### Identification and analysis of inherited and zygotic promoters

We selected the 5861 promoters accessible in the somatic cells of 4-cell stage embryos (earlyEMS, EMS and ABx cell types), and measured and visualised the ATAC-seq signal from adult germline cells (Serizay et al. 2020), and from cells from 2- and 4-cell stage embryos using the deepTools2 suite (Ramírez et al. 2016). We used k-means to cluster the ATAC-seq signal from germline cells, thus distinguishing a set of promoters with levels of ATAC-seq coverage comparable to the background (zygotic promoters) from other accessible sites (inherited promoters). Maternal RNA levels of genes regulated by inherited or zygotic sites (maternal and zygotic genes) were obtained from scRNA-seq data from (Cole et al. 2024). Gene ontology enrichment of maternal and zygotic genes was performed using WormCat (Holdorf et al. 2020)(version 2.0).

Motifs enriched in inherited and zygotic promoters were detected using MEME-ChIP (Machanick & Bailey 2011)(version 5.5.2) using settings “-meme-minsites 10 -streme-pvt 0.05 -maxw 15 -minw 6 -filter-thresh 0.05 -meme-mod zoops”. MEME and STREME significant hits from both sets of promoters were mapped on all promoters using FIMO (Grant et al. 2011), keeping hits with a P-value < 0.00001. We combined hits from similar motifs with the following approach. First, all motifs from both promoter sets were aligned against each other using Tomtom (Gupta et al. 2007). Motif pairs with a Tomtom E-value < 0.001 were then clustered using the NetworkX python package (Hagberg et al. 2008). We then quantified the presence of a given motif cluster in any inherited or zygotic promoter using BEDtools.

### Transient genes

We identified transient genes - i.e. genes expressed in every pre-gastrulation cell type based on their differential expression in pre- and post-gastrulation cell types. Specifically, we compared their expression by subsampling 200 cells from the following cell lineages:

- Pre-gastrulation lineages: ABx, ABxx, ABxxx, EMS, E, MS, MSx, C, Cx
- Post-gastrulation lineages: ABxxxxx, ABxxxxxx, Exx, Exxx, MSxxx, MSxxxx, Cxxx, Dx, DxxCxxx

The subsampling was performed to avoid over-representation of data from lineages with more cells. Cell lineages present around the gastrulation onset were excluded to more clearly detect differences between the two sets. Aggregate counts from the subsampled sets of each lineage were then used as input for DEseq2, in which each lineage was used as an independent sample. Genes were considered transient if they showed a significantly higher expression in the pre-gastrulation set (adjusted P-value < 0.001, LFC > 2, lfcSE < 0.7).

Requiring a low lfcSE allowed to include genes upregulated across the pre-gastrulation cells.

We used MEME-ChIP to identify motifs enriched in the promoters of transient genes as well as their putative regulators using the motif-to-transcription factor assignment from CisBP2.0 (Weirauch et al. 2014).

### CUT proteins regulatory network

Binding sites of CEH-38 and CEH-49 overlapping promoters were merged and then distinguished in two sets (CEH-49-enriched and CEH-49-depleted) based on k-means clustering of their signal with DeepTools2.

We determined sets of genes differentially expressed between different CUT protein mutant strains using DESeq2 (|LFC| > 1 and adjusted P-value < 0.001). VolcanoPlots were generated with the EnhancedVolcano R package (version 1.12.0). We defined direct targets as genes downregulated in a mutant and bound by a CUT protein on their promoter(s).

### Data availability

An R object (C_elegans_wt_early_embryo_GEX_ATAC_multiome.rds) containing the processed single-nucleus 10X Genomics multiome dataset is available from at DOI 10.5281/zenodo.14261731.

## Supporting information

Code and external data

## Acknowledgements

This work was supported by a Wellcome Investigator award (217170) to JA. Some strains were provided by the CGC, which is funded by NIH Office of Research Infrastructure Programs (P40 OD010440). We would like to acknowledge staff at the Gurdon Institute Imaging Facility for microscopy support.

---

Supplementary Figure 1

UMAP visualisation and annotations of all cells that passed quality controls.

Supplementary Figure 2

HCR RNA-FISH validations of marker genes.

For each gene assayed, the feature plot shows the gene expression signal on the UMAP and the chart displays the detection of the RNA in the nucleus per cell type.

Supplementary Figure 3

Cell cluster metrics. Violin plots showing distributions of the number of RNA UMIs, the number of detected genes, and the number of ATAC fragments per cell cluster.

Supplementary Figure 4

Characterisation of putative early cell cycle clusters.

(A) Heatmap of expression of histone genes in putative early cell clusters (left) and putative late cell clusters (right).

(B) *pes-10* levels, quantified by RNA-FISH, decrease across the cell cycles of ABaraa and ABarap, timed using the number of cells in the embryo. RNA-FISH for the D cell used as an internal control.

Supplementary Figure 5

Differences in chromatin accessibility and gene expression across individual cell cycles

Supplementary Figure 6

Number of genes expressed per embryo stage separated by gene expression level

Supplementary Figure 7

Expression of genes with inherited vs zygotic promoters across early embryo stages.

Supplementary Figure 8

Stage and tissue specificity of expression of genes with inherited or zygotic promoters. Left heatmaps plot gene expression across developmental stages (Janes paper), right heatmaps plot expression in adult tissues (Serizay et al. 2020), and the gene expression class plots gene expression class in adult tissues (Serizay et al. 2020). (A) Most genes with inherited promoters are broadly expressed across stages and have ubiquitous expression across tissues ; Bottom, genes with zygotic promoters show less uniform expression across stages and tissues, and most show tissue restricted expression in adults.

Supplementary Figure 9

Functional classes of inherited and zygotic genes.

Supplementary Figure 10

Feature plots showing expression of CUT family members.

Supplementary Figure 11

Violin plots showing gene expression changes in CUT mutants

Supplementary Figure 12

Functional classes of transient genes with and without CUT bound promoters

Supplementary Figure 13

*med-1* promoter accessibility in EMS and P2

