## Supplementary material for "A lineage-resolved multimodal single-cell atlas reveals the genomic dynamics of early *C. elegans* development": Code and external data: wormcat_function.R

```
library("stats")
library(ggplot2)
library(ggthemes)
library("plyr")
library("stats")


# Worm Cat Function

worm_cat_fun <- function(file_to_process, title = "rgs", output_dir = NULL, rm_dir = FALSE, annotation_file = "whole_genome_v2_nov-11-2021.csv", input_type = "Sequence.ID", zip_files = FALSE) {
    # Check the input file for validity

    # Check special case for help documentation
    if (substr(file_to_process, 1, 7) == "WORMCAT") {
        wormcat_package_path <- system.file(package = "wormcat")
        file_to_process <- gsub("WORMCAT", wormcat_package_path, file_to_process)
        print(sprintf("WORMCAT substituted for %s", file_to_process))
    }

    if (!file.exists(file_to_process)) {
        print(sprintf("The file %s cannot be found.", file_to_process))
        print("EXITING!")
        return()
    }

    # If output_dir is not given, create one using a timestamp
    if (is.null(output_dir)) {
        output_dir <- paste("worm-cat_", format(Sys.time(), "%b-%d-%Y-%H:%M:%S"), sep = "")
    }

    # Check if the output_dir directory exists
    if (dir.exists(output_dir)) {
        # Check if the directory is empty
        if (length(list.files(output_dir)) == 0) {
            message(sprintf("The directory %s exists and is Empty.", output_dir))
        } else {
            print(sprintf("The directory %s is not Empty.", output_dir))
            print("EXITING!")
            return()
        }
    } else {
        # Create the output directory
        dir.create(output_dir, recursive = TRUE)
    }


    # If annotation_file starts with HTTP assume we have a URL
    if (grepl("^http:|^https:", tolower(annotation_file))) {
      worm_cat_annotations <- annotation_file
    } else {
        separator <- .get_system_path_separator()
        # If annotation_file contains a file system specific separator assume an external annotation file is being used
        if (grepl(separator, annotation_file)) {
            worm_cat_annotations <- annotation_file
        } else {
            # Get full path to annotations file
            worm_cat_annotations <- system.file("extdata", annotation_file, package = "wormcat")
        }

        if (!file.exists(worm_cat_annotations)) {
          print(sprintf("The annotation file %s cannot be found.", worm_cat_annotations))
          print("EXITING!")
          return()
        }

    }


    # Create the categories file and save it to CSV output
    .worm_cat_add_categories(file_to_process, output_dir, worm_cat_annotations, input_type)
    # Run a fisher test
    .worm_cat_fisher_test(output_dir, worm_cat_annotations)

    # For each of the three files created above:
    # 1. Parse the file only to include the entries with "acceptable p-values."
    # 2. Create bubble plots for each of the three categories based on the acceptable p-vlaues.
    for (i in 1:3) {
        rgs_fisher_cat_csv <- .create_file_nm(output_dir, "rgs_fisher_cat%d.csv", i)
        .worm_cat_acceptable_pvalues(rgs_fisher_cat_csv)

        rgs_fisher_cat_apv_csv <- .create_file_nm(output_dir, "rgs_fisher_cat%d_apv.csv", i)

        plot_title <- paste(title, sprintf("category%d", i), sep = ":")
        .worm_cat_bubble_plot(rgs_fisher_cat_apv_csv, plot_title)
    }

    # Capture basic information about this run write output to run_data.txt
    run_data <- .create_file_nm(output_dir, "run_data.txt")
    runtime_l <- paste("runtime", format(Sys.time(), "%b-%d-%Y-%H:%M:%S"), sep = ": ")
    annotation_file_l <- paste("annotation_version", annotation_file, sep = ": ")
    input_type_l <- paste("input_type", input_type, sep = ": ")

    cat(runtime_l, annotation_file_l, input_type_l, file = run_data, sep = "\n", append = TRUE)

    # Create a zip file as the final output
    zip_ext <- ""
    if (zip_files) {
       files2zip <- dir(output_dir, full.names = TRUE)
       utils::zip(zipfile = output_dir, files = files2zip)
       zip_ext <- ".zip"
    }

    if (rm_dir & zip_files) {
        message("Cleaning up the working directory")
        unlink(output_dir, TRUE)
    }
    print("Wormcat Execution completed!")
    print(sprintf("Data is available %s%s", output_dir, zip_ext))
}

.get_system_path_separator <- function() {
    separator <- file.path("dummy", "file")
    # Extract the separator from the generated path
    separator <- substr(separator, nchar("dummy") + 1, nchar(separator) - nchar("file"))
    return(separator)
}

.create_file_nm <- function(output_dir, name_format, n = 0) {
    if (n == 0) {
        file_nm <- name_format
    } else {
        file_nm <- sprintf(name_format, n)
    }
    file_path <- file.path(output_dir, file_nm)
    return(file_path)
}


get_local_annotation_file_names <- function() {

    # Get the path to the "extdata" directory within the wormcat package
    extdata_dir <- system.file("extdata", package = "wormcat")

    # List all files in the "extdata" directory
    files_in_extdata <- list.files(extdata_dir, full.names = TRUE)
    annotation_files <- basename(files_in_extdata)

    return(annotation_files)
}


get_remote_annotation_file_names <- function() {
  # URL of the CSV data
  base_url <- "https://dphiggs01.github.io/Wormcat_data/"
  annotation_list_url <- paste0(base_url, "annotation_list.csv")

    # Read CSV data from the URL
  data <- utils::read.csv(annotation_list_url, stringsAsFactors = FALSE)

  # Sort the data by the "order" column
  sorted_data <- data[order(data$order), ]

  # Create a list of tuples
  tuples_list <- lapply(1:nrow(sorted_data), function(i) {
    short_desc <- sorted_data$short_desc[i]
    file_name <- paste0(base_url, sorted_data$location_suffix[i], "/", sorted_data$file_name[i])
    return(list(short_desc, file_name))
  })

  return(tuples_list)
}


copy_annotation_files_to_extdata <- function() {
  # Call the function to get the list of tuples
  annotation_files <- get_remote_annotation_file_names()

  # Directory to copy the files to
  extdata_dir <- system.file("extdata", package = "wormcat")

  # Process and copy files
  for (tuple in annotation_files) {
    short_desc <- tuple[[1]]
    file_url <- tuple[[2]]

    file_name <- gsub("^.*/", "", file_url)
    dest_file <- file.path(extdata_dir, file_name)

    if (!file.exists(dest_file)) {
      utils::download.file(file_url, dest_file)
      cat("Copied:", file_name, "\n")
    } else {
      message(sprintf("Skipped (already exists): %s", file_name))
    }
  }
}

.worm_cat_acceptable_pvalues <- function(rgs_fisher_cat_csv) {
    Bonferroni <- NULL

    rgs_fisher_cat  <- utils::read.csv(rgs_fisher_cat_csv, row.names = 1, header = TRUE, sep = ",")
    rgs_fisher_cat <- na.omit(rgs_fisher_cat)

    rgs_fisher_cat[order(rgs_fisher_cat$PValue), ]

    bonferroni <- p.adjust(rgs_fisher_cat$PValue, method = "bonferroni")
    rgs_fisher_cat <- data.frame(rgs_fisher_cat, Bonferroni = bonferroni)

    ### Acceptable is to be 0.01 on Bonferroni
    # rgs_fisher_cat <- subset(rgs_fisher_cat, PValue < 0.05)
    rgs_fisher_cat <- subset(rgs_fisher_cat, Bonferroni < 0.01)

    csv_out_file_nm <- sprintf("%s_apv.csv", substr(rgs_fisher_cat_csv, 0, nchar(rgs_fisher_cat_csv) - 4))
    utils::write.csv(rgs_fisher_cat, file = csv_out_file_nm, row.names = FALSE)
}

.worm_cat_add_categories <- function(file_to_process, out_dir, worm_cat_annotations, input_type) {
    #log_file <- file("/Users/dan/delme/log.txt", open = "a")
    #message_to_log <- sprintf("Calling .worm_cat_add_categories input_type[%s]",input_type)
    #cat(message_to_log, "\n", file = log_file)
    #close(log_file)

    # Read in the file to process
    file_to_process  <- utils::read.csv(file_to_process, header = TRUE, sep = ",")

    # Read in the annotation file
    annotations  <- utils::read.csv(worm_cat_annotations, header = TRUE, sep = ",")

    # Keep only the first column
    first_column <- file_to_process[, 1]
    file_to_process_df <- data.frame(FirstColumn = first_column)
    colnames(file_to_process_df)[1] <- input_type

    # Remove duplicates from file_to_process
    file_to_process_df <- data.frame(unique(file_to_process_df))

    # Remove duplicates from annotations (there really should not be any)
    # But we have found some custom Annotation files have duplicates
    annotations_df <- annotations[!duplicated(annotations[, input_type]), ]

    # Remove other columns from annotations (there really should not be any)
    # But we have found some custom Annotation files have additional columns
    annotations_df <- annotations_df[c(1, 2, 3, 4, 5, 6)]

    # Combine the file_to_process with the annotation file
    rgs_merge <- merge(file_to_process_df, annotations_df, by = input_type, all.x = TRUE)

    # Save merged data as CSV file
    rgs_and_categories <- file.path(out_dir, "rgs_and_categories.csv")
    utils::write.csv(rgs_merge, file = rgs_and_categories)
}

.worm_cat_bubble_plot <- function(csv_file_name, plot_title) {
    Category <- NULL
    bubbles_z <- NULL

    bubbles <- utils::read.csv(csv_file_name, header = TRUE, sep = ",")

    bubbles <- na.omit(bubbles)

    bubbles <- rbind(bubbles, data.frame("Category" = "calibration high", "RGS" = 250, "AC" = 0, "PValue" = 1.00E-50, "Bonferroni" = 1.00E-50))
    bubbles <- rbind(bubbles, data.frame("Category" = "calibration low", "RGS" = 1, "AC" = 0, "PValue" = 1, "Bonferroni" = 1))


    bubbles$bubbles_z <- round(0.001 * (bubbles$PValue - mean(bubbles$PValue)) / sd(bubbles$PValue), 2)  # compute normalized value as a placeholder

    bubbles$p_value_type <- ifelse(bubbles$Bonferroni < 1e-40, "Col1",
                                   ifelse(bubbles$Bonferroni < 1e-20, "Col2",
                                        ifelse(bubbles$Bonferroni < 1e-10, "Col3",
                                               ifelse(bubbles$Bonferroni < 1e-5, "Col4",
                                                      ifelse(bubbles$Bonferroni < 1e-2, "Col5",
                                                             ifelse(bubbles$Bonferroni < 5e-2, "Col6", "NS"))))))


    bubbles$RGS_size <- ifelse(bubbles$RGS > 150, "Size1",
                               ifelse(bubbles$RGS > 100, "Size2",
                                    ifelse(bubbles$RGS > 75, "Size3",
                                           ifelse(bubbles$RGS > 50, "Size4",
                                                  ifelse(bubbles$RGS > 25, "Size5",
                                                         ifelse(bubbles$RGS > 10, "Size6",
                                                                ifelse(bubbles$RGS > 5, "Size7",
                                                                       ifelse(bubbles$RGS > 2, "Size8", "Size9"))))))))


    bubbles_plot <- bubbles[order(bubbles$PValue), ]  # sort

    myplot <- ggplot(.reorder_by(Category, ~ -PValue, bubbles_plot),
                     aes(x = Category, y = bubbles_z, size = bubbles_plot$RGS_size)) +

                    geom_point(stat = "identity", aes(col = bubbles_plot$p_value_type))  +

                    scale_color_manual(name = "P value",
                                       labels = c("10-40", "10-20", "10-10", "10-5", "0.001", "0.05", "NS"),
                                       values = c("Col1" = "red4", "Col2" = "orangered3", "Col3" = "orangered1",
                                                  "Col4" = "orange", "Col5" = "gold", "Col6" = "yellow", "NS" = "grey48"),
                                       limits = c("Col1", "Col2", "Col3", "Col4", "Col5", "Col6", "NS")) +
                    scale_size_manual(name = "Count",
                                      labels = c("150", "100", "75", "50", "25", "10", "5", "2", "Zero"),
                                      values = c("Size1" = 10, "Size2" = 9,
                                                 "Size3" = 8, "Size4" = 7, "Size5" = 6, "Size6" = 5, "Size7" = 2.5, "Size8" = 1, "Size9" = 0.1),
                                      limits = c("Size1", "Size2",
                                                 "Size3", "Size4", "Size5", "Size6", "Size7", "Size8", "Size9")
                                      ) +


                    labs(title = plot_title) +
                    ylim(-1.0, 1.0) +
                    coord_flip() +
                    theme(panel.grid = element_blank(),
                          panel.background = element_blank(),
                          legend.key = element_rect(fill = "white", colour = "white"),
                          text = element_text(family = "Arial", size = 14))

    s_from <- 0
    s_to <- nchar(csv_file_name) - 4
    file_out_name <- sprintf("%s.svg", substr(csv_file_name, s_from, s_to))
    ggsave(filename = file_out_name,
           plot = myplot,
           device = "svg",
           width = 8, height = 7)
    file_out_name <- sprintf("%s.png", substr(csv_file_name, s_from, s_to))
    ggsave(filename = file_out_name,
           plot = myplot,
           device = "png",
           width = 8, height = 7)
    #message(paste("Plot saved to:", file_out_name))

}

.reorder_by <- function(fact, by, data, FUN = NULL, df = TRUE){

  if(by[[1]] != "~") {
    stop("Argument 'by' must be a one-sided formula.")
  }

  x <- data

  form1 <- as.character(substitute(fact))
  form2 <- as.character(by[[2]])
  check1 <- c(length(form2) == 1 && form1 == form2)
  check2 <- c(length(form2) == 2 && form1 == form2[2] && form2[1] == "-")

  if(check1 | check2) {
    if (is.null(FUN)) {
      warning("FUN not provided: `length` assumed")
      FUN <- length
    }
    ord <- match.fun(ifelse(check2, "rev", "c"))
    lvls <- ord(sort(tapply(data[, form1], data[, form1], FUN = FUN)))
    data[, form1] <- factor(data[, form1], levels = names(lvls))
    return(data)
  }


  fact <- as.character(substitute(fact))
  # Make the formula into character and remove spaces
  formc <- as.character(by[2])
  formc <- gsub(" ", "", formc)
  # If the first character is not + or -, add +
  if(!is.element(substring(formc, 1, 1), c("+", "-")))
    formc <- paste("+", formc, sep = "")

  # Extract the variables from the formula
  vars <- unlist(strsplit(formc, "[\\+\\-]"))
  vars <- vars[vars != ""] # Remove any extra "" terms

  ## use for aggregating
  if (!is.null(FUN)) {
    x <- eval(parse(text=paste0("aggregate(cbind(", paste(vars, collapse = ", "), ") ~",
                                fact, ", data = data, FUN = \"", substitute(FUN), "\")")))
  }

  # Build a list of arguments to pass to "order" function
  calllist <- list()
  pos <- 1 # Position of + or -
  for(i in 1:length(vars)){
    varsign <- substring(formc, pos, pos)
    pos <- pos + 1 + nchar(vars[i])
    if(is.factor(x[, vars[i]])){
      if(varsign == "-") {
        calllist[[i]] <- -rank(x[, vars[i]])
      } else {
        calllist[[i]] <- rank(x[, vars[i]])
      }
    } else {
      if(varsign == "-") {
        calllist[[i]] <- -x[, vars[i]]
      } else {
        calllist[[i]] <- x[,vars[i]]
      }
    }
  }
  data[, fact] <- factor(data[, fact], levels = x[do.call("order", calllist), fact])
  if (df) {
    data
  } else {
    data[, fact]
  }
}


.worm_cat_fisher_test <- function(output_dir, annotations_csv) {
    # Read in annotations file
    annotations <- utils::read.csv(annotations_csv, header = TRUE, sep = ",")

    # Read in rgs mapped to annotations
    rgs_and_categories_csv <- file.path(output_dir, "rgs_and_categories.csv")
    regulated_gene_set <- utils::read.csv(rgs_and_categories_csv, header = TRUE, sep = ",")

    # Step 1 Count categories in annotation files *AC*, will stay static

    total_annotations_count <- data.frame(nrow(annotations))

    annotated_cat1 <- data.frame(table(annotations$Category.1))
    annotated_cat2 <- data.frame(table(annotations$Category.2))
    annotated_cat3 <- data.frame(table(annotations$Category.3))

    # Step 2/3: Count categories in regulated gene sets *RGS*

    total_rgs_count <- data.frame(nrow(regulated_gene_set))

    rgs_annotated_cat1 <- data.frame(table(regulated_gene_set$Category.1))
    rgs_annotated_cat2 <- data.frame(table(regulated_gene_set$Category.2))
    rgs_annotated_cat3 <- data.frame(table(regulated_gene_set$Category.3))

    .merger_cats(rgs_annotated_cat1, annotated_cat1, total_annotations_count$nrow, total_rgs_count$nrow, output_dir, 1)
    .merger_cats(rgs_annotated_cat2, annotated_cat2, total_annotations_count$nrow, total_rgs_count$nrow, output_dir, 2)
    .merger_cats(rgs_annotated_cat3, annotated_cat3, total_annotations_count$nrow, total_rgs_count$nrow, output_dir, 3)
}

.merger_cats <- function(rgs_annotated_cat, annotated_cat, total_annotations_count, total_rgs_count, output_dir, n) {

    merged_cats <- merge(rgs_annotated_cat, annotated_cat, by = "Var1", all.x = TRUE)

    merged_cats <- plyr::rename(merged_cats, c("Var1" = "Category", "Freq.x" = "RGS", "Freq.y" = "AC"))

    # Step 5: Build contingency table for each category in RGS vs AC

    # con <- file("test.log")
    # sink(con, append=TRUE)
    # sink(con, append=TRUE, type="message")

    df <- data.frame(Category = character(),
                     RGS = double(),
                     AC = double(),
                     PValue = double(),
                     stringsAsFactors = FALSE)

    fact_character <- levels(merged_cats$Category)[as.numeric(merged_cats$Category)]

    for (i in 1:nrow(merged_cats)) {
        if (is.na(merged_cats$RGS[i]) | is.na(merged_cats$AC[i])) {
            pvalue <- NA
        } else {
            stat <- fisher.test(matrix(c(merged_cats$RGS[i], total_rgs_count,
                                         merged_cats$AC[i],  total_annotations_count),
                                 nrow = 2, ncol = 2),
                          alternative = "greater")
            pvalue <- stat$p.value
        }

        df[nrow(df) + 1, ] <- list(Category = fact_character[i],
                                   RGS = merged_cats$RGS[i],
                                   AC = merged_cats$AC[i], pvalue)
    }

    sorted_df <- df[with(df, order(PValue)), ]
    file_nm <- .create_file_nm(output_dir, "rgs_fisher_cat%d.csv", n)
    utils::write.csv(sorted_df, file = file_nm)
}
```
