## Supplementary material for "A lineage-resolved multimodal single-cell atlas reveals the genomic dynamics of early *C. elegans* development": Code and external data: AS_downstream_coverage_combined.py

```
import sys

AS_coverage = sys.argv[1]
flank_coverage = sys.argv[2]
output_file = sys.argv[3]
strand = sys.argv[4]

new = open(output_file, "w")
new.write("rep1_AS\trep2_AS\trep1_downstream\trep2_downstream\n")

if strand == "fwd":
    flank = "right"
else:
    flank = "left"

AS_cvg_dict = {}
for line in open(AS_coverage):
    G = line.rstrip().split("\t")
    AS_cvg_dict[G[3]] = [G[4], G[5]]

for line in open(flank_coverage):
    G =	line.rstrip().split("\t")
    K = G[3].split("_")
    if K[3] == flank:
        AS_name = "_".join(K[0:3])
        AS_cvg_dict[AS_name].extend(G[4:])
        new.write(AS_name + "\t" + "\t".join(AS_cvg_dict[AS_name]) + "\n")

new.close()
```
