## Supplementary material for "A lineage-resolved multimodal single-cell atlas reveals the genomic dynamics of early *C. elegans* development": Code and external data: atac_fragment_sample_barcode.20231107.sh

```
#!/bin/bash
#
#SBATCH --job-name=frags_name
#SBATCH --output=slurm_out/frags_name.%N.%j.out
#SBATCH --error=slurm_err/frags_name.%N.%j.err
#SBATCH --ntasks=1
#SBATCH -N 1
#SBATCH -n 1
#SBATCH --mem-per-cpu=4000
#SBATCH --mail-type=END,FAIL
#SBATCH --mail-user=

sample_id=$1

input_fragments="data/sc/cellranger_arc/"$sample_id"/outs/atac_fragments_no_dash.tsv.gz"
output_fragments="data/sc/cellranger_arc/"$sample_id"/outs/atac_fragments_barcode_corrected.tsv"

zcat $input_fragments | grep "#" > $output_fragments
zcat $input_fragments | grep -v "#" | awk -v var="$sample_id" 'BEGIN{OFS="\t";}{print $1, $2, $3, var "_" $4, $5}' >> $output_fragments
bgzip $output_fragments
tabix -p bed $output_fragments.gz
```
