## Supplementary material for "A lineage-resolved multimodal single-cell atlas reveals the genomic dynamics of early *C. elegans* development": Code and external data: atac_fragments_tsv_parser.20230502.py

```
import sys
import os

input_fragments=sys.argv[1]
output_fragments=sys.argv[2]

cmd = "gunzip -c " + input_fragments + ".gz > " + input_fragments
os.system(cmd)

new=open(output_fragments, "w")
for line in open(input_fragments):
    if line[0] == "#":
        new.write(line)
    else:
        G=line.split("\t")
        G[3] = G[3].split("-")[0]
        new.write("\t".join(G))

new.close()
cmd = "bgzip " + output_fragments
os.system(cmd)
cmd = "tabix -p bed " + output_fragments + ".gz"
os.system(cmd)
```
