## Supplementary material for "A lineage-resolved multimodal single-cell atlas reveals the genomic dynamics of early *C. elegans* development": Code and external data: atac_fragments_tsv_parser.sh

```
#!/bin/bash
#
#SBATCH --job-name=STARsolo
#SBATCH --output=slurm_out/STARsolo.%N.%j.out
#SBATCH --error=slurm_err/STARsolo.%N.%j.err
#SBATCH --ntasks=1
#SBATCH -N 1
#SBATCH -n 1
#SBATCH --mem=32G
#SBATCH --mail-type=END,FAIL
#SBATCH --mail-user=

input_frags=$1
output_frags=$2

python scripts/atac_fragments_tsv_parser.20230502.py $input_frags $output_frags
```
