## Supplementary material for "A lineage-resolved multimodal single-cell atlas reveals the genomic dynamics of early *C. elegans* development": Code and external data: ATAC_frags_for_IDR.R

```
# compare gene expression in 2-cell and 4-cell clusters to identify sets of newly transcribed genes (ZGA)
library(Seurat)
library(Signac)
library(Hmisc)
library(RColorBrewer)
library(presto)
library(rtracklayer)

source("data/external_data/wormcat_function.R")

current_date= paste(unlist(strsplit(as.character(Sys.Date()), "-")), collapse="")
plot_dir_w_date = paste0("plots/", current_date)
dir.create(plot_dir_w_date, recursive=TRUE)

# load RDS and WBid table
combined_seurat = readRDS("seurat_objects/wt_all.all_samples.all_cells.WS285_extended_no_ovlp.20240911.rds")
wb_id_common_names = read.table("species/elegans/gene_annotation/c_elegans.PRJNA13758.WS285.canonical_geneset.filtered.gene_name.txt")


# load new MACS peaks
macs_peaks = import.bed("cluster_specific_peaks/round_2_all_samples/WS285_extended_no_ovlp/all_peaks.pre_IDR.bed")

macs_peaks_matrix <- FeatureMatrix(combined_seurat[["WNN"]]@fragments, sep = c("-", "-"), features=macs_peaks, cells = colnames(combined_seurat))
combined_seurat[["macs_peaks"]] <- CreateChromatinAssay(counts = macs_peaks_matrix, sep = c(":", "-"), fragments = combined_seurat[["WNN"]]@fragments)

# get aggregate ATAC-seq fragment counts from a given cell type after dividing the object in cells belonging to a two sets of samples
# split object and get aggregate counts
combined_seurat_1 = subset(combined_seurat, orig.ident %in% c("exp024", "exp031", "exp042"))
combined_seurat_2 = subset(combined_seurat, orig.ident %in% c("exp024", "exp031", "exp042"), invert = TRUE)

combined_seurat_1_agg_atac = as.data.frame(AggregateExpression(combined_seurat_1, assays = "macs_peaks", group.by = "cell_type"))
names(combined_seurat_1_agg_atac) = sapply(strsplit(names(combined_seurat_1_agg_atac), split = "macs_peaks."), "[[", 2)

combined_seurat_2_agg_atac = as.data.frame(AggregateExpression(combined_seurat_2, assays = "macs_peaks", group.by = "cell_type"))
names(combined_seurat_2_agg_atac) = sapply(strsplit(names(combined_seurat_2_agg_atac), split = "macs_peaks."), "[[", 2)

# output aggregate counts as score filed of a bed file
dir.create("cluster_specific_peaks/round_2_all_samples/WS285_extended_no_ovlp/cell_type/macs_peak_frag_counts", recursive=TRUE)

for (cell_type in names(combined_seurat_1_agg_atac)){
  cell_type_rep = as.data.frame(matrix(unlist(strsplit(row.names(combined_seurat_1_agg_atac), split = "-")), ncol = 3, byrow = T))
  cell_type_rep$V2 = as.numeric(cell_type_rep$V2) - 1
  cell_type_rep = cbind(cell_type_rep, paste0("embryo_AS_", as.character(c(1:dim(cell_type_rep)[1]))))
  
  cell_type_rep_1 = cbind(cell_type_rep, combined_seurat_1_agg_atac[,cell_type])
  cell_type_rep_1 = cbind(cell_type_rep_1, rep(".", dim(cell_type_rep)[1]))
  cell_type_rep_1 = cbind(cell_type_rep_1, combined_seurat_1_agg_atac[,cell_type])
  cell_type_rep_1 = cbind(cell_type_rep_1, combined_seurat_1_agg_atac[,cell_type])
  cell_type_rep_1 = cbind(cell_type_rep_1, combined_seurat_1_agg_atac[,cell_type])
  cell_type_rep_1 = cbind(cell_type_rep_1, 100)
  names(cell_type_rep_1) = c("chr", "start", "end", "name", "score", "strand", "signalValue", "pValue", "qValue", "peak")
  
  cell_type_rep_2 = cbind(cell_type_rep, combined_seurat_2_agg_atac[,cell_type])
  cell_type_rep_2 = cbind(cell_type_rep_2, rep(".", dim(cell_type_rep)[1]))
  cell_type_rep_2 = cbind(cell_type_rep_2, combined_seurat_2_agg_atac[,cell_type])
  cell_type_rep_2 = cbind(cell_type_rep_2, combined_seurat_2_agg_atac[,cell_type])
  cell_type_rep_2 = cbind(cell_type_rep_2, combined_seurat_2_agg_atac[,cell_type])
  cell_type_rep_2 = cbind(cell_type_rep_2, 100)
  names(cell_type_rep_2) = c("chr", "start", "end", "name", "score", "strand", "signalValue", "pValue", "qValue", "peak")
  
  write.table(cell_type_rep_1, file = paste0("cluster_specific_peaks/round_2_all_samples/WS285_extended_no_ovlp/cell_type/macs_peak_frag_counts/", cell_type, "_rep_1.bed"), append = F, quote = F, sep = "\t", row.names = F, col.names = F)
  write.table(cell_type_rep_2, file = paste0("cluster_specific_peaks/round_2_all_samples/WS285_extended_no_ovlp/cell_type/macs_peak_frag_counts/", cell_type, "_rep_2.bed"), append = F, quote = F, sep = "\t", row.names = F, col.names = F)
}

write.table(names(combined_seurat_1_agg_atac), file = paste0("cluster_specific_peaks/round_2_all_samples/WS285_extended_no_ovlp/cell_type/macs_peak_frag_counts/cell_types.txt"), append = F, quote = F, sep = "\t", row.names = F, col.names = F)

k1 = table(combined_seurat_1$cell_type)
k2 = table(combined_seurat_2$cell_type)
k = matrix(c(k1, k2), ncol = 2, byrow = F, dimnames = list(names(k1), c("rep1", "rep2")))

write.table(k, file = paste0("cluster_specific_peaks/round_2_all_samples/WS285_extended_no_ovlp/cell_type/macs_peak_frag_counts/cell_number_by_type.txt"), append = F, quote = F, sep = "\t", row.names = T, col.names = T)
```
