## Supplementary material for "A lineage-resolved multimodal single-cell atlas reveals the genomic dynamics of early *C. elegans* development": Code and external data: bam_barcode_split.nosuffix.sh

```
#!/bin/bash
#
#SBATCH --job-name=sinto
#SBATCH --output=slurm_out/sinto.%N.%j.out
#SBATCH --error=slurm_err/sinto.%N.%j.err
#SBATCH --ntasks=1
#SBATCH -N 1
#SBATCH -n 12
#SBATCH --mem-per-cpu=8000
#SBATCH --mail-type=END,FAIL
#SBATCH --mail-user=

input_bam=$1
input_barcodes=$2
output_dir=$3

mkdir -p $output_dir
samtools index $input_bam
sinto filterbarcodes -b $input_bam -c $input_barcodes --outdir $output_dir -p 12
```
