## Supplementary material for "A lineage-resolved multimodal single-cell atlas reveals the genomic dynamics of early *C. elegans* development": Code and external data: batch_enriched_clusters_removal.20240430.R

```
# QC on the merged dataset and clusters removal
library(Seurat)
library(Signac)
library(ggplot2)
library(Hmisc)
library(SoupX)
library(glmGamPoi)	
library(presto)	

args = commandArgs(trailingOnly=TRUE)

current_date= paste(unlist(strsplit(as.character(Sys.Date()), "-")), collapse="")
plot_dir_w_date = paste0("plots/", current_date)
dir.create(plot_dir_w_date, recursive=TRUE)

seurat_object = args[1]
annotation = args[2]

combined_seurat = readRDS(seurat_object)

### identify clusters to remove as those showing a strong skew in their batch of origin
## increase resolution of clustering
combined_seurat <- FindClusters(combined_seurat, graph.name = "wsnn", algorithm = 3, verbose = T, resolution = 5)

combined_seurat_UMAP = DimPlot(combined_seurat, reduction = "umapWNN", label = T) & NoLegend()

## mark the very early cells (2-cell and ABx) to avoid their removal (underrepresented due to differences in stageing)
#combined_seurat = PrepSCTFindMarkers(combined_seurat)
DefaultAssay(combined_seurat) = "RNA"
all_markers = FindAllMarkers(combined_seurat,assay = "RNA", logfc.threshold = 0.5, only.pos = T)

# get the early clusters based on enrichment of pes-10 and/or vet-2
early_clusters = all_markers[all_markers$gene %in% c("pes-10", "vet-2"),6]

# flag the 2-cell and ABx clusters
twocell_cluster = all_markers[all_markers$gene == "cpg-3",6]
abx_cluster = all_markers[all_markers$gene == "ccch-2",6][which(all_markers[all_markers$gene == "ccch-2",6] %nin% early_clusters)]

## determine proportion of cells from different experiments for each cluster
exp_cell_mtx = matrix(rep(NA, 7*length(unique(combined_seurat$seurat_clusters))), ncol = length(unique(combined_seurat$seurat_clusters)))
exp_cell_mtx[1,] = table(combined_seurat$seurat_clusters[combined_seurat$exp == "exp031"])
exp_cell_mtx[2,] = table(combined_seurat$seurat_clusters[combined_seurat$exp == "exp032"])
exp_cell_mtx[3,] = table(combined_seurat$seurat_clusters[combined_seurat$exp == "exp042"])
exp_cell_mtx[4,] = table(combined_seurat$seurat_clusters[combined_seurat$exp == "exp043"])
exp_cell_mtx[5,] = table(combined_seurat$seurat_clusters[combined_seurat$exp == "exp024"])
exp_cell_mtx[6,] = table(combined_seurat$seurat_clusters[combined_seurat$exp == "exp025"])
exp_cell_mtx[7,] = table(combined_seurat$seurat_clusters[combined_seurat$exp == "exp027"])
row.names(exp_cell_mtx) = c("exp031", "exp032", "exp042", "exp043", "exp024", "exp025", "exp027")
colnames(exp_cell_mtx) = seq(0, length(unique(combined_seurat$seurat_clusters))-1)

exp_cell_mtx_fraction = apply(exp_cell_mtx, 2, function(x){x/sum(x)})

batch_cell_mtx = rbind(exp_cell_mtx[1,] + exp_cell_mtx[2,], exp_cell_mtx[3,] + exp_cell_mtx[4,], exp_cell_mtx[5,] + exp_cell_mtx[6,] + exp_cell_mtx[7,])
row.names(batch_cell_mtx) = c("batch1", "batch2","batch3")
batch_cell_mtx_fraction = apply(batch_cell_mtx, 2, function(x){x/sum(x)})

#batch_cell_df_fraction = as.data.frame(batch_cell_mtx_fraction)
batch_enriched_clusters_EE = names(which(batch_cell_mtx_fraction[1,] > batch_cell_mtx_fraction[2,] * 4 | batch_cell_mtx_fraction[2,] > batch_cell_mtx_fraction[1,] * 4))
batch_enriched_clusters_LE = names(which(batch_cell_mtx_fraction[3,] > 0.8))

batch_enriched_clusters = sort(unique(c(batch_enriched_clusters_EE, batch_enriched_clusters_LE)))
# remove 2-cell/ABx if present
batch_enriched_clusters = batch_enriched_clusters[as.character(batch_enriched_clusters) %nin% c(twocell_cluster, abx_cluster)]

batch_cell_mtx_fraction_clean = batch_cell_mtx_fraction[, colnames(batch_cell_mtx_fraction) %nin% batch_enriched_clusters]

batch_enriched_barcodes = as.character(combined_seurat$seurat_clusters)
batch_enriched_barcodes[batch_enriched_barcodes %in% batch_enriched_clusters] = "batch_enriched"
batch_enriched_barcodes[combined_seurat$seurat_clusters %nin% batch_enriched_clusters] = "not_enriched"

combined_seurat$batch_enrichment = batch_enriched_barcodes
p_batch = DimPlot(combined_seurat, group.by = "batch_enrichment", order=T, reduction = "umapWNN")

# plot batch enrichment
pdf(file=paste0(plot_dir_w_date, "/merged_seurat.", annotation, ".batch_enrichment.pdf"), width=16, height = 8)
par(mfrow=c(1,1))
barplot(exp_cell_mtx_fraction, beside = F, las = 1, main = "exp of origin")
abline(h=0.80, lwd = 2, lty = 2)
abline(h=0.20, lwd = 2, lty = 2)
barplot(batch_cell_mtx_fraction, beside = F, las = 1, main = "batch of origin")
abline(h=0.80, lwd = 2, lty = 2)
abline(h=0.20, lwd = 2, lty = 2)
print(combined_seurat_UMAP | p_batch)
dev.off()

# pre-processing: normalise RNA for FeaturePlots
DefaultAssay(combined_seurat) = "RNA"
combined_seurat = NormalizeData(combined_seurat)

# QC 1: P-granule gene expression
P_granule_genes = read.table("data/external_data/P_granule_transcripts.sorted.txt")
DefaultAssay(combined_seurat) = "RNA"
combined_seurat = AddModuleScore(combined_seurat, features = list(P_granule_genes$V1), name = "Pgranule")

p_pcell = FeaturePlot(combined_seurat, label = T, repel = T, label.size = 2, label.color = 2, 
            features = c("Pgranule1"), reduction = "umapWNN",
            order=T) & scale_colour_gradientn(colours = c("grey90", "cyan", "deepskyblue", "darkorange", "darkred", "darkorchid3"), na.value = "grey90")

# QC 2: ambient RNA
# read in all raw matrices
rnaseq_data_raw_exp031 <- Seurat::Read10X(paste0("data/sc/rnaseq/exp031/star_", annotation, "/star_", annotation, "_Solo.out/GeneFull/raw/um_reads"))
rnaseq_data_raw_exp032 <- Seurat::Read10X(paste0("data/sc/rnaseq/exp032/star_", annotation, "/star_", annotation, "_Solo.out/GeneFull/raw/um_reads"))
rnaseq_data_raw_exp042 <- Seurat::Read10X(paste0("data/sc/rnaseq/exp042/star_", annotation, "/star_", annotation, "_Solo.out/GeneFull/raw/um_reads"))
rnaseq_data_raw_exp043 <- Seurat::Read10X(paste0("data/sc/rnaseq/exp043/star_", annotation, "/star_", annotation, "_Solo.out/GeneFull/raw/um_reads"))

rnaseq_data_raw_exp024 <- Seurat::Read10X(paste0("data/sc/rnaseq/exp024/star_", annotation, "/star_", annotation, "_Solo.out/GeneFull/raw/um_reads"))
rnaseq_data_raw_exp025 <- Seurat::Read10X(paste0("data/sc/rnaseq/exp025/star_", annotation, "/star_", annotation, "_Solo.out/GeneFull/raw/um_reads"))
rnaseq_data_raw_exp027 <- Seurat::Read10X(paste0("data/sc/rnaseq/exp027/star_", annotation, "/star_", annotation, "_Solo.out/GeneFull/raw/um_reads"))

colnames(rnaseq_data_raw_exp031) = paste0("exp031_", colnames(rnaseq_data_raw_exp031))
colnames(rnaseq_data_raw_exp032) = paste0("exp032_", colnames(rnaseq_data_raw_exp032))
colnames(rnaseq_data_raw_exp042) = paste0("exp042_", colnames(rnaseq_data_raw_exp042))
colnames(rnaseq_data_raw_exp043) = paste0("exp043_", colnames(rnaseq_data_raw_exp043))

colnames(rnaseq_data_raw_exp024) = paste0("exp024_", colnames(rnaseq_data_raw_exp024))
colnames(rnaseq_data_raw_exp025) = paste0("exp025_", colnames(rnaseq_data_raw_exp025))
colnames(rnaseq_data_raw_exp027) = paste0("exp027_", colnames(rnaseq_data_raw_exp027))

rnaseq_data_raw_batch1 = cbind(rnaseq_data_raw_exp031, rnaseq_data_raw_exp032)
rnaseq_data_raw_batch2 = cbind(rnaseq_data_raw_exp042, rnaseq_data_raw_exp043)
rnaseq_data_raw_batch3 = cbind(rnaseq_data_raw_exp024, rnaseq_data_raw_exp025, rnaseq_data_raw_exp027)

rnaseq_data_raw_batch1_filtered <- rnaseq_data_raw_batch1[row.names(rnaseq_data_raw_batch1) %in% row.names(combined_seurat@assays$RNA@counts),]
rnaseq_data_raw_batch2_filtered <- rnaseq_data_raw_batch2[row.names(rnaseq_data_raw_batch2) %in% row.names(combined_seurat@assays$RNA@counts),]
rnaseq_data_raw_batch3_filtered <- rnaseq_data_raw_batch3[row.names(rnaseq_data_raw_batch3) %in% row.names(combined_seurat@assays$RNA@counts),]

counts_batch1_combined_seurat = combined_seurat[,combined_seurat$exp %in% c("exp031", "exp032")]@assays$RNA@counts
counts_batch1_combined_seurat = counts_batch1_combined_seurat[row.names(counts_batch1_combined_seurat) %in% row.names(rnaseq_data_raw_batch1_filtered),]
counts_batch2_combined_seurat = combined_seurat[,combined_seurat$exp %in% c("exp042", "exp043")]@assays$RNA@counts
counts_batch2_combined_seurat = counts_batch2_combined_seurat[row.names(counts_batch2_combined_seurat) %in% row.names(rnaseq_data_raw_batch2_filtered),]

counts_batch3_combined_seurat = combined_seurat[,combined_seurat$exp %in% c("exp024", "exp025", "exp027")]@assays$RNA@counts
counts_batch3_combined_seurat = counts_batch3_combined_seurat[row.names(counts_batch3_combined_seurat) %in% row.names(rnaseq_data_raw_batch3_filtered),]

sc_batch1 = SoupChannel(rnaseq_data_raw_batch1_filtered, counts_batch1_combined_seurat, calcSoupProfile=F)
sc_batch1 = estimateSoup(sc_batch1, soupRange=c(100, 400))
sc_batch2 = SoupChannel(rnaseq_data_raw_batch2_filtered, counts_batch2_combined_seurat, calcSoupProfile=F)
sc_batch2 = estimateSoup(sc_batch2, soupRange=c(20, 150))
sc_batch3 = SoupChannel(rnaseq_data_raw_batch3_filtered, counts_batch3_combined_seurat, calcSoupProfile=F)
sc_batch3 = estimateSoup(sc_batch3, soupRange=c(100, 200))

high_ambient_batch1 = row.names(sc_batch1$soupProfile[sc_batch1$soupProfile$est > 0.0005,])
high_ambient_batch2 = row.names(sc_batch2$soupProfile[sc_batch2$soupProfile$est > 0.0005,])
high_ambient_batch3 = row.names(sc_batch3$soupProfile[sc_batch3$soupProfile$est > 0.0005,])

high_ambient = high_ambient_batch2[high_ambient_batch2 %in% high_ambient_batch1 & high_ambient_batch2 %in% high_ambient_batch3]

DefaultAssay(combined_seurat) = "RNA"
combined_seurat = AddModuleScore(combined_seurat, features = list(high_ambient), name = "ambient")
p_ambient = FeaturePlot(combined_seurat, label = T, repel = T, label.size = 2, label.color = 2, 
            features = c("ambient1"), reduction = "umapWNN",
            order=T) & scale_colour_gradientn(colours = c("grey90", "cyan", "deepskyblue", "darkorange", "darkred", "darkorchid3"), na.value = "grey90")

# QC 3: rRNA, lincRNA and mito RNA expression
mt_genes <- read.table("species/elegans/gene_annotation/c_elegans.PRJNA13758.WS285.canonical_geneset.filtered.MtDNA_genes.genes")
gene_type = read.table("species/elegans/gene_annotation/c_elegans.PRJNA13758.WS285.canonical_geneset.gene_name_gene_type.txt")

rrna_genes <-  gene_type$V1[gene_type$V2  == "rRNA" & gene_type$V1 %nin% mt_genes$V1]
rrna_genes_expression_pct_exp031 = colSums(rnaseq_data_raw_exp031[row.names(rnaseq_data_raw_exp031) %in% rrna_genes,])/colSums(rnaseq_data_raw_exp031)
rrna_genes_expression_pct_exp032 = colSums(rnaseq_data_raw_exp032[row.names(rnaseq_data_raw_exp032) %in% rrna_genes,])/colSums(rnaseq_data_raw_exp032)
rrna_genes_expression_pct_exp042 = colSums(rnaseq_data_raw_exp042[row.names(rnaseq_data_raw_exp042) %in% rrna_genes,])/colSums(rnaseq_data_raw_exp042)
rrna_genes_expression_pct_exp043 = colSums(rnaseq_data_raw_exp043[row.names(rnaseq_data_raw_exp043) %in% rrna_genes,])/colSums(rnaseq_data_raw_exp043)
rrna_genes_expression_pct_exp024 = colSums(rnaseq_data_raw_exp024[row.names(rnaseq_data_raw_exp024) %in% rrna_genes,])/colSums(rnaseq_data_raw_exp024)
rrna_genes_expression_pct_exp025 = colSums(rnaseq_data_raw_exp025[row.names(rnaseq_data_raw_exp025) %in% rrna_genes,])/colSums(rnaseq_data_raw_exp025)
rrna_genes_expression_pct_exp027 = colSums(rnaseq_data_raw_exp027[row.names(rnaseq_data_raw_exp027) %in% rrna_genes,])/colSums(rnaseq_data_raw_exp027)

rrna_pct = rrna_genes_expression_pct_exp031[names(rrna_genes_expression_pct_exp031) %in% colnames(combined_seurat)]
rrna_pct = append(rrna_pct, rrna_genes_expression_pct_exp032[names(rrna_genes_expression_pct_exp032) %in% colnames(combined_seurat)])
rrna_pct = append(rrna_pct, rrna_genes_expression_pct_exp042[names(rrna_genes_expression_pct_exp042) %in% colnames(combined_seurat)])
rrna_pct = append(rrna_pct, rrna_genes_expression_pct_exp043[names(rrna_genes_expression_pct_exp043) %in% colnames(combined_seurat)])
rrna_pct = append(rrna_pct, rrna_genes_expression_pct_exp024[names(rrna_genes_expression_pct_exp024) %in% colnames(combined_seurat)])
rrna_pct = append(rrna_pct, rrna_genes_expression_pct_exp025[names(rrna_genes_expression_pct_exp025) %in% colnames(combined_seurat)])
rrna_pct = append(rrna_pct, rrna_genes_expression_pct_exp027[names(rrna_genes_expression_pct_exp027) %in% colnames(combined_seurat)])


combined_seurat$rrna_pct = rrna_pct
lincRNA_genes <- gene_type$V1[gene_type$V2  == "lincRNA" & gene_type$V1 %nin% mt_genes$V1]
combined_seurat = AddModuleScore(combined_seurat, features = list(c(lincRNA_genes)), assay = "RNA", name = "lincRNA_RNA", nbin = 10)

p_mito = FeaturePlot(combined_seurat, label = T, repel = T, label.size = 2, label.color = 2, 
            features = c("mito_pct"), reduction = "umapWNN",
            order=T) & scale_colour_gradientn(colours = c("grey90", "cyan", "deepskyblue", "darkorange", "darkred", "darkorchid3"), na.value = "grey90")

p_rrna = FeaturePlot(combined_seurat, label = T, repel = T, label.size = 2, label.color = 2, 
                     features = c("rrna_pct"), reduction = "umapWNN",
                     order=T) & scale_colour_gradientn(colours = c("grey90", "cyan", "deepskyblue", "darkorange", "darkred", "darkorchid3"), na.value = "grey90")

p_lncrna = FeaturePlot(combined_seurat, label = T, repel = T, label.size = 2, label.color = 2, 
                     features = c("lincRNA_RNA1"), reduction = "umapWNN",
                     order=T) & scale_colour_gradientn(colours = c("grey90", "cyan", "deepskyblue", "darkorange", "darkred", "darkorchid3"), na.value = "grey90")


# QC 4: FRiP/ATAC feature n
tot_frag = CountFragments(list("data/sc/cellranger_arc/exp031/outs/atac_fragments_barcode_corrected.tsv.gz",
                               "data/sc/cellranger_arc/exp032/outs/atac_fragments_barcode_corrected.tsv.gz",
                               "data/sc/cellranger_arc/exp042/outs/atac_fragments_barcode_corrected.tsv.gz",
                               "data/sc/cellranger_arc/exp043/outs/atac_fragments_barcode_corrected.tsv.gz",
                               "data/sc/cellranger_arc/exp024/outs/atac_fragments_barcode_corrected.tsv.gz",
                               "data/sc/cellranger_arc/exp025/outs/atac_fragments_barcode_corrected.tsv.gz",
                               "data/sc/cellranger_arc/exp027/outs/atac_fragments_barcode_corrected.tsv.gz"), cells = colnames(combined_seurat), max_lines = NULL, verbose = TRUE)
combined_seurat$tot_frag = tot_frag$frequency_count
combined_seurat = FRiP(object = combined_seurat, assay = "ATAC", total.fragments = "tot_frag", col.name = "FRiP")

p_frip = FeaturePlot(combined_seurat, label = T, repel = T, label.size = 2, label.color = 2, 
                       features = c("FRiP"), reduction = "umapWNN",
                       order=T) & scale_colour_gradientn(colours = c("grey90", "cyan", "deepskyblue", "darkorange", "darkred", "darkorchid3"), na.value = "grey90")

p_wnnFeat = FeaturePlot(combined_seurat, label = T, repel = T, label.size = 2, label.color = 2, 
                        features = c("nFeature_WNN"), reduction = "umapWNN",
                        order=T, max.cutoff = 5000) & scale_colour_gradientn(colours = c("grey90", "cyan", "deepskyblue", "darkorange", "darkred", "darkorchid3"), na.value = "grey90")


# plotting QC
pdf(file=paste0(plot_dir_w_date, "/merged_seurat.", annotation, ".QC.pdf"), width=16, height = 8)

print(combined_seurat_UMAP | p_pcell)

pcell_module = as.data.frame(cbind(combined_seurat$Pgranule1, combined_seurat$seurat_clusters))
pcell_module_mean = aggregate(pcell_module$V1~pcell_module$V2, pcell_module, mean)
pcell_module_mean = pcell_module_mean[order(pcell_module_mean$`pcell_module$V1`, decreasing = T),]
pcell_module_col = as.character(pcell_module_mean$`pcell_module$V2` - 1)
pcell_module_col[pcell_module_col %nin% batch_enriched_clusters] = "deepskyblue"
pcell_module_col[pcell_module_col %in% batch_enriched_clusters] = "darkred"
VlnPlot(combined_seurat, features = "Pgranule1", sort = "increasing", cols = pcell_module_col) & NoLegend()

print(combined_seurat_UMAP | p_ambient)

ambient_module = as.data.frame(cbind(combined_seurat$ambient1, combined_seurat$seurat_clusters))
ambient_module_mean = aggregate(ambient_module$V1~ambient_module$V2, ambient_module, mean)
ambient_module_mean = ambient_module_mean[order(ambient_module_mean$`ambient_module$V1`, decreasing = T),]
ambient_module_col = as.character(ambient_module_mean$`ambient_module$V2` - 1)
ambient_module_col[ambient_module_col %nin% batch_enriched_clusters] = "deepskyblue"
ambient_module_col[ambient_module_col %in% batch_enriched_clusters] = "darkred"
VlnPlot(combined_seurat, features = "ambient1", sort = "increasing", cols = ambient_module_col) & NoLegend()

print(combined_seurat_UMAP | p_mito)

mito_module = as.data.frame(cbind(combined_seurat$mito_pct, combined_seurat$seurat_clusters))
mito_module_mean = aggregate(mito_module$V1~mito_module$V2, mito_module, mean)
mito_module_mean = mito_module_mean[order(mito_module_mean$`mito_module$V1`, decreasing = T),]
mito_module_col = as.character(mito_module_mean$`mito_module$V2` - 1)
mito_module_col[mito_module_col %nin% batch_enriched_clusters] = "deepskyblue"
mito_module_col[mito_module_col %in% batch_enriched_clusters] = "darkred"
VlnPlot(combined_seurat, features = "mito_pct", sort = "increasing", cols = mito_module_col) & NoLegend()

print(combined_seurat_UMAP | p_rrna)

rrna_module = as.data.frame(cbind(combined_seurat$rrna_pct, combined_seurat$seurat_clusters))
rrna_module_mean = aggregate(rrna_module$V1~rrna_module$V2, rrna_module, mean)
rrna_module_mean = rrna_module_mean[order(rrna_module_mean$`rrna_module$V1`, decreasing = T),]
rrna_module_col = as.character(rrna_module_mean$`rrna_module$V2` - 1)
rrna_module_col[rrna_module_col %nin% batch_enriched_clusters] = "deepskyblue"
rrna_module_col[rrna_module_col %in% batch_enriched_clusters] = "darkred"
VlnPlot(combined_seurat, features = "rrna_pct", sort = "increasing", cols = rrna_module_col) & NoLegend()

print(combined_seurat_UMAP | p_lncrna)

lincrna_module = as.data.frame(cbind(combined_seurat$lincRNA_RNA1, combined_seurat$seurat_clusters))
lincrna_module_mean = aggregate(lincrna_module$V1~lincrna_module$V2, lincrna_module, mean)
lincrna_module_mean = lincrna_module_mean[order(lincrna_module_mean$`lincrna_module$V1`, decreasing = T),]
lincrna_module_col = as.character(lincrna_module_mean$`lincrna_module$V2` - 1)
lincrna_module_col[lincrna_module_col %nin% batch_enriched_clusters] = "deepskyblue"
lincrna_module_col[lincrna_module_col %in% batch_enriched_clusters] = "darkred"
VlnPlot(combined_seurat, features = "lincRNA_RNA1", sort = "increasing", cols = lincrna_module_col) & NoLegend()

print(combined_seurat_UMAP | p_frip)

frip_module = as.data.frame(cbind(combined_seurat$FRiP, combined_seurat$seurat_clusters))
frip_module_mean = aggregate(frip_module$V1~frip_module$V2, frip_module, mean)
frip_module_mean = frip_module_mean[order(frip_module_mean$`frip_module$V1`, decreasing = T),]
frip_module_col = as.character(frip_module_mean$`frip_module$V2` - 1)
frip_module_col[frip_module_col %nin% batch_enriched_clusters] = "deepskyblue"
frip_module_col[frip_module_col %in% batch_enriched_clusters] = "darkred"
VlnPlot(combined_seurat, features = "FRiP", sort = "increasing", cols = frip_module_col) & NoLegend()

print(combined_seurat_UMAP | p_wnnFeat)

atacfrag_module = as.data.frame(cbind(combined_seurat$nFeature_WNN, combined_seurat$seurat_clusters))
atacfrag_module_mean = aggregate(atacfrag_module$V1~atacfrag_module$V2, atacfrag_module, mean)
atacfrag_module_mean = atacfrag_module_mean[order(atacfrag_module_mean$`atacfrag_module$V1`, decreasing = T),]
atacfrag_module_col = as.character(atacfrag_module_mean$`atacfrag_module$V2` - 1)
atacfrag_module_col[atacfrag_module_col %nin% batch_enriched_clusters] = "deepskyblue"
atacfrag_module_col[atacfrag_module_col %in% batch_enriched_clusters] = "darkred"
VlnPlot(combined_seurat, features = "nFeature_WNN", sort = "increasing", log=T, cols = atacfrag_module_col) & NoLegend()
dev.off()

# remove batch-enriched cells and cells with < 200 WNN features and re-cluster the data
cells_to_keep = colnames(subset(combined_seurat, subset = nFeature_WNN > 200 & seurat_clusters %nin% batch_enriched_clusters))

# extract original RNA and WNN count matrix and fragments of the selected cells for each experiment
exp1_RNA_mtx = combined_seurat[,combined_seurat$exp == "exp031" & colnames(combined_seurat) %in% cells_to_keep]@assays$RNA@counts
exp2_RNA_mtx = combined_seurat[,combined_seurat$exp == "exp032" & colnames(combined_seurat) %in% cells_to_keep]@assays$RNA@counts
exp3_RNA_mtx = combined_seurat[,combined_seurat$exp == "exp042" & colnames(combined_seurat) %in% cells_to_keep]@assays$RNA@counts
exp4_RNA_mtx = combined_seurat[,combined_seurat$exp == "exp043" & colnames(combined_seurat) %in% cells_to_keep]@assays$RNA@counts
exp5_RNA_mtx = combined_seurat[,combined_seurat$exp == "exp024" & colnames(combined_seurat) %in% cells_to_keep]@assays$RNA@counts
exp6_RNA_mtx = combined_seurat[,combined_seurat$exp == "exp025" & colnames(combined_seurat) %in% cells_to_keep]@assays$RNA@counts
exp7_RNA_mtx = combined_seurat[,combined_seurat$exp == "exp027" & colnames(combined_seurat) %in% cells_to_keep]@assays$RNA@counts

exp1_ATAC_mtx = combined_seurat[,combined_seurat$exp == "exp031" & colnames(combined_seurat) %in% cells_to_keep]@assays$WNN@counts
exp2_ATAC_mtx = combined_seurat[,combined_seurat$exp == "exp032" & colnames(combined_seurat) %in% cells_to_keep]@assays$WNN@counts
exp3_ATAC_mtx = combined_seurat[,combined_seurat$exp == "exp042" & colnames(combined_seurat) %in% cells_to_keep]@assays$WNN@counts
exp4_ATAC_mtx = combined_seurat[,combined_seurat$exp == "exp043" & colnames(combined_seurat) %in% cells_to_keep]@assays$WNN@counts
exp5_ATAC_mtx = combined_seurat[,combined_seurat$exp == "exp024" & colnames(combined_seurat) %in% cells_to_keep]@assays$WNN@counts
exp6_ATAC_mtx = combined_seurat[,combined_seurat$exp == "exp025" & colnames(combined_seurat) %in% cells_to_keep]@assays$WNN@counts
exp7_ATAC_mtx = combined_seurat[,combined_seurat$exp == "exp027" & colnames(combined_seurat) %in% cells_to_keep]@assays$WNN@counts

all_fragments = combined_seurat[,colnames(combined_seurat) %in% cells_to_keep]@assays$WNN@fragments

# create separate Seurat files and normalise RNA and ATAC counts
exp1_seurat <- CreateSeuratObject(counts = exp1_RNA_mtx, project = "exp031_clean")
exp1_seurat <- SCTransform(exp1_seurat, vst.flavor = "v2", method = "glmGamPoi")
exp1_seurat[["WNN"]] <- CreateChromatinAssay(counts = exp1_ATAC_mtx, sep = c(":", "-"), fragments = all_fragments[[1]])
DefaultAssay(exp1_seurat) = "WNN"
exp1_seurat <- RunTFIDF(exp1_seurat)

exp2_seurat <- CreateSeuratObject(counts = exp2_RNA_mtx, project = "exp032_clean")
exp2_seurat <- SCTransform(exp2_seurat, vst.flavor = "v2", method = "glmGamPoi")
exp2_seurat[["WNN"]] <- CreateChromatinAssay(counts = exp2_ATAC_mtx, sep = c(":", "-"), fragments = all_fragments[[2]])
DefaultAssay(exp2_seurat) = "WNN"
exp2_seurat <- RunTFIDF(exp2_seurat)

exp3_seurat <- CreateSeuratObject(counts = exp3_RNA_mtx, project = "exp042_clean")
exp3_seurat <- SCTransform(exp3_seurat, vst.flavor = "v2", method = "glmGamPoi")
exp3_seurat[["WNN"]] <- CreateChromatinAssay(counts = exp3_ATAC_mtx, sep = c(":", "-"), fragments = all_fragments[[3]])
DefaultAssay(exp3_seurat) = "WNN"
exp3_seurat <- RunTFIDF(exp3_seurat)

exp4_seurat <- CreateSeuratObject(counts = exp4_RNA_mtx, project = "exp043_clean")
exp4_seurat <- SCTransform(exp4_seurat, vst.flavor = "v2", method = "glmGamPoi")
exp4_seurat[["WNN"]] <- CreateChromatinAssay(counts = exp4_ATAC_mtx, sep = c(":", "-"), fragments = all_fragments[[4]])
DefaultAssay(exp4_seurat) = "WNN"
exp4_seurat <- RunTFIDF(exp4_seurat)

exp5_seurat <- CreateSeuratObject(counts = exp5_RNA_mtx, project = "exp024_clean")
exp5_seurat <- SCTransform(exp5_seurat, vst.flavor = "v2", method = "glmGamPoi")
exp5_seurat[["WNN"]] <- CreateChromatinAssay(counts = exp5_ATAC_mtx, sep = c(":", "-"), fragments = all_fragments[[5]])
DefaultAssay(exp5_seurat) = "WNN"
exp5_seurat <- RunTFIDF(exp5_seurat)

exp6_seurat <- CreateSeuratObject(counts = exp6_RNA_mtx, project = "exp025_clean")
exp6_seurat <- SCTransform(exp6_seurat, vst.flavor = "v2", method = "glmGamPoi")
exp6_seurat[["WNN"]] <- CreateChromatinAssay(counts = exp6_ATAC_mtx, sep = c(":", "-"), fragments = all_fragments[[6]])
DefaultAssay(exp6_seurat) = "WNN"
exp6_seurat <- RunTFIDF(exp6_seurat)

exp7_seurat <- CreateSeuratObject(counts = exp7_RNA_mtx, project = "exp027_clean")
exp7_seurat <- SCTransform(exp7_seurat, vst.flavor = "v2", method = "glmGamPoi")
exp7_seurat[["WNN"]] <- CreateChromatinAssay(counts = exp7_ATAC_mtx, sep = c(":", "-"), fragments = all_fragments[[7]])
DefaultAssay(exp7_seurat) = "WNN"
exp7_seurat <- RunTFIDF(exp7_seurat)

# merge all objects - RNA
ee.list <- list(exp1_seurat@assays[["RNA"]], exp2_seurat@assays[["RNA"]], exp3_seurat@assays[["RNA"]], exp4_seurat@assays[["RNA"]], 
                exp5_seurat@assays[["RNA"]], exp6_seurat@assays[["RNA"]], exp7_seurat@assays[["RNA"]])

exp1_seurat@meta.data$exp <- "exp031"
exp2_seurat@meta.data$exp <- "exp032"
exp3_seurat@meta.data$exp <- "exp042"
exp4_seurat@meta.data$exp <- "exp043"
exp5_seurat@meta.data$exp <- "exp024"
exp6_seurat@meta.data$exp <- "exp025"
exp7_seurat@meta.data$exp <- "exp027"

gcdata <- merge(exp1_seurat, y=c(exp2_seurat, exp3_seurat, exp4_seurat, exp5_seurat, exp6_seurat, exp7_seurat), merge.data = T)

VariableFeatures(gcdata[["SCT"]]) <- rownames(gcdata[["SCT"]]@scale.data)
DefaultAssay(gcdata) = "SCT"

gcdata <- RunPCA(gcdata, seed.use = 42, features = VariableFeatures(object = gcdata), npcs = 100)
gcdata <- FindNeighbors(gcdata, dims = 1:50)
gcdata <- FindClusters(gcdata, resolution = 3, random.seed = 0)
gcdata <- RunUMAP(gcdata, seed.use = 42, dims = 1:50, return.model=TRUE)

# logNorm the RNA counts - useful for plot features
DefaultAssay(gcdata) = "RNA"
gcdata = NormalizeData(gcdata)

# merge all objects - ATAC
DefaultAssay(gcdata) = "WNN"

gcdata <- FindTopFeatures(gcdata, min.cutoff = 'q0')
gcdata <- RunSVD(gcdata)
gcdata <- RunUMAP(gcdata, reduction = 'lsi', dims = 2:50, reduction.name = "umapATAC")
gcdata <- FindNeighbors(gcdata, reduction = 'lsi', dims = 2:50)
gcdata <- FindClusters(gcdata, verbose = T, algorithm = 3, resolution = 3)

# cluster barcodes based on both modalities
gcdata <- FindMultiModalNeighbors(gcdata, reduction.list = list("pca", "lsi"), dims.list = list(1:50, 2:50))
gcdata <- RunUMAP(gcdata, nn.name = "weighted.nn", reduction.name = "umapWNN")
gcdata <- FindClusters(gcdata, graph.name = "wsnn", algorithm = 3, verbose = T, resolution = 4)

gcdata <- PrepSCTFindMarkers(gcdata)

# plotting
p1 <- DimPlot(gcdata, reduction = "umap", group.by = "wsnn_res.4", label = TRUE, label.size = 4)
p2 <- DimPlot(gcdata, reduction = "umapATAC", group.by = "wsnn_res.4", label = TRUE, label.size = 4)
p3 <- DimPlot(gcdata, reduction = "umapWNN", group.by = "wsnn_res.4", label = TRUE, label.size = 4)

pdf(file=paste0(plot_dir_w_date, "/exp031_exp032_exp042_exp043_exp024_exp025_exp027_SCT_clean.", annotation, ".umap_WNN.pdf"), width=24, height = 8)
p1 + p2 + p3 & NoLegend() & theme(plot.title = element_text(hjust = 0.5))
dev.off()

pdf(file=paste0(plot_dir_w_date, "/exp031_exp032_exp042_exp043_exp024_exp025_exp027_SCT_clean.", annotation, ".umap_split.WNN.pdf"), width=24, height = 8)
p1 = DimPlot(gcdata, reduction = "umapWNN", label = TRUE, label.size = 4) 
p2 = DimPlot(gcdata, reduction = "umapWNN", group.by = "exp")
p1 & NoLegend() & theme(plot.title = element_text(hjust = 0.5)) | p2  & theme(plot.title = element_text(hjust = 0.5))
DimPlot(gcdata, reduction = "umapWNN", split.by = "exp") & NoLegend() & theme(plot.title = element_text(hjust = 0.5))
dev.off()

DefaultAssay(gcdata) = "SCT"

pdf(file=paste0("plots/", current_date, "/exp031_exp032_exp042_exp043_exp024_exp025_exp027_SCT.", annotation, ".key_lineage_markers.WNN_clean.pdf"), width=8, height = 8)
FeaturePlot(gcdata, order = T, features = c("med-1"), reduction = "umapWNN") & scale_colour_gradientn(colours = c("grey90", "cyan", "deepskyblue", "forestgreen", "darkorange", "darkred"))
FeaturePlot(gcdata, order = T, features = c("tbx-40"), reduction = "umapWNN") & scale_colour_gradientn(colours = c("grey90", "cyan", "deepskyblue", "forestgreen", "darkorange", "darkred"))
FeaturePlot(gcdata, order = T, features = c("tbx-35"), reduction = "umapWNN") & scale_colour_gradientn(colours = c("grey90", "cyan", "deepskyblue", "forestgreen", "darkorange", "darkred"))
FeaturePlot(gcdata, order = T, features = c("tbx-33"), reduction = "umapWNN") & scale_colour_gradientn(colours = c("grey90", "cyan", "deepskyblue", "forestgreen", "darkorange", "darkred"))
FeaturePlot(gcdata, order = T, features = c("end-3"), reduction = "umapWNN") & scale_colour_gradientn(colours = c("grey90", "cyan", "deepskyblue", "forestgreen", "darkorange", "darkred"))
FeaturePlot(gcdata, order = T, features = c("hnd-1"), reduction = "umapWNN") & scale_colour_gradientn(colours = c("grey90", "cyan", "deepskyblue", "forestgreen", "darkorange", "darkred"))
FeaturePlot(gcdata, order = T, features = c("pes-10"), reduction = "umapWNN") & scale_colour_gradientn(colours = c("grey90", "cyan", "deepskyblue", "forestgreen", "darkorange", "darkred"))
FeaturePlot(gcdata, order = T, features = c("vet-2"), reduction = "umapWNN") & scale_colour_gradientn(colours = c("grey90", "cyan", "deepskyblue", "forestgreen", "darkorange", "darkred"))
FeaturePlot(gcdata, order = T, features = c("ceh-51"), reduction = "umapWNN") & scale_colour_gradientn(colours = c("grey90", "cyan", "deepskyblue", "forestgreen", "darkorange", "darkred"))
FeaturePlot(gcdata, order = T, features = c("oma-1"), reduction = "umapWNN") & scale_colour_gradientn(colours = c("grey90", "cyan", "deepskyblue", "forestgreen", "darkorange", "darkred"))
FeaturePlot(gcdata, order = T, features = c("oma-2"), reduction = "umapWNN") & scale_colour_gradientn(colours = c("grey90", "cyan", "deepskyblue", "forestgreen", "darkorange", "darkred"))
FeaturePlot(gcdata, order = T, features = c("lsl-1"), reduction = "umapWNN") & scale_colour_gradientn(colours = c("grey90", "cyan", "deepskyblue", "forestgreen", "darkorange", "darkred"))
FeaturePlot(gcdata, order = T, features = c("skr-7"), reduction = "umapWNN") & scale_colour_gradientn(colours = c("grey90", "cyan", "deepskyblue", "forestgreen", "darkorange", "darkred"))
FeaturePlot(gcdata, order = T, features = c("ccch-2"), reduction = "umapWNN") & scale_colour_gradientn(colours = c("grey90", "cyan", "deepskyblue", "forestgreen", "darkorange", "darkred"))
FeaturePlot(gcdata, order = T, features = c("ref-1"), reduction = "umapWNN") & scale_colour_gradientn(colours = c("grey90", "cyan", "deepskyblue", "forestgreen", "darkorange", "darkred"))
FeaturePlot(gcdata, order = T, features = c("cpg-3"), reduction = "umapWNN") & scale_colour_gradientn(colours = c("grey90", "cyan", "deepskyblue", "forestgreen", "darkorange", "darkred"))
dev.off()

# save RDS
saveRDS(gcdata, file = paste0("seurat_objects/combined_exp031_exp032_exp042_exp043_exp024_exp025_exp027_SCT.", annotation, ".WNN_clean.rds"))
write.table(gcdata[[]], file = paste0("seurat_objects/combined_exp031_exp032_exp042_exp043_exp024_exp025_exp027_SCT.", annotation, ".WNN_clean.table.txt"), quote = F, sep = "\t", row.names = T, col.names = T)
```
