## Supplementary material for "A lineage-resolved multimodal single-cell atlas reveals the genomic dynamics of early *C. elegans* development": Code and external data: cell_cycle_random_forest.R

```
library(Seurat)
library(Signac)
library(Hmisc)
library(RColorBrewer)
library(presto)
library(caret)

args = commandArgs(trailingOnly=TRUE)

combined_seurat_rds = args[1]

combined_seurat = readRDS(combined_seurat_rds)

# define train set: get sample of 500 early and 500 late cells from the AB/E/MS/C lineages
train_early = c("earlyABxxx", "earlyEMS", "earlyCx", "earlyMSxx")
train_late = c("ABala", "ABalp", "ABpxa", "EMS", "Ca", "Cp", "MSxa", "MSxp")

test_early = c("earlyABxxxx", "earlyEandMS", "earlyCxx", "earlyMSx")
test_late = c("ABaraa", "ABarpa", "ABpxap", "E", "MS", "Cxa", "Cxp", "MSa", "MSp")

train_early_bc = colnames(combined_seurat[,combined_seurat$cell_type %in% train_early])
train_late_bc = colnames(combined_seurat[,combined_seurat$cell_type %in% train_late])
test_early_bc = colnames(combined_seurat[,combined_seurat$cell_type %in% test_early])
test_late_bc = colnames(combined_seurat[,combined_seurat$cell_type %in% test_late])

set.seed(10)
train_early_bc_subs = sample(train_early_bc, size = 200, replace = F)
train_late_bc_subs = sample(train_late_bc, size = 200, replace = F)

train_set = t(combined_seurat@assays$SCT@scale.data[,colnames(combined_seurat@assays$SCT@scale.data) %in% train_early_bc_subs])
train_set = rbind(train_set, t(combined_seurat@assays$SCT@scale.data[,colnames(combined_seurat@assays$SCT@scale.data) %in% train_late_bc_subs]))
train_set = as.data.frame(train_set)
train_set$early_late = c(rep("early", length(train_early_bc_subs)), rep("late", length(train_late_bc_subs)))

test_set = t(combined_seurat@assays$SCT@scale.data[,colnames(combined_seurat@assays$SCT@scale.data) %in% test_early_bc])
test_set = rbind(test_set, t(combined_seurat@assays$SCT@scale.data[,colnames(combined_seurat@assays$SCT@scale.data) %in% test_late_bc]))
test_set = as.data.frame(test_set)
test_set$early_late = c(rep("early", length(test_early_bc)), rep("late", length(test_late_bc)))

# training
#set.seed(6)

# randomForest training
#train_rf <- train(early_late ~ .,
#                  method = "rf",
#                  tuneGrid = data.frame(mtry = seq(10, 50, 5)), ntree=100,
#                  data = train_set)

#rf_preds <- predict(train_rf, test_set)
#mean(rf_preds == test_set$early_late)

# predict on entire set
#rf_preds_whole <- predict(train_rf, t(combined_seurat@assays$SCT@scale.data))
#combined_seurat$cell_cycle_rf = rf_preds_whole

#p1 = DimPlot(combined_seurat, reduction = "umapWNN", group.by = "cell_cycle_rf", order = T)


# train using RF (mtry=50) on whole dataset
train_set_whole = t(combined_seurat@assays$SCT@scale.data[,colnames(combined_seurat@assays$SCT@scale.data) %in% train_early_bc])
train_set_whole = rbind(train_set_whole, t(combined_seurat@assays$SCT@scale.data[,colnames(combined_seurat@assays$SCT@scale.data) %in% train_late_bc]))
train_set_whole = as.data.frame(train_set_whole)
train_set_whole$early_late = c(rep("early", length(train_early_bc)), rep("late", length(train_late_bc)))

test_set_whole = t(combined_seurat@assays$SCT@scale.data[,colnames(combined_seurat@assays$SCT@scale.data) %in% test_early_bc])
test_set_whole = rbind(test_set_whole, t(combined_seurat@assays$SCT@scale.data[,colnames(combined_seurat@assays$SCT@scale.data) %in% test_late_bc]))
test_set_whole = as.data.frame(test_set_whole)
test_set_whole$early_late = c(rep("early", length(test_early_bc)), rep("late", length(test_late_bc)))

set.seed(6)
train_rf_all <- train(early_late ~ .,
                      method = "rf",
                      tuneGrid = data.frame(mtry = 50), ntree=100,
                      data = train_set_whole)

# accuracy
rf_preds <- predict(train_rf_all, test_set_whole)
mean(rf_preds == test_set$early_late)

# prediction
rf_preds_all <- predict(train_rf_all, t(combined_seurat@assays$SCT@scale.data))

combined_seurat$cell_cycle_rf = rf_preds_all

prediction_df = data.frame("cell_cycle"=rf_preds_all, row.names = colnames(combined_seurat))
write.table(rf_preds_all, file = "cell_cycle_prediction/cell_cycle_RF.txt", quote = F, sep = "\t", row.names = T, col.names = F)
p1 = DimPlot(combined_seurat, reduction = "umapWNN", group.by = "cell_cycle_rf", order = T)

dir.create("cell_cycle_prediction", recursive = T)

pdf("cell_cycle_prediction/cell_cycle_RF.200cells.pdf", width = 6, height = 6, useDingbats = F)
p1
dev.off()

saveRDS(combined_seurat, file = paste0("seurat_objects/wt_all.all_samples.all_cells.WS285_extended_no_ovlp.final_peakset_IDR_cell_cycle.rds"))
```
