## Supplementary material for "A lineage-resolved multimodal single-cell atlas reveals the genomic dynamics of early *C. elegans* development": Code and external data: cellranger_arc_launcher.sh

```
#!/bin/bash
#
#SBATCH --job-name=cellranger_arc
#SBATCH --output=slurm_out/cellranger_arc.%N.%j.out
#SBATCH --error=slurm_err/cellranger_arc.%N.%j.err
#SBATCH --ntasks=1
#SBATCH -N 1
#SBATCH -n 12
#SBATCH --mem-per-cpu=8000
#SBATCH --mail-type=END,FAIL
#SBATCH --mail-user=

exp_id=$1

cellranger-arc count --id=$exp_id --reference=WS285_cellranger_arc --libraries=cellranger_arc_library_files/$exp_id.cr_arc_library_file.csv --localcores=12

mv $exp_id data/sc/cellranger_arc/
```
