## Supplementary material for "A lineage-resolved multimodal single-cell atlas reveals the genomic dynamics of early *C. elegans* development": Code and external data: chipseq_alignment.sh

```
#!/bin/bash
#
#SBATCH --job-name=chip_alignment
#SBATCH --output=slurm_out/chip_alignment.%N.%j.out
#SBATCH --error=slurm_err/chip_alignment.%N.%j.err
#SBATCH --ntasks=1
#SBATCH -N 1
#SBATCH -n 10
#SBATCH --mem-per-cpu=10000
#SBATCH --mail-type=END,FAIL
#SBATCH --mail-user=

sample=$1

read1="data/chipseq/fastq/"$sample"_R1_001.fastq.gz"
read2="data/chipseq/fastq/"$sample"_R2_001.fastq.gz"

outdir_trimmed=data/chipseq/fastq_trimmed
outdir_alignment=data/chipseq/alignment
outdir_bw=data/chipseq/bw

bwa_index=bwa_idx/elegans
genome_chr=species/elegans/genome/elegans.fa.fai
blacklist=data/external_data/ce11-blacklist.bed

mkdir $outdir_alignment
mkdir $outdir_bw

if [ -f $read2 ]
then
    trim_galore --gzip --fastqc --paired --trim-n -j 10 -o $outdir_trimmed $read1 $read2
    bwa mem -t 10 $bwa_index $outdir_trimmed"/"$sample"_R1_001_val_1.fq.gz" $outdir_trimmed"/"$sample"_R2_001_val_2.fq.gz" | samtools view -@ 10 -q 10 -bT $genome_chr - | samtools sort -@ 10 - > $outdir_alignment/$sample.mapq10.bam
    samtools index $outdir_alignment/$sample.mapq10.bam
    bamCoverage --bam $outdir_alignment/$sample.mapq10.bam -o $outdir_bw/$sample.mapq10.bw --binSize 1 --normalizeUsing CPM --effectiveGenomeSize 100286401 -bl $blacklist --extendReads -p 6
else
    trim_galore --gzip --fastqc --trim-n -j 10 -o $outdir_trimmed $read1
    bwa mem -t 10 $bwa_index $outdir_trimmed"/"$sample"_R1_001_trimmed.fq.gz" | samtools view -@ 10 -q 10 -bT $genome_chr - | samtools sort -@ 10 - > $outdir_alignment/$sample.mapq10.bam
    samtools index $outdir_alignment/$sample.mapq10.bam
    bamCoverage --bam $outdir_alignment/$sample.mapq10.bam -o $outdir_bw/$sample.mapq10.bw --binSize 1 --normalizeUsing CPM --effectiveGenomeSize 100286401 -bl $blacklist -p 6 
fi
```
