## Supplementary material for "A lineage-resolved multimodal single-cell atlas reveals the genomic dynamics of early *C. elegans* development": Code and external data: chipseq_peaks.sh

```
#!/bin/bash
#
#SBATCH --job-name=chip_alignment
#SBATCH --output=slurm_out/chip_alignment.%N.%j.out
#SBATCH --error=slurm_err/chip_alignment.%N.%j.err
#SBATCH --ntasks=1
#SBATCH -N 1
#SBATCH -n 1
#SBATCH --mem-per-cpu=10000
#SBATCH --mail-type=END,FAIL
#SBATCH --mail-user=

sample=$1
model=$2

outdir_alignment=data/chipseq/alignment
outdir_macs=data/chipseq/macs

genome_chr=species/elegans/genome/elegans.chrom.sizes.txt

mkdir -p $outdir_macs

if [ -z "$model" ]
then
    macs2 callpeak --bdg --call-summits --SPMR --gsize ce --keep-dup all --nomodel -n $sample --outdir $outdir_macs/$sample.nomodel -t $outdir_alignment/$sample.mapq10.bam
    sort -k 1,1 -k2,2n $outdir_macs"/"$sample".nomodel/"$sample"_treat_pileup.bdg" > $outdir_macs"/"$sample".nomodel/"$sample"_treat_pileup_sorted.bdg"
    bedGraphToBigWig $outdir_macs"/"$sample".nomodel/"$sample"_treat_pileup_sorted.bdg" $genome_chr $outdir_macs"/"$sample".nomodel/"$sample"_treat_pileup_sorted.bw"
else
    macs2 callpeak --bdg --call-summits --SPMR --gsize ce --keep-dup all -n $sample --outdir $outdir_macs/$sample.model -t $outdir_alignment/$sample.mapq10.bam
    sort -k 1,1 -k2,2n $outdir_macs"/"$sample".model/"$sample"_treat_pileup.bdg" > $outdir_macs"/"$sample".model/"$sample"_treat_pileup_sorted.bdg"
    bedGraphToBigWig $outdir_macs"/"$sample".model/"$sample"_treat_pileup_sorted.bdg" $genome_chr $outdir_macs"/"$sample".model/"$sample"_treat_pileup_sorted.bw"
fi
```
