## Supplementary material for "A lineage-resolved multimodal single-cell atlas reveals the genomic dynamics of early *C. elegans* development": Code and external data: combine_bw_tracks_stage.20240916.sh

```
#!/bin/bash
#
#SBATCH --job-name=bw_merge
#SBATCH --output=slurm_out/bw_merge.%N.%j.out
#SBATCH --error=slurm_err/bw_merge.%N.%j.err
#SBATCH --ntasks=1
#SBATCH -N 1
#SBATCH -n 12
#SBATCH --mem-per-cpu=8000
#SBATCH --mail-type=END,FAIL
#SBATCH --mail-user=

mkdir cluster_specific_peaks/round_2_all_samples/WS285_extended_no_ovlp/embryo_stage_tracks

cp cluster_specific_peaks/round_2_all_samples/WS285_extended_no_ovlp/cell_type/bw/twocell.cpm.bw cluster_specific_peaks/round_2_all_samples/WS285_extended_no_ovlp/embryo_stage_tracks/2cell.cpm.bw
cp cluster_specific_peaks/round_2_all_samples/WS285_extended_no_ovlp/cell_type/bw/Z2Z3.cpm.bw cluster_specific_peaks/round_2_all_samples/WS285_extended_no_ovlp/embryo_stage_tracks/Z2Z3.cpm.bw

bigwigAverage --bigwigs \
              cluster_specific_peaks/round_2_all_samples/WS285_extended_no_ovlp/cell_type/bw/P2.cpm.bw \
              cluster_specific_peaks/round_2_all_samples/WS285_extended_no_ovlp/cell_type/bw/P3.cpm.bw \
              cluster_specific_peaks/round_2_all_samples/WS285_extended_no_ovlp/cell_type/bw/P4.cpm.bw \
              -bs 1 -p 12 -o cluster_specific_peaks/round_2_all_samples/WS285_extended_no_ovlp/embryo_stage_tracks/Pcell.cpm.bw

bigwigAverage --bigwigs \
	      cluster_specific_peaks/round_2_all_samples/WS285_extended_no_ovlp/cell_type/bw/ABx.cpm.bw \
	      cluster_specific_peaks/round_2_all_samples/WS285_extended_no_ovlp/cell_type/bw/earlyEMS.cpm.bw \
	      cluster_specific_peaks/round_2_all_samples/WS285_extended_no_ovlp/cell_type/bw/EMS.cpm.bw \
	      -bs 1 -p 12 -o cluster_specific_peaks/round_2_all_samples/WS285_extended_no_ovlp/embryo_stage_tracks/4cell.cpm.bw

bigwigAverage --bigwigs \
	      cluster_specific_peaks/round_2_all_samples/WS285_extended_no_ovlp/cell_type/bw/ABax.cpm.bw \
	      cluster_specific_peaks/round_2_all_samples/WS285_extended_no_ovlp/cell_type/bw/ABpx.cpm.bw \
	      cluster_specific_peaks/round_2_all_samples/WS285_extended_no_ovlp/cell_type/bw/earlyEandMS.cpm.bw \
	      cluster_specific_peaks/round_2_all_samples/WS285_extended_no_ovlp/cell_type/bw/E.cpm.bw \
	      cluster_specific_peaks/round_2_all_samples/WS285_extended_no_ovlp/cell_type/bw/MS.cpm.bw \
	      cluster_specific_peaks/round_2_all_samples/WS285_extended_no_ovlp/cell_type/bw/earlyC.cpm.bw \
	      cluster_specific_peaks/round_2_all_samples/WS285_extended_no_ovlp/cell_type/bw/C.cpm.bw \
	      -bs 1 -p 12 -o cluster_specific_peaks/round_2_all_samples/WS285_extended_no_ovlp/embryo_stage_tracks/8cell.cpm.bw

bigwigAverage --bigwigs \
	      cluster_specific_peaks/round_2_all_samples/WS285_extended_no_ovlp/cell_type/bw/earlyABxxx.cpm.bw \
	      cluster_specific_peaks/round_2_all_samples/WS285_extended_no_ovlp/cell_type/bw/ABxxx.cpm.bw \
	      cluster_specific_peaks/round_2_all_samples/WS285_extended_no_ovlp/cell_type/bw/ABala.cpm.bw \
	      cluster_specific_peaks/round_2_all_samples/WS285_extended_no_ovlp/cell_type/bw/ABara.cpm.bw \
	      cluster_specific_peaks/round_2_all_samples/WS285_extended_no_ovlp/cell_type/bw/ABalp.cpm.bw \
	      cluster_specific_peaks/round_2_all_samples/WS285_extended_no_ovlp/cell_type/bw/ABarp.cpm.bw \
	      cluster_specific_peaks/round_2_all_samples/WS285_extended_no_ovlp/cell_type/bw/ABpxa.cpm.bw \
	      cluster_specific_peaks/round_2_all_samples/WS285_extended_no_ovlp/cell_type/bw/ABpxp.cpm.bw \
	      cluster_specific_peaks/round_2_all_samples/WS285_extended_no_ovlp/cell_type/bw/earlyMSx.cpm.bw \
	      cluster_specific_peaks/round_2_all_samples/WS285_extended_no_ovlp/cell_type/bw/MSx.cpm.bw \
	      cluster_specific_peaks/round_2_all_samples/WS285_extended_no_ovlp/cell_type/bw/MSa.cpm.bw \
	      cluster_specific_peaks/round_2_all_samples/WS285_extended_no_ovlp/cell_type/bw/MSp.cpm.bw \
	      cluster_specific_peaks/round_2_all_samples/WS285_extended_no_ovlp/cell_type/bw/earlyEx.cpm.bw \
	      cluster_specific_peaks/round_2_all_samples/WS285_extended_no_ovlp/cell_type/bw/earlyCx.cpm.bw \
	      -bs 1 -p 12 -o cluster_specific_peaks/round_2_all_samples/WS285_extended_no_ovlp/embryo_stage_tracks/15cell.cpm.bw

bigwigAverage --bigwigs \
	      cluster_specific_peaks/round_2_all_samples/WS285_extended_no_ovlp/cell_type/bw/earlyABxxxx.cpm.bw \
	      cluster_specific_peaks/round_2_all_samples/WS285_extended_no_ovlp/cell_type/bw/ABxxxx.cpm.bw \
	      cluster_specific_peaks/round_2_all_samples/WS285_extended_no_ovlp/cell_type/bw/ABalaa.cpm.bw \
	      cluster_specific_peaks/round_2_all_samples/WS285_extended_no_ovlp/cell_type/bw/ABalap.cpm.bw \
	      cluster_specific_peaks/round_2_all_samples/WS285_extended_no_ovlp/cell_type/bw/ABalpa.cpm.bw \
	      cluster_specific_peaks/round_2_all_samples/WS285_extended_no_ovlp/cell_type/bw/ABalpp.cpm.bw \
	      cluster_specific_peaks/round_2_all_samples/WS285_extended_no_ovlp/cell_type/bw/ABaraa.cpm.bw \
	      cluster_specific_peaks/round_2_all_samples/WS285_extended_no_ovlp/cell_type/bw/ABarap.cpm.bw \
	      cluster_specific_peaks/round_2_all_samples/WS285_extended_no_ovlp/cell_type/bw/ABarpa.cpm.bw \
	      cluster_specific_peaks/round_2_all_samples/WS285_extended_no_ovlp/cell_type/bw/ABarpp.cpm.bw \
	      cluster_specific_peaks/round_2_all_samples/WS285_extended_no_ovlp/cell_type/bw/ABpxaa.cpm.bw \
	      cluster_specific_peaks/round_2_all_samples/WS285_extended_no_ovlp/cell_type/bw/ABpxap.cpm.bw \
	      cluster_specific_peaks/round_2_all_samples/WS285_extended_no_ovlp/cell_type/bw/ABpxpa.cpm.bw \
	      cluster_specific_peaks/round_2_all_samples/WS285_extended_no_ovlp/cell_type/bw/ABpxpp.cpm.bw \
	      cluster_specific_peaks/round_2_all_samples/WS285_extended_no_ovlp/cell_type/bw/earlyMSxx.cpm.bw \
	      cluster_specific_peaks/round_2_all_samples/WS285_extended_no_ovlp/cell_type/bw/Ex.cpm.bw \
	      cluster_specific_peaks/round_2_all_samples/WS285_extended_no_ovlp/cell_type/bw/Ca.cpm.bw \
	      cluster_specific_peaks/round_2_all_samples/WS285_extended_no_ovlp/cell_type/bw/Cp.cpm.bw \
	      cluster_specific_peaks/round_2_all_samples/WS285_extended_no_ovlp/cell_type/bw/D.cpm.bw \
	      -bs 1 -p 12 -o cluster_specific_peaks/round_2_all_samples/WS285_extended_no_ovlp/embryo_stage_tracks/26cell.cpm.bw

bigwigAverage --bigwigs \
	      cluster_specific_peaks/round_2_all_samples/WS285_extended_no_ovlp/cell_type/bw/ABxxxxx[0-99]*.cpm.bw \
	      cluster_specific_peaks/round_2_all_samples/WS285_extended_no_ovlp/cell_type/bw/earlyExx.cpm.bw \
	      cluster_specific_peaks/round_2_all_samples/WS285_extended_no_ovlp/cell_type/bw/MSxa.cpm.bw \
	      cluster_specific_peaks/round_2_all_samples/WS285_extended_no_ovlp/cell_type/bw/MSxp.cpm.bw \
	      cluster_specific_peaks/round_2_all_samples/WS285_extended_no_ovlp/cell_type/bw/earlyCxx.cpm.bw \
	      cluster_specific_peaks/round_2_all_samples/WS285_extended_no_ovlp/cell_type/bw/Cxa.cpm.bw \
	      cluster_specific_peaks/round_2_all_samples/WS285_extended_no_ovlp/cell_type/bw/Cxp.cpm.bw \
	      -bs 1 -p 12 -o cluster_specific_peaks/round_2_all_samples/WS285_extended_no_ovlp/embryo_stage_tracks/46cell.cpm.bw 

bigwigAverage --bigwigs \
              cluster_specific_peaks/round_2_all_samples/WS285_extended_no_ovlp/cell_type/bw/ABxxxxxx*.cpm.bw \
              cluster_specific_peaks/round_2_all_samples/WS285_extended_no_ovlp/cell_type/bw/Exx.cpm.bw \
              cluster_specific_peaks/round_2_all_samples/WS285_extended_no_ovlp/cell_type/bw/Exxx.cpm.bw \
              cluster_specific_peaks/round_2_all_samples/WS285_extended_no_ovlp/cell_type/bw/MSxxx*.cpm.bw \
              cluster_specific_peaks/round_2_all_samples/WS285_extended_no_ovlp/cell_type/bw/Cxxx*.cpm.bw \
              cluster_specific_peaks/round_2_all_samples/WS285_extended_no_ovlp/cell_type/bw/Dx.cpm.bw \
              cluster_specific_peaks/round_2_all_samples/WS285_extended_no_ovlp/cell_type/bw/DxxCxxx*.cpm.bw \
              -bs 1 -p 12 -o cluster_specific_peaks/round_2_all_samples/WS285_extended_no_ovlp/embryo_stage_tracks/87morecell.cpm.bw

bigwigAverage --bigwigs \
              cluster_specific_peaks/round_2_all_samples/WS285_extended_no_ovlp/cell_type/bw/unassigned*.cpm.bw \
              -bs 1 -p 12 -o cluster_specific_peaks/round_2_all_samples/WS285_extended_no_ovlp/embryo_stage_tracks/unassigned.cpm.bw
```
