## Supplementary material for "A lineage-resolved multimodal single-cell atlas reveals the genomic dynamics of early *C. elegans* development": Code and external data: coveragebed_launcher.20231101.sh

```
#!/bin/bash
#
#SBATCH --job-name=coveragebed
#SBATCH --output=slurm_out/coveragebed.%N.%j.out
#SBATCH --error=slurm_err/coveragebed.%N.%j.err
#SBATCH --ntasks=1
#SBATCH -N 1
#SBATCH -n 1
#SBATCH --mem=64G
#SBATCH --mail-type=END,FAIL
#SBATCH --mail-user=

bam_input=$1
bed_input=$2
bed_output=$3
remove_blacklist=$4

chr_size=species/elegans/genome/elegans.chrom.sizes.txt
blacklist=data/external_data/ce11-blacklist.bed

if [ $remove_blacklist = "no" ]
then
    sort -k 1,1 -k2,2n $bed_input | coverageBed -a stdin -b $bam_input -counts -s -g $chr_size -sorted > $bed_output
else
    intersectBed -a $bed_input -b $blacklist -v | sort -k 1,1 -k2,2n | coverageBed -a stdin -b $bam_input -counts -s -g $chr_size -sorted > $bed_output
fi
```
