## Supplementary material for "A lineage-resolved multimodal single-cell atlas reveals the genomic dynamics of early *C. elegans* development": Code and external data: dedup_launcher.clusters.20240226.sh

```
#!/bin/bash
#
#SBATCH --job-name=dedup_clusters
#SBATCH --output=slurm_out/dedup_clusters.%N.%j.out
#SBATCH --error=slurm_err/dedup_clusters.%N.%j.err
#SBATCH --ntasks=1
#SBATCH -N 1
#SBATCH -n 8
#SBATCH --mem-per-cpu=4000
#SBATCH --mail-type=END,FAIL
#SBATCH --mail-user=

sample_name=$1
alignment_path=$2

mkdir -p $alignment_path/dedup_err
mkdir -p $alignment_path/dedup_log
mkdir -p $alignment_path/bw
mkdir -p $alignment_path/bw/unique
mkdir -p $alignment_path/dedup_bam/

blacklist=data/external_data/ce11-blacklist.bed

samtools index $alignment_path/bam/$sample_name.bam
umi_tools dedup --stdin=$alignment_path/bam/$sample_name.bam --log=$alignment_path/dedup_log/$sample_name.dedup.log --error=$alignment_path/dedup_err/$sample_name.dedup.err --extract-umi-method=tag --umi-tag=UR --cell-tag=CR --per-cell > $alignment_path/dedup_bam/$sample_name.dedup.bam
samtools index $alignment_path/dedup_bam/$sample_name.dedup.bam
bamCoverage -b $alignment_path/dedup_bam/$sample_name.dedup.bam -o $alignment_path/bw/$sample_name.dedup.rev.bw --filterRNAstrand forward -bs 1 -p 8 --effectiveGenomeSize 100286401 --normalizeUsing CPM --blackListFileName=$blacklist
bamCoverage -b $alignment_path/dedup_bam/$sample_name.dedup.bam -o $alignment_path/bw/unique/$sample_name.unique.dedup.rev.bw --filterRNAstrand forward -bs 1 -p 8 --effectiveGenomeSize 100286401 --normalizeUsing CPM --samFlagExclude 256 --blackListFileName=$blacklist
bamCoverage -b $alignment_path/dedup_bam/$sample_name.dedup.bam -o $alignment_path/bw/$sample_name.dedup.for.bw --filterRNAstrand reverse -bs 1 -p 8 --effectiveGenomeSize 100286401 --normalizeUsing CPM --blackListFileName=$blacklist
bamCoverage -b $alignment_path/dedup_bam/$sample_name.dedup.bam -o $alignment_path/bw/unique/$sample_name.unique.dedup.for.bw --filterRNAstrand reverse -bs 1 -p 8 --effectiveGenomeSize 100286401 --normalizeUsing CPM --samFlagExclude 256 --blackListFileName=$blacklist
```
