## Supplementary material for "A lineage-resolved multimodal single-cell atlas reveals the genomic dynamics of early *C. elegans* development": Code and external data: DESeq_embryo_expression.20241202.R

```
library('EnhancedVolcano')
library(rhdf5)
library("DESeq2")
library("GenomicAlignments")
library("tximport")
library("readr")
library("tximportData")
library("pheatmap")
library("RColorBrewer")
library('Hmisc')

source("data/external_data/wormcat_function.R")

current_date= paste(unlist(strsplit(as.character(Sys.Date()), "-")), collapse="")
plot_dir_w_date = paste0("paper_figures/", current_date)
dir.create(plot_dir_w_date, recursive=TRUE)

# expression correlation between all samples
#samples_list = t(sapply(strsplit(list.files("data/rnaseq/fastq/"), "[_]"), '[', c(1:3)))
samples_list_cat = c("rnaseq_wt_EE_rep2", "rnaseq_wt_EE_rep3", "rnaseq_wt_EE_rep4", "rnaseq_wt_EE_rep5",
                     "rnaseq_ceh49_EE_rep1", "rnaseq_ceh49_EE_rep2", "rnaseq_ceh49_EE_rep3", "rnaseq_ceh49_EE_rep4",
                     "rnaseq_6xCUT_EE_rep1", "rnaseq_6xCUT_EE_rep2", "rnaseq_6xCUT_EE_rep3",
                     "rnaseq_7xCUT_EE_rep1", "rnaseq_7xCUT_EE_rep2", "rnaseq_7xCUT_EE_rep3")
#samples_list_cat = unique(paste(samples_list[,1], samples_list[,2], samples_list[,3], sep="_"))
sample_ID_list=data.frame("sample" = samples_list_cat, "strain" = unlist(lapply(strsplit(samples_list_cat, "[_]"), '[[', 2)), stringsAsFactors = T)

sample_ID_list$exp_files <- file.path("data/rnaseq/kallisto/", sample_ID_list$sample, "abundance.genes.txt")
sample_ID_list$replicates <- c(2,3,4,5,1,2,3,4,1,2,3,1,2,3)

remove(gene_exp_all_samples)
for (sample_rnaseq in c(1:dim(sample_ID_list)[1])){
  if (exists("gene_exp_all_samples") == FALSE){
    gene_exp_all_samples = read.table(sample_ID_list[sample_rnaseq,3], row.names = 1)
  }
  else{
    k = read.table(sample_ID_list[sample_rnaseq,3])
    gene_exp_all_samples = cbind(gene_exp_all_samples, k$V2)
  }
}

names(gene_exp_all_samples) = sample_ID_list$sample

k = pheatmap(cor(gene_exp_all_samples[,], method = "pearson"))

pdf(file=paste0(plot_dir_w_date, "/embryo_RNAseq_heatmap_samples.pdf"), width=8, height = 8)
print(k)
dev.off()

strains=levels(sample_ID_list$strain)

# tx2gene table
gene2tx <- read.table("species/elegans/gene_annotation/c_elegans.PRJNA13758.WS285.canonical_geneset.gene_transcript_id.sorted.txt")
tx2gene = as.data.frame(cbind(gene2tx$V2, gene2tx$V1))


# all samples
files <- file.path("data/rnaseq/kallisto/", sample_ID_list$sample, "abundance.h5")
names(files) <- paste0("sample", 1:length(files))
txi.kallisto <- tximport(files, type = "kallisto", tx2gene = tx2gene)

dds <- DESeqDataSetFromTximport(txi.kallisto, colData = sample_ID_list, design = ~strain)

#PCA 
pdf(file=paste0(plot_dir_w_date, "/embryo_RNAseq_PCA_.pdf"), width=8, height = 8)
vsd <- vst(dds, blind=FALSE)
plotPCA(vsd, intgroup=c("sample"))
dev.off()

de1 <- DESeq(dds)

all_mut = c("ceh49", "6xCUT", "6xCUT", "7xCUT", "7xCUT", "7xCUT")
all_ctrl = c("wt", "wt", "ceh49", "wt", "ceh49", "6xCUT")

dir.create("DE_analysis")
wb_id_common_names = read.table("species/elegans/gene_annotation/c_elegans.PRJNA13758.WS285.canonical_geneset.gene_name.txt")
CUT_targets_bed = read.table("CUT_targets/CUT_targets_embryo.promoters.bed")
CUT_targets = unique(wb_id_common_names[wb_id_common_names$V1 %in% CUT_targets_bed$V5, 2])
transient_genes_wb = read.table("transient_genes/transient_genes.all.txt", header = T)
transient_genes = unique(wb_id_common_names[wb_id_common_names$V1 %in% transient_genes_wb$gene_name, 2])

for (test_n in c(1:length(all_mut))){
  res <- results(de1, c('strain', all_mut[test_n], all_ctrl[test_n]))
  downreg=res[res$log2FoldChange < -1 & res$padj < 0.001 & !is.na(res$padj),]
  upreg=res[res$log2FoldChange > 1 & res$padj < 0.001 & !is.na(res$padj),]
  write.table(downreg, file = paste0("DE_analysis/genes_downregulated_", all_mut[test_n], "_vs_", all_ctrl[test_n], ".txt"), quote=F, col.names = F, row.names = T, sep="\t")
  write.table(upreg, file = paste0("DE_analysis/genes_upregulated_", all_mut[test_n], "_vs_", all_ctrl[test_n], ".txt"), quote=F, col.names = F, row.names = T, sep="\t")
  write.table(res, file = paste0("DE_analysis/genes_DESeq_", all_mut[test_n], "_vs_", all_ctrl[test_n], ".txt"), quote=F, row.names = T, sep="\t")
  # run WormCAT on the downregulated genes (all and direct targets)
  downreg_genes = c("gene_name", row.names(downreg))
  write.table(downreg_genes, file = paste0("DE_analysis/genes_downregulated_", all_mut[test_n], "_vs_", all_ctrl[test_n], ".genes"), append = F, quote = F, row.names = F, col.names = F, sep="\t")
  worm_cat_fun(file_to_process = paste0("DE_analysis/genes_downregulated_", all_mut[test_n], "_vs_", all_ctrl[test_n], ".genes"),
               output_dir = paste0("DE_analysis/genes_downregulated_", all_mut[test_n], "_vs_", all_ctrl[test_n], ".wormcat"), 
               title = paste0(all_mut[test_n], "_vs_", all_ctrl[test_n], "_all"), 
               annotation_file = "data/external_data/whole_genome_v2_nov-11-2021.csv", 
               input_type = "Wormbase.ID", zip_files = F)
  downreg_targets = c("gene_name", row.names(downreg)[row.names(downreg) %in% CUT_targets_bed$V5])
  write.table(downreg_targets, file = paste0("DE_analysis/genes_downregulated_", all_mut[test_n], "_vs_", all_ctrl[test_n], ".direct_targets"), append = F, quote = F, row.names = F, col.names = F, sep="\t")
  worm_cat_fun(file_to_process = paste0("DE_analysis/genes_downregulated_", all_mut[test_n], "_vs_", all_ctrl[test_n], ".direct_targets"),
               output_dir = paste0("DE_analysis/genes_downregulated_", all_mut[test_n], "_vs_", all_ctrl[test_n], ".direct_targets.wormcat"), 
               title = paste0(all_mut[test_n], "_vs_", all_ctrl[test_n], "_direct_targets"), 
               annotation_file = "data/external_data/whole_genome_v2_nov-11-2021.csv", 
               input_type = "Wormbase.ID", zip_files = F)
  # volcanoplot - all genes
  res_all <- lfcShrink(de1, contrast = c('strain', all_mut[test_n], all_ctrl[test_n]), res=res, type = 'normal')
  row.names(res_all) = wb_id_common_names$V2
  res_all <- res_all[row.names(res_all) != "ceh-49",]
  pdf(paste0(plot_dir_w_date, "/embryo_DESeq_all_genes_", all_mut[test_n], "_vs_", all_ctrl[test_n], ".volcano.pdf"), width = 10, height = 8, useDingbats = F)
  print(EnhancedVolcano(res_all,
                  lab = rownames(res_all),x = 'log2FoldChange',
                  y = 'padj', xlab = bquote(~Log[2]~ 'fold change'),
                  pCutoffCol='padj', pCutoff = 1e-3,
                  FCcutoff = 1, pointSize = 1.5, labSize = 3.0, colAlpha = 1,
                  legendPosition = 'right', legendLabSize = 14, legendIconSize = 5.0,
                  title = paste0(all_mut[test_n], " vs ", all_ctrl[test_n]), subtitle = "all genes",
                  legendLabels = c("NS", "Log2FC", "adj-P", "Log2FC & adj-P")))
  dev.off()
  # volcanoplot - CEH-49 targets
  res_direct <- lfcShrink(de1, contrast = c('strain',all_mut[test_n], all_ctrl[test_n]), res=res, type = 'normal')
  row.names(res_direct) = wb_id_common_names$V2
  res_direct <- res_direct[row.names(res_direct) != "ceh-49" & row.names(res_direct) %in% CUT_targets,]
  pdf(paste0(plot_dir_w_date, "/embryo_DESeq_direct_targets_", all_mut[test_n], "_vs_", all_ctrl[test_n], ".volcano.pdf"), width = 10, height = 8, useDingbats = F)
  print(EnhancedVolcano(res_direct,
                  lab = rownames(res_direct),
                  x = 'log2FoldChange', y = 'padj',
                  xlab = bquote(~Log[2]~ 'fold change'),
                  pCutoffCol='padj', pCutoff = 1e-3, FCcutoff = 1, pointSize = 1.5, 
                  labSize = 3.0, colAlpha = 1, legendPosition = 'right', legendLabSize = 14, legendIconSize = 5.0,
                  title = paste0(all_mut[test_n], " vs ", all_ctrl[test_n]), subtitle = "CEH-49 targets", 
                  legendLabels = c("NS", "Log2FC", "adj-P", "Log2FC & adj-P")))
  dev.off()
  # volcanoplot - transient genes
  res_direct <- lfcShrink(de1, contrast = c('strain',all_mut[test_n], all_ctrl[test_n]), res=res, type = 'normal')
  row.names(res_direct) = wb_id_common_names$V2
  res_direct <- res_direct[row.names(res_direct) != "ceh-49" & row.names(res_direct) %in% transient_genes,]
  pdf(paste0(plot_dir_w_date, "/embryo_DESeq_transient_genes_", all_mut[test_n], "_vs_", all_ctrl[test_n], ".volcano.pdf"), width = 10, height = 8, useDingbats = F)
  print(EnhancedVolcano(res_direct,
                        lab = rownames(res_direct),
                        x = 'log2FoldChange', y = 'padj',
                        xlab = bquote(~Log[2]~ 'fold change'),
                        pCutoffCol='padj', pCutoff = 1e-3, FCcutoff = 1, pointSize = 1.5, 
                        labSize = 3.0, colAlpha = 1, legendPosition = 'right', legendLabSize = 14, legendIconSize = 5.0,
                        title = paste0(all_mut[test_n], " vs ", all_ctrl[test_n]), subtitle = "transient genes", 
                        legendLabels = c("NS", "Log2FC", "adj-P", "Log2FC & adj-P")))
  dev.off()
}
```
