## Supplementary material for "A lineage-resolved multimodal single-cell atlas reveals the genomic dynamics of early *C. elegans* development": Code and external data: emptydrops.20231106.R

```
library(DropletUtils)
library(Hmisc)

set.seed(42)

args = commandArgs(trailingOnly=TRUE)
current_date= paste(unlist(strsplit(as.character(Sys.Date()), "-")), collapse="")
dir.create(paste0("plots/", current_date))

sample_id = args[1]
barhop = as.numeric(args[2])
ambient = as.numeric(args[3])
wb_annotation = args[4]

# run EmptyDrops on the sample
sample_tod = Seurat::Read10X(paste0("data/sc/rnaseq/", sample_id, "/star_", wb_annotation, "/star_", wb_annotation, "_Solo.out/GeneFull/raw/um_reads"))

genes_type = read.table("species/elegans/gene_annotation/c_elegans.PRJNA13758.WS285.canonical_geneset.gene_type.txt")
gene_name = read.table("species/elegans/gene_annotation/c_elegans.PRJNA13758.WS285.canonical_geneset.gene_name.txt")
genes_type_filtered = genes_type[genes_type$V2 %in% c("lincRNA", "protein_coding", "pseudogene"),]
genes_name_filtered = gene_name[gene_name$V1 %in% genes_type_filtered$V1,]
sample_tod_filtered = sample_tod[row.names(sample_tod) %in% genes_name_filtered$V2,]
sample_counts_per_cell <- Matrix::colSums(sample_tod_filtered)

# inspect the UMI distribution to define the set of putative barhops
pdf(file=paste0("plots/", current_date, "/", sample_id, ".", wb_annotation, ".counts_per_cell.pdf"), width=6, height = 6)
hist(sample_counts_per_cell, breaks = seq(0, ceiling(max(sample_counts_per_cell)), by=1), xlim=c(0, 500), ylim=c(0, 10000), las=1)
dev.off()

# remove from the matrix the barcodes with fewer than N UMIs (N = max of barhop peak)
putative_bh = sample_counts_per_cell[sample_counts_per_cell < barhop]
sample_tod_no_bh = sample_tod_filtered[,colnames(sample_tod_filtered) %nin% names(putative_bh)]

# run ED
ed_sample_l_ambient <- emptyDrops(sample_tod_no_bh, lower=ambient, test.ambient=T)
dim(ed_sample_l_ambient[ed_sample_l_ambient$Total >= ambient & ed_sample_l_ambient$FDR < 0.001 & !is.na(ed_sample_l_ambient$FDR),])

pdf(file=paste0("plots/", current_date, "/", sample_id, ".", wb_annotation, ".ambient_P.pdf"), width=10, height = 6)
ed_sample_l_ambient_ambient = ed_sample_l_ambient[ed_sample_l_ambient$Total < ambient,]
hist(ed_sample_l_ambient_ambient$PValue, breaks = 1000)
dev.off()


# output the set of barcodes from real cells in two distinct files
sample_cells = ed_sample_l_ambient[ed_sample_l_ambient$Total >= ambient & ed_sample_l_ambient$FDR < 0.001 & !is.na(ed_sample_l_ambient$FDR),]
dir.create(paste0("data/sc/rnaseq/", sample_id, "/ED_RNA_", wb_annotation), recursive = T)
write.table(sample_cells, file = paste0("data/sc/rnaseq/", sample_id, "/ED_RNA_", wb_annotation, "/ED_RNA_barcodes.txt"), sep = "\t", quote = F, row.names = T, col.names = F)
```
