## Supplementary material for "A lineage-resolved multimodal single-cell atlas reveals the genomic dynamics of early *C. elegans* development": Code and external data: emptydrops_launcher.sh

```
#!/bin/bash
#
#SBATCH --job-name=emptydrops
#SBATCH --output=slurm_out/emptydrops.%N.%j.out
#SBATCH --error=slurm_err/emptydrops.%N.%j.err
#SBATCH --ntasks=1
#SBATCH -N 1
#SBATCH -n 1
#SBATCH --mem-per-cpu=8000
#SBATCH --mail-type=END,FAIL
#SBATCH --mail-user=

sample=$1
barhops=$2
ambient=$3
annotation=$4

Rscript scripts/emptydrops.20231106.R $sample $barhops $ambient $annotation
```
