## Supplementary material for "A lineage-resolved multimodal single-cell atlas reveals the genomic dynamics of early *C. elegans* development": Code and external data: fimo_clusters_hmm.sh

```
#!/bin/bash
#
#SBATCH --job-name=fimo_hmm
#SBATCH --output=slurm_out/fimo_hmm.%N.%j.out
#SBATCH --error=slurm_err/fimo_hmm.%N.%j.err
#SBATCH -n 8
#SBATCH -N 1
#SBATCH --mem=32G
#SBATCH --mail-type=END,FAIL
#SBATCH --mail-user=

clustered_motifs=$1
promoters=$2
output_dir=$3

genome_fa=species/elegans/genome/elegans.fa

sort -k 1,1 $output_dir/$clustered_motifs.motif_cluster | join -1 1 -2 4 - $promoters | awk 'BEGIN{OFS="\t";}{print $3, $4, $5, $1, $6, $7}' | sort -k 1,1 -k2,2n > $output_dir/$clustered_motifs.to_merge_motifs.bed

max_len=$(awk '{print $3-$2}' $output_dir/$clustered_motifs.to_merge_motifs.bed | sort -rg | head -n 1)
echo $max_len
mergeBed -i $output_dir/$clustered_motifs.to_merge_motifs.bed -c 4,6 -o first,first | awk -v var="$max_len" '{if ($3-$2 < var * 1.5) print}' | sort -k 4,4 > $output_dir/$clustered_motifs.merged_motifs.to_orient.bed

sort -k 1,1 $output_dir/$clustered_motifs.motif_cluster | join -1 1 -2 4 - $output_dir/$clustered_motifs.merged_motifs.to_orient.bed | awk 'BEGIN{OFS="\t";}{if ($2 == "+") print $3, $4, $5, $1, 0, $6; else if ($6 == "+") print $3, $4, $5, $1, 0, "-"; else print $3, $4, $5, $1, 0, "+"}' | sort -k 1,1 -k2,2n | fastaFromBed -fi $genome_fa -bed stdin -fo $output_dir/$clustered_motifs.merged_motifs.fa -s

mafft --thread 8 --localpair --maxiterate 1000 $output_dir/$clustered_motifs.merged_motifs.fa > $output_dir/$clustered_motifs.merged_motifs.mafft_out
hmmbuild $output_dir/$clustered_motifs.merged_motifs.mafft_out.hmm $output_dir/$clustered_motifs.merged_motifs.mafft_out

echo ">"$clustered_motifs > $output_dir/$clustered_motifs.merged_motifs.mafft_out.chen
hmmlogo --height_relent_all --no_indel $output_dir/$clustered_motifs.merged_motifs.mafft_out.hmm | sed '1,2d' | awk 'BEGIN{OFS = "\t";}{print $2, $3, $4, $5}' >> $output_dir/$clustered_motifs.merged_motifs.mafft_out.chen

chen2meme $output_dir/$clustered_motifs.merged_motifs.mafft_out.chen > $output_dir/$clustered_motifs.merged_motifs.mafft_out.meme
rm $output_dir/$clustered_motifs.merged_motifs.mafft_out.chen

ceqlogo -i $output_dir/$clustered_motifs.merged_motifs.mafft_out.meme -m $clustered_motifs -o $output_dir/$clustered_motifs.merged_motifs.mafft_out.logo.fwd.png -f PNG
ceqlogo -i $output_dir/$clustered_motifs.merged_motifs.mafft_out.meme -m $clustered_motifs -o $output_dir/$clustered_motifs.merged_motifs.mafft_out.logo.rev.png -f PNG -r
```
