## Supplementary material for "A lineage-resolved multimodal single-cell atlas reveals the genomic dynamics of early *C. elegans* development": Code and external data: final_cell_type_assignment.20240911.R

```
library(ggplot2)
library(dplyr)
library(Seurat)
library(Signac)
library(Hmisc)
library(stringr)
library(multimode)
library(rtracklayer)

source("scripts/second_derivative_binarisation.R")

current_date= paste(unlist(strsplit(as.character(Sys.Date()), "-")), collapse="")
plot_dir_w_date = paste0("/mnt/home3/ahringer/fnc21/10x/plots/", current_date)
dir.create(plot_dir_w_date, recursive=TRUE)

args = commandArgs(trailingOnly=TRUE)
seurat_object_file = args[1]
annotation = args[2]
late_clusters = args[3]

AB_barcodes_file = paste0("cell_type_annotation_all/AB.cell_assignment.", annotation, ".txt")
E_MS_C_D_barcodes_file = paste0("cell_type_annotation_all/E_MS_C_D.cell_assignment.", annotation, ".txt")
P_granule_file = "data/external_data/P_granule_transcripts.txt"

combined_seurat = readRDS(seurat_object_file)

# add cell_type information for somatic early lineage 
AB_barcodes = read.table(AB_barcodes_file)
E_MS_C_D_barcodes = read.table(E_MS_C_D_barcodes_file)

# remove underscores from cell type names, and change the "early_E_MS" name to "earlyEandMS" (to avoid mixups with "earlyEMS")
AB_barcodes$cell_type <- gsub('_','',AB_barcodes$cell_type)
E_MS_C_D_barcodes$cell_type[E_MS_C_D_barcodes$cell_type == "early_E_MS"] = "earlyEandMS"
E_MS_C_D_barcodes$cell_type <- gsub('_','',E_MS_C_D_barcodes$cell_type)

all_barcodes = as.vector(colnames(combined_seurat))

all_barcodes[all_barcodes %in% row.names(AB_barcodes)] <- AB_barcodes$cell_type
all_barcodes[all_barcodes %in% row.names(E_MS_C_D_barcodes)] <- E_MS_C_D_barcodes$cell_type
all_barcodes[grep(all_barcodes, pattern = "exp")] <- as.character(combined_seurat$seurat_clusters[grep(all_barcodes, pattern = "exp")])

combined_seurat$cell_type = all_barcodes

# subcluster the putative 2-cell cluster and retain only the cpg-3-enriched subclusters as "2_cell"
combined_seurat_avgExp_cpg3 = as.numeric(as.data.frame(AverageExpression(combined_seurat, slot = "data", assays = "RNA", features = "cpg-3")))
putative_twocell = which(combined_seurat_avgExp_cpg3 == max(combined_seurat_avgExp_cpg3))-1

combined_seurat = FindSubCluster(combined_seurat, cluster = putative_twocell, graph.name = "wsnn", resolution = 0.5, subcluster.name = "twocell")
combined_seurat$twocell = as.character(combined_seurat$twocell)
twocell_sucl_names = unique(combined_seurat@meta.data$twocell)[grep(unique(combined_seurat@meta.data$twocell), pattern = "_")]
twocell_subclusters = c()
for (i in twocell_sucl_names){
  if ("cpg-3" %in% row.names(FindMarkers(combined_seurat, ident.1 = i, group.by = "twocell", only.pos = T, assay = "RNA", logfc.threshold = 2))){
    twocell_subclusters = append(twocell_subclusters, i)
  }
}

combined_seurat$cell_type[combined_seurat$twocell %in% twocell_subclusters] = "twocell"
#DimPlot(combined_seurat, reduction = "umapWNN", group.by = "cell_type", order = T, label = T, label.size = 3) & NoLegend()

# identify putative Z2/Z3 cluster
combined_seurat_avgExp_prg2 = as.numeric(as.data.frame(AverageExpression(combined_seurat, slot = "data", assays = "RNA", features = "prg-2")))
putative_z2z3 = which(combined_seurat_avgExp_prg2 == max(combined_seurat_avgExp_prg2))-1

combined_seurat$cell_type[combined_seurat$twocell %in% putative_z2z3] = "Z2Z3"
#DimPlot(combined_seurat, reduction = "umapWNN", group.by = "cell_type", order = T, label = T, label.size = 3) & NoLegend()

### define identity of P cells based on increased accessibility of m1m2 motifs
# define P cells based on enrichment of P-granule module, then subcluster them
p_granule_genes = read.table(P_granule_file)
combined_seurat = AddModuleScore(combined_seurat, features = list(p_granule_genes$V1), assay = "RNA")
combined_seurat$p_granules = combined_seurat$Cluster1
combined_seurat$Cluster1 = NULL

all_clusters = unique(combined_seurat@meta.data$twocell)
all_clusters = all_clusters[all_clusters %nin% twocell_subclusters]

combined_seurat_avg_p_granule_exp = c()
for (cl in all_clusters){
  combined_seurat_avg_p_granule_exp = append(combined_seurat_avg_p_granule_exp, mean(combined_seurat$p_granules[combined_seurat$twocell == cl]))
}

loc <- locmodes(combined_seurat_avg_p_granule_exp, mod0 = 2, display = T)
low_high_exp_loc = second_derivative_binarization(combined_seurat_avg_p_granule_exp, loc$cbw$bw, loc$locations, "P-granules")
p_cell_clusters <- all_clusters[which(combined_seurat_avg_p_granule_exp >= low_high_exp_loc[2])]

combined_seurat$cell_type[combined_seurat$twocell %in% p_cell_clusters] = "Pcell"

Idents(combined_seurat) = "cell_type" 
combined_seurat = FindSubCluster(combined_seurat, cluster = "Pcell", graph.name = "wsnn", resolution = 0.3, subcluster.name = "pcell")
combined_seurat$pcell = as.character(combined_seurat$pcell)
pcell_sucl_names = unique(combined_seurat@meta.data$pcell)[grep(unique(combined_seurat@meta.data$pcell), pattern = "Pcell_")]
combined_seurat$cell_type = combined_seurat$pcell

#DimPlot(combined_seurat, reduction = "umapWNN", group.by = "cell_type", order = T, label = T, label.size = 3) & NoLegend()
# load the m1m2 promoters, then quantify their accessibility in P cells to distinguish P2, P3 and P4
m1m2_elements = import.bed("data/external_data/reg_elements_all.elegans.gl_specific.m1m2.bed")

m1m2_elements_matrix <- FeatureMatrix(combined_seurat[["WNN"]]@fragments, sep = c("-", "-"), features=m1m2_elements, cells = colnames(combined_seurat))
combined_seurat[["m1m2_elements"]] <- CreateChromatinAssay(counts = m1m2_elements_matrix, sep = c(":", "-"), fragments = combined_seurat[["WNN"]]@fragments)

mean_counts = c()
for (cl in c(0:2)){
  mean_counts[paste0("Pcell_", cl)] = mean(combined_seurat$nCount_m1m2_elements[combined_seurat$cell_type == paste0("Pcell_", cl)])
}

P2 = names(which(mean_counts == min(mean_counts)))
P3 = names(which(mean_counts < max(mean_counts) & mean_counts > min(mean_counts)))
P4 = names(which(mean_counts == max(mean_counts)))

combined_seurat$cell_type[combined_seurat$cell_type == P2] = "P2"
combined_seurat$cell_type[combined_seurat$cell_type == P3] = "P3"
combined_seurat$cell_type[combined_seurat$cell_type == P4] = "P4"

pdf(file=paste0(plot_dir_w_date, "/Pcell_annotation.pdf"), width=12, height = 6)
par(mfrow=c(1,2))
boxplot(combined_seurat$nFeature_m1m2_elements[combined_seurat$cell_type == "P2"],
        combined_seurat$nFeature_m1m2_elements[combined_seurat$cell_type == "P3"],
        combined_seurat$nFeature_m1m2_elements[combined_seurat$cell_type == "P4"],
        combined_seurat$nFeature_m1m2_elements[combined_seurat$cell_type == "Z2Z3"], 
        outline = F, main = "Accessible m1m2 sites in germline precursors", las = 1, names = c("P2", "P3", "P4", "Z2-Z3"))

plot(density(combined_seurat$nFeature_m1m2_elements[combined_seurat$cell_type == "P2"], bw = 3), col = 1, lwd = 2, lty = 1, las = 1, main = "Accessible m1m2 sites in germline precursors")
lines(density(combined_seurat$nFeature_m1m2_elements[combined_seurat$cell_type == "P3"], bw = 3), col = 2, lwd = 2, lty = 2)
lines(density(combined_seurat$nFeature_m1m2_elements[combined_seurat$cell_type == "P4"], bw = 3), col = 3, lwd = 2, lty = 3)
lines(density(combined_seurat$nFeature_m1m2_elements[combined_seurat$cell_type == "Z2Z3"], bw = 3), col = 4, lwd = 2, lty = 4)
legend(60, 0.05, legend = c("P2", "P3", "P4", "Z2-Z3"), fill = c(1:4), bty = "n")
dev.off()

# add late cells
cell_type_seurat_cluster = combined_seurat$cell_type

reassigned_clusters_table = read.table(late_clusters)
reassigned_cell_type = cell_type_seurat_cluster

for (cluster_n in reassigned_clusters_table$V1){
  reassigned_cell_type[cell_type_seurat_cluster == cluster_n] = paste0(reassigned_clusters_table[reassigned_clusters_table$V1 == cluster_n, 2], cluster_n)
}

combined_seurat$cell_type = reassigned_cell_type
cl_n = max(as.numeric(combined_seurat$seurat_clusters))-1
combined_seurat$cell_type[combined_seurat$cell_type %in% c(0:cl_n)] = "unassigned"

#DimPlot(combined_seurat, reduction = "umapWNN", group.by = "cell_type", order = T, label = T, label.size = 3) & NoLegend()

# as the Cxx cluster has high ATAC fragment counts, low RNA complexity and does not seem to reflect the characteristics of a Cxx cell, lable these cells as unassigned
combined_seurat$cell_type[combined_seurat$cell_type == "Cxx"] = "unassigned"

# add lineage information 
cell_lineage = combined_seurat$cell_type

cell_lineage = unlist(strsplit(cell_lineage, "early"))
cell_lineage = cell_lineage[cell_lineage != ""]

new_cell_lineage = cell_lineage
new_cell_lineage[str_detect(cell_lineage, "AB[a-z]{2}$")] = "ABxx"
new_cell_lineage[str_detect(cell_lineage, "AB[a-z]{3}$")] = "ABxxx"
new_cell_lineage[str_detect(cell_lineage, "AB[a-z]{4}$")] = "ABxxxx"
new_cell_lineage[str_detect(cell_lineage, "AB[a-z]{5}")] = "ABxxxxx"
new_cell_lineage[str_detect(cell_lineage, "AB[a-z]{6}")] = "ABxxxxxx"
new_cell_lineage[str_detect(cell_lineage, "MS[a-z]{1}$")] = "MSx"
new_cell_lineage[str_detect(cell_lineage, "MS[a-z]{2}$")] = "MSxx"
new_cell_lineage[str_detect(cell_lineage, "MS[a-z]{3}")] = "MSxxx"
new_cell_lineage[str_detect(cell_lineage, "MS[a-z]{4}")] = "MSxxxx"
new_cell_lineage[str_detect(cell_lineage, "C[a-z]{1}$")] = "Cx"
new_cell_lineage[str_detect(cell_lineage, "C[a-z]{2}$")] = "Cxx"
new_cell_lineage[str_detect(cell_lineage, "Cxxx")] = "Cxxx"
new_cell_lineage[str_detect(cell_lineage, "D[a-z]{1}$")] = "Dx"
new_cell_lineage[str_detect(cell_lineage, "DxxCxxx")] = "DxxCxxx"
new_cell_lineage[str_detect(cell_lineage, "unassigned")] = "unassigned"

combined_seurat$cell_lineage = as.character(new_cell_lineage)
#DimPlot(combined_seurat, reduction = "umapWNN", group.by = "cell_lineage", order = T, label = T, label.size = 3) & NoLegend()

pdf(file=paste0(plot_dir_w_date, "/wt.all_samples.all_cells.", annotation, ".UMAP.pdf"), width=16, height = 8)
p1 = DimPlot(combined_seurat, reduction = "umapWNN", group.by = "cell_type", order = T, label = T, label.size = 4) & NoLegend()
p2 = DimPlot(combined_seurat, reduction = "umapWNN", group.by = "cell_lineage", order = T, label = T, label.size = 4) & NoLegend()
print(p1 | p2)
dev.off()

combined_seurat$p_granules <- NULL
combined_seurat$twocell <- NULL
combined_seurat$SCT_snn_res.3 <- NULL
combined_seurat$WNN_snn_res.3 <- NULL

# output distinct fragment files per cluster, cell lineage, and cell type to create bigWig tracks and call peaks
DefaultAssay(combined_seurat) <- "WNN"

dir.create(paste0("cluster_specific_peaks/round_2_all_samples/", annotation, "/cell_lineage/bed"), recursive = T)
dir.create(paste0("cluster_specific_peaks/round_2_all_samples/", annotation, "/cell_type/bed"), recursive = T)

SplitFragments(combined_seurat, group.by = "cell_lineage", outdir = paste0("cluster_specific_peaks/round_2_all_samples/", annotation, "/cell_lineage/bed"), append = T)
SplitFragments(combined_seurat, group.by = "cell_type", outdir = paste0("cluster_specific_peaks/round_2_all_samples/", annotation, "/cell_type/bed"), append = T)

# output number of cells per cluster
n_cell_lineage = unique(combined_seurat$cell_lineage)
n_cell_lineage = n_cell_lineage[order(n_cell_lineage, as.numeric(n_cell_lineage))]
cells_lineage=data.frame(row.names=n_cell_lineage, "n_cells" = rep(NA, length(n_cell_lineage)))
for (n_c in c(1:length(n_cell_lineage))){
  cells_lineage[n_c,1]=dim(combined_seurat[,combined_seurat$cell_lineage == n_cell_lineage[n_c]])[2]
}
write.table(cells_lineage, file = paste0("seurat_objects/wt_all.all_samples.all_cells.", annotation, ".cells_per_cell_lineage.20240911.txt"), quote = F, sep = "\t", row.names = T, col.names = T)

n_cell_type = unique(combined_seurat$cell_type)
n_cell_type = n_cell_type[order(n_cell_type, as.numeric(n_cell_type))]
cells_type=data.frame(row.names=n_cell_type, "n_cells" = rep(NA, length(n_cell_type)))
for (n_c in c(1:length(n_cell_type))){
  cells_type[n_c,1]=dim(combined_seurat[,combined_seurat$cell_type == n_cell_type[n_c]])[2]
}
write.table(cells_type, file = paste0("seurat_objects/wt_all.all_samples.all_cells.", annotation, ".cells_per_cell_type.20240911.txt"), quote = F, sep = "\t", row.names = T, col.names = T)

# export dataset
saveRDS(combined_seurat, file = paste0("seurat_objects/wt_all.all_samples.all_cells.", annotation, ".20240911.rds"))
write.table(combined_seurat[[]], file = paste0("seurat_objects/wt_all.all_samples.all_cells.", annotation, ".table.20240911.txt"), quote = F, sep = "\t", row.names = T, col.names = T)

#saveRDS(combined_seurat_early, file = paste0("seurat_objects/wt.all_samples.early_cells.", annotation, ".rds"))
#write.table(combined_seurat_early[[]], file = paste0("seurat_objects/wt.all_samples.early_cells.", annotation, ".table.txt"), quote = F, sep = "\t", row.names = T, col.names = T)
```
