## Supplementary material for "A lineage-resolved multimodal single-cell atlas reveals the genomic dynamics of early *C. elegans* development": Code and external data: gene_promoter_locus_extension.py

```
import sys

gene_locus = sys.argv[1]
promoters = sys.argv[2]
extended_gene_locus = sys.argv[3]
gene_type= sys.argv[4]

# select only coding genes, pseudogenes, and lincRNAs
filtered_genes = []
gene_types_allowed = ["protein_coding", "pseudogene", "lincRNA"]
for line in open(gene_type):
    G=line.rstrip().split("\t")
    if G[1] in gene_types_allowed:
        filtered_genes.append(G[0])

# get longest transcript locus for each filtered gene
gene_locus_dict = {}
for line in open(gene_locus):
    G=line.rstrip().split("\t")
    if G[3] in filtered_genes:
        gene_locus_dict[G[3]] = [G[0], G[1], G[2], G[5]]

# extend gene locus if a promoters is annotated
for line in open(promoters):
    G=line.rstrip().split("\t")
    if G[4] in gene_locus_dict:
        if G[5] == "+" and eval(gene_locus_dict[G[4]][1]) > eval(G[1]):
            gene_locus_dict[G[4]][1] = G[1]
        elif G[5] == "-" and eval(gene_locus_dict[G[4]][2]) < eval(G[2]):
            gene_locus_dict[G[4]][2] = G[2]

# output extended locus annotation
new = open(extended_gene_locus, "w")
for gene in gene_locus_dict:
    temp = gene_locus_dict[gene][0] + "\t" + gene_locus_dict[gene][1] + "\t" + gene_locus_dict[gene][2] + "\t" + gene + "\t0\t" + gene_locus_dict[gene][3] + "\n"
    new.write(temp)

new.close()
```
