## Supplementary material for "A lineage-resolved multimodal single-cell atlas reveals the genomic dynamics of early *C. elegans* development": Code and external data: gtf_3_utr_extension.20231108.py

```
import sys
import os

input_gtf = sys.argv[1]
bin_size = sys.argv[2]
distance_from_tts = sys.argv[3]
genome_chr = sys.argv[4]
gene_transcript = sys.argv[5]
file_path = sys.argv[6]
extension_type=sys.argv[7]

# convert GTF annotation in BED12 format (coding genes)
# modification 20231106: keep MtDNA genes
input_genePred = ".".join(input_gtf.split(".")[:-1]) + ".genePred"
input_bed = ".".join(input_gtf.split(".")[:-1]) + ".bed"
cmd="""gtfToGenePred """ + file_path + "/" + input_gtf + """ """ + file_path + "/" + input_genePred + """; genePredToBed """ + file_path + "/" + input_genePred + """ """ + file_path + "/" + input_bed + """.temp; awk 'BEGIN{{FS="\t";OFS="\t";}}{{print $1, $2, $3, $4, $5, $6, $7, $8, $9, $10, $11, $12}}' """ + file_path + "/" + input_bed + """.temp | sort -k 4,4 > """ + file_path + "/" + input_bed
os.system(cmd)

# extract coordinates of last exon in each transcript
input_bed6 = ".".join(input_gtf.split(".")[:-1]) + ".bed6"
cmd = "bed12ToBed6 -i " + file_path + "/" + input_bed + " -n | sort -k 4,4 -k 5,5rn > " + file_path + "/" + input_bed6
os.system(cmd)

# get chromosomes size in a dictionary
chr_size = {}
for line in open(genome_chr):
    chr_size[line.split("\t")[0]] = eval(line.split("\t")[1])

extensions = distance_from_tts.split(",")
for extension in extensions:
    print(extension)
    extended_exons = ".".join(input_gtf.split(".")[:-1]) + ".3prime_" + extension + "_" + str(eval(extension) + eval(bin_size))
    new = open(file_path + "/" + extension_type + "/" + extended_exons, "w")
    for line in open(file_path + "/" + input_bed):
        G = line.rstrip().split("\t")
        if G[5] == "+":
            if eval(G[2]) + eval(extension) + eval(bin_size) < chr_size[G[0]]:
                new.write(G[0] + "\t" + G[1] + "\t" + str(eval(G[2]) + eval(extension) + eval(bin_size)) + "\t" + G[3] + "\t0\t" + G[5] + "\n")
        else:
            if eval(G[1]) - eval(extension) - eval(bin_size) > 0:
                new.write(G[0] + "\t" + str(eval(G[1]) - eval(extension) - eval(bin_size)) + "\t" + G[2] + "\t" + G[3] + "\t0\t" + G[5] + "\n")
    new.close()
    # replace transcript id with gene id in extended exons and gene annotation
    extended_exons_genename = ".".join(input_gtf.split(".")[:-1]) + ".genes.3prime_" + extension + "_" + str(eval(extension) + eval(bin_size))
    cmd = """join -1 2 -2 4 """ + file_path + "/" + gene_transcript + """ """ + file_path + "/" + extension_type + "/" + extended_exons + """ | awk 'BEGIN{OFS="\t"}{print $3, $4, $5, $2, $6, $7, $1}' > """ + file_path + "/" + extension_type + "/" + extended_exons_genename
    os.system(cmd)
    input_bed_genename = ".".join(input_gtf.split(".")[:-1]) + ".genes.bed"
    cmd = """join -1 2 -2 4 """ + file_path + "/" + gene_transcript + """ """ + file_path + "/" + input_bed + """ | awk 'BEGIN{OFS="\t"}{print $3, $4, $5, $2, $6, $7, $8, $9, $10, $11, $12, $1}' > """ + file_path + "/" + extension_type + "/" + input_bed_genename
    os.system(cmd)
    # extract only bins not overlapping with any downstream gene (in ANY orientation)
    extended_exons_to_filter = ".".join(input_gtf.split(".")[:-1]) + ".genes.3prime_" + extension + "_" + str(eval(extension) + eval(bin_size)) + "_overlapping_genes"
    cmd = """intersectBed -a """ + file_path + "/" + extension_type + "/" + extended_exons_genename + """ -b """ + file_path + "/" + extension_type + "/" + input_bed_genename + """ -wa -wb | awk '{if ($4 != $11) print $7}' | sort | uniq > """ + file_path + "/" + extension_type + "/" + extended_exons_to_filter
    os.system(cmd)
    # modified 20231101: simply output new transcripts in bed6 format; coverage will be calculated on whole locus (not only exons) and no GTF will be needed
    input_bed_corrected = ".".join(input_gtf.split(".")[:-1]) + ".3prime_" + extension + "_" + str(eval(extension) + eval(bin_size)) + "_extended.bed"
    cmd = """sort -k 7,7 """ + file_path + "/" + extension_type + "/" + extended_exons_genename + """ | join -v 1 -1 7 -2 1 - """ + file_path + "/" + extension_type + "/" + extended_exons_to_filter + """ | awk 'BEGIN{OFS="\t";}{print $2, $3, $4, $1 "_" """ + str(eval(extension)) + """ "_" """ + str(eval(extension) + eval(bin_size)) + """, $6, $7}' > """ + file_path + "/" + extension_type + "/" + input_bed_corrected
    os.system(cmd)
```
