## Supplementary material for "A lineage-resolved multimodal single-cell atlas reveals the genomic dynamics of early *C. elegans* development": Code and external data: gtf_3_utr_extension.bin_selection.20231108.py

```
import sys
import os

original_counts = sys.argv[1]
extended_counts = sys.argv[2]
bin_size = sys.argv[3]
distance_from_tts = sys.argv[4]
transcript_extension = sys.argv[5]

# import the counts per original transcript and store them in a dictionary (key: transcript, value: ordered list of counts)
# modified on 20231101: include extended transcripts only if they have > 100 reads
original_counts_dict = {}
for line in open(original_counts):
    G=line.rstrip().split("\t")
    if eval(G[6]) > 100:
        original_counts_dict[G[3]] = eval(G[6])

# define max extension of each transcript
max_extension_dict = {}
for line in open(extended_counts):
    G=line.rstrip().split("\t")
    transcript_id = "_".join(G[3].split("_")[:-2])
    transcript_bin = int(eval("_".join(G[3].split("_")[-1:]))/100)
    if transcript_id not in max_extension_dict:
        max_extension_dict[transcript_id] = 0
    if max_extension_dict[transcript_id] < transcript_bin:
        max_extension_dict[transcript_id] = transcript_bin

# import the counts of each extended transcript and store them in a dictionary (key: transcript, value: ordered list of counts)
extended_counts_dict = {}
for line in open(extended_counts):
    G=line.rstrip().split("\t")
    transcript_id = "_".join(G[3].split("_")[:-2])
    transcript_bin = int(eval("_".join(G[3].split("_")[-1:]))/100)
    if transcript_id not in extended_counts_dict:
#        print(transcript_id)
#        print(max_extension_dict[transcript_id])
        extended_counts_dict[transcript_id] = list([0] * max_extension_dict[transcript_id])
    extended_counts_dict[transcript_id][transcript_bin-1] = eval(G[6])

# transform the counts of the extended transcripts in per-bin counts
# modified on 20231101: exclude extended transcripts if original transcript had less than 100 reads
extended_counts_bin = {}
for transcript in extended_counts_dict:
    # exclude transcript if original transcript had less than 100 reads
    if transcript not in original_counts_dict:
        continue
    extended_counts_bin[transcript] = [extended_counts_dict[transcript][0] - original_counts_dict[transcript]]
    for i in range(1, len(extended_counts_dict[transcript])):
        extended_counts_bin[transcript].append(extended_counts_dict[transcript][i] - extended_counts_dict[transcript][i-1])


# output the list of extensions, based on these rules:
## if the transcript could not be shrinked at its 3', report the original
## extend a transcript if the counts per extension bin do not exceed the count in the previous bin by more than 10%
## if the tentative extension exceed by at least 10% the coverage of the original transcript, keep the shortest extension retaining more than 90% of the additional reads.

new = open(transcript_extension, "w")
final_bins={1:0, 2:0, 3:0, 4:0, 5:0}

print(original_counts_dict["K04C2.6.1"])
print(extended_counts_dict["K04C2.6.1"])
print(extended_counts_bin["K04C2.6.1"])

#print(original_counts_dict["Y74C9A.2a.3"])
#print(extended_counts_dict["Y74C9A.2a.3"])
#print(extended_counts_bin["Y74C9A.2a.3"])

#print(original_counts_dict["Y74C9A.1.1"])
#print(extended_counts_dict["Y74C9A.1.1"])
#print(extended_counts_bin["Y74C9A.1.1"])

for transcript in extended_counts_bin:
    if extended_counts_dict[transcript][-1] < original_counts_dict[transcript]*1.2 or original_counts_dict[transcript] < 100:
        new.write(transcript + "\t0\n")
    else:
        final_bin = 0
        for i in range(len(extended_counts_bin[transcript]), 0, -1):
            if extended_counts_dict[transcript][i-1] < extended_counts_dict[transcript][-1]*0.9:
                final_bin = 1
                new.write(transcript + "\t" + str(eval(distance_from_tts.split(",")[i]) + eval(bin_size)) + "\n")
                break
        if final_bin == 0:
            new.write(transcript + "\t100\n")

new.close()
```
