## Supplementary material for "A lineage-resolved multimodal single-cell atlas reveals the genomic dynamics of early *C. elegans* development": Code and external data: gtf_3_utr_extension.rnaseq_extended.20231108.py

```
import sys
import os

input_bed = sys.argv[1]
input_extension = sys.argv[2]
bin_size = sys.argv[3]
distance_from_tts = sys.argv[4]
input_gene_transcripts = sys.argv[5]
gene_annotation_path = sys.argv[6]
extension_type = sys.argv[7]

extended_genes = {}
for line in open(gene_annotation_path + "/" + extension_type + "/" + input_extension):
    if line.rstrip().split("\t")[1] != "0":
        extended_genes[line.split("\t")[0]] = eval(line.rstrip().split("\t")[1])

out_corrected_bed = ".".join(input_bed.split(".")[:-1]) + "." + extension_type + ".3prime_extended.rnaseq_corrected.bed"
new = open(gene_annotation_path + "/" + out_corrected_bed, "w")
for line in open(gene_annotation_path + "/" + input_bed):
    G=line.split("\t")
    if G[3] not in extended_genes or extended_genes[G[3]] == 0:
        new.write(line)
    else:
        if G[5] == "+":
            corrected_start = G[1]
            corrected_end = str(eval(G[2]) + extended_genes[G[3]])
            corrected_exons = ",".join(G[10].split(",")[:-2]) + "," + str(eval(G[10].split(",")[-2]) + extended_genes[G[3]]) + ","
            corrected_exons = corrected_exons.lstrip(",")
            corrected_introns = G[11]
            corrected_CDS_start = G[6]
            corrected_CDS_end = G[7]
            if corrected_CDS_start == corrected_CDS_end:
                corrected_CDS_start = corrected_end
                corrected_CDS_end = corrected_end
        else:
            corrected_start = str(eval(G[1]) - extended_genes[G[3]])
            corrected_end = G[2]
            corrected_exons = str(eval(G[10].split(",")[0]) + extended_genes[G[3]]) + "," + ",".join(G[10].split(",")[1:])
            corrected_introns = "0," + ",".join([str(eval(x) + extended_genes[G[3]]) for x in G[11].split(",")[1:-1]])
            corrected_introns = corrected_introns.rstrip(",") + ",\n"
            corrected_CDS_start = G[6]
            corrected_CDS_end =	G[7]
            if corrected_CDS_start == corrected_CDS_end:
                corrected_CDS_start = corrected_start
                corrected_CDS_end = corrected_start
        G[1] = corrected_start
        G[2] = corrected_end
        G[10] = corrected_exons
        G[11] = corrected_introns
        G[6] = corrected_CDS_start
        G[7] = corrected_CDS_end
        new.write("\t".join(G))


new.close()

out_corrected_gtf = ".".join(input_bed.split(".")[:-1]) + "." + extension_type + ".3prime_extended.rnaseq_corrected.gtf"
cmd = """sort -k 4,4 """ + gene_annotation_path + "/" + out_corrected_bed + """ > """ + gene_annotation_path + "/" + out_corrected_bed + """.sorted; bedToGenePred """ + gene_annotation_path + "/" + out_corrected_bed + """.sorted """ + gene_annotation_path + "/" + out_corrected_bed + """.genePred; join -1 2 -2 1 """ + gene_annotation_path + "/" + input_gene_transcripts + """ """ + gene_annotation_path + "/" + out_corrected_bed + """.genePred | awk 'BEGIN{OFS="\t";}{print $1, $3, $4, $5, $6, $7, $8, $9, $10, $11, 0, $2}' > """ + gene_annotation_path + "/" + out_corrected_bed + """.genePredExt ; genePredToGtf "file" """ + gene_annotation_path + "/" + out_corrected_bed + """.genePredExt """ + gene_annotation_path + "/" + out_corrected_gtf
os.system(cmd)
```
