## Supplementary material for "A lineage-resolved multimodal single-cell atlas reveals the genomic dynamics of early *C. elegans* development": Code and external data: gtf_gene_type_filter.py

```
import sys

input_file=sys.argv[1]
gene_types=sys.argv[2]
output_file=sys.argv[3]
genes=sys.argv[4]

allowed_types = gene_types.split(",")

genes_to_retain=[]
for line in open(genes):
    if line.rstrip().split("\t")[1] in allowed_types:
        genes_to_retain.append(line.rstrip().split("\t")[0])

new=open(output_file, "w")
for line in open(input_file):
    if line[0] == "#":
        continue
    G=line.rstrip().split("\t")
    K = G[8].split('"')
    gene_name=K[1]
    if gene_name in genes_to_retain:
        new.write(line)

new.close()
```
