## Supplementary material for "A lineage-resolved multimodal single-cell atlas reveals the genomic dynamics of early *C. elegans* development": Code and external data: idr_atac.sh

```
#!/bin/bash
#
#SBATCH --job-name=idr
#SBATCH --output=slurm_out/idr.%N.%j.out
#SBATCH --error=slurm_err/idr.%N.%j.err
#SBATCH --ntasks=1
#SBATCH -N 1
#SBATCH -n 8
#SBATCH --mem-per-cpu=4000
#SBATCH --mail-type=END,FAIL
#SBATCH --mail-user=

input_filename=$1
input_path=$2

mkdir -p $input_path/IDR
mkdir -p $input_path/IDR_logs

idr --samples $input_path"/"$input_filename"_rep_1.bed" $input_path"/"$input_filename"_rep_2.bed" --input-file-type narrowPeak --output-file $input_path"/IDR/"$input_filename"_idr.txt" --output-file-type bed --log-output-file $input_path"/IDR_logs/"$input_filename"_idr.log"  --plot --rank signal.value

awk -v var="$input_filename" 'BEGIN{OFS="\t";}{if($5 >= 830) print $1, $2, $3, var}' $input_path"/IDR/"$input_filename"_idr.txt" > $input_path"/IDR/"$input_filename"_idr.0.01.bed"
awk -v var="$input_filename" 'BEGIN{OFS="\t";}{if($5 >= 540) print $1, $2, $3, var}' $input_path"/IDR/"$input_filename"_idr.txt" > $input_path"/IDR/"$input_filename"_idr.0.05.bed"
```
