## Supplementary material for "A lineage-resolved multimodal single-cell atlas reveals the genomic dynamics of early *C. elegans* development": Code and external data: idr_vs_all_macs_peaks_atac_comparison.sh

```
#!/bin/bash
#
#SBATCH --job-name=idr_vs_macs
#SBATCH --output=slurm_out/idr_vs_macs.%N.%j.out
#SBATCH --error=slurm_err/idr_vs_macs.%N.%j.err
#SBATCH --ntasks=1
#SBATCH -N 1
#SBATCH -n 8
#SBATCH --mem-per-cpu=4000
#SBATCH --mail-type=END,FAIL
#SBATCH --mail-user=

sample=$1
input_path=$2
idr_threshold=$3

mkdir -p $input_path/IDR_vs_MACS

intersectBed -a $input_path"/IDR/"$sample"_idr."$idr_threshold".bed" -b cluster_specific_peaks/round_2_all_samples/WS285_extended_no_ovlp/all_peaks.pre_IDR.bed -u | cut -f 1,2,3 > $input_path/IDR_vs_MACS/$sample.idr.$idr_threshold.macs.bed
echo "#IDR" >>  $input_path/IDR_vs_MACS/$sample.idr.$idr_threshold.macs.bed
intersectBed -b $input_path"/IDR/"$sample"_idr."$idr_threshold".bed" -a cluster_specific_peaks/round_2_all_samples/WS285_extended_no_ovlp/all_peaks.pre_IDR.bed -v | cut -f 1,2,3 >> $input_path/IDR_vs_MACS/$sample.idr.$idr_threshold.macs.bed
echo "#no_IDR" >>  $input_path/IDR_vs_MACS/$sample.idr.$idr_threshold.macs.bed

computeMatrix scale-regions -R $input_path/IDR_vs_MACS/$sample.idr.$idr_threshold.macs.bed -S cluster_specific_peaks/round_2_all_samples/WS285_extended_no_ovlp/cell_type/bw/$sample.cpm.bw -o $input_path/IDR_vs_MACS/$sample.idr.$idr_threshold.macs.mtx.gz -a 500 -b 500 -m 200 --binSize 10 -p 12 -bl data/external_data/ce11-blacklist.bed

plotHeatmap -m $input_path/IDR_vs_MACS/$sample.idr.$idr_threshold.macs.mtx.gz -o $input_path/IDR_vs_MACS/$sample.idr.$idr_threshold.macs.pdf --heatmapHeight 25 --heatmapWidth 5 --colorMap jet --whatToShow 'heatmap and colorbar' --zMin 0 --zMax 20
```
