## Supplementary material for "A lineage-resolved multimodal single-cell atlas reveals the genomic dynamics of early *C. elegans* development": Code and external data: inherited_zygotic_promoters.gene_expression.20241127.R

```
library(cluster)
library(Seurat)
library(Signac)
library(Hmisc)
library(RColorBrewer)
library(presto)
library(rtracklayer)
library(ComplexHeatmap)
library(UpSetR)

source("data/external_data/wormcat_function.R")

current_date= paste(unlist(strsplit(as.character(Sys.Date()), "-")), collapse="")
plot_dir_w_date = paste0("paper_figures/", current_date)
dir.create(plot_dir_w_date, recursive=TRUE)

# load RDS and WBid table
combined_seurat = readRDS("seurat_objects/wt_all.all_samples.all_cells.WS285_extended_no_ovlp.final_peakset_IDR.rds")
wb_id_common_names = read.table("species/elegans/gene_annotation/c_elegans.PRJNA13758.WS285.canonical_geneset.filtered.gene_name.txt")

# look at germline expression of genes regulated by promoters not/accessible in the germline
inherited_promoters = read.table("inherited_zygotic_program/4cell_promoters.inherited.bed")
zygotic_promoters = read.table("inherited_zygotic_program/4cell_promoters.zygotic.bed")

inherited_promoters_genes = unique(inherited_promoters$V5)
zygotic_promoters_genes_all = unique(zygotic_promoters$V5)

zygotic_promoters_genes = zygotic_promoters_genes_all[zygotic_promoters_genes_all %nin% inherited_promoters_genes]

inherited_promoters_genes_name = wb_id_common_names[wb_id_common_names$V1 %in% inherited_promoters_genes, 2]
zygotic_promoters_genes_name = wb_id_common_names[wb_id_common_names$V1 %in% zygotic_promoters_genes, 2]

# split the inherited genes based on their expression in 2 vs 4 cells; output gene lists
two_four_de = FindMarkers(combined_seurat, ident.1 = "twocell", ident.2 = c("ABx", "EMS"), group.by = "cell_lineage", 
                          assay = "RNA", features = inherited_promoters_genes_name, 
                          logfc.threshold = 0, min.pct = 0, only.pos = F)

#maternally_enriched_inherited_genes = row.names(two_four_de[two_four_de$p_val_adj < 0.01 & !is.na(two_four_de$p_val_adj) & two_four_de$avg_log2FC > 0,])
zygotic_enriched_inherited_genes = row.names(two_four_de[two_four_de$p_val_adj < 0.01 & !is.na(two_four_de$p_val_adj) & two_four_de$avg_log2FC < 0,])
#non_biased_enriched_inherited_genes = row.names(two_four_de[row.names(two_four_de) %nin% c(zygotic_enriched_inherited_genes, maternally_enriched_inherited_genes),])
non_zygotic_enriched_inherited_genes = row.names(two_four_de[row.names(two_four_de) %nin% c(zygotic_enriched_inherited_genes),])


#maternally_enriched_inherited_genes_wb = wb_id_common_names[wb_id_common_names$V2 %in% maternally_enriched_inherited_genes, 1]
zygotic_enriched_inherited_genes_wb = wb_id_common_names[wb_id_common_names$V2 %in% zygotic_enriched_inherited_genes, 1]
#non_biased_enriched_inherited_genes_wb = wb_id_common_names[wb_id_common_names$V2 %in% non_biased_enriched_inherited_genes, 1]
non_zygotic_enriched_inherited_genes_wb = wb_id_common_names[wb_id_common_names$V2 %in% non_zygotic_enriched_inherited_genes, 1]

# write.table(inherited_promoters_genes_only, file = "inherited_zygotic_program/4cell_promoters.inherited.specific.genes", append = F, quote = F, row.names = F, col.names = "Gene_name")
# write.table(zygotic_promoters_genes_only, file = "inherited_zygotic_program/4cell_promoters.zygotic.specific.genes", append = F, quote = F, row.names = F, col.names = "Gene_name")
# write.table(inh_zyg_promoters_genes, file = "inherited_zygotic_program/4cell_promoters.inherited.zygotic.genes", append = F, quote = F, row.names = F, col.names = "Gene_name")
# write.table(maternally_enriched_inherited_genes_wb, file = "inherited_zygotic_program/4cell_promoters.inherited.maternal_bias.genes", append = F, quote = F, row.names = F, col.names = "Gene_name")
# write.table(zygotic_enriched_inherited_genes_wb, file = "inherited_zygotic_program/4cell_promoters.inherited.zygotic_bias.genes", append = F, quote = F, row.names = F, col.names = "Gene_name")
# write.table(non_biased_enriched_inherited_genes_wb, file = "inherited_zygotic_program/4cell_promoters.inherited.no_bias.genes", append = F, quote = F, row.names = F, col.names = "Gene_name")

# plot gene expression: GL tomography
gl_tomography = read.table("data/external_data/gonad_RNAseq_Colaiacovo.txt")

# gl_tomography_inh_zyg = gl_tomography[gl_tomography$V16 %in% maternally_enriched_inherited_genes_wb,c(2:11)]
# gl_tomography_inh_zyg = rbind(gl_tomography_inh_zyg, gl_tomography[gl_tomography$V16 %in% non_biased_enriched_inherited_genes_wb,c(2:11)])
gl_tomography_inh_zyg = gl_tomography[gl_tomography$V16 %in% non_zygotic_enriched_inherited_genes_wb,c(2:11)]
gl_tomography_inh_zyg = rbind(gl_tomography_inh_zyg, gl_tomography[gl_tomography$V16 %in% zygotic_enriched_inherited_genes_wb,c(2:11)])
gl_tomography_inh_zyg = rbind(gl_tomography_inh_zyg, gl_tomography[gl_tomography$V16 %in% zygotic_promoters_genes,c(2:11)])

gl_tomography_inh_zyg_heatmap = Heatmap(as.matrix(log10(gl_tomography_inh_zyg+1)), name = "inherited/zygotic gene exp", cluster_columns = F, 
                                        cluster_rows = T, use_raster = F, row_dend_reorder = TRUE, row_title_rot = 0, column_title_rot = 90, 
                                        column_title_gp = gpar(cex = 0.8), row_title_gp = gpar(cex = 0.8), show_row_dend = F, show_row_names = F,
                                        row_split = c(rep("inh - no zyg bias", dim(gl_tomography[gl_tomography$V16 %in% non_zygotic_enriched_inherited_genes_wb,c(2:11)])[1]),
                                                      rep("inh - zygotic", dim(gl_tomography[gl_tomography$V16 %in% zygotic_enriched_inherited_genes_wb,c(2:11)])[1]),
                                                      rep("zygotic", dim(gl_tomography[gl_tomography$V16 %in% zygotic_promoters_genes,c(2:11)])[1])))
gl_tomography_inh_zyg_heatmap = draw(gl_tomography_inh_zyg_heatmap, padding = unit(c(8, 5, 10, 4), "mm"))

pdf(file=paste0(plot_dir_w_date, "/gene_expression_inherited_zygotic_promoters.tomography.pdf"), width=12, height = 6, useDingbats = F)
par(mar=c(8, 5, 3, 3), xpd=TRUE)
boxplot(log10(gl_tomography[gl_tomography$V16 %in% inherited_promoters_genes,c(2:11)] + 1), notch=T, at = seq(1,28,by=3), xaxt = "n", las = 1, main = "gene expression adult germline", xlim = c(0, 30), col = "deepskyblue", ylab = "log10(TPM + 1)")
boxplot(log10(gl_tomography[gl_tomography$V16 %in% zygotic_promoters_genes,c(2:11)] + 1), add = T, yaxt = "n", xaxt = "n", notch=T, at = seq(2,29,by=3), col = "darkviolet")
axis(1, at = seq(1.5,28.5,by=3), labels = c("distal end", rep("", 8), "proximal end"))
legend(15, -0.5, xjust = 0.5, legend=c("inherited","zygotic"), fill = c("deepskyblue", "darkviolet"), title="Gene promoter", bty = "n", ncol = 2)

boxplot(log10(gl_tomography[gl_tomography$V16 %in% non_zygotic_enriched_inherited_genes_wb,c(2:11)] + 1), notch=T, at = seq(1,37,by=4), xaxt = "n", las = 1, main = "gene expression adult germline", xlim = c(0, 40), col = "deepskyblue", ylab = "log10(TPM + 1)")
#boxplot(log10(gl_tomography[gl_tomography$V16 %in% non_biased_enriched_inherited_genes_wb,c(2:11)] + 1), add = T, yaxt = "n", xaxt = "n", notch=T, at = seq(2,47,by=5), col = "deepskyblue3")
boxplot(log10(gl_tomography[gl_tomography$V16 %in% zygotic_enriched_inherited_genes_wb,c(2:11)] + 1), add = T, yaxt = "n", xaxt = "n", notch=T, at = seq(2,38,by=4), col = "deepskyblue4")
boxplot(log10(gl_tomography[gl_tomography$V16 %in% zygotic_promoters_genes,c(2:11)] + 1), add = T, yaxt = "n", xaxt = "n", notch=T, at = seq(3,39,by=4), col = "darkviolet")
axis(1, at = seq(2.5,38.5,by=4), labels = c("distal end", rep("", 8), "proximal end"))
legend(25, -0.5, xjust = 0.5, legend=c("inherited - no zygotic bias", "inherited - zygotic bias", "zygotic"), fill = c("deepskyblue", "deepskyblue4", "darkviolet"), title="Gene promoter", bty = "n", ncol = 3)

gl_tomography_inh_zyg_heatmap
dev.off()

# adult germline
combined_gl_exp = read.table("data/external_data/adult_gl.tsv")

# gl_adult_inh_zyg = combined_gl_exp[row.names(combined_gl_exp) %in% maternally_enriched_inherited_genes_wb,c(2:4)]
# gl_adult_inh_zyg = rbind(gl_adult_inh_zyg, combined_gl_exp[row.names(combined_gl_exp) %in% non_biased_enriched_inherited_genes_wb,c(2:4)])
gl_adult_inh_zyg = combined_gl_exp[row.names(combined_gl_exp) %in% non_zygotic_enriched_inherited_genes_wb,c(2:4)]
gl_adult_inh_zyg = rbind(gl_adult_inh_zyg, combined_gl_exp[row.names(combined_gl_exp) %in% zygotic_enriched_inherited_genes_wb,c(2:4)])
gl_adult_inh_zyg = rbind(gl_adult_inh_zyg, combined_gl_exp[row.names(combined_gl_exp) %in% zygotic_promoters_genes,c(2:4)])

gl_adult_inh_zyg_heatmap = Heatmap(as.matrix(log10(gl_adult_inh_zyg+1)), name = "inherited/zygotic gene exp", cluster_columns = F, 
                                   cluster_rows = T, use_raster = F, row_dend_reorder = TRUE, row_title_rot = 0, column_title_rot = 90, 
                                   column_title_gp = gpar(cex = 0.8), row_title_gp = gpar(cex = 0.8), show_row_dend = F, show_row_names = F,
                                   row_split = c(rep("inh - no zyg bias", dim(combined_gl_exp[row.names(combined_gl_exp) %in% non_zygotic_enriched_inherited_genes_wb,])[1]),
                                                 rep("inh - zyg bias", dim(combined_gl_exp[row.names(combined_gl_exp) %in% zygotic_enriched_inherited_genes_wb,])[1]),
                                                 rep("zygotic", dim(combined_gl_exp[row.names(combined_gl_exp) %in% zygotic_promoters_genes,])[1])))
gl_adult_inh_zyg_heatmap = draw(gl_adult_inh_zyg_heatmap, padding = unit(c(8, 5, 10, 4), "mm"))

pdf(file=paste0(plot_dir_w_date, "/gene_expression_inherited_zygotic_promoters.scRNAseq.pdf"), width=6, height = 6, useDingbats = F)
par(mfrow = c(1,1), mar=c(8, 5, 3, 3), xpd=TRUE)
boxplot(log10(combined_gl_exp[row.names(combined_gl_exp) %in% inherited_promoters_genes,c(2:4)] + 1), notch=T, at = c(1,4,7), xaxt = "n", las = 1, main = "gene expression adult germline", xlim = c(0, 9), col = "deepskyblue", ylab = "log10(TPM + 1)")
boxplot(log10(combined_gl_exp[row.names(combined_gl_exp) %in% zygotic_promoters_genes,c(2:4)] + 1), add = T, yaxt = "n", xaxt = "n", notch=T, at = c(2,5,8), col = "darkviolet")
axis(1, at = c(1.5,4.5,7.5), labels = c("mitotic", "meiotic", "oocyte"))
legend(4.5, -1, xjust = 0.5, legend=c("inherited","zygotic"), fill = c("deepskyblue", "darkviolet"), title="Gene promoter", bty = "n", ncol = 2)

# boxplot(log10(combined_gl_exp[row.names(combined_gl_exp) %in% maternally_enriched_inherited_genes_wb,c(2:4)] + 1), notch=T, at = c(1,6,11), xaxt = "n", las = 1, main = "gene expression adult germline", xlim = c(0, 15), col = "deepskyblue", ylab = "log10(TPM + 1)")
# boxplot(log10(combined_gl_exp[row.names(combined_gl_exp) %in% non_biased_enriched_inherited_genes_wb,c(2:4)] + 1), add = T, yaxt = "n", xaxt = "n", notch=T, at = c(2,7,12), col = "deepskyblue3")
boxplot(log10(combined_gl_exp[row.names(combined_gl_exp) %in% non_zygotic_enriched_inherited_genes_wb,c(2:4)] + 1), notch=T, at = c(1,5,9), xaxt = "n", las = 1, main = "gene expression adult germline", xlim = c(0, 12), col = "deepskyblue", ylab = "log10(TPM + 1)")
boxplot(log10(combined_gl_exp[row.names(combined_gl_exp) %in% zygotic_enriched_inherited_genes_wb,c(2:4)] + 1), add = T, yaxt = "n", xaxt = "n", notch=T, at = c(2,6,10), col = "deepskyblue4")
boxplot(log10(combined_gl_exp[row.names(combined_gl_exp) %in% zygotic_promoters_genes,c(2:4)] + 1), add = T, yaxt = "n", xaxt = "n", notch=T, at = c(3,7,11), col = "darkviolet")
axis(1, at = c(2,6,10), labels = c("mitotic", "meiotic", "oocyte"))
legend(6, -1, xjust = 0.5, legend=c("inherited - no zygotic bias", "inherited - zygotic bias", "zygotic"), fill = c("deepskyblue", "deepskyblue4", "darkviolet"), title="Gene promoter", bty = "n", ncol = 3)

gl_adult_inh_zyg_heatmap
dev.off()

# look into CelSeq data
cole_exp = read.table("data/external_data/Yanai_single_cell.tsv", header=T)

# celseq_inh_zyg = cole_exp[cole_exp$WB_ID %in% maternally_enriched_inherited_genes_wb,c(3:5)]
# celseq_inh_zyg = rbind(celseq_inh_zyg, cole_exp[cole_exp$WB_ID %in% non_biased_enriched_inherited_genes_wb,c(3:5)])
celseq_inh_zyg = cole_exp[cole_exp$WB_ID %in% non_zygotic_enriched_inherited_genes_wb,c(3:5)]
celseq_inh_zyg = rbind(celseq_inh_zyg, cole_exp[cole_exp$WB_ID %in% zygotic_enriched_inherited_genes_wb,c(3:5)])
celseq_inh_zyg = rbind(celseq_inh_zyg, cole_exp[cole_exp$WB_ID %in% zygotic_promoters_genes,c(3:5)])

celseq_inh_zyg_heatmap = Heatmap(as.matrix(log10(celseq_inh_zyg+1)), name = "inherited/zygotic gene exp", cluster_columns = F, 
                                 cluster_rows = T, use_raster = F, row_dend_reorder = TRUE, row_title_rot = 0, column_title_rot = 90, 
                                 column_title_gp = gpar(cex = 0.8), row_title_gp = gpar(cex = 0.8), show_row_dend = F, show_row_names = F,
                                 row_split = c(rep("inh - non zygotic", dim(cole_exp[cole_exp$WB_ID %in% non_zygotic_enriched_inherited_genes_wb,c(3:5)])[1]),
                                               rep("inh - zygotic", dim(cole_exp[cole_exp$WB_ID %in% zygotic_enriched_inherited_genes_wb,c(3:5)])[1]),
                                               rep("zygotic", dim(cole_exp[cole_exp$WB_ID %in% zygotic_promoters_genes,c(3:5)])[1])))
celseq_inh_zyg_heatmap = draw(celseq_inh_zyg_heatmap, padding = unit(c(8, 5, 10, 4), "mm"))

pdf(file=paste0(plot_dir_w_date, "/gene_expression_inherited_zygotic_promoters.preZGA_CELSeq.pdf"), width=6, height = 6, useDingbats = F)
par(mar=c(8, 5, 3, 3), xpd=TRUE)
boxplot(log10(cole_exp[cole_exp$WB_ID %in% inherited_promoters_genes,c(3:5)] + 1), notch=T, at = c(1,4,7), xaxt = "n", las = 1, main = "gene expression pre-ZGA", xlim = c(0, 9), col = "deepskyblue", ylab = "log10(TPM + 1)")
boxplot(log10(cole_exp[cole_exp$WB_ID %in% zygotic_promoters_genes,c(3:5)] + 1), add = T, yaxt = "n", xaxt = "n", notch=T, at = c(2,5,8), col = "darkviolet")
axis(1, at = c(1.5,4.5,7.5), labels = c("P0", "AB", "P1"))
legend(4.5, -1, xjust = 0.5, legend=c("inherited", "zygotic"), fill = c("deepskyblue", "darkviolet"), title="Gene promoter", bty = "n", ncol = 2)

boxplot(log10(cole_exp[cole_exp$WB_ID %in% non_zygotic_enriched_inherited_genes_wb,c(3:5)] + 1), notch=T, at = c(1,5,9), xaxt = "n", las = 1, main = "gene expression pre-ZGA", xlim = c(0, 12), col = "deepskyblue", ylab = "log10(TPM + 1)")
boxplot(log10(cole_exp[cole_exp$WB_ID %in% zygotic_enriched_inherited_genes_wb,c(3:5)] + 1), add = T, yaxt = "n", xaxt = "n", notch=T, at = c(2,6,10), col = "deepskyblue4")
boxplot(log10(cole_exp[cole_exp$WB_ID %in% zygotic_promoters_genes,c(3:5)] + 1), add = T, yaxt = "n", xaxt = "n", notch=T, at = c(3,7,11), col = "darkviolet")
axis(1, at = c(2,6,10), labels = c("P0", "AB", "P1"))
legend(6, -1, xjust = 0.5, legend=c("inherited - no zygotic bias","inherited - zygotic bias", "zygotic"), fill = c("deepskyblue", "deepskyblue4", "darkviolet"), title="Gene promoter", bty = "n", ncol = 3)

celseq_inh_zyg_heatmap
dev.off()

# expression in 10x data
combined_seurat_avgExp_stage_all = as.data.frame(AverageExpression(combined_seurat, layer = "data", assays = "RNA", group.by = "stage"))
combined_seurat_avgExp_stage_all = cbind(combined_seurat_avgExp_stage_all[,c(8,4,6,1,2,3,5,7)])
names(combined_seurat_avgExp_stage_all) = c("2cell", "4cell", "8cell", "15cell", "26cell", "46cell", "87cell", "Pcell")


pdf(file=paste0(plot_dir_w_date, "/gene_expression_inherited_zygotic_promoters.10x_data.pdf"), width=18, height = 6, useDingbats = F)
par(mfrow=c(1,2))
boxplot(log10(combined_seurat_avgExp_stage_all[row.names(combined_seurat_avgExp_stage_all) %in% inherited_promoters_genes_name,]+1), notch = T, ylim = c(0, 2), main = paste0("inherited (N = ", length(inherited_promoters_genes_name), ")"), las = 1, ylab = "log10(avg exp + 1)", col = "deepskyblue")
boxplot(log10(combined_seurat_avgExp_stage_all[row.names(combined_seurat_avgExp_stage_all) %in% zygotic_promoters_genes_name,]+1), notch = T, ylim = c(0, 2), main = paste0("zygotic (N = ", length(zygotic_promoters_genes_name), ")"), las = 1, ylab = "log10(avg exp + 1)", col = "deepskyblue4")

par(mfrow=c(1,3))
boxplot(log10(combined_seurat_avgExp_stage_all[row.names(combined_seurat_avgExp_stage_all) %in% non_zygotic_enriched_inherited_genes,]+1), notch = T, ylim = c(0, 2), main = paste0("inherited - no zygotic bias (N = ", length(non_zygotic_enriched_inherited_genes), ")"), las = 1, ylab = "log10(avg exp + 1)", col = "deepskyblue")
boxplot(log10(combined_seurat_avgExp_stage_all[row.names(combined_seurat_avgExp_stage_all) %in% zygotic_enriched_inherited_genes,]+1), notch = T, ylim = c(0, 2), main = paste0("inherited - zygotic bias (N = ", length(zygotic_enriched_inherited_genes), ")"), las = 1, ylab = "log10(avg exp + 1)", col = "deepskyblue4")
boxplot(log10(combined_seurat_avgExp_stage_all[row.names(combined_seurat_avgExp_stage_all) %in% zygotic_promoters_genes_name,]+1), notch = T, ylim = c(0, 2), main = paste0("zygotic (N = ", length(zygotic_promoters_genes_name), ")"), las = 1, ylab = "log10(avg exp + 1)", col = "darkviolet")
dev.off()

# exclude from the zygotic set the genes with any expression in P0, P1 and/or AB in Cole
# maternal_cole = wb_id_common_names[wb_id_common_names$V1 %in% cole_exp[cole_exp$P0 > 0 | cole_exp$AB > 0 | cole_exp$P1 > 0, 2],2]
# 
# zygotic_promoters_genes_name_no_cole = zygotic_promoters_genes_name[zygotic_promoters_genes_name %nin% maternal_cole]
# 
# pdf(file=paste0(plot_dir_w_date, "/gene_expression_inherited_zygotic_promoters.10x_data.zyg_no_cole.pdf"), width=18, height = 6, useDingbats = F)
# par(mfrow=c(1,2))
# boxplot(log10(combined_seurat_avgExp_stage_all[row.names(combined_seurat_avgExp_stage_all) %in% inherited_promoters_genes_name,]+1), notch = T, ylim = c(0, 2), main = paste0("inherited (N = ", length(inherited_promoters_genes_name), ")"), las = 1, ylab = "log10(avg exp + 1)", col = "deepskyblue")
# boxplot(log10(combined_seurat_avgExp_stage_all[row.names(combined_seurat_avgExp_stage_all) %in% zygotic_promoters_genes_name_no_cole,]+1), notch = T, ylim = c(0, 2), main = paste0("zygotic (N = ", length(zygotic_promoters_genes_name_no_cole), ")"), las = 1, ylab = "log10(avg exp + 1)", col = "deepskyblue4")
# 
# par(mfrow=c(1,4))
# boxplot(log10(combined_seurat_avgExp_stage_all[row.names(combined_seurat_avgExp_stage_all) %in% maternally_enriched_inherited_genes,]+1), notch = T, ylim = c(0, 2), main = paste0("inherited - maternal bias (N = ", length(maternally_enriched_inherited_genes), ")"), las = 1, ylab = "log10(avg exp + 1)", col = "deepskyblue")
# boxplot(log10(combined_seurat_avgExp_stage_all[row.names(combined_seurat_avgExp_stage_all) %in% non_biased_enriched_inherited_genes,]+1), notch = T, ylim = c(0, 2), main = paste0("inherited - no bias (N = ", length(non_biased_enriched_inherited_genes), ")"), las = 1, ylab = "log10(avg exp + 1)", col = "deepskyblue3")
# boxplot(log10(combined_seurat_avgExp_stage_all[row.names(combined_seurat_avgExp_stage_all) %in% zygotic_enriched_inherited_genes,]+1), notch = T, ylim = c(0, 2), main = paste0("inherited - zygotic bias (N = ", length(zygotic_enriched_inherited_genes), ")"), las = 1, ylab = "log10(avg exp + 1)", col = "deepskyblue4")
# boxplot(log10(combined_seurat_avgExp_stage_all[row.names(combined_seurat_avgExp_stage_all) %in% zygotic_promoters_genes_name_no_cole,]+1), notch = T, ylim = c(0, 2), main = paste0("zygotic (N = ", length(zygotic_promoters_genes_name_no_cole), ")"), las = 1, ylab = "log10(avg exp + 1)", col = "darkviolet")
# dev.off()
# 
```
