## Supplementary material for "A lineage-resolved multimodal single-cell atlas reveals the genomic dynamics of early *C. elegans* development": Code and external data: inherited_zygotic_promoters.motif_enrichment.20241119.R

```
# plot relative enrichment of motifs in inherited and zygotic sites
require(png)

current_date= paste(unlist(strsplit(as.character(Sys.Date()), "-")), collapse="")
plot_dir_w_date = paste0("paper_figures/", current_date)
dir.create(plot_dir_w_date, recursive=TRUE)

graphical_pvalue <- function(x){
  x[x < 0.001] = "***"
  x[x > 0.001 & x < 0.01] = "**"
  x[x > 0.01 & x < 0.05] = "*"
  x[x >= 0.05] = "ns"
  print(x)
}

# read in the enrichment table
all_motifs_to_invert = read.table("inherited_zygotic_program/memechip/4cell_promoters.inherited.zygotic.processed/motifs_enriched.all.enrichment.txt")

# invert the enrichment table (to plot top to bottom)
all_motifs_to_test_to_order = all_motifs_to_invert[c(dim(all_motifs_to_invert)[1]:1),]

# order motif table: split between motifs enriched in inherited and zygotic, then order from higher fraction of inherited sites to higher fraction of zygotic sites
all_motifs_to_test_inherited = all_motifs_to_test_to_order[all_motifs_to_test_to_order$inherited_ratio/all_motifs_to_test_to_order$zygotic_ratio > 1,]
all_motifs_to_test_zygotic = all_motifs_to_test_to_order[all_motifs_to_test_to_order$inherited_ratio/all_motifs_to_test_to_order$zygotic_ratio < 1,]

all_motifs_to_test = rbind(all_motifs_to_test_zygotic[order(all_motifs_to_test_zygotic$zygotic_ratio, decreasing = T),],
                           all_motifs_to_test_inherited[order(all_motifs_to_test_inherited$inherited_ratio),])


# calculate the significance of the difference in enrichment with a chisq.test
all_motifs_chisq = p.adjust(apply(all_motifs_to_test[,c(1,2,4,5)], 1, function(x){chisq.test(matrix(as.numeric(x), ncol=2, byrow = T))$p.value}), method = "BH")

# only retain motifs with a significant difference
all_motifs = all_motifs_to_test[which(all_motifs_chisq < 0.05),]

# plot enrichment
pdf(file=paste0(plot_dir_w_date, "/all_promoters_motifs.enrichment.pdf"), width=12, height = 10, useDingbats = F)
par(mar=c(10, 3, 4, 4), xpd=TRUE, mfrow = c(1,3))
k = barplot(t(matrix(c(all_motifs$inherited_ratio, all_motifs$zygotic_ratio), ncol = 2, byrow = F)), beside = T, horiz = T, plot = F)
plot(1, type="n", xlab="", ylab="", xlim=c(0, 10), ylim=c(1, k[2, dim(k)[2] + 0.5]), xaxt = "n", yaxt = "n", bty = "n")
# read in sequentially all fwd/rev logos and add them to the left of the barplot
for (motif_n in c(1:dim(k)[2])){
  motif_name = row.names(all_motifs)[motif_n]
  img_fwd<-readPNG(paste0("inherited_zygotic_program/memechip/4cell_promoters.inherited.zygotic.processed/motif_clusters/", motif_name, ".merged_motifs.mafft_out.logo.fwd.png"))
  img_rev<-readPNG(paste0("inherited_zygotic_program/memechip/4cell_promoters.inherited.zygotic.processed/motif_clusters/", motif_name, ".merged_motifs.mafft_out.logo.rev.png"))
  rasterImage(img_fwd,1, k[1,motif_n]-0.5, 4, k[2,motif_n]+0.5)
  rasterImage(img_rev,5, k[1,motif_n]-0.5, 9, k[2,motif_n]+0.5)
}
barplot(t(matrix(c(all_motifs$inherited_ratio, all_motifs$zygotic_ratio), ncol = 2, byrow = F)), beside = T, horiz = T, 
            main = "all_motifs motifs - enrichment", col = c("darkviolet", "deepskyblue3"), xlab = "fraction of promoters with motif")
legend(0, 0-ceiling(k[2,dim(k)[2]]/10), xjust = 0, legend=c("inherited", "zygotic"), fill = c("darkviolet", "deepskyblue3"), bty = "n", ncol = 2)

# plot ratio (use rect to specify bar location)
ratio_all_inh_zyg = t(matrix(all_motifs$inherited_ratio/all_motifs$zygotic_ratio, ncol = 1))
ratio_all_zyg_inh = t(matrix(all_motifs$zygotic_ratio/all_motifs$inherited_ratio, ncol = 1))

ratio_all = ratio_all_inh_zyg
ratio_all[ratio_all_inh_zyg < 1] = 0-ratio_all_zyg_inh[ratio_all_inh_zyg < 1]

ratio_all_left = ratio_all
ratio_all_left[ratio_all_left > 0] = 0
ratio_all_right = ratio_all
ratio_all_right[ratio_all_right < 0] = 0

barplot_col = as.character(ratio_all_left)
barplot_col[barplot_col == "0"] = "darkviolet"
barplot_col[barplot_col != "darkviolet"] = "deepskyblue3"

plot(1, ylim=c(1, k[2, dim(k)[2] + 0.5]), xlim = c(round(min(ratio_all) - 0.1, 1), round(max(ratio_all) + 0.1, 1)), 
     type = "n", bty = "n", yaxt = "n", xlab = "fold change", xaxt = "n")
axis(1, at = seq(round(min(ratio_all)), round(max(ratio_all)), by=1), labels = abs(seq(round(min(ratio_all)), round(max(ratio_all)))))
rect(xleft= ratio_all_left, ybottom=k[1,], xright=ratio_all_right, ytop=k[2,], col=barplot_col)
text(ratio_all_right, k[1,] + 0.5, labels = graphical_pvalue(all_motifs_chisq[which(all_motifs_chisq < 0.05)]), pos = 4)
abline(v=0, lty = 2, lwd = 2, col = 2, xpd = F)
dev.off()
```
