## Supplementary material for "A lineage-resolved multimodal single-cell atlas reveals the genomic dynamics of early *C. elegans* development": Code and external data: lcap_alignment.20240228.sh

```
#!/bin/bash
#
#SBATCH --job-name=STAR
#SBATCH --output=slurm_out/STAR.%N.%j.out
#SBATCH --error=slurm_err/STAR.%N.%j.err
#SBATCH --ntasks=1
#SBATCH -N 1
#SBATCH -n 12
#SBATCH --mem-per-cpu=8000
#SBATCH --mail-type=END,FAIL
#SBATCH --mail-user=

genome_index=$1
sample_name=$2
max_frag_size=$3

mkdir data/lcap/alignment
blacklist=data/external_data/ce11-blacklist.bed
chr_size=species/elegans/genome/elegans.chrom.sizes.txt

if [ ! -f "data/lcap/fastq_trimmed/"$sample_name"_L001_R1_001_val_1.fq.gz" ]
then
    trim_galore --gzip --trim-n -j 4 --paired -o data/lcap/fastq_trimmed "data/lcap/fastq/"$sample_name"_L001_R1_001.fastq.gz" "data/lcap/fastq/"$sample_name"_L001_R2_001.fastq.gz"
fi


STAR --genomeDir star_idx/$genome_index --readFilesIn "data/lcap/fastq_trimmed/"$sample_name"_L001_R1_001_val_1.fq.gz" "data/lcap/fastq_trimmed/"$sample_name"_L001_R2_001_val_2.fq.gz" --readFilesCommand zcat --runThreadN 12 --outFileNamePrefix "data/lcap/alignment/"$sample_name"_" --outSAMtype BAM SortedByCoordinate --outWigType wiggle --twopassMode None --limitBAMsortRAM 6274374343 --outSAMattrIHstart 0 --outReadsUnmapped Fastx --alignIntronMax 20000 

# only keep uniquely mapped reads
samtools index "data/lcap/alignment/"$sample_name"_Aligned.sortedByCoord.out.bam"
samtools view -b -q 255 "data/lcap/alignment/"$sample_name"_Aligned.sortedByCoord.out.bam" > "data/lcap/alignment/"$sample_name"_Aligned.sortedByCoord.out.unique.bam"
samtools index "data/lcap/alignment/"$sample_name"_Aligned.sortedByCoord.out.unique.bam"

# output tracks of unique and multimapping reads
bamCoverage --bam "data/lcap/alignment/"$sample_name"_Aligned.sortedByCoord.out.unique.bam" -o "data/lcap/alignment/"$sample_name"_Aligned.unique.fwd.bw" --filterRNAstrand forward -bs 1 -bl $blacklist -p 12 --effectiveGenomeSize 100286401 --normalizeUsing BPM -e
bamCoverage --bam "data/lcap/alignment/"$sample_name"_Aligned.sortedByCoord.out.unique.bam" -o "data/lcap/alignment/"$sample_name"_Aligned.unique.rev.bw" --filterRNAstrand reverse -bs 1 -bl $blacklist -p 12 --effectiveGenomeSize 100286401 --normalizeUsing BPM -e
bamCoverage --bam "data/lcap/alignment/"$sample_name"_Aligned.sortedByCoord.out.bam" -o "data/lcap/alignment/"$sample_name"_Aligned.fwd.bw" --filterRNAstrand forward -bs 1 -bl $blacklist -p 12 --effectiveGenomeSize 100286401 --normalizeUsing BPM -e
bamCoverage --bam "data/lcap/alignment/"$sample_name"_Aligned.sortedByCoord.out.bam" -o "data/lcap/alignment/"$sample_name"_Aligned.rev.bw" --filterRNAstrand reverse -bs 1 -bl $blacklist -p 12 --effectiveGenomeSize 100286401 --normalizeUsing BPM -e

# convert bam file into bed, then output bedgraph file or fragments - used in promoter annotation
samtools sort -n "data/lcap/alignment/"$sample_name"_Aligned.sortedByCoord.out.unique.bam" | bamToBed -i stdin -bedpe -mate1 | awk 'BEGIN{OFS="\t";}{if ($2 < $5) print $1, $2, $6, $7, 0, "-"; else print $1, $5, $3, $7, 0, "+"}' | awk -v var="$max_frag_size" '{if ($3 - $2 < var) print}' | sort -k 1,1 -k2,2n > "data/lcap/alignment/"$sample_name"_Aligned.sortedByCoord.out.unique.fragments.bed"
genomeCoverageBed -i "data/lcap/alignment/"$sample_name"_Aligned.sortedByCoord.out.unique.fragments.bed" -bga -strand + -g $chr_size > "data/lcap/alignment/"$sample_name"_Aligned.sortedByCoord.out.unique.fragments.fwd.bg"
genomeCoverageBed -i "data/lcap/alignment/"$sample_name"_Aligned.sortedByCoord.out.unique.fragments.bed" -bga -strand - -g $chr_size > "data/lcap/alignment/"$sample_name"_Aligned.sortedByCoord.out.unique.fragments.rev.bg"
```
