## Supplementary material for "A lineage-resolved multimodal single-cell atlas reveals the genomic dynamics of early *C. elegans* development": Code and external data: lcap_DESeq_jump.R

```
library("DESeq2")
args = commandArgs(trailingOnly=TRUE)

input_counts = args[1]
deseq_out=args[2]

set.seed(42)

countData <- read.table(input_counts, header=TRUE, sep="\t", row.names=1)
colData <- data.frame(row.names=colnames(countData),
                      condition = relevel(factor(c("cond1", "cond1", "cond2", "cond2")), "cond1"))
dds <- DESeqDataSetFromMatrix(countData=countData, colData=colData, design=~condition)
dds <- DESeq(dds)
res <- results(dds, altHypothesis="greater")

write.table(res, deseq_out, quote = F, row.names=T, col.names=T, sep="\t")
```
