## Supplementary material for "A lineage-resolved multimodal single-cell atlas reveals the genomic dynamics of early *C. elegans* development": Code and external data: left_right_coverage_combined.py

```
import sys

input_file = sys.argv[1]
output_file = sys.argv[2]
strand = sys.argv[3]

new = open(output_file, "w")
new.write("rep1_left\trep2_left\trep1_right\trep2_right\n")

temp = []
for line in open(input_file):
    if temp == []:
        temp = line.rstrip().split("\t")
    else:
        K=line.rstrip().split("\t")
        temp += [K[4], K[5]]
        if strand == "fwd":
            new.write("embryo_AS_" + temp[3].split("_")[2] + "\t" + temp[4] + "\t" + temp[5] + "\t" + temp[6] + "\t" + temp[7] + "\n")
        else:
            new.write("embryo_AS_" + temp[3].split("_")[2] + "\t" + temp[6] + "\t" + temp[7] + "\t" + temp[4] + "\t" + temp[5] + "\n")
        temp = []

new.close()
```
