## Supplementary material for "A lineage-resolved multimodal single-cell atlas reveals the genomic dynamics of early *C. elegans* development": Code and external data: lineage_specific_annotation.20240207.R

```
# annotate cell types in Seurat object based on expression of specific markers
# requires subclustering and DE test
library(Seurat)
library(Signac)
library(ggplot2)
library(multimode)
library(Hmisc)
library(stringr)

source("scripts/second_derivative_binarisation.R")

current_date= paste(unlist(strsplit(as.character(Sys.Date()), "-")), collapse="")
plot_dir_w_date = paste0("plots/", current_date)
dir.create(plot_dir_w_date, recursive=TRUE)

args = commandArgs(trailingOnly=TRUE)
seurat_object_file = args[1]
output_dir = args[2]
annotation = args[3]

### scoring function
scoring_markers = function(x, y){
  score_cluster_cell = length(x)
  malus_value = score_cluster_cell * 0.1
  kill_value = malus_value * 5
  for (marker_gene in c(1:length(x))){
    if (x[marker_gene] == y[marker_gene]){
      score_cluster_cell = score_cluster_cell
    }
    else if (abs(x[marker_gene] - y[marker_gene]) == 2){
      score_cluster_cell = score_cluster_cell - kill_value
    }
    else if (abs(x[marker_gene] - y[marker_gene]) == 1){
      score_cluster_cell = score_cluster_cell - malus_value
    }
  }
  return(score_cluster_cell)
}

combined_seurat = readRDS(seurat_object_file)

# reduce resolution to facilitate identification of AB, E, MS, C and D lineages based on marker genes
combined_seurat <- FindClusters(combined_seurat, resolution = 0.5, graph.name = "wsnn")
combined_seurat_UMAP = DimPlot(combined_seurat, reduction = "umapWNN", label = T) & NoLegend()

AB_mkrs = c("sea-2","T25E12.6","sre-45")
E_MS_C_D_mkrs = c("F28H6.8","elt-7","tbx-35", "ceh-51", "hnd-1", "nlp-11", "cbs-1")

# find clusters corresponding to the AB or E/MS/C/D lineages based on enrichment of specific markers
if (file.exists(paste0(output_dir, "/all_clusters_markers.", annotation, ".txt"))){
  all_mrkrs = read.table(paste0(output_dir, "/all_clusters_markers.", annotation, ".txt"))
  } else {
    all_mrkrs = FindAllMarkers(combined_seurat, assay = "SCT", logfc.threshold = 0.5, only.pos = T)
    write.table(all_mrkrs, file = paste0(output_dir, "/all_clusters_markers.", annotation, ".txt"), quote = F, sep = "\t", row.names = T, col.names = T)
  }

AB_clusters = unique(all_mrkrs[all_mrkrs$gene %in% c("sea-2","T25E12.6","sre-45"),6])
E_MS_C_D_clusters = unique(all_mrkrs[all_mrkrs$gene %in% c("F28H6.8","elt-7","tbx-35", "ceh-51", "hnd-1", "nlp-11", "cbs-1"),6])

pdf(file=paste0(plot_dir_w_date, "/early_linege_markers.", annotation, ".pdf"), width=18, height = 9)
print(FeaturePlot(combined_seurat, reduction = "umapWNN", order = T, features = AB_mkrs, ncol = 3)  & 
  scale_colour_gradientn(colours = c("grey90", "cyan", "deepskyblue", "forestgreen", "darkorange", "darkred")))
print(FeaturePlot(combined_seurat, reduction = "umapWNN", order = T, features = E_MS_C_D_mkrs, ncol = 4)  & 
  scale_colour_gradientn(colours = c("grey90", "cyan", "deepskyblue", "forestgreen", "darkorange", "darkred")))
AB_UMAP = DimPlot(combined_seurat, reduction = "umapWNN", label = T, sizes.highlight = 0.4, pt.size = 0.4, cells.highlight = 
          colnames(combined_seurat[,combined_seurat$seurat_clusters %in% AB_clusters])) & NoLegend()
E_MS_C_D_UMAP = DimPlot(combined_seurat, reduction = "umapWNN", label = T, sizes.highlight = 0.4, pt.size = 0.4, cells.highlight = 
          colnames(combined_seurat[,combined_seurat$seurat_clusters %in% E_MS_C_D_clusters])) & NoLegend()
AB_UMAP | E_MS_C_D_UMAP
dev.off()

lineage_clusters = list("AB" = AB_clusters, "E_MS_C_D" = E_MS_C_D_clusters)

# annotate AB and E_MS_C_D lineages
for (lineage in c("AB", "E_MS_C_D")){
  markers_table = read.table(paste0("data/external_data/markers_UMAP_annotation.", lineage, ".20240207.tsv"))
  # subcluster the lineage
  combined_seurat_subcluster = subset(combined_seurat, idents = lineage_clusters[[lineage]])
  DefaultAssay(combined_seurat_subcluster) = "WNN"
  combined_seurat_subcluster <- FindMultiModalNeighbors(combined_seurat_subcluster, reduction.list = list("pca", "lsi"), dims.list = list(1:50, 2:50), weighted.nn.name = "clusterwnn")
  combined_seurat_subcluster <- FindClusters(combined_seurat_subcluster, resolution = 5.5, graph.name = "wsnn")
  combined_seurat_subcluster <- RunUMAP(combined_seurat_subcluster, nn.name = "clusterwnn", reduction.name = "clusterumapWNN")
  # print subclustering
  pdf(file = paste0(plot_dir_w_date, "/", lineage, "_subclustering.", annotation, ".pdf"), width = 22, height = 6, useDingbats = F)
  p1 = DimPlot(combined_seurat_subcluster, reduction = "clusterumapWNN", label = T) & NoLegend()
  p2 = DimPlot(combined_seurat_subcluster, reduction = "clusterumapWNN", label = T, group.by = "wsnn_res.4") & NoLegend()
  p3 = DimPlot(combined_seurat, reduction = "umapWNN", label = T, group.by = "wsnn_res.4") & NoLegend()
  print(p1 | p2 | p3)
  dev.off()
  # identify modes of expression and divide genes into high/low/no expression
  cl_n = max(as.numeric(combined_seurat_subcluster$seurat_clusters))-1
  combined_seurat_subcluster_avgExp = as.data.frame(AverageExpression(combined_seurat_subcluster, slot = "data", assays = "RNA", features = row.names(markers_table)))
  combined_seurat_subcluster_avgExp_log = log10(combined_seurat_subcluster_avgExp + 1)
  combined_seurat_subcluster_avgExp_quant = combined_seurat_subcluster_avgExp_log
  pdf(file = paste0(plot_dir_w_date, "/", lineage, "_marker_binarisation.", annotation, ".pdf"), width = 8, height = 8, useDingbats = F)
  for (gene in c(1:dim(combined_seurat_subcluster_avgExp_log)[1])){
    loc <- locmodes(as.numeric(combined_seurat_subcluster_avgExp_log[gene,]), mod0 = 2, display = F)
    low_high_exp_loc = second_derivative_binarization(as.numeric(combined_seurat_subcluster_avgExp_log[gene,]), loc$cbw$bw, loc$locations, row.names(combined_seurat_subcluster_avgExp_log)[gene])
    combined_seurat_subcluster_avgExp_quant[gene,which(as.numeric(combined_seurat_subcluster_avgExp_log[gene,]) < low_high_exp_loc[1])] <- 0
    combined_seurat_subcluster_avgExp_quant[gene,which(as.numeric(combined_seurat_subcluster_avgExp_log[gene,]) >= low_high_exp_loc[1] & as.numeric(combined_seurat_subcluster_avgExp_log[gene,]) < low_high_exp_loc[2])] <- 1
    combined_seurat_subcluster_avgExp_quant[gene,which(as.numeric(combined_seurat_subcluster_avgExp_log[gene,]) >= low_high_exp_loc[2])] <- 2
  }
  dev.off()
  # binarize expression
  for (gene in c(1:dim(combined_seurat_subcluster_avgExp_quant)[1])){
    marker_gene_exp = as.numeric(combined_seurat_subcluster$seurat_clusters)-1
    for (cluster_n in c(0:cl_n)){
      marker_gene_exp[as.numeric(combined_seurat_subcluster$seurat_clusters)-1 == cluster_n] = combined_seurat_subcluster_avgExp_quant[gene, cluster_n+1]
    }
    combined_seurat_subcluster@meta.data[[row.names(combined_seurat_subcluster_avgExp_quant)[gene]]] = marker_gene_exp
  }
  # plot binarized expression
  pdf(file = paste0(plot_dir_w_date, "/", lineage, "_marker_binarisation.UMAP.", annotation, ".pdf"), width = 22, height = 6, useDingbats = F)
  for (gene in row.names(markers_table)){
    p1 = FeaturePlot(combined_seurat_subcluster, reduction = "clusterumapWNN", order = T, features = paste0("rna_", gene)) & scale_colour_gradientn(colours = c("grey90", "cyan", "deepskyblue", "darkorange", "darkviolet"))
    p2 = FeaturePlot(combined_seurat_subcluster, reduction = "clusterumapWNN", order = T, features = gene) & scale_colour_gradientn(colours = c("grey90", "cyan", "deepskyblue", "darkorange", "darkviolet"))
    p3 = DimPlot(combined_seurat_subcluster, reduction = "clusterumapWNN", label = T) & NoLegend()
    print(p1 | p2 | p3)
  }
  dev.off()
  # score subclusters based on markers expression
  scoring_cluster = matrix(NA, nrow = dim(markers_table)[2], ncol = dim(combined_seurat_subcluster_avgExp_quant)[2], dimnames = list(colnames(markers_table), colnames(combined_seurat_subcluster_avgExp_quant)))
  for (cluster_name in c(0:cl_n)){
    for (cell_type in c(1:dim(markers_table)[2])){
      scoring_cluster[cell_type, cluster_name+1] = scoring_markers(markers_table[,cell_type],combined_seurat_subcluster_avgExp_quant[, cluster_name+1])
    }
  }
  # output scoring table
  write.table(scoring_cluster, file = paste0(output_dir, "/", lineage, "_clusters_scoring.", annotation, ".txt"), quote = F, row.names = T, col.names = T)
  # assign subclusters to cell types
  n_markers = dim(markers_table)[1]
  marker_assignment = rep(NA, cluster_n + 1)
  for (cluster_n in c(1:dim(scoring_cluster)[2])){
    candidate_cell = which(scoring_cluster[,cluster_n] == max(scoring_cluster[,cluster_n]))
    if (max(scoring_cluster[,cluster_n]) >= n_markers*0.7 & length(candidate_cell) == 1){
      marker_assignment[cluster_n] = names(candidate_cell)
    }
    else if (scoring_cluster[,cluster_n][order(scoring_cluster[,cluster_n])][dim(scoring_cluster)[1]] > 0 &
             scoring_cluster[,cluster_n][order(scoring_cluster[,cluster_n])][dim(scoring_cluster)[1]] - scoring_cluster[,cluster_n][order(scoring_cluster[,cluster_n])][dim(scoring_cluster)[1]-1] > n_markers*0.5){
      marker_assignment[cluster_n] = names(candidate_cell)
    }
  }
  new_metadata = as.numeric(combined_seurat_subcluster$seurat_clusters)
  for (cluster_n in c(1:length(marker_assignment))){
    if (!is.na(marker_assignment[cluster_n])){
      new_metadata[as.numeric(combined_seurat_subcluster$seurat_clusters) == cluster_n] = marker_assignment[cluster_n]
    }
    else{
      new_metadata[as.numeric(combined_seurat_subcluster$seurat_clusters) == cluster_n] = NA
    }
  }
  combined_seurat_subcluster$cell_assignment = new_metadata
  p1 = DimPlot(combined_seurat_subcluster, reduction = "clusterumapWNN", order = T, label = T, group.by = "cell_assignment")
  p2 = DimPlot(combined_seurat_subcluster, reduction = "clusterumapWNN", order = T, label = T, group.by = "wsnn_res.5.5") & NoLegend()
  p3 = DimPlot(combined_seurat_subcluster, reduction = "clusterumapWNN", order = T, label = T, group.by = "wsnn_res.4") & NoLegend()
  
  # plot initial cell type assignment
  pdf(file = paste0(plot_dir_w_date, "/", lineage, "_full_annotation.before_rescue.UMAP.", annotation, ".pdf"), width = 14, height = 6, useDingbats = F)
  print(p1 | p2 | p3)
  dev.off()
  
  # rescue unassigned clusters: assign them to lineages based on lower resolution clustering
  new_marker_assignment = marker_assignment
  for (i in c(1:length(marker_assignment))){
    if (is.na(marker_assignment[i])){
      original_clusters = table(combined_seurat_subcluster$wsnn_res.4[combined_seurat_subcluster$wsnn_res.5.5 == i-1])
      original_clusters = names(original_clusters[original_clusters > sum(original_clusters) * 0.1])
      original_clusters_cell_types = table(combined_seurat_subcluster$cell_assignment[combined_seurat_subcluster$wsnn_res.4 %in% original_clusters])
      original_clusters_cell_types = names(original_clusters_cell_types[original_clusters_cell_types > sum(original_clusters_cell_types) * 0.1])
      original_clusters_cell_types = original_clusters_cell_types[!is.na(original_clusters_cell_types)]
      if (is.null(original_clusters_cell_types)){
        next
      }
      original_clusters_cell_types = unlist(strsplit(original_clusters_cell_types, split = "early_"))[unlist(strsplit(original_clusters_cell_types, split = "early_")) != ""]
      original_clusters_cell_types = unique(str_replace_all(original_clusters_cell_types, "[a-z]", "x"))
      if (length(original_clusters_cell_types) == 1){
        new_marker_assignment[i] = original_clusters_cell_types
      }
    }
  }
  #new_marker_assignment = cell_id_rescue(marker_assignment, markers_table[,cell_type])
  new_metadata = as.numeric(combined_seurat_subcluster$seurat_clusters)
  for (cluster_n in c(1:length(new_marker_assignment))){
    if (!is.na(new_marker_assignment[cluster_n])){
      new_metadata[as.numeric(combined_seurat_subcluster$seurat_clusters) == cluster_n] = new_marker_assignment[cluster_n]
    }
    else{
      new_metadata[as.numeric(combined_seurat_subcluster$seurat_clusters) == cluster_n] = NA
    }
  }
  combined_seurat_subcluster$cell_assignment = new_metadata
  p1 = DimPlot(combined_seurat_subcluster, reduction = "clusterumapWNN", order = T, label = T, group.by = "cell_assignment")
  p2 = DimPlot(combined_seurat_subcluster, reduction = "clusterumapWNN", order = T, label = T, group.by = "seurat_clusters") & NoLegend()
  p3 = DimPlot(combined_seurat_subcluster, reduction = "clusterumapWNN", order = T, label = T, group.by = "wsnn_res.4") & NoLegend()
  
  # plot cell annotation in subclustered UMAP
  pdf(file = paste0(plot_dir_w_date, "/", lineage, "_full_annotation.UMAP.", annotation, ".pdf"), width = 14, height = 6, useDingbats = F)
  print(p1 | p2 | p3)
  dev.off()
  
  # output cell annotation per barcode
  output_table = data.frame("cell_type" = combined_seurat_subcluster$cell_assignment, row.names = colnames(combined_seurat_subcluster))
  write.table(output_table, file = paste0(output_dir, "/", lineage, ".cell_assignment.", annotation, ".txt"), quote = F, sep = "\t", row.names = T, col.names = T)
}
```
