## Supplementary material for "A lineage-resolved multimodal single-cell atlas reveals the genomic dynamics of early *C. elegans* development": Code and external data: macs2_cellranger_fragments_all.20231103.sh

```
#!/bin/bash
#
#SBATCH --job-name=macs_atac
#SBATCH --output=slurm_out/macs_atac.%N.%j.out
#SBATCH --error=slurm_err/macs_atac.%N.%j.err
#SBATCH --ntasks=1
#SBATCH -N 1
#SBATCH -n 1
#SBATCH --mem=32G
#SBATCH --mail-type=END,FAIL
#SBATCH --mail-user=

input_filename_list=$1
output_dir=$2

chr_sizes=species/elegans/genome/elegans.chrom.sizes.txt
genome_index=species/elegans/genome/elegans.fa.fai
ce11_blacklist=data/external_data/ce11-blacklist.bed

input_filename=$(echo $input_filename_list | tr "," " ")

mkdir -p $output_dir

# split each fragment into two separate reads with --extsize 150 --shift -75
zcat $input_filename | grep -v "#" | awk 'BEGIN{OFS="\t";}{if ($3 - $2 < 150) print; else if ($2 - 75 < 0) print $1, 0, $2 + 75, $4, $5 "\n" $1, $3-75, $3 + 75, $4, $5; else print $1, $2 - 75, $2 + 75, $4, $5 "\n" $1, $3-75, $3 + 75, $4, $5}' | sort -k 1,1 -k2,2n > $output_dir/resized_fragments.all.bed

# call peaks using MACS
macs2 callpeak --call-summits --bdg --SPMR --extsize 150 --shift 0 --gsize ce --keep-dup all --nomodel -n resized_fragments.all_samples --outdir $output_dir/macs -t $output_dir/resized_fragments.all.bed
sort -k 1,1 -k2,2n $output_dir/macs/resized_fragments.all_samples_treat_pileup.bdg > $output_dir/macs/resized_fragments.all_samples_treat_pileup.sorted.bdg
awk 'BEGIN{OFS="\t";}{print $1, 1, $2}' $chr_sizes | intersectBed -a $output_dir/macs/resized_fragments.all_samples_treat_pileup.sorted.bdg -b stdin -u -f 1 > $output_dir/macs/resized_fragments.all_samples_treat_pileup.sorted.filtered.bdg
bedGraphToBigWig $output_dir/macs/resized_fragments.all_samples_treat_pileup.sorted.filtered.bdg $chr_sizes $output_dir/macs/resized_fragments.all_samples_treat_pileup.sorted.bw
rm $output_dir/macs/resized_fragments.all_samples_treat_pileup.bdg
rm $output_dir/macs/resized_fragments.all_samples_treat_pileup.sorted.bdg

# re-define MACS2 peaks by setting their size to 200bp centered around the summit
slopBed -i $output_dir/macs/resized_fragments.all_samples_summits.bed -b 100 -g $chr_sizes > $output_dir/macs/resized_fragments.all_samples_peaks.summits_centered.bed
```
