## Supplementary material for "A lineage-resolved multimodal single-cell atlas reveals the genomic dynamics of early *C. elegans* development": Code and external data: macs2_clusters.from_signac.20240207.sh

```
#!/bin/bash
#
#SBATCH --job-name=sinto
#SBATCH --output=slurm_out/macs2.%N.%j.out
#SBATCH --error=slurm_err/macs2.%N.%j.err
#SBATCH --ntasks=1
#SBATCH -N 1
#SBATCH -n 8
#SBATCH --mem-per-cpu=4000
#SBATCH --mail-type=END,FAIL
#SBATCH --mail-user=

input_filename=$1
input_path=$2
input_cell_n=$3

output_macs=$input_path/macs
output_bam=$input_path/bam
output_bw=$input_path/bw
output_resized_bed=$input_path/resized_bed

chr_sizes=species/elegans/genome/elegans.chrom.sizes.txt
genome_index=species/elegans/genome/elegans.fa.fai
ce11_blacklist=data/external_data/ce11-blacklist.bed

output_macs_dir=$output_macs/$input_filename

mkdir -p $output_resized_bed
mkdir -p $output_macs_dir
mkdir -p $output_bam
mkdir -p $output_bw

# split each fragment into two separate reads with --extsize 150 --shift -75
awk 'BEGIN{OFS="\t";}{if ($3 - $2 < 150) print; else if ($2 - 75 < 0) print $1, 0, $2 + 75, $4, $5 "\n" $1, $3-75, $3 + 75, $4, $5; else print $1, $2 - 75, $2 + 75, $4, $5 "\n" $1, $3-75, $3 + 75, $4, $5}' $input_path/bed/$input_filename.bed | sort -k 1,1 -k2,2n > $output_resized_bed/$input_filename.resized.bed

# call peaks using MACS
macs2 callpeak --llocal 2000 --call-summits --bdg --SPMR --extsize 150 --shift 0 --gsize ce --keep-dup all --nomodel -n $input_filename --outdir $output_macs_dir -t $output_resized_bed/$input_filename.resized.bed
sort -k 1,1 -k2,2n $output_macs_dir"/"$input_filename"_treat_pileup.bdg" > $output_macs_dir"/"$input_filename"_treat_pileup.sorted.bdg"
bedGraphToBigWig $output_macs_dir"/"$input_filename"_treat_pileup.sorted.bdg" $chr_sizes $output_macs_dir"/"$input_filename"_treat_pileup.sorted.bw"
rm $output_macs_dir"/"$input_filename"_treat_pileup.bdg"

# re-define MACS2 peaks by setting their size to 100bp centered around the summit and remove blacklisted regions
### peaks will be re-sized later on to avoid losing close, partially overlapping peaks
slopBed -i $output_macs_dir"/"$input_filename"_summits.bed" -b 50 -g $chr_sizes | intersectBed -a stdin -b $ce11_blacklist -v > $output_macs_dir"/"$input_filename"_peaks.summits_centered.bed"

# generate bw tracks of absolute number of ATAC-seq fragments normalized by the number of cells in each cluster x mean fragment n. per cell using bamCoverage 
cell_n=$(awk -v var="$input_filename" '{if ($1 == var) print 1/$2}' $input_cell_n)

bedToBam -i $output_resized_bed/$input_filename.resized.bed -g $genome_index > $output_bam/$input_filename.bam
samtools index $output_bam/$input_filename.bam
bamCoverage -b $output_bam/$input_filename.bam -o $output_bw/$input_filename.cell_n_norm.bw --scaleFactor $cell_n -bs 1 -bl $ce11_blacklist -p 8 --effectiveGenomeSize 100286401 --normalizeUsing None
bamCoverage -b $output_bam/$input_filename.bam -o $output_bw/$input_filename.cpm.bw -bs 1 -bl $ce11_blacklist -p 8 --effectiveGenomeSize 100286401 --normalizeUsing CPM
```
