## Supplementary material for "A lineage-resolved multimodal single-cell atlas reveals the genomic dynamics of early *C. elegans* development": Code and external data: memechip.20240521.sh

```
#!/bin/bash
#
#SBATCH --job-name=streme
#SBATCH --output=slurm_out/streme.%N.%j.out
#SBATCH --error=slurm_err/streme.%N.%j.err
#SBATCH -n 8
#SBATCH -N 1
#SBATCH --mem=32G
#SBATCH --mail-type=END,FAIL
#SBATCH --mail-user=

input_bed=$1
output_fa=$2
output_dir=$3
min_w=$4
max_w=$5
n_motifs=$6
markov_bkg=$7

genome_fa=species/elegans/genome/elegans.fa
motif_names=motif_names_cisbp2_elegans.txt

fastaFromBed -fi $genome_fa -bed $input_bed  -fo $output_fa

if [ -z "${markov_bkg}" ]
then
    meme-chip -ccut 0 --oc $output_dir".w_"$min_w"_"$max_w -minw $min_w -maxw $max_w -seed 32 -filter-thresh 0.05 -meme-mod zoops -meme-nmotifs $n_motifs -meme-minsites 10 -meme-p 8 -streme-pvt 0.05 -db motif_databases/CIS-BP_2.00/Caenorhabditis_elegans.meme $output_fa
else
    meme-chip -ccut 0 --oc $output_dir".w_"$min_w"_"$max_w -minw $min_w -maxw $max_w -seed 32 -filter-thresh 0.05 -meme-mod zoops -meme-nmotifs $n_motifs -meme-minsites 10 -meme-p 8 -streme-pvt 0.05 -db motif_databases/CIS-BP_2.00/Caenorhabditis_elegans.meme -bfile $markov_bkg $output_fa
fi

rm $output_fa
```
