## Supplementary material for "A lineage-resolved multimodal single-cell atlas reveals the genomic dynamics of early *C. elegans* development": Code and external data: merge_dedup_bam.20231031.sh

```
#!/bin/bash
#
#SBATCH --job-name=bam_dedup_merge
#SBATCH --output=slurm_out/dedup_merge_bam.%N.%j.out
#SBATCH --error=slurm_err/dedup_merge_bam.%N.%j.err
#SBATCH --ntasks=1
#SBATCH -N 1
#SBATCH -n 8
#SBATCH --mem-per-cpu=4000
#SBATCH --mail-type=END,FAIL
#SBATCH --mail-user=

output_dedup=$1
input_dedup_list=$2
dup_input=$3

input_dedup=$(echo $input_dedup_list | tr "," " ")

samtools merge -h $dup_input -@ 8 $output_dedup $input_dedup
```
