## Supplementary material for "A lineage-resolved multimodal single-cell atlas reveals the genomic dynamics of early *C. elegans* development": Code and external data: merge_multiple_seurat_objects.20240430.R

```
# combine two seurat objects, re-cluster cells based on RNA-seq data, create a shared seurat object, then integrate the ATAC -seq data based on the merged MACS peaks called on all cells from each experiment
library(ggplot2)
library(dplyr)
library(Seurat)
library(patchwork)
library(DropletUtils)
library(pheatmap)
library(RColorBrewer)
library(scDblFinder)
library(scater)
library(glmGamPoi)
library(Signac)
library(rtracklayer)

set.seed(42)
args = commandArgs(trailingOnly=TRUE)

exp_1_id = args[1]
exp_2_id = args[2]
exp_3_id = args[3]
exp_4_id = args[4]
exp_5_id = args[5]
exp_6_id = args[6]
exp_7_id = args[7]
exp_1_id_seurat_rds = args[8]
exp_2_id_seurat_rds = args[9]
exp_3_id_seurat_rds = args[10]
exp_4_id_seurat_rds = args[11]
exp_5_id_seurat_rds = args[12]
exp_6_id_seurat_rds = args[13]
exp_7_id_seurat_rds = args[14]

merged_macs = args[15]
fragments_out_dir = args[16]
annotation = args[17]

cR_output = "data/sc/cellranger_arc/"

cR_output_exp1 = paste0(cR_output, exp_1_id, "/outs/raw_feature_bc_matrix.h5")
cR_output_exp2 = paste0(cR_output, exp_2_id, "/outs/raw_feature_bc_matrix.h5")
cR_output_exp3 = paste0(cR_output, exp_3_id, "/outs/raw_feature_bc_matrix.h5")
cR_output_exp4 = paste0(cR_output, exp_4_id, "/outs/raw_feature_bc_matrix.h5")
cR_output_exp5 = paste0(cR_output, exp_5_id, "/outs/raw_feature_bc_matrix.h5")
cR_output_exp6 = paste0(cR_output, exp_6_id, "/outs/raw_feature_bc_matrix.h5")
cR_output_exp7 = paste0(cR_output, exp_7_id, "/outs/raw_feature_bc_matrix.h5")

fragpath_exp_1 = paste0(cR_output, exp_1_id, "/outs/atac_fragments_barcode_corrected.tsv.gz")
fragpath_exp_2 = paste0(cR_output, exp_2_id, "/outs/atac_fragments_barcode_corrected.tsv.gz")
fragpath_exp_3 = paste0(cR_output, exp_3_id, "/outs/atac_fragments_barcode_corrected.tsv.gz")
fragpath_exp_4 = paste0(cR_output, exp_4_id, "/outs/atac_fragments_barcode_corrected.tsv.gz")
fragpath_exp_5 = paste0(cR_output, exp_5_id, "/outs/atac_fragments_barcode_corrected.tsv.gz")
fragpath_exp_6 = paste0(cR_output, exp_6_id, "/outs/atac_fragments_barcode_corrected.tsv.gz")
fragpath_exp_7 = paste0(cR_output, exp_7_id, "/outs/atac_fragments_barcode_corrected.tsv.gz")

current_date= paste(unlist(strsplit(as.character(Sys.Date()), "-")), collapse="")

dir.create(paste0("plots/", current_date))

# exp_1_id_seurat_rds="seurat_objects/exp031_postSoupX_before_reseq.ATAC_MACS.20230530.rds"
# exp_2_id_seurat_rds="seurat_objects/exp032_postSoupX_before_reseq.ATAC_MACS.20230530.rds"
# exp_3_id_seurat_rds="seurat_objects/exp042_postSoupX.ATAC_MACS.20230502.rds"
# exp_4_id_seurat_rds="seurat_objects/exp043_postSoupX.ATAC_MACS.20230502.rds"
# exp_1_id="exp031"
# exp_2_id="exp032"
# exp_3_id="exp042"
# exp_4_id="exp043"

exp_1_id_seurat <- readRDS(exp_1_id_seurat_rds)
exp_2_id_seurat <- readRDS(exp_2_id_seurat_rds)
exp_3_id_seurat <- readRDS(exp_3_id_seurat_rds)
exp_4_id_seurat <- readRDS(exp_4_id_seurat_rds)
exp_5_id_seurat <- readRDS(exp_5_id_seurat_rds)
exp_6_id_seurat <- readRDS(exp_6_id_seurat_rds)
exp_7_id_seurat <- readRDS(exp_7_id_seurat_rds)

#ee.list <- list(exp_1_id_rnaseq_data_seurat_soupx, exp_2_id_rnaseq_data_seurat_soupx, exp_3_id_rnaseq_data_seurat_soupx, exp_4_id_rnaseq_data_seurat_soupx)
ee.list <- list(exp_1_id_seurat@assays[["RNA"]], exp_2_id_seurat@assays[["RNA"]], exp_3_id_seurat@assays[["RNA"]], exp_4_id_seurat@assays[["RNA"]], exp_5_id_seurat@assays[["RNA"]], exp_6_id_seurat@assays[["RNA"]], exp_7_id_seurat@assays[["RNA"]])

exp_1_id_seurat@meta.data$exp <- exp_1_id
exp_2_id_seurat@meta.data$exp <- exp_2_id
exp_3_id_seurat@meta.data$exp <- exp_3_id
exp_4_id_seurat@meta.data$exp <- exp_4_id
exp_5_id_seurat@meta.data$exp <- exp_5_id
exp_6_id_seurat@meta.data$exp <- exp_6_id
exp_7_id_seurat@meta.data$exp <- exp_7_id

#gcdata <- merge(exp_1_id_seurat@assays[["RNA"]], y=c(exp_2_id_seurat@assays[["RNA"]], exp_3_id_seurat@assays[["RNA"]], exp_4_id_seurat@assays[["RNA"]]),
#                add.cell.ids=c(exp_1_id, exp_2_id, exp_3_id, exp_4_id), merge.data = T)
gcdata <- merge(exp_1_id_seurat, y=c(exp_2_id_seurat, exp_3_id_seurat, exp_4_id_seurat, exp_5_id_seurat, exp_6_id_seurat, exp_7_id_seurat),
                add.cell.ids=c(exp_1_id, exp_2_id, exp_3_id, exp_4_id, exp_5_id, exp_6_id, exp_7_id), merge.data = T)

VariableFeatures(gcdata[["SCT"]]) <- rownames(gcdata[["SCT"]]@scale.data)
DefaultAssay(gcdata) = "SCT"
#all.genes <- rownames(gcdata)

gcdata <- RunPCA(gcdata, seed.use = 42, features = VariableFeatures(object = gcdata), npcs = 100)

gcdata <- FindNeighbors(gcdata, dims = 1:50)
gcdata <- FindClusters(gcdata, resolution = 3, random.seed = 0)

gcdata <- RunUMAP(gcdata, seed.use = 42, dims = 1:50, return.model=TRUE)

pdf(file=paste0("plots/", current_date, "/", exp_1_id, "_", exp_2_id, "_", exp_3_id, "_", exp_4_id, "_", exp_5_id, "_", exp_6_id,  "_", exp_7_id, "_SCT.", annotation, ".umap_split.50PCA.pdf"), width=18, height = 8)
ElbowPlot(gcdata, ndims = 100)
DimPlot(gcdata, reduction = "umap", group.by = "orig.ident")
DimPlot(gcdata, reduction = "umap", split.by = "orig.ident")
DimPlot(gcdata, reduction = "umap", label = TRUE )
dev.off()

p1 = DimPlot(gcdata, reduction = "umap", group.by = "orig.ident")
p2 = DimPlot(gcdata, reduction = "umap", split.by = "orig.ident")
p3 = DimPlot(gcdata, reduction = "umap", label = TRUE )
(p1 | p3) / p2

pdf(file=paste0("plots/", current_date, "/", exp_1_id, "_", exp_2_id, "_", exp_3_id, "_", exp_4_id, "_", exp_5_id, "_", exp_6_id, "_", exp_7_id, "_SCT.", annotation, ".key_lineage_markers.50PCA.pdf"), width=8, height = 8)
FeaturePlot(gcdata, order = T, features = c("med-1")) & scale_colour_gradientn(colours = c("grey90", "cyan", "deepskyblue", "forestgreen", "darkorange", "darkred"))
FeaturePlot(gcdata, order = T, features = c("tbx-40")) & scale_colour_gradientn(colours = c("grey90", "cyan", "deepskyblue", "forestgreen", "darkorange", "darkred"))
FeaturePlot(gcdata, order = T, features = c("tbx-35")) & scale_colour_gradientn(colours = c("grey90", "cyan", "deepskyblue", "forestgreen", "darkorange", "darkred"))
FeaturePlot(gcdata, order = T, features = c("tbx-33")) & scale_colour_gradientn(colours = c("grey90", "cyan", "deepskyblue", "forestgreen", "darkorange", "darkred"))
FeaturePlot(gcdata, order = T, features = c("end-3")) & scale_colour_gradientn(colours = c("grey90", "cyan", "deepskyblue", "forestgreen", "darkorange", "darkred"))
FeaturePlot(gcdata, order = T, features = c("hnd-1")) & scale_colour_gradientn(colours = c("grey90", "cyan", "deepskyblue", "forestgreen", "darkorange", "darkred"))
FeaturePlot(gcdata, order = T, features = c("pes-10")) & scale_colour_gradientn(colours = c("grey90", "cyan", "deepskyblue", "forestgreen", "darkorange", "darkred"))
FeaturePlot(gcdata, order = T, features = c("vet-2")) & scale_colour_gradientn(colours = c("grey90", "cyan", "deepskyblue", "forestgreen", "darkorange", "darkred"))
FeaturePlot(gcdata, order = T, features = c("ceh-51")) & scale_colour_gradientn(colours = c("grey90", "cyan", "deepskyblue", "forestgreen", "darkorange", "darkred"))
FeaturePlot(gcdata, order = T, features = c("oma-1")) & scale_colour_gradientn(colours = c("grey90", "cyan", "deepskyblue", "forestgreen", "darkorange", "darkred"))
FeaturePlot(gcdata, order = T, features = c("oma-2")) & scale_colour_gradientn(colours = c("grey90", "cyan", "deepskyblue", "forestgreen", "darkorange", "darkred"))
FeaturePlot(gcdata, order = T, features = c("lsl-1")) & scale_colour_gradientn(colours = c("grey90", "cyan", "deepskyblue", "forestgreen", "darkorange", "darkred"))
FeaturePlot(gcdata, order = T, features = c("skr-7")) & scale_colour_gradientn(colours = c("grey90", "cyan", "deepskyblue", "forestgreen", "darkorange", "darkred"))
FeaturePlot(gcdata, order = T, features = c("ccch-2")) & scale_colour_gradientn(colours = c("grey90", "cyan", "deepskyblue", "forestgreen", "darkorange", "darkred"))
FeaturePlot(gcdata, order = T, features = c("ref-1")) & scale_colour_gradientn(colours = c("grey90", "cyan", "deepskyblue", "forestgreen", "darkorange", "darkred"))
dev.off()

pdf(file=paste0("plots/", current_date, "/", exp_1_id, "_", exp_2_id, "_", exp_3_id, "_", exp_4_id, "_", exp_5_id, "_", exp_6_id, "_", exp_7_id, "_SCT.", annotation, ".umap.50PCA.pdf"), width=8, height = 8)
DimPlot(gcdata, reduction = "umap", label = TRUE)
dev.off()

# add ATAC data
counts_exp_1 <- Read10X_h5(cR_output_exp1)
colnames(counts_exp_1$Peaks) <- sapply(strsplit(colnames(counts_exp_1$Peaks), split='-', fixed=TRUE), `[`, 1)
rna_barcodes_exp_1 <- sapply(strsplit(colnames(gcdata)[grep(exp_1_id, colnames(gcdata))], "_"), "[[", 2)
counts_new_exp_1 <- counts_exp_1$Peaks[,colnames(counts_exp_1$Peaks) %in% rna_barcodes_exp_1]

counts_exp_2 <- Read10X_h5(cR_output_exp2)
colnames(counts_exp_2$Peaks) <- sapply(strsplit(colnames(counts_exp_2$Peaks), split='-', fixed=TRUE), `[`, 1)
rna_barcodes_exp_2 <- sapply(strsplit(colnames(gcdata)[grep(exp_2_id, colnames(gcdata))], "_"), "[[", 2)
counts_new_exp_2 <- counts_exp_2$Peaks[,colnames(counts_exp_2$Peaks) %in% rna_barcodes_exp_2]

counts_exp_3 <- Read10X_h5(cR_output_exp3)
colnames(counts_exp_3$Peaks) <- sapply(strsplit(colnames(counts_exp_3$Peaks), split='-', fixed=TRUE), `[`, 1)
rna_barcodes_exp_3 <- sapply(strsplit(colnames(gcdata)[grep(exp_3_id, colnames(gcdata))], "_"), "[[", 2)
counts_new_exp_3 <- counts_exp_3$Peaks[,colnames(counts_exp_3$Peaks) %in% rna_barcodes_exp_3]

counts_exp_4 <- Read10X_h5(cR_output_exp4)
colnames(counts_exp_4$Peaks) <- sapply(strsplit(colnames(counts_exp_4$Peaks), split='-', fixed=TRUE), `[`, 1)
rna_barcodes_exp_4 <- sapply(strsplit(colnames(gcdata)[grep(exp_4_id, colnames(gcdata))], "_"), "[[", 2)
counts_new_exp_4 <- counts_exp_4$Peaks[,colnames(counts_exp_4$Peaks) %in% rna_barcodes_exp_4]

counts_exp_5 <- Read10X_h5(cR_output_exp5)
colnames(counts_exp_5$Peaks) <- sapply(strsplit(colnames(counts_exp_5$Peaks), split='-', fixed=TRUE), `[`, 1)
rna_barcodes_exp_5 <- sapply(strsplit(colnames(gcdata)[grep(exp_5_id, colnames(gcdata))], "_"), "[[", 2)
counts_new_exp_5 <- counts_exp_5$Peaks[,colnames(counts_exp_5$Peaks) %in% rna_barcodes_exp_5]

counts_exp_6 <- Read10X_h5(cR_output_exp6)
colnames(counts_exp_6$Peaks) <- sapply(strsplit(colnames(counts_exp_6$Peaks), split='-', fixed=TRUE), `[`, 1)
rna_barcodes_exp_6 <- sapply(strsplit(colnames(gcdata)[grep(exp_6_id, colnames(gcdata))], "_"), "[[", 2)
counts_new_exp_6 <- counts_exp_6$Peaks[,colnames(counts_exp_6$Peaks) %in% rna_barcodes_exp_6]

counts_exp_7 <- Read10X_h5(cR_output_exp7)
colnames(counts_exp_7$Peaks) <- sapply(strsplit(colnames(counts_exp_7$Peaks), split='-', fixed=TRUE), `[`, 1)
rna_barcodes_exp_7 <- sapply(strsplit(colnames(gcdata)[grep(exp_7_id, colnames(gcdata))], "_"), "[[", 2)
counts_new_exp_7 <- counts_exp_7$Peaks[,colnames(counts_exp_7$Peaks) %in% rna_barcodes_exp_7]

#fragpath_exp_1 <- "temp_analysis/20230219/exp031_atac_fragment_barcode_corrected.tsv.gz"
#fragpath_exp_2 <- "temp_analysis/20230219/exp032_atac_fragment_barcode_corrected.tsv.gz"

merged_peaks = import.bed(merged_macs)

fragments_exp_1 <- CreateFragmentObject(fragpath_exp_1, cells = colnames(gcdata)[grep(exp_1_id, colnames(gcdata))])
fragments_exp_2 <- CreateFragmentObject(fragpath_exp_2, cells = colnames(gcdata)[grep(exp_2_id, colnames(gcdata))])
fragments_exp_3 <- CreateFragmentObject(fragpath_exp_3, cells = colnames(gcdata)[grep(exp_3_id, colnames(gcdata))])
fragments_exp_4 <- CreateFragmentObject(fragpath_exp_4, cells = colnames(gcdata)[grep(exp_4_id, colnames(gcdata))])
fragments_exp_5 <- CreateFragmentObject(fragpath_exp_5, cells = colnames(gcdata)[grep(exp_5_id, colnames(gcdata))])
fragments_exp_6 <- CreateFragmentObject(fragpath_exp_6, cells = colnames(gcdata)[grep(exp_6_id, colnames(gcdata))])
fragments_exp_7 <- CreateFragmentObject(fragpath_exp_7, cells = colnames(gcdata)[grep(exp_7_id, colnames(gcdata))])

merged_peaks_matrix_filtered <- FeatureMatrix(list(fragments_exp_1,fragments_exp_2,fragments_exp_3,fragments_exp_4,fragments_exp_5,fragments_exp_6,fragments_exp_7), sep = c("-", "-"), features=merged_peaks, cells = colnames(gcdata))
merged_peaks_matrix_filtered1 <- merged_peaks_matrix_filtered[,colnames(merged_peaks_matrix_filtered) %in% colnames(gcdata)]

gcdata[["ATAC"]] <- CreateChromatinAssay(
  counts = merged_peaks_matrix_filtered1,
  sep = c(":", "-"),
  fragments = list(fragments_exp_1, fragments_exp_2, fragments_exp_3, fragments_exp_4, fragments_exp_5, fragments_exp_6, fragments_exp_7))


pdf(file=paste0("plots/", current_date, "/combined_", exp_1_id, "_", exp_2_id, "_", exp_3_id, "_", exp_4_id, "_", exp_5_id, "_", exp_6_id, "_", exp_7_id, "_SCT.", annotation, ".all_cells.MACS_peaks.nfeature_atac_rna.pdf"), width=24, height = 8)
p1 = DimPlot(gcdata, reduction = "umap", group.by = "seurat_clusters", label = TRUE, label.size = 4)
p2 = FeaturePlot(gcdata, features = "nFeature_ATAC", min.cutoff = 0, max.cutoff = 2000, order=T) & scale_colour_gradientn(colours = c("grey90", "cyan", "deepskyblue", "forestgreen", "darkorange", "darkred"))
p3 = FeaturePlot(gcdata, features = "nFeature_RNA", min.cutoff = 0, max.cutoff = 1000, order=T) & scale_colour_gradientn(colours = c("grey90", "cyan", "deepskyblue", "forestgreen", "darkorange", "darkred"))
p1 + p2 + p3
dev.off()

pdf(file=paste0("plots/", current_date, "/combined_", exp_1_id, "_", exp_2_id, "_", exp_3_id, "_", exp_4_id, "_", exp_5_id, "_", exp_6_id, "_", exp_7_id, "_SCT.", annotation, ".all_cells.MACS_peaks.ncounts_atac_rna.pdf"), width=24, height = 8)
p1 = DimPlot(gcdata, reduction = "umap", group.by = "seurat_clusters", label = TRUE, label.size = 4)
p2 = FeaturePlot(gcdata, features = "nCount_ATAC", min.cutoff = 0, max.cutoff = 3000, order=T) & scale_colour_gradientn(colours = c("grey90", "cyan", "deepskyblue", "forestgreen", "darkorange", "darkred"))
p3 = FeaturePlot(gcdata, features = "nCount_RNA", min.cutoff = 0, max.cutoff = 2000, order=T) & scale_colour_gradientn(colours = c("grey90", "cyan", "deepskyblue", "forestgreen", "darkorange", "darkred"))
p1 + p2 + p3
dev.off()

saveRDS(gcdata, file = paste0("seurat_objects/combined_", exp_1_id, "_", exp_2_id, "_", exp_3_id, "_", exp_4_id, "_", exp_5_id, "_", exp_6_id, "_", exp_7_id, "_SCT.", annotation, ".all_cells.MACS_peaks.rds"))
write.table(gcdata[[]], file = paste0("seurat_objects/combined_", exp_1_id, "_", exp_2_id, "_", exp_3_id, "_", exp_4_id, "_", exp_5_id, "_", exp_6_id, "_", exp_7_id, "_SCT.", annotation, ".all_cells.MACS_peaks.table.txt"), quote = F, sep = "\t", row.names = T, col.names = T)

DefaultAssay(gcdata) <- "ATAC"

# output distinct fragment files per cluster to create bigWig tracks
dir.create(fragments_out_dir, recursive = T)

SplitFragments(gcdata, group.by = "seurat_clusters",
               outdir = fragments_out_dir, append = T)

# output number of cells per cluster
n_clusters = unique(gcdata$seurat_clusters)
n_clusters = n_clusters[order(n_clusters, as.numeric(n_clusters))]
cells_cluster=data.frame(row.names=n_clusters, "n_cells" = rep(NA, length(n_clusters)))

for (n_c in c(1:length(n_clusters))){
  cells_cluster[n_c,1]=dim(gcdata[,gcdata$seurat_clusters == n_clusters[n_c]])[2]
}
write.table(cells_cluster, file = paste0("seurat_objects/combined_", exp_1_id, "_", exp_2_id, "_", exp_3_id, "_", exp_4_id, "_", exp_5_id, "_", exp_6_id, "_", exp_7_id, "_SCT.", annotation, ".all_cells.MACS_peaks.cells_per_cluster.txt"), quote = F, sep = "\t", row.names = T, col.names = T)
```
