## Supplementary material for "A lineage-resolved multimodal single-cell atlas reveals the genomic dynamics of early *C. elegans* development": Code and external data: merge_multiple_seurat_objects.cluster_peaks_step_2.20240430.R

```
# upload the cluster-specific peaks in the new merged object
library(ggplot2)
library(dplyr)
library(Seurat)
library(patchwork)
library(RColorBrewer)
library(glmGamPoi)
library(Signac)
library(rtracklayer)

set.seed(42)
args = commandArgs(trailingOnly=TRUE)

combined_seurat_rds = args[1]
merged_peaks_bed = args[2]
exp_1_id = args[3]
exp_2_id = args[4]
exp_3_id = args[5]
exp_4_id = args[6]
exp_5_id = args[7]
exp_6_id = args[8]
exp_7_id = args[9]
new_ATAC_assay = args[10]
annotation = args[11]

current_date= paste(unlist(strsplit(as.character(Sys.Date()), "-")), collapse="")
plot_dir_w_date = paste0("plots/", current_date)

dir.create(plot_dir_w_date, recursive = T)

combined_seurat <- readRDS(combined_seurat_rds)
merged_peaks = import.bed(merged_peaks_bed)

merged_peaks_matrix_filtered <- FeatureMatrix(combined_seurat[["ATAC"]]@fragments, sep = c("-", "-"), features=merged_peaks, cells = colnames(combined_seurat))

combined_seurat[[new_ATAC_assay]] <- CreateChromatinAssay(
  counts = merged_peaks_matrix_filtered,
  sep = c(":", "-"),
  fragments = combined_seurat[["ATAC"]]@fragments)

# cluster barcodes based on ATAC data 
DefaultAssay(combined_seurat) <- new_ATAC_assay
combined_seurat <- RunTFIDF(combined_seurat)
combined_seurat <- FindTopFeatures(combined_seurat, min.cutoff = 'q0')
combined_seurat <- RunSVD(combined_seurat)
combined_seurat <- RunUMAP(combined_seurat, reduction = 'lsi', dims = 2:50, reduction.name = "umapATAC")
combined_seurat <- FindNeighbors(combined_seurat, reduction = 'lsi', dims = 2:50)
combined_seurat <- FindClusters(combined_seurat, verbose = T, algorithm = 3, resolution = 3)

# cluster barcodes based on both modalities
combined_seurat <- FindMultiModalNeighbors(combined_seurat, reduction.list = list("pca", "lsi"), dims.list = list(1:50, 2:50))
combined_seurat <- RunUMAP(combined_seurat, nn.name = "weighted.nn", reduction.name = "umapWNN")
combined_seurat <- FindClusters(combined_seurat, graph.name = "wsnn", algorithm = 3, verbose = T, resolution = 4)

p1 <- DimPlot(combined_seurat, reduction = "umap", group.by = "wsnn_res.4", label = TRUE, label.size = 4)
p2 <- DimPlot(combined_seurat, reduction = "umapATAC", group.by = "wsnn_res.4", label = TRUE, label.size = 4)
p3 <- DimPlot(combined_seurat, reduction = "umapWNN", group.by = "wsnn_res.4", label = TRUE, label.size = 4)

pdf(file=paste0(plot_dir_w_date, "/",exp_1_id,"_", exp_2_id, "_",exp_3_id,"_", exp_4_id,"_", exp_5_id,"_", exp_6_id,"_", exp_7_id, "_SCT.", annotation, ".umap_WNN.pdf"), width=24, height = 8)
p1 + p2 + p3 & NoLegend() & theme(plot.title = element_text(hjust = 0.5))
dev.off()

pdf(file=paste0(plot_dir_w_date, "/",exp_1_id,"_", exp_2_id, "_",exp_3_id,"_", exp_4_id,"_", exp_5_id,"_", exp_6_id,"_", exp_7_id, "_SCT.", annotation, ".umap_split.WNN.pdf"), width=24, height = 8)
p1 = DimPlot(combined_seurat, reduction = "umapWNN", label = TRUE, label.size = 4) 
p2 = DimPlot(combined_seurat, reduction = "umapWNN", group.by = "exp")
p1 & NoLegend() & theme(plot.title = element_text(hjust = 0.5)) | p2  & theme(plot.title = element_text(hjust = 0.5))
DimPlot(combined_seurat, reduction = "umapWNN", split.by = "exp") & NoLegend() & theme(plot.title = element_text(hjust = 0.5))
dev.off()

DefaultAssay(combined_seurat) = "SCT"

  pdf(file=paste0("plots/", current_date, "/", exp_1_id, "_", exp_2_id, "_", exp_3_id, "_", exp_4_id,"_", exp_5_id,"_", exp_6_id,"_", exp_7_id, "_SCT.", annotation, ".key_lineage_markers.WNN.pdf"), width=8, height = 8)
  FeaturePlot(combined_seurat, order = T, features = c("med-1"), reduction = "umapWNN") & scale_colour_gradientn(colours = c("grey90", "cyan", "deepskyblue", "forestgreen", "darkorange", "darkred"))
  FeaturePlot(combined_seurat, order = T, features = c("tbx-40"), reduction = "umapWNN") & scale_colour_gradientn(colours = c("grey90", "cyan", "deepskyblue", "forestgreen", "darkorange", "darkred"))
  FeaturePlot(combined_seurat, order = T, features = c("tbx-35"), reduction = "umapWNN") & scale_colour_gradientn(colours = c("grey90", "cyan", "deepskyblue", "forestgreen", "darkorange", "darkred"))
  FeaturePlot(combined_seurat, order = T, features = c("tbx-33"), reduction = "umapWNN") & scale_colour_gradientn(colours = c("grey90", "cyan", "deepskyblue", "forestgreen", "darkorange", "darkred"))
  FeaturePlot(combined_seurat, order = T, features = c("end-3"), reduction = "umapWNN") & scale_colour_gradientn(colours = c("grey90", "cyan", "deepskyblue", "forestgreen", "darkorange", "darkred"))
  FeaturePlot(combined_seurat, order = T, features = c("hnd-1"), reduction = "umapWNN") & scale_colour_gradientn(colours = c("grey90", "cyan", "deepskyblue", "forestgreen", "darkorange", "darkred"))
  FeaturePlot(combined_seurat, order = T, features = c("pes-10"), reduction = "umapWNN") & scale_colour_gradientn(colours = c("grey90", "cyan", "deepskyblue", "forestgreen", "darkorange", "darkred"))
  FeaturePlot(combined_seurat, order = T, features = c("vet-2"), reduction = "umapWNN") & scale_colour_gradientn(colours = c("grey90", "cyan", "deepskyblue", "forestgreen", "darkorange", "darkred"))
  FeaturePlot(combined_seurat, order = T, features = c("ceh-51"), reduction = "umapWNN") & scale_colour_gradientn(colours = c("grey90", "cyan", "deepskyblue", "forestgreen", "darkorange", "darkred"))
  FeaturePlot(combined_seurat, order = T, features = c("oma-1"), reduction = "umapWNN") & scale_colour_gradientn(colours = c("grey90", "cyan", "deepskyblue", "forestgreen", "darkorange", "darkred"))
  FeaturePlot(combined_seurat, order = T, features = c("oma-2"), reduction = "umapWNN") & scale_colour_gradientn(colours = c("grey90", "cyan", "deepskyblue", "forestgreen", "darkorange", "darkred"))
  FeaturePlot(combined_seurat, order = T, features = c("lsl-1"), reduction = "umapWNN") & scale_colour_gradientn(colours = c("grey90", "cyan", "deepskyblue", "forestgreen", "darkorange", "darkred"))
  FeaturePlot(combined_seurat, order = T, features = c("skr-7"), reduction = "umapWNN") & scale_colour_gradientn(colours = c("grey90", "cyan", "deepskyblue", "forestgreen", "darkorange", "darkred"))
  FeaturePlot(combined_seurat, order = T, features = c("ccch-2"), reduction = "umapWNN") & scale_colour_gradientn(colours = c("grey90", "cyan", "deepskyblue", "forestgreen", "darkorange", "darkred"))
  FeaturePlot(combined_seurat, order = T, features = c("ref-1"), reduction = "umapWNN") & scale_colour_gradientn(colours = c("grey90", "cyan", "deepskyblue", "forestgreen", "darkorange", "darkred"))
  FeaturePlot(combined_seurat, order = T, features = c("cpg-3"), reduction = "umapWNN") & scale_colour_gradientn(colours = c("grey90", "cyan", "deepskyblue", "forestgreen", "darkorange", "darkred"))
  dev.off()

# save RDS
saveRDS(combined_seurat, file = paste0("seurat_objects/combined_",exp_1_id,"_", exp_2_id, "_",exp_3_id,"_", exp_4_id,"_", exp_5_id,"_", exp_6_id,"_", exp_7_id, "_SCT.", annotation, ".WNN.rds"))

# output distinct fragment files per cluster to create bigWig tracks
#DefaultAssay(combined_seurat) <- new_ATAC_assay

#fragments_out_dir = paste0("cluster_specific_peaks/round_2/", annotation, "/bed")
#dir.create(fragments_out_dir, recursive = T)

#SplitFragments(combined_seurat, group.by = "wsnn_res.4",
#               outdir = fragments_out_dir, append = T)

#n_clusters = unique(combined_seurat$wsnn_res.4)
#n_clusters = n_clusters[order(n_clusters, as.numeric(n_clusters))]
#cells_cluster=data.frame(row.names=n_clusters, "n_cells" = rep(NA, length(n_clusters)))

#for (n_c in c(1:length(n_clusters))){
#  cells_cluster[n_c,1]=dim(combined_seurat[,combined_seurat$wsnn_res.4 == n_clusters[n_c]])[2]
#}
#write.table(cells_cluster, file = paste0("cluster_specific_peaks/round_2/", annotation, "/",exp_1_id,"_", exp_2_id, "_",exp_3_id,"_", exp_4_id, "_SCT.WNN.cells_per_cluster.txt"), quote = F, sep = "\t", row.names = T, col.names = T)
```
