## Supplementary material for "A lineage-resolved multimodal single-cell atlas reveals the genomic dynamics of early *C. elegans* development": Code and external data: merge_multiple_seurat_objects.cluster_peaks_step_2.20240430.sh

```
#!/bin/bash
#
#SBATCH --job-name=seurat_merge
#SBATCH --output=slurm_out/seurat_merge.%N.%j.out
#SBATCH --error=slurm_err/seurat_merge.%N.%j.err
#SBATCH --ntasks=1
#SBATCH -N 1
#SBATCH -n 1
#SBATCH --mem=64G
#SBATCH --mail-type=END,FAIL
#SBATCH --mail-user=

seurat_obj=$1
peak_set=$2
sample_1=$3
sample_2=$4
sample_3=$5
sample_4=$6
sample_5=$7
sample_6=$8
sample_7=$9
assay_name=${10}
annotation=${11}

Rscript scripts/merge_multiple_seurat_objects.cluster_peaks_step_2.20240430.R $seurat_obj $peak_set $sample_1 $sample_2 $sample_3 $sample_4 $sample_5 $sample_6 $sample_7 $assay_name $annotation
```
