## Supplementary material for "A lineage-resolved multimodal single-cell atlas reveals the genomic dynamics of early *C. elegans* development": Code and external data: merge_multiple_seurat_objects_launcher.20240430.sh

```
#!/bin/bash
#
#SBATCH --job-name=seurat_merge
#SBATCH --output=slurm_out/seurat_merge.%N.%j.out
#SBATCH --error=slurm_err/seurat_merge.%N.%j.err
#SBATCH --ntasks=1
#SBATCH -N 1
#SBATCH -n 1
#SBATCH --mem=64G
#SBATCH --mail-type=END,FAIL
#SBATCH --mail-user=

sample_1=$1
sample_2=$2
sample_3=$3
sample_4=$4
sample_5=$5
sample_6=$6
sample_7=$7
seurat_sample_1=$8
seurat_sample_2=$9
seurat_sample_3=${10}
seurat_sample_4=${11}
seurat_sample_5=${12}
seurat_sample_6=${13}
seurat_sample_7=${14}
peak_set=${15}
frags_out=${16}
annotation=${17}

Rscript scripts/merge_multiple_seurat_objects.20240430.R $sample_1 $sample_2 $sample_3 $sample_4 $sample_5 $sample_6 $sample_7 $seurat_sample_1 $seurat_sample_2 $seurat_sample_3 $seurat_sample_4 $seurat_sample_5 $seurat_sample_6 $seurat_sample_7 $peak_set $frags_out $annotation
```
