## Supplementary material for "A lineage-resolved multimodal single-cell atlas reveals the genomic dynamics of early *C. elegans* development": Code and external data: merge_replicate_bam.20240502.sh

```
#!/bin/bash
#
#SBATCH --job-name=sinto
#SBATCH --output=slurm_out/merge_bam.%N.%j.out
#SBATCH --error=slurm_err/merge_bam.%N.%j.err
#SBATCH --ntasks=1
#SBATCH -N 1
#SBATCH -n 8
#SBATCH --mem-per-cpu=4000
#SBATCH --mail-type=END,FAIL
#SBATCH --mail-user=

input_filename=$1
output_path=$2
input_reps=$3

input_reps_all=$(echo $input_reps | tr "," " ")

mkdir -p $output_path
samtools merge -@ 8 $output_path/$input_filename.bam $input_reps_all
samtools index -@ 8 $output_path/$input_filename.bam
```
