## Supplementary material for "A lineage-resolved multimodal single-cell atlas reveals the genomic dynamics of early *C. elegans* development": Code and external data: operon_parser.py

```
import sys

operons_bed=sys.argv[1]
canonical_genes_bed=sys.argv[2]
gene_tx=sys.argv[3]
genes_operons_out=sys.argv[4]

# dictionary of all tx to genes
gene_tx_dict = {}
for line in open(gene_tx):
    G=line.rstrip().split("\t")
    gene_tx_dict[G[1]] = G[0]

# dictionary of most upstream start for each gene
gene_start = {}
for line in open(canonical_genes_bed):
    G=line.rstrip().split("\t")
    if gene_tx_dict[G[3]] not in gene_start:
        if G[5] == "+":
            gene_start[gene_tx_dict[G[3]]] = G[1]
        else:
            gene_start[gene_tx_dict[G[3]]] = G[2]
    else:
        if G[5] == "+":
            if eval(G[1]) < eval(gene_start[gene_tx_dict[G[3]]]):
                gene_start[gene_tx_dict[G[3]]] = G[1]
        else:
            if eval(G[2]) > eval(gene_start[gene_tx_dict[G[3]]]):
                gene_start[gene_tx_dict[G[3]]] = G[2]

new = open(genes_operons_out, "w")
for line in open(operons_bed):
    G=line.rstrip().split("\t")
    operon_dict = {}
    for gene in G[4].split(","):
        if gene not in gene_start:
            print(gene)
            operon_dict = {}
            continue
        operon_dict[gene] = eval(gene_start[gene])
    if G[5] == "+":
        operon_position = 0
        for gene in sorted(operon_dict, key=operon_dict.get):
            operon_position += 1
            new.write(gene + "\t" + G[3] + "\t" + str(operon_position) + "\n")
    else:
        operon_position = 0
        for gene in sorted(operon_dict, key=operon_dict.get, reverse=True):
            operon_position += 1
            new.write(gene + "\t" + G[3] + "\t" + str(operon_position) + "\n")

new.close()
```
