## Supplementary material for "A lineage-resolved multimodal single-cell atlas reveals the genomic dynamics of early *C. elegans* development": Code and external data: operon_promoter_annotation.20240313.py

```
# if genes in operon, assign promoter of first gene also to other downstream genes
import sys

operons=sys.argv[1]
promoter_no_operons=sys.argv[2]
promoters_out=sys.argv[3]
added_genes=sys.argv[4]

operons_genes = {}
for line in open(operons):
    G=line.rstrip().split("\t")
    if G[1] not in operons_genes:
        operons_genes[G[1]] = [G[0]]
    else:
        operons_genes[G[1]].append(G[0])

operon_ordered_genes = {}
for operon in operons_genes:
    operon_ordered_genes[operons_genes[operon][0]] = operons_genes[operon][1:]

new = open(promoters_out, "w")
added_genes_out = open(added_genes, "w")
for line in open(promoter_no_operons):
    G=line.split("\t")
    new.write(line)
    if G[4] in operon_ordered_genes:
        for downstream_gene in operon_ordered_genes[G[4]]:
            added_genes_out.write(downstream_gene + "\n")
            new.write(G[0] + "\t" + G[1] + "\t" + G[2] + "\t" + G[3] + "\t" + downstream_gene + "\t" + G[5])

new.close()
added_genes_out.close()
```
