## Supplementary material for "A lineage-resolved multimodal single-cell atlas reveals the genomic dynamics of early *C. elegans* development": Code and external data: pairwise_clustering.orientation.py

```
import sys
import itertools

input_file = sys.argv[1]
output_dir = sys.argv[2]

flag = 0
motif_pair_orientation = {}
dict_pairwise = {}
for line in open(input_file):
    G = line.rstrip().split("\t")
    if G[0] != "Query_ID" and "#" not in G[0] and G[0] != "" and eval(G[4]) < 0.001:
        flag += 1
        if G[0] not in motif_pair_orientation:
            motif_pair_orientation[G[0]] = {}
        motif_pair_orientation[G[0]][G[1]] = G[9]
        dict_pairwise[flag] = [G[0], G[1]]

singletons=[]
for motif_id in motif_pair_orientation:
    if len(motif_pair_orientation[motif_id]) == 1:
        singletons.append(motif_id)

list_pairwise = []
print(dict_pairwise)
for pair_n in dict_pairwise:
    list_pairwise.append(dict_pairwise[pair_n])


import networkx as nx
G = nx.Graph()
for component in list_pairwise:
    G.add_edges_from(itertools.pairwise(component))

components = (G.subgraph(c) for c in nx.connected_components(G))
comp_dict = {idx: comp.nodes() for idx, comp in enumerate(components)}

opposite_orientation = {"+": "-", "-" : "+"}
for cluster_n in comp_dict:
    print(cluster_n)
    clustered_motifs = list(comp_dict[cluster_n])
    new = open(output_dir + "/" + str(cluster_n) + ".motif_cluster", "w")
    cluster_n_first_motif = clustered_motifs[0]
    cluster_n_motif_orientation = ["+"]
    for additional_motif in clustered_motifs[1:]:
        if additional_motif in motif_pair_orientation[cluster_n_first_motif]:
            cluster_n_motif_orientation.append(motif_pair_orientation[cluster_n_first_motif][additional_motif])
        else:
            cluster_n_motif_orientation.append("NA")
    print("initial_motif_orientation")
    if "NA" in cluster_n_motif_orientation:
        print("deal with NAs")
    while "NA" in cluster_n_motif_orientation:
        #print("iterate NAs")
        na_indices = [i for i, x in enumerate(cluster_n_motif_orientation) if x == "NA"]
        not_na_indices = [i for i, x in enumerate(cluster_n_motif_orientation) if x != "NA"]
        print(cluster_n_motif_orientation)
        print(na_indices)
        print(clustered_motifs)
        for motif_to_rescue_idx in na_indices:
            motif_to_rescue = clustered_motifs[motif_to_rescue_idx]
            for other_motif_idx in not_na_indices:
                other_motif = clustered_motifs[other_motif_idx]
                if other_motif in motif_pair_orientation[motif_to_rescue]:
                    if cluster_n_motif_orientation[other_motif_idx] == "+":
                        cluster_n_motif_orientation[motif_to_rescue_idx] = motif_pair_orientation[motif_to_rescue][other_motif]
                        print(cluster_n_motif_orientation)
                    else:
                        cluster_n_motif_orientation[motif_to_rescue_idx] = opposite_orientation[motif_pair_orientation[motif_to_rescue][other_motif]]
                        print(cluster_n_motif_orientation)
    print("print output")
    for motif_n in range(len(clustered_motifs)):
        new.write(clustered_motifs[motif_n] + "\t" + cluster_n_motif_orientation[motif_n] + "\n")
    new.close()
```
