## Supplementary material for "A lineage-resolved multimodal single-cell atlas reveals the genomic dynamics of early *C. elegans* development": Code and external data: promoter_annotation.20240624.sh

```
#!/bin/bash
#
#SBATCH --job-name=jump
#SBATCH --output=slurm_out/jump.%N.%j.out
#SBATCH --error=slurm_err/jump.%N.%j.err
#SBATCH --ntasks=1
#SBATCH -N 1
#SBATCH -n 1
#SBATCH --mem-per-cpu=8000
#SBATCH --mail-type=END,FAIL
#SBATCH --mail-user=

accessible_sites=$1
output_dir=$2
lcap_rep1=$3
lcap_rep2=$4
pseudoreplicates=$5
pseudoreplicate_sample=$6

lcap_rep1_frags="data/lcap/alignment/"$lcap_rep1"_Aligned.sortedByCoord.out.unique.fragments.bed"
lcap_rep2_frags="data/lcap/alignment/"$lcap_rep2"_Aligned.sortedByCoord.out.unique.fragments.bed"
lcap_rep1_bg_fwd="data/lcap/alignment/"$lcap_rep1"_Aligned.sortedByCoord.out.unique.fragments.fwd.bg"
lcap_rep1_bg_rev="data/lcap/alignment/"$lcap_rep1"_Aligned.sortedByCoord.out.unique.fragments.rev.bg"
lcap_rep2_bg_fwd="data/lcap/alignment/"$lcap_rep2"_Aligned.sortedByCoord.out.unique.fragments.fwd.bg"
lcap_rep2_bg_rev="data/lcap/alignment/"$lcap_rep2"_Aligned.sortedByCoord.out.unique.fragments.rev.bg"

lcap_bg_fwd="data/lcap/alignment/"$pseudoreplicate_sample"_Aligned.sortedByCoord.out.unique.fragments.fwd.bg"
lcap_bg_rev="data/lcap/alignment/"$pseudoreplicate_sample"_Aligned.sortedByCoord.out.unique.fragments.rev.bg"

genome_bed=species/elegans/genome/elegans.bed
tx_id=species/elegans/gene_annotation/c_elegans.PRJNA13758.WS285.canonical_geneset.gene_transcript_id.sorted.txt
tx_id_WB_sorted=species/elegans/gene_annotation/c_elegans.PRJNA13758.WS285.canonical_geneset.gene_transcript_id.WB_sorted.txt
gene_type=species/elegans/gene_annotation/c_elegans.PRJNA13758.WS285.canonical_geneset.gene_type.txt
tx_bed_original=species/elegans/gene_annotation/c_elegans.PRJNA13758.WS285.canonical_geneset.extended_no_ovlp.3prime_extended.rnaseq_corrected.bed
chr_size=species/elegans/genome/elegans.chrom.sizes.txt

mkdir $output_dir
sort -k 4,4 $accessible_sites > $output_dir/all_peaks.bed

tx_bed=$output_dir/c_elegans.PRJNA13758.WS285.canonical_geneset.extended_no_ovlp.3prime_extended.rnaseq_corrected.sorted.bed
sort -k 4,4 $tx_bed_original > $tx_bed

# distinguish accessible sites COMPLETELY within or outside filtered genes (coding, pseudogenes, lincRNAs)
grep -E 'protein_coding|pseudogene|lincRNA' $gene_type | join - $tx_id_WB_sorted | awk '{print $3}' | sort | join -1 1 -2 4 - $tx_bed_original | awk 'BEGIN{OFS="\t";}{print $2, $3, $4}' | intersectBed -a $output_dir/all_peaks.bed -b stdin -f 1 -u | sort -k 4,4 > $output_dir/all_peaks.genic.bed
grep -E 'protein_coding|pseudogene|lincRNA' $gene_type | join - $tx_id_WB_sorted | awk '{print $3}' | sort | join -1 1 -2 4 - $tx_bed_original | awk 'BEGIN{OFS="\t";}{print $2, $3, $4}' | intersectBed -a $output_dir/all_peaks.bed -b stdin -f 1 -v | sort -k 4,4 > $output_dir/all_peaks.intergenic.bed

### 1. jump test
# extract lcap reads up/downstream of any accessible site and test for increase in coverage
echo "preparing jump"
awk 'BEGIN{OFS="\t";}{if ($2 - 200 > 0) print $1, $2-200, $2, $4 "_left\n" $1, $3, $3+200, $4 "_right"; else print $1, 0, $2, $4 "_left\n" $1, $3, $3+200, $4 "_right"}' $output_dir/all_peaks.bed | intersectBed -a stdin -b $genome_bed > $output_dir/all_peaks.left_right_flank.bed

awk '{if ($6 == "+") print}' $lcap_rep1_frags | coverageBed -a $output_dir/all_peaks.left_right_flank.bed -b stdin -counts > $output_dir/all_peaks.left_right_flank.rep1_fwd.bed
awk '{if ($6 == "-") print}' $lcap_rep1_frags | coverageBed -a $output_dir/all_peaks.left_right_flank.bed -b stdin -counts > $output_dir/all_peaks.left_right_flank.rep1_rev.bed
awk '{if ($6 == "+") print}' $lcap_rep2_frags | coverageBed -a $output_dir/all_peaks.left_right_flank.bed -b stdin -counts > $output_dir/all_peaks.left_right_flank.rep2_fwd.bed
awk '{if ($6 == "-") print}' $lcap_rep2_frags | coverageBed -a $output_dir/all_peaks.left_right_flank.bed -b stdin -counts > $output_dir/all_peaks.left_right_flank.rep2_rev.bed

paste $output_dir/all_peaks.left_right_flank.rep1_fwd.bed $output_dir/all_peaks.left_right_flank.rep2_fwd.bed | cut -f 1,2,3,4,5,10 > $output_dir/all_peaks.left_right_flank.fwd.bed
paste $output_dir/all_peaks.left_right_flank.rep1_rev.bed $output_dir/all_peaks.left_right_flank.rep2_rev.bed | cut -f 1,2,3,4,5,10 > $output_dir/all_peaks.left_right_flank.rev.bed

python scripts/left_right_coverage_combined.py $output_dir/all_peaks.left_right_flank.fwd.bed $output_dir/all_peaks.left_right_flank.fwd_combined.bed fwd
python scripts/left_right_coverage_combined.py $output_dir/all_peaks.left_right_flank.rev.bed $output_dir/all_peaks.left_right_flank.rev_combined.bed rev

# NEW: also extract lcap signal WITHIN the accessible site and use it to test a strong jump if the normal jump test fails (e.g. due to a gene transcribed in same orientation immediately upstream of the accessible site)
awk '{if ($6 == "+") print}' $lcap_rep1_frags | coverageBed -a $output_dir/all_peaks.bed -b stdin -counts > $output_dir/all_peaks.AS.rep1_fwd.bed
awk '{if ($6 == "-") print}' $lcap_rep1_frags | coverageBed -a $output_dir/all_peaks.bed -b stdin -counts > $output_dir/all_peaks.AS.rep1_rev.bed
awk '{if ($6 == "+") print}' $lcap_rep2_frags | coverageBed -a $output_dir/all_peaks.bed -b stdin -counts > $output_dir/all_peaks.AS.rep2_fwd.bed
awk '{if ($6 == "-") print}' $lcap_rep2_frags | coverageBed -a $output_dir/all_peaks.bed -b stdin -counts > $output_dir/all_peaks.AS.rep2_rev.bed

paste $output_dir/all_peaks.AS.rep1_fwd.bed $output_dir/all_peaks.AS.rep2_fwd.bed | cut -f 1,2,3,4,5,10 > $output_dir/all_peaks.AS.fwd.bed
paste $output_dir/all_peaks.AS.rep1_rev.bed $output_dir/all_peaks.AS.rep2_rev.bed | cut -f 1,2,3,4,5,10 > $output_dir/all_peaks.AS.rev.bed

python scripts/AS_downstream_coverage_combined.py $output_dir/all_peaks.AS.fwd.bed $output_dir/all_peaks.left_right_flank.fwd.bed $output_dir/all_peaks.AS.fwd_combined.bed fwd
python scripts/AS_downstream_coverage_combined.py $output_dir/all_peaks.AS.fwd.bed $output_dir/all_peaks.left_right_flank.fwd.bed $output_dir/all_peaks.AS.rev_combined.bed rev

# jump test
echo "start jump"
Rscript scripts/lcap_DESeq_jump.R $output_dir/all_peaks.left_right_flank.fwd_combined.bed $output_dir/all_peaks.left_right_flank.fwd.deseq.out
Rscript scripts/lcap_DESeq_jump.R $output_dir/all_peaks.left_right_flank.rev_combined.bed $output_dir/all_peaks.left_right_flank.rev.deseq.out

Rscript scripts/lcap_DESeq_jump.R $output_dir/all_peaks.AS.fwd_combined.bed $output_dir/all_peaks.AS.fwd.deseq.out
Rscript scripts/lcap_DESeq_jump.R $output_dir/all_peaks.AS.rev_combined.bed $output_dir/all_peaks.AS.rev.deseq.out

# jump test for intergenic peaks
sed '1d' $output_dir/all_peaks.left_right_flank.fwd.deseq.out | sort -k 1,1 | join -1 1 -2 4 - $output_dir/all_peaks.intergenic.bed | awk '{if ($3 > 1.5 && $7 < 0.1) print $1 "\tfwd"}' > $output_dir/all_peaks.left_right_flank.jump_test.temp.txt 
sed '1d' $output_dir/all_peaks.left_right_flank.rev.deseq.out | sort -k 1,1 | join -1 1 -2 4 - $output_dir/all_peaks.intergenic.bed | awk '{if ($3 > 1.5 && $7 < 0.1) print $1 "\trev"}' >> $output_dir/all_peaks.left_right_flank.jump_test.temp.txt 

sed '1d' $output_dir/all_peaks.left_right_flank.fwd_combined.bed | join -1 1 -2 4 - $output_dir/all_peaks.intergenic.bed | awk '{if ($2 == 0 && $3 == 0 && $4 > 0 && $5 > 0 && $4 + $5 > 3) print $1 "\tfwd"}' >> $output_dir/all_peaks.left_right_flank.jump_test.temp.txt
sed '1d' $output_dir/all_peaks.left_right_flank.rev_combined.bed | join -1 1 -2 4 - $output_dir/all_peaks.intergenic.bed | awk '{if ($2 == 0 && $3 == 0 && $4 > 0 && $5 > 0 && $4 + $5 > 3) print $1 "\trev"}' >> $output_dir/all_peaks.left_right_flank.jump_test.temp.txt

# jump test for intra-genic peaks
sed '1d' $output_dir/all_peaks.left_right_flank.fwd.deseq.out | sort -k 1,1 | join -1 1 -2 4 - $output_dir/all_peaks.genic.bed | awk '{if ($3 > 1.5 && $7 < 0.05) print $1 "\tfwd"}' >> $output_dir/all_peaks.left_right_flank.jump_test.temp.txt
sed '1d' $output_dir/all_peaks.left_right_flank.rev.deseq.out | sort -k 1,1 | join -1 1 -2 4 - $output_dir/all_peaks.genic.bed | awk '{if ($3 > 1.5 && $7 < 0.05) print $1 "\trev"}' >> $output_dir/all_peaks.left_right_flank.jump_test.temp.txt

# jump test: AS vs downstream coverage
sed '1d' $output_dir/all_peaks.AS.fwd.deseq.out | sort -k 1,1 | join -1 1 -2 4 - $output_dir/all_peaks.bed | awk '{if ($3 > 1.5 && $7 < 0.05) print $1 "\tfwd"}' >> $output_dir/all_peaks.left_right_flank.jump_test.temp.txt
sed '1d' $output_dir/all_peaks.AS.rev.deseq.out | sort -k 1,1 | join -1 1 -2 4 - $output_dir/all_peaks.bed | awk '{if ($3 > 1.5 && $7 < 0.05) print $1 "\trev"}' >> $output_dir/all_peaks.left_right_flank.jump_test.temp.txt

sort $output_dir/all_peaks.left_right_flank.jump_test.temp.txt | uniq | sort -k 1,1 > $output_dir/all_peaks.left_right_flank.jump_test.txt
rm $output_dir/all_peaks.left_right_flank.jump_test.temp.txt

echo "finished jump"

### 2. summit of elongating AS located upstream of a first exon, and at most 250bp downstream of a 5'UTR
# define coding start for protein coding genes, or TSS+250bp for lincRNAs/pseudogenes (use TSS is shorter than 250)
sort -k 1,1 $tx_id | join - $gene_type | grep "lincRNA\|protein_coding\|pseudogene" | awk '{print $2}' | sort | join -1 1 -2 4 - $tx_bed | awk 'BEGIN{OFS="\t";}{if ($7 != $8 && $6 == "+") print $2, $7, $7+1, $1, 0, $6; else if ($7 != $8 && $6 == "-") print $2, $8-1, $8, $1, 0, $6; else if ($7 == $8 && $6 == "+" && $3 + 250 < $4) print $2, $3 + 250, $3 + 251, $1, 0, $6; else if ($7 == $8 && $6 == "+" && $3 + 250 > $4) print $2, $3, $3 + 1, $1, 0, $6; else if ($7 == $8 && $6 == "-" && $3 + 250 < $4) print $2, $4 - 251, $4 - 250, $1, 0, $6; else if ($7 == $8 && $6 == "-" && $3 + 250 > $4) print $2, $4 - 1, $4, $1, 0, $6}' > $output_dir/coding_noncoding_start.bed

sort -k 1,1 -k2,2n $output_dir/coding_noncoding_start.bed > $output_dir/coding_noncoding_start.sorted.bed

# convert bed12 to bed6 for coding/pseudogenes/lincRNAs for the first and any downstream exon
cut -f 4 $output_dir/coding_noncoding_start.bed | join -1 1 -2 4 - $tx_bed | awk 'BEGIN{OFS="\t";}{print $2, $3, $4, $1, $5, $6, $7, $8, $9, $10, $11, $12}' | bed12ToBed6 -i stdin -n | awk 'BEGIN{OFS="\t";}{if ($5 == 1) print $1, $2, $3, $4, 0, $6}' | sort -k 1,1 -k2,2n > $output_dir/coding_noncoding_first_exon.bed
cut -f 4 $output_dir/coding_noncoding_start.bed | join -1 1 -2 4 - $tx_bed | awk 'BEGIN{OFS="\t";}{print $2, $3, $4, $1, $5, $6, $7, $8, $9, $10, $11, $12}' | bed12ToBed6 -i stdin -n | awk 'BEGIN{OFS="\t";}{if ($5 != 1 && $6 == "+") print $1, $2, $2 + 1, $4, 0, $6; else if ($5 != 1 && $6 == "-") print $1, $3 - 1, $3, $4, 0, $6}' | sort -k 1,1 -k2,2n > $output_dir/coding_noncoding_following_exon.bed

# extract the summit of all elongating ASs, then get their closest downstream coding/noncoding start, then confirm that the summits overlaps or lays upstream of the first exon of those transcripts (this will get rid of sites located in an intron downstream of the first exon)
echo "closest exon"
sort -k 4,4 $output_dir/all_peaks.bed | join -1 1 -2 4 $output_dir/all_peaks.left_right_flank.jump_test.txt - | awk 'BEGIN{OFS="\t";}{if ($2 == "fwd") print $3, $4 + 100, $4 + 101, $1, 0, "+"; else print $3, $4 + 100, $4 + 101, $1, 0, "-"}' | sort -k 1,1 -k2,2n > $output_dir/all_peaks.left_right_flank.jump_test.bed

closestBed -a $output_dir/all_peaks.left_right_flank.jump_test.bed -b $output_dir/coding_noncoding_start.sorted.bed -io -iu -D "a" -t "all" -s | awk 'BEGIN{OFS="\t";}{if ($10 != ".") print $4, $6, $10}' | sort -k 3,3 | join -1 3 -2 2 - $tx_id | awk 'BEGIN{OFS="\t";}{print $2, $3, $4}' | sort -k 1,1 > $output_dir/all_peaks.left_right_flank.closest_downstream_start_site.txt

closestBed -a $output_dir/all_peaks.left_right_flank.jump_test.bed -b $output_dir/coding_noncoding_first_exon.bed -iu -D "a" -t "all" -s | awk 'BEGIN{OFS="\t";}{if ($10 != ".") print $4, $6, $7, $8, $9, $10, $11, $12}' | sort -k 6,6 | join -1 6 -2 2 - $tx_id | awk 'BEGIN{OFS="\t";}{print $2, $3, $4, $5, $6, $9, $7, $8}' | sort -k 1,1 > $output_dir/all_peaks.left_right_flank.closest_downstream_first_exon.txt

awk '{if ($2 == "+") print}' $output_dir/all_peaks.left_right_flank.closest_downstream_start_site.txt > $output_dir/all_peaks.left_right_flank.closest_downstream_start_site.fwd.txt
awk '{if ($2 == "-") print}' $output_dir/all_peaks.left_right_flank.closest_downstream_start_site.txt > $output_dir/all_peaks.left_right_flank.closest_downstream_start_site.rev.txt
awk '{if ($2 == "+") print}' $output_dir/all_peaks.left_right_flank.closest_downstream_first_exon.txt > $output_dir/all_peaks.left_right_flank.closest_downstream_first_exon.fwd.txt
awk '{if ($2 == "-") print}' $output_dir/all_peaks.left_right_flank.closest_downstream_first_exon.txt > $output_dir/all_peaks.left_right_flank.closest_downstream_first_exon.rev.txt

join $output_dir/all_peaks.left_right_flank.closest_downstream_start_site.fwd.txt $output_dir/all_peaks.left_right_flank.closest_downstream_first_exon.fwd.txt | awk 'BEGIN{OFS="\t";}{if ($3 == $8) print $1, $2, $5, $6, $7, $8, $9, $10}' > $output_dir/all_peaks.left_right_flank.closest_downstream_first_exon.true.temp.txt
join $output_dir/all_peaks.left_right_flank.closest_downstream_start_site.rev.txt $output_dir/all_peaks.left_right_flank.closest_downstream_first_exon.rev.txt | awk 'BEGIN{OFS="\t";}{if ($3 == $8) print $1, $2, $5, $6, $7, $8, $9, $10}' >> $output_dir/all_peaks.left_right_flank.closest_downstream_first_exon.true.temp.txt

sort -k 1,1 $output_dir/all_peaks.left_right_flank.closest_downstream_first_exon.true.temp.txt > $output_dir/all_peaks.left_right_flank.closest_downstream_first_exon.true.txt

# remove sites if 5' of a non-first exon is located between the AS and the 5' of the first downstream exon
sort -k 4,4 $output_dir/all_peaks.bed | join -1 1 -2 4 $output_dir/all_peaks.left_right_flank.closest_downstream_first_exon.true.txt - | awk 'BEGIN{OFS = "\t";}{if ($2 == "+" && $11 >= $4) print $3, $11, $11, $1, $6, $2; else if ($2 == "+" && $11 < $4) print $3, $11, $4, $1, $6, $2; else if ($2 == "-" && $10 <= $5) print $3, $10, $10, $1, $6, $2; else if ($2 == "-" && $10 > $5) print $3, $5, $10, $1, $6, $2}' | awk 'BEGIN{OFS = "\t";}{if ($3 - $2 < 10000) print}' | intersectBed -a stdin -b $output_dir/coding_noncoding_following_exon.bed -s -v > $output_dir/all_peaks.left_right_flank.closest_downstream_first_exon.true_no_other_exon.span.bed

### 3. require a lower than 200bp non-coverage gap between the accessible site and the downstream first exon
# combine the bedgraph of the lcap replicates and find the gaps in coverage (0-coverage regions longer than 50bp)
echo "preparing maxgap"
if [ "$pseudoreplicates" == "True" ]
then
    awk '{if ($4 == 0 && $3 - $2 > 50) print}' $lcap_bg_fwd > $output_dir/lcap_EE.fwd.gaps.bed
    awk '{if ($4 == 0 && $3 - $2 > 50) print}' $lcap_bg_rev > $output_dir/lcap_EE.rev.gaps.bed
else
    unionBedGraphs -g $chr_size -empty -i $lcap_rep1_bg_fwd $lcap_rep2_bg_fwd | awk 'BEGIN{OFS="\t";}{print $1, $2, $3, $4 + $5}' > $output_dir/lcap_EE.fwd.bg
    unionBedGraphs -g $chr_size -empty -i $lcap_rep1_bg_rev $lcap_rep2_bg_rev | awk 'BEGIN{OFS="\t";}{print $1, $2, $3, $4 + $5}' > $output_dir/lcap_EE.rev.bg
    awk '{if ($4 == 0 && $3 - $2 > 50) print}' $output_dir/lcap_EE.fwd.bg > $output_dir/lcap_EE.fwd.gaps.bed
    awk '{if ($4 == 0 && $3 - $2 > 50) print}' $output_dir/lcap_EE.rev.bg > $output_dir/lcap_EE.rev.gaps.bed
fi

# keep all putative promoter closer than 200bp to the TSS; then exclude from putative promoter list sites were the intervening region between the AS and the transcript's TSS has non-covered (0 lcap coverage) regions longer than 50bp; note, exclude the 200bp downstream of the AS (where the jump test has been done, the lcap signal could actually start a bit downstream to the peak as peak location could be inaccurate)
echo "run maxgap"
awk '{if ($3 - $2 > 200 && $6 == "+") print}' $output_dir/all_peaks.left_right_flank.closest_downstream_first_exon.true_no_other_exon.span.bed | awk 'BEGIN{OFS="\t";}{print $1, $2 + 200, $3, $4, $5, $6}' | intersectBed -a stdin -b $output_dir/lcap_EE.fwd.gaps.bed | awk '{if ($3 - $2 > 50) print $4 "\t" $6}' > $output_dir/all_peaks.left_right_flank.closest_downstream_first_exon.true_no_other_exon.span.gapped_sites.txt.temp
awk '{if ($3 - $2 > 200 && $6 == "-") print}' $output_dir/all_peaks.left_right_flank.closest_downstream_first_exon.true_no_other_exon.span.bed | awk 'BEGIN{OFS="\t";}{print $1, $2, $3 - 200, $4, $5, $6}' | intersectBed -a stdin -b $output_dir/lcap_EE.rev.gaps.bed | awk '{if ($3 - $2 > 50) print $4 "\t" $6}' >> $output_dir/all_peaks.left_right_flank.closest_downstream_first_exon.true_no_other_exon.span.gapped_sites.txt.temp

sort -k 1,1 -k 2,2 $output_dir/all_peaks.left_right_flank.closest_downstream_first_exon.true_no_other_exon.span.gapped_sites.txt.temp | uniq > $output_dir/all_peaks.left_right_flank.closest_downstream_first_exon.true_no_other_exon.span.gapped_sites.txt; rm $output_dir/all_peaks.left_right_flank.closest_downstream_first_exon.true_no_other_exon.span.gapped_sites.txt.temp

python scripts/ungapped_promoter_selection.py $output_dir/all_peaks.left_right_flank.closest_downstream_first_exon.true_no_other_exon.span.bed $output_dir/all_peaks.left_right_flank.closest_downstream_first_exon.true_no_other_exon.span.gapped_sites.txt $output_dir/all_peaks.left_right_flank.closest_downstream_first_exon.true_no_other_exon_no_gap.span.bed

sort -k 4,4 $output_dir/all_peaks.bed | join -1 4 -2 4 - $output_dir/all_peaks.left_right_flank.closest_downstream_first_exon.true_no_other_exon_no_gap.span.bed | awk 'BEGIN{OFS="\t";}{print $2, $3, $4, $1, $8, $9}' | sort -k 1,1 -k2,2n -k 4,4 | uniq > $output_dir/promoters.bed

rm $tx_bed
```
