## Supplementary material for "A lineage-resolved multimodal single-cell atlas reveals the genomic dynamics of early *C. elegans* development": Code and external data: rnaseq_alignment.20240126.sh

```
#!/bin/bash
#
#SBATCH --job-name=star_kallisto
#SBATCH --output=slurm_out/star_kallisto.%N.%j.out
#SBATCH --error=slurm_err/star_kallisto.%N.%j.err
#SBATCH --ntasks=1
#SBATCH -N 1
#SBATCH -n 6
#SBATCH --mem-per-cpu=4000
#SBATCH --mail-type=END,FAIL
#SBATCH --mail-user=

sample=$1

fastq_dir=data/rnaseq/fastq
read1=$fastq_dir"/"$sample"_R1_001.fastq.gz"
read2=$fastq_dir"/"$sample"_R2_001.fastq.gz"

outdir_star=data/rnaseq/alignment
outdir_kallisto=data/rnaseq/kallisto
outdir_fastq_trimmed=data/rnaseq/fastq_trimmed

star_index=star_idx/WS285_extended_no_ovlp
genome_chr=species/elegans/genome/elegans.chrom.sizes.txt
kallisto_idx=kallisto_index/WS285_extended_no_ovlp
gene_transcripts=species/elegans/gene_annotation/c_elegans.PRJNA13758.WS285.canonical_geneset.gene_transcript_id.sorted.txt

mkdir $outdir_star
mkdir $outdir_kallisto
mkdir $outdir_fastq_trimmed

if [ ! -f $outdir_fastq_trimmed"/"$sample_name"_R1_001_val_1.fq.gz" ]
then
    trim_galore --gzip --trim-n -j 4 --paired -o $outdir_fastq_trimmed $read1 $read2
fi

if [ ! -s $outdir_star"/"$sample".Aligned.sortedByCoord.out.bam" ]
then
STAR --genomeDir $star_index --readFilesIn $outdir_fastq_trimmed"/"$sample"_R1_001_val_1.fq.gz" $outdir_fastq_trimmed"/"$sample"_R2_001_val_2.fq.gz" --readFilesCommand zcat --runThreadN 12 --outFileNamePrefix $outdir_star"/"$sample"." --outSAMtype BAM SortedByCoordinate --outWigType wiggle --twopassMode None --limitBAMsortRAM 6274374343 --outSAMattrIHstart 0 --outReadsUnmapped Fastx --alignIntronMax 20000
fi

wigToBigWig $outdir_star/$sample.Signal.UniqueMultiple.str1.out.wig $genome_chr $outdir_star/$sample.Signal.UniqueMultiple.str1.out.bw
wigToBigWig $outdir_star/$sample.Signal.UniqueMultiple.str2.out.wig $genome_chr $outdir_star/$sample.Signal.UniqueMultiple.str2.out.bw
wigToBigWig $outdir_star/$sample.Signal.Unique.str1.out.wig $genome_chr $outdir_star/$sample.Signal.Unique.str1.out.bw
wigToBigWig $outdir_star/$sample.Signal.Unique.str2.out.wig $genome_chr $outdir_star/$sample.Signal.Unique.str2.out.bw

mkdir -p $outdir_kallisto
kallisto quant -i $kallisto_idx -b 100 -o $outdir_kallisto/$sample -t 6 --rf-stranded $read1 $read2
python scripts/sum_TPM_per_gene.py $gene_transcripts $outdir_kallisto/$sample/abundance.tsv $outdir_kallisto/$sample/abundance.genes.txt
sort -k 1,1 $outdir_kallisto/$sample/abundance.genes.txt > $outdir_kallisto/$sample/abundance.genes.txt.sorted
mv $outdir_kallisto/$sample/abundance.genes.txt.sorted $outdir_kallisto/$sample/abundance.genes.txt
```
