## Supplementary material for "A lineage-resolved multimodal single-cell atlas reveals the genomic dynamics of early *C. elegans* development": Code and external data: rnaseq_idx.20240126.sh

```
#!/bin/bash
#
#SBATCH --job-name=star_kallisto_idx
#SBATCH --output=/mnt/home1/ahringer/fnc21/spt2/slurm_out/star_kallisto_idx.out
#SBATCH --error=/mnt/home1/ahringer/fnc21/spt2/slurm_err/star_kallisto_idx.err
#SBATCH --ntasks=1
#SBATCH -N 1
#SBATCH -n 6
#SBATCH --mem-per-cpu=4000
#SBATCH --mail-type=END,FAIL
#SBATCH --mail-user=

genome=species/elegans/genome/elegans.fa
coding_genes=species/elegans/gene_annotation/c_elegans.PRJNA13758.WS285.canonical_geneset.extended_no_ovlp.3prime_extended.rnaseq_corrected.bed
coding_genes_fa=species/elegans/gene_annotation/c_elegans.PRJNA13758.WS285.canonical_geneset.extended_no_ovlp.3prime_extended.rnaseq_corrected.fa
kallisto_idx=kallisto_index/WS285_extended_no_ovlp

fastaFromBed -fi $genome -bed $coding_genes -nameOnly -split -s | awk 'BEGIN{{FS="(";}}{{print $1}}' > $coding_genes_fa
mkdir kallisto_index
kallisto index -i $kallisto_idx $coding_genes_fa
```
