## Supplementary material for "A lineage-resolved multimodal single-cell atlas reveals the genomic dynamics of early *C. elegans* development": Code and external data: second_derivative_binarisation.R

```
# function to determine stes of clusters with no/low/high expression of a given marker gene
second_derivative_binarization = function(exp_vector, bw, modes, gene_name){
  kdde_2 <- ks::kdde(x = exp_vector, h = bw, deriv.order = 2)
  mode_1 = modes[1]
  kdde_2_loc_mode1 = max(which(kdde_2$eval.points <= mode_1))+1
  kdde_2_est_mode1 = kdde_2$estimate[kdde_2_loc_mode1]
  # define point changing direction of second derivative
  for (mode_1_end in c(kdde_2_loc_mode1:length(kdde_2$eval.points))){
    if (kdde_2$estimate[mode_1_end + 1] < kdde_2$estimate[mode_1_end] & kdde_2$estimate[mode_1_end] > 0){
      break
    }
  }
  # define location and estimate of mode_2 in 2nd derivative (shifted by 1 upstream)
  mode_2 = modes[3]
  kdde_2_loc_mode2 = max(which(kdde_2$eval.points <= mode_2))-1
  kdde_2_est_mode2 = kdde_2$estimate[kdde_2_loc_mode2]
  # define point changing direction of second derivative
  for (mode_2_end in seq(kdde_2_loc_mode2, 1, by = -1)){
    if (kdde_2$estimate[mode_2_end - 1] < kdde_2$estimate[mode_2_end] & kdde_2$estimate[mode_2_end] > 0){
      break
    }
  }
  par(mfrow = c(3,1))
  hist(exp_vector, breaks = 50, main = paste0(gene_name, " cluster expression"), las = 1, xlab = "log10(avg exp + 1)", xlim = c(min(kdde_2$eval.points), max(kdde_2$eval.points)))
  plot(density(exp_vector, bw = bw), main = paste0(gene_name, " cluster expression density"), las = 1, xlab = "log10(avg exp + 1)", xlim = c(min(kdde_2$eval.points), max(kdde_2$eval.points)))
  abline(v = modes[c(1,3)], col = c(3, 3))
  abline(v = kdde_2$eval.points[mode_1_end], col = 4, lwd = 2)
  abline(v = kdde_2$eval.points[mode_2_end], col = 4, lwd = 2)
  plot(kdde_2, main = paste0(gene_name, " density second derivative estimation"), las = 1, xlab = "log10(avg exp + 1)", xlim = c(min(kdde_2$eval.points), max(kdde_2$eval.points)))
  abline(v = modes[c(1,3)], col = c(3, 3))
  abline(v = kdde_2$eval.points[mode_1_end], col = 4, lwd = 2)
  abline(v = kdde_2$eval.points[mode_2_end], col = 4, lwd = 2)
  return(c(kdde_2$eval.points[mode_1_end], kdde_2$eval.points[mode_2_end]))
}
```
