## Supplementary material for "A lineage-resolved multimodal single-cell atlas reveals the genomic dynamics of early *C. elegans* development": Code and external data: seurat_integrate_cluster_peaks.20240913.R

```
# upload the cluster-specific peaks in the new merged object
library(ggplot2)
library(dplyr)
library(Seurat)
library(patchwork)
library(RColorBrewer)
library(glmGamPoi)
library(Signac)
library(rtracklayer)
library(Hmisc)

set.seed(42)
args = commandArgs(trailingOnly=TRUE)

combined_seurat_rds = args[1]
merged_peaks_bed = args[2]
new_ATAC_assay = args[3]
final_seurat_rds = args[4]
gene_annotation = args[5]

current_date= paste(unlist(strsplit(as.character(Sys.Date()), "-")), collapse="")
plot_dir_w_date = paste0("plots/", current_date)

dir.create(plot_dir_w_date, recursive = T)

combined_seurat <- readRDS(combined_seurat_rds)

# create new metadata column specifying embryo stage
cell_types_by_stage =list("twocell" = "twocell",
                          "4cell" = c("ABx", "earlyEMS", "EMS"),
                          "8cell" = c("ABax", "ABpx", "earlyEandMS", "E", "MS", "earlyC", "C"),
                          "15cell" = c("earlyABxxx", "ABxxx", "ABala", "ABalp", "ABara", "ABarp", "ABpxa", "ABpxp", "earlyEx", "earlyMSx", "MSx", "MSa", "MSp", "earlyCx"),
                          "26cell" = c("earlyABxxxx", "ABxxxx", "ABalaa", "ABalap", "ABalpa", "ABalpp", "ABaraa", "ABarap", "ABarpa", "ABarpp", "ABpxaa", "ABpxap", "ABpxpa", "ABpxpp", "Ex", "earlyMSxx", "Ca", "Cp", "D"),
                          "46cell" = c("MSxx", "MSxa", "MSxp", "earlyExx", "earlyCxx", "Cxa", "Cxp",
                                        unique(combined_seurat$cell_type)[intersect(grep(pattern = "ABxxxxx", unique(combined_seurat$cell_type)), grep(pattern = "ABxxxxxx", unique(combined_seurat$cell_type), invert = T))]),
                          "87morecell" = c(unique(combined_seurat$cell_type)[grep(pattern = "ABxxxxxx", unique(combined_seurat$cell_type))],
                                            "Exx", "Exxx",
                                            unique(combined_seurat$cell_type)[grep(pattern = "MSxxx", unique(combined_seurat$cell_type))],
                                            unique(combined_seurat$cell_type)[grep(pattern = "MSxxxx", unique(combined_seurat$cell_type))],
                                            unique(combined_seurat$cell_type)[intersect(grep(pattern = "Cxxx", unique(combined_seurat$cell_type)), grep(pattern = "DxxCxxx", unique(combined_seurat$cell_type), invert = T))],
                                            unique(combined_seurat$cell_type)[grep(pattern = "DxxCxxx", unique(combined_seurat$cell_type))],
                                            "Dx"),
                          "Pcell" = c("P2", "P3", "P4"),
                          "Z2Z3" = "Z2Z3",
                          "unassigned" = unique(combined_seurat$cell_type)[grep(pattern = "unassigned", unique(combined_seurat$cell_type))]
)
cell_types_cols = c("black", 
                    "cyan", "darkorange", "darkorange1",
                    "lightblue1", "lightblue3", "darkorange3", "palegreen", "indianred1", "plum", "plum3",
                    "lightskyblue1", "lightskyblue2", "lightskyblue3", "lightskyblue4", "lightskyblue3", "lightskyblue4", "lightskyblue3", "lightskyblue4", "palegreen2", "indianred1", "indianred2", "indianred3", "indianred4", "mediumpurple1",
                    "deepskyblue1", "deepskyblue2", "deepskyblue3", "deepskyblue4", "deepskyblue3", "deepskyblue4", "deepskyblue3", "deepskyblue4", "deepskyblue3", "deepskyblue4", "deepskyblue3", "deepskyblue4", "deepskyblue3", "deepskyblue4", "palegreen3", "red1", "mediumpurple3", "mediumpurple4", "palevioletred2",
                    "red2", "red3", "red4", "seagreen1", "purple1", "purple3", "purple4",
                    rep("royalblue2", length(unique(combined_seurat$cell_type)[grep(pattern = "ABxxxxx_", unique(combined_seurat$cell_type))])),
                    rep("royalblue4", length(unique(combined_seurat$cell_type)[grep(pattern = "ABxxxxxx_", unique(combined_seurat$cell_type))])),
                    "seagreen3", "seagreen4",
                    rep("orangered2",length(unique(combined_seurat$cell_type)[grep(pattern = "MSxxx_", unique(combined_seurat$cell_type))])),
                    rep("orangered4", length(unique(combined_seurat$cell_type)[grep(pattern = "MSxxxx_", unique(combined_seurat$cell_type))])),
                    rep("plum1", length(unique(combined_seurat$cell_type)[intersect(grep(pattern = "Cxxx_", unique(combined_seurat$cell_type)), grep(pattern = "DxxCxxx_", unique(combined_seurat$cell_type), invert = T))])),
                    rep("plum4", length(unique(combined_seurat$cell_type)[grep(pattern = "DxxCxxx_", unique(combined_seurat$cell_type))])),
                    "palevioletred4",
                    "gold",
                    "deeppink2",
                    rep("grey80", length(unique(combined_seurat$cell_type)[grep(pattern = "unassigned", unique(combined_seurat$cell_type))]))
)


stage_metadata = combined_seurat$cell_type
stage_metadata[combined_seurat$cell_type %in% cell_types_by_stage[["twocell"]]] = "twocell"
stage_metadata[combined_seurat$cell_type %in% cell_types_by_stage[["4cell"]]] = "4cell"
stage_metadata[combined_seurat$cell_type %in% cell_types_by_stage[["8cell"]]] = "8cell"
stage_metadata[combined_seurat$cell_type %in% cell_types_by_stage[["15cell"]]] = "15cell"
stage_metadata[combined_seurat$cell_type %in% cell_types_by_stage[["26cell"]]] = "26cell"
stage_metadata[combined_seurat$cell_type %in% cell_types_by_stage[["46cell"]]] = "46cell"
stage_metadata[combined_seurat$cell_type %in% cell_types_by_stage[["87morecell"]]] = "87morecell"
stage_metadata[combined_seurat$cell_type %in% cell_types_by_stage[["Pcell"]]] = "Pcell"
stage_metadata[combined_seurat$cell_type %in% cell_types_by_stage[["Z2Z3"]]] = "Z2Z3"
stage_metadata[combined_seurat$cell_type %in% cell_types_by_stage[["unassigned"]]] = "unassigned"

combined_seurat$stage = stage_metadata

# generate metadata of stems and tips
#stem_cell_types = c("ABxxx" = "early_ABxxx", "ABxxxx" = "early_ABxxxx", "C" = "early_C", "Cx" = "early_Cx","Cxx" = "early_Cxx",
#                    "E_MS" = "early_E_MS","EMS" = "early_EMS","Ex" = "early_Ex","Exx" = "early_Exx","MSx" = "early_MSx","MSxx" = "early_MSxx")
#tip_cell_types = list("ABxxx" = c("ABala", "ABalp", "ABara", "ABarp", "ABpxa", "ABpxp"),
#                      "ABxxxx" = c("ABalaa", "ABalap", "ABalpa", "ABalpp", "ABaraa", "ABarap","ABarpa", "ABarpp",
#                                   "ABpxaa", "ABpxap", "ABpxpa", "ABpxpp"),"C" = "C","Cx" = c("Ca", "Cp"),"Cxx" = c("Cxa", "Cxp"),"D" = "D",
#                      "E_MS" = c("E", "MS"),"EMS" = "EMS","Ex" = "Ex","Exx" = "Exx","MSx" = c("MSa", "MSp"),"MSxx" = c("MSxa", "MSxp"))

#stem_tip = as.character(combined_seurat$cell_type)
#stem_tip[which(stem_tip %in% unlist(stem_cell_types))] = "stem"
#stem_tip[stem_tip %in% unlist(tip_cell_types)] = "tip"
#stem_tip[stem_tip %nin% c("stem", "tip")] = "other"
#combined_seurat$stem_tip = stem_tip


# add new peaks
merged_peaks = import.bed(merged_peaks_bed)

merged_peaks_matrix_filtered <- FeatureMatrix(combined_seurat[["WNN"]]@fragments, sep = c("-", "-"), features=merged_peaks, cells = colnames(combined_seurat))

combined_seurat[[new_ATAC_assay]] <- CreateChromatinAssay(
  counts = merged_peaks_matrix_filtered,
  sep = c(":", "-"),
  fragments = combined_seurat[["WNN"]]@fragments)

# output fragment files per cell type/seurat cluster
DefaultAssay(combined_seurat) <- new_ATAC_assay
combined_seurat <- RunTFIDF(combined_seurat)

# add FRiP information
tot_frag = CountFragments(list("data/sc/cellranger_arc/exp031/outs/atac_fragments_barcode_corrected.tsv.gz",
                               "data/sc/cellranger_arc/exp032/outs/atac_fragments_barcode_corrected.tsv.gz",
                               "data/sc/cellranger_arc/exp042/outs/atac_fragments_barcode_corrected.tsv.gz",
                               "data/sc/cellranger_arc/exp043/outs/atac_fragments_barcode_corrected.tsv.gz",
                               "data/sc/cellranger_arc/exp024/outs/atac_fragments_barcode_corrected.tsv.gz",
                               "data/sc/cellranger_arc/exp025/outs/atac_fragments_barcode_corrected.tsv.gz",
                               "data/sc/cellranger_arc/exp027/outs/atac_fragments_barcode_corrected.tsv.gz"), cells = colnames(combined_seurat), max_lines = NULL, verbose = TRUE)
combined_seurat$tot_frag = tot_frag$frequency_count
combined_seurat = FRiP(object = combined_seurat, assay = "final_peakset", total.fragments = "tot_frag", col.name = "FRiP")

# save RDS
saveRDS(combined_seurat, file = final_seurat_rds)
```
