## Supplementary material for "A lineage-resolved multimodal single-cell atlas reveals the genomic dynamics of early *C. elegans* development": Code and external data: seurat_soupx.20240207.R

```
library(ggplot2)
library(dplyr)
library(Seurat)
library(Signac)
library(patchwork)
library(DropletUtils)
library(SoupX)
library(pheatmap)
library(RColorBrewer)
library(scDblFinder)
library(scater)
library(Hmisc)
library(rtracklayer)
library(glmGamPoi)	
library(multimode)

source("scripts/second_derivative_binarisation.R")

args = commandArgs(trailingOnly=TRUE)

sample_id <- args[1]
barhop_rna <- args[2]
lower_rna <- args[3]
annotation <- args[4]
min_frags = as.numeric(args[5])
merged_peaks_file = args[6]

ED_barcodes = paste0("data/sc/rnaseq/", sample_id, "/ED_RNA_", annotation, "/ED_RNA_barcodes.txt")
set.seed(42)

current_date= paste(unlist(strsplit(as.character(Sys.Date()), "-")), collapse="")
plot_dir_w_date = paste0("/mnt/home3/ahringer/fnc21/10x/plots/", current_date)
dir.create(plot_dir_w_date, recursive=TRUE)

raw_matrix_fran = paste0("data/sc/rnaseq/",sample_id, "/star_", annotation, "/star_", annotation, "_Solo.out/GeneFull/raw/um_reads")
rnaseq_data_raw <- Seurat::Read10X(raw_matrix_fran)

# read in the barcodes obtained by EmptyDrops Multi and subset the object
combined_barcodes <- read.table(ED_barcodes)
rnaseq_data <- rnaseq_data_raw[,colnames(rnaseq_data_raw) %in% combined_barcodes$V1]
combined_data_seurat <- CreateSeuratObject(counts = rnaseq_data, project = sample_id )

### inspect mitochondrial contamination
# get mitochondrial genes
mt_genes <- read.table("species/elegans/gene_annotation/c_elegans.PRJNA13758.WS285.canonical_geneset.filtered.MtDNA_genes.genes")
gene_type = read.table("species/elegans/gene_annotation/c_elegans.PRJNA13758.WS285.canonical_geneset.gene_name_gene_type.txt")

mt_genes_expression_pct = colSums(rnaseq_data_raw[row.names(rnaseq_data_raw) %in% mt_genes$V1,])/colSums(rnaseq_data_raw)

#rrna_genes <-  gene_type$V1[gene_type$V2  == "rRNA" & gene_type$V1 %nin% mt_genes$V1]
#rrna_genes_expression_pct = colSums(rnaseq_data_raw[row.names(rnaseq_data_raw) %in% rrna_genes,])/colSums(rnaseq_data_raw)

sce.rnaseq_data <- as.SingleCellExperiment(combined_data_seurat)
is.mito <- rownames(sce.rnaseq_data) %in% mt_genes$V1
sce.rnaseq_data <- addPerCellQC(sce.rnaseq_data, subsets=list(Mt=is.mito))
summary(sce.rnaseq_data$subsets_Mt_percent == 0)

stats <- quickPerCellQC(colData(sce.rnaseq_data), sub.fields="subsets_Mt_percent")

pdf(file=paste0(plot_dir_w_date, "/", sample_id, ".", annotation, ".mitochondrial_pct_all_barcodes.pdf"), width=8, height = 8)
plotColData(sce.rnaseq_data, y="subsets_Mt_percent",
            colour_by=I(stats$high_subsets_Mt_percent))
dev.off()

#mt_pct_threshold = attr(stats$high_subsets_Mt_percent,"thresholds")[2]
mt_pct_threshold = 5
high_mt_cells = colnames(sce.rnaseq_data[,sce.rnaseq_data$subsets_Mt_percent >= mt_pct_threshold])

# remove high-Mito cells
rnaseq_data <- rnaseq_data[,colnames(rnaseq_data) %nin% high_mt_cells]

# remove MT genes, rRNAs and small RNAs from matrix
genes_to_retain = gene_type$V1[gene_type$V2 %in% c("lincRNA", "protein_coding", "pseudogene") & gene_type$V1 %nin% mt_genes$V1]

rnaseq_data <- rnaseq_data[row.names(rnaseq_data) %in% genes_to_retain,]

### remove cells wih low or too high ATAC fragment counts
# load fragment count matrix from CellRanger
counts <- Read10X_h5(paste0("data/sc/cellranger_arc/", sample_id, "/outs/raw_feature_bc_matrix.h5"))
colnames(counts$Peaks) <- sapply(strsplit(colnames(counts$Peaks), split='-', fixed=TRUE), `[`, 1)
atac_count = counts$Peaks

# retain barcodes with a minimum number of fragments
min_frags_barcodes = colnames(atac_count[,colSums(atac_count) > min_frags])
atac_count = atac_count[,colnames(atac_count) %in% min_frags_barcodes]

# remove barcodes with a number of fragments exceeding 5 MADs
high_frags_barcodes = names(colSums(atac_count)[which(isOutlier(colSums(atac_count), nmads = 5))])
counts = NULL
atac_count = NULL

rnaseq_data <- rnaseq_data[,colnames(rnaseq_data) %in% min_frags_barcodes & colnames(rnaseq_data) %nin% high_frags_barcodes]

# Initialize the Seurat object with the raw (non-normalized data).
combined_data_seurat <- CreateSeuratObject(counts = rnaseq_data, project = sample_id)

combined_data_seurat <- SCTransform(combined_data_seurat, vst.flavor = "v2", method = "glmGamPoi")

combined_data_seurat <- RunPCA(combined_data_seurat, seed.use=42, features = VariableFeatures(object = combined_data_seurat))
combined_data_seurat <- FindNeighbors(combined_data_seurat, dims = 1:50)
combined_data_seurat <- FindClusters(combined_data_seurat, resolution = 2, random.seed = 0)
combined_data_seurat <- RunUMAP(combined_data_seurat, dims = 1:50, seed.use = 42)

pdf(file=paste0(plot_dir_w_date, "/", sample_id, ".", annotation, ".umap_pre_doublets.50PCA.pdf"), width=8, height = 8)
DimPlot(combined_data_seurat, reduction = "umap", label=T)
dev.off()

# identify doublets
sce <- as.SingleCellExperiment(combined_data_seurat)

set.seed(42)
sce.mam.dbl <- scDblFinder(sce, clusters=colData(sce)$ident)
pdf(file=paste0(plot_dir_w_date, "/", sample_id, ".", annotation, ".seurat.scDblFinder.scores.pdf"), width=8, height = 8)
plotUMAP(sce.mam.dbl, colour_by="scDblFinder.score")
dev.off()
table(sce.mam.dbl$scDblFinder.class)

# extract singlets, then re-cluster
singlet_cells = colnames(sce.mam.dbl[,sce.mam.dbl$scDblFinder.class == "singlet"])

rnaseq_data_nodub = rnaseq_data[,colnames(rnaseq_data) %in% singlet_cells]

### additionally, remove cells wih extremely high RNA counts
rnaseq_data_nodub = rnaseq_data_nodub[,colSums(rnaseq_data_nodub) < 10000]

combined_data_seurat_nodoub <- CreateSeuratObject(counts = rnaseq_data_nodub, project = sample_id)

combined_data_seurat_nodoub <- SCTransform(combined_data_seurat_nodoub, vst.flavor = "v2", method = "glmGamPoi")
#combined_data_seurat_nodoub <- SCTransform(combined_data_seurat_nodoub)
combined_data_seurat_nodoub <- RunPCA(combined_data_seurat_nodoub, seed.use=42)
combined_data_seurat_nodoub <- FindNeighbors(combined_data_seurat_nodoub, dims = 1:50)
combined_data_seurat_nodoub <- FindClusters(combined_data_seurat_nodoub, resolution = 2, random.seed = 0)
combined_data_seurat_nodoub <- RunUMAP(combined_data_seurat_nodoub, dims = 1:50, seed.use = 42)

pdf(file=paste0(plot_dir_w_date, "/", sample_id, ".", annotation, ".umap_pre_soupx.50PCA.pdf"), width=8, height = 8)
DimPlot(combined_data_seurat_nodoub, reduction = "umap", label=T)
dev.off()

# pass clustering to SoupX to clean the data
# rnaseq_data_raw_filtered <- rnaseq_data_raw[row.names(rnaseq_data_raw) %in% genes_to_retain,]
# 
# sc_all = SoupChannel(rnaseq_data_raw_filtered, rnaseq_data_nodub, calcSoupProfile=F)
# sc_all = estimateSoup(sc_all,soupRange=c(as.numeric(barhop_rna) ,as.numeric(lower_rna))  )
# 
# # new: remove clusters whose markers contribute the most to the soup
# ### get top 5% genes by soup expression
# sorted_sc = sc_all$soupProfile[order(sc_all$soupProfile$est, decreasing = T), ]
# high_soup_exp = row.names(sorted_sc[c(1:ceiling(dim(sc_all$soupProfile)[1]/20)),])
# 
# ### run marker enrichment 
# marker_genes = quickMarkers(rnaseq_data_nodub, clusters = as.vector(combined_data_seurat_nodoub$seurat_clusters), N = 100, FDR = 0.05, expressCut = 0.9)
# marker_enrichment = data.frame(matrix(NA, ncol = 2, nrow = length(unique(marker_genes$cluster)), 
#                            dimnames = list(c(0:(length(unique(marker_genes$cluster))-1)), c("all_markers", "ambient_markers"))))
# 
# for (cl_name in c(0:(dim(marker_enrichment)[1]-1))){
#   marker_enrichment[cl_name+1,1] = dim(marker_genes[marker_genes$cluster == cl_name,])[1]
#   marker_enrichment[cl_name+1,2] = dim(marker_genes[marker_genes$cluster == cl_name & marker_genes$gene %in% high_soup_exp,])[1]
# }
# 
# ### flag clusters with: a) fewer than 20 markers; b) more than 10% markers among the high ambient genes
# clusters_to_remove = rownames(marker_enrichment[marker_enrichment$all_markers < 20 | marker_enrichment$ambient_markers/marker_enrichment$all_markers > 0.1,])
# clusters_to_remove = c("10", clusters_to_remove)
# ### remove barcodes from flagged clusters from input matrix, then create new SoupX object and run SoupX analysis
# flagged_barcodes = colnames(combined_data_seurat_nodoub[,combined_data_seurat_nodoub$seurat_clusters %in% clusters_to_remove])
# rnaseq_data_nodub_filtered = rnaseq_data_nodub[,colnames(rnaseq_data_nodub) %nin% flagged_barcodes]
# 
# sc = SoupChannel(rnaseq_data_raw_filtered, rnaseq_data_nodub_filtered, calcSoupProfile=F)
# sc = estimateSoup(sc,soupRange=c(as.numeric(barhop_rna) ,as.numeric(lower_rna))  )
# 
# meta    <- combined_data_seurat_nodoub@meta.data
# umap    <- combined_data_seurat_nodoub@reductions$umap@cell.embeddings
# sc  <- setClusters(sc, setNames(meta$seurat_clusters, rownames(meta)))
# sc  <- setDR(sc, umap)
# 
# # automatic estimate
# par(mfrow=c(1,1), mar=c(6, 4, 3, 3), oma=c(0,0,0,0))
# pdf(file=paste0(plot_dir_w_date, "/",sample_id,".soupx_contamination_rate.pdf"), width=8, height = 8)
# 
# markers_soupx <- quickMarkers(rnaseq_data_nodub_filtered, combined_data_seurat_nodoub$seurat_clusters[combined_data_seurat_nodoub$seurat_clusters %nin% clusters_to_remove],
#                              N = 100, FDR = 0.01, expressCut = 0.9)
# # reorder and filter marker table 
# markers_soupx = markers_soupx[order(markers_soupx$gene,-markers_soupx$tfidf),]
# markers_soupx = markers_soupx[!duplicated(markers_soupx$gene),]
# markers_soupx = markers_soupx[order(-markers_soupx$tfidf),]
# markers_soupx = markers_soupx[markers_soupx$tfidf > 1,]
# 
# sc  <- autoEstCont(sc, topMarkers = markers_soupx, soupQuantile = 0.8, maxMarkers = 100)
# 
# sc  <- autoEstCont(sc, soupQuantile = 0.8)

# new: remove P-cell clusters
# get average expression per cluster of P-granule genes
rnaseq_data_raw_filtered <- rnaseq_data_raw[row.names(rnaseq_data_raw) %in% genes_to_retain,]
P_cells = read.table("data/external_data/P_granule_transcripts.sorted.txt")

combined_data_seurat_nodoub = AddModuleScore(combined_data_seurat_nodoub, features = list(P_cells$V1), nbin = 10)

DefaultAssay(combined_data_seurat_nodoub) = "RNA"
combined_data_seurat_nodoub_avgExp = as.data.frame(AverageExpression(combined_data_seurat_nodoub, slot = "data", assays = "RNA", features = P_cells$V1, group.by = "seurat_clusters"))
combined_data_seurat_nodoub_avgExp_log = log10(combined_data_seurat_nodoub_avgExp + 1)
combined_data_seurat_nodoub_avgExp_log_avg = apply(combined_data_seurat_nodoub_avgExp_log, 2, mean)
combined_data_seurat_nodoub_avgExp_quant = combined_data_seurat_nodoub_avgExp_log_avg

pdf(file = paste0(plot_dir_w_date, "/", sample_id, ".", annotation, ".P_cell_binarisation.pdf"), width = 8, height = 8, useDingbats = F)

loc <- locmodes(as.numeric(combined_data_seurat_nodoub_avgExp_log_avg), mod0 = 2, display = F)
low_high_exp_loc = second_derivative_binarization(as.numeric(combined_data_seurat_nodoub_avgExp_log_avg), loc$cbw$bw, loc$locations, "P_granule_genes")
combined_data_seurat_nodoub_avgExp_quant[which(as.numeric(combined_data_seurat_nodoub_avgExp_log_avg) < low_high_exp_loc[1])] <- 0
combined_data_seurat_nodoub_avgExp_quant[which(as.numeric(combined_data_seurat_nodoub_avgExp_log_avg) >= low_high_exp_loc[1] & as.numeric(combined_data_seurat_nodoub_avgExp_log_avg) < low_high_exp_loc[2])] <- 1
combined_data_seurat_nodoub_avgExp_quant[which(as.numeric(combined_data_seurat_nodoub_avgExp_log_avg) >= low_high_exp_loc[2])] <- 2

dev.off()

cl_n = max(as.numeric(combined_data_seurat_nodoub$seurat_clusters))-1
names(combined_data_seurat_nodoub_avgExp_quant) = c(0:cl_n)
#DefaultAssay(combined_data_seurat_nodoub) = "SCT"
#combined_data_seurat_nodoub = AddModuleScore(combined_data_seurat_nodoub, features = list(P_cells$V1), nbin = 10)

# flag clusters if > 33% of their cells have an P-cell module score > 3*sd above the global average P-cell module score
# threshold_P_cell = sd(combined_data_seurat_nodoub$Cluster1)*3+mean(combined_data_seurat_nodoub$Cluster1)
# P_cell_fraction = data.frame(matrix(NA, ncol = 2, nrow = length(unique(combined_data_seurat_nodoub$seurat_clusters)), 
#                                     dimnames = list(c(0:(length(unique(combined_data_seurat_nodoub$seurat_clusters))-1)), c("all_cells", "putative_P_cells"))))
# 
# combined_data_seurat_nodoub_df = combined_data_seurat_nodoub@meta.data
# 
# for (cl_name in c(0:(length(unique(combined_data_seurat_nodoub_df$seurat_clusters))-1))){
#   P_cell_fraction[cl_name+1,1] = dim(combined_data_seurat_nodoub_df[combined_data_seurat_nodoub_df$seurat_clusters == cl_name,])[1]
#   P_cell_fraction[cl_name+1,2] = dim(combined_data_seurat_nodoub_df[combined_data_seurat_nodoub_df$seurat_clusters == cl_name & combined_data_seurat_nodoub_df$Cluster1 > threshold_P_cell,])[1]
# }

#clusters_to_remove = rownames(P_cell_fraction[P_cell_fraction$putative_P_cells/P_cell_fraction$all_cells > 0.33,])
clusters_to_remove = names(combined_data_seurat_nodoub_avgExp_quant[combined_data_seurat_nodoub_avgExp_quant == 2])

pdf(file=paste0(plot_dir_w_date, "/", sample_id, ".", annotation, ".seurat_P_cell_clusters.noDub.pdf"), width=16, height = 8)
p1 = DimPlot(combined_data_seurat_nodoub, reduction = "umap", label=T, cells.highlight = colnames(combined_data_seurat_nodoub[,combined_data_seurat_nodoub$seurat_clusters %in% clusters_to_remove]), pt.size = 0.3, sizes.highlight = 0.3) & NoLegend()
p2 = FeaturePlot(combined_data_seurat_nodoub, reduction = "umap", features = "Cluster1")  & scale_colour_gradientn(colours = c("grey90", "cyan", "deepskyblue", "forestgreen", "darkorange", "darkred"))
p1 | p2
dev.off()

### remove barcodes from flagged clusters from input matrix, then create new SoupX object and run SoupX analysis
flagged_barcodes = colnames(combined_data_seurat_nodoub[,combined_data_seurat_nodoub$seurat_clusters %in% clusters_to_remove])
rnaseq_data_nodub_filtered = rnaseq_data_nodub[,colnames(rnaseq_data_nodub) %nin% flagged_barcodes]
combined_data_seurat_nodoub_filtered = subset(x = combined_data_seurat_nodoub, idents = clusters_to_remove, invert = TRUE)

sc = SoupChannel(rnaseq_data_raw_filtered, rnaseq_data_nodub_filtered, calcSoupProfile=F)
sc = estimateSoup(sc,soupRange=c(as.numeric(barhop_rna) ,as.numeric(lower_rna))  )

meta    <- combined_data_seurat_nodoub_filtered@meta.data
umap    <- combined_data_seurat_nodoub_filtered@reductions$umap@cell.embeddings
sc  <- setClusters(sc, setNames(meta$seurat_clusters, rownames(meta)))
sc  <- setDR(sc, umap)

print("SoupX object ready")
# automatic estimate
# set marker genes
# markers_soupx <- quickMarkers(rnaseq_data_nodub_filtered, combined_data_seurat_nodoub$seurat_clusters[combined_data_seurat_nodoub$seurat_clusters %nin% clusters_to_remove],
#                               N = 100, FDR = 0.01, expressCut = 0.9)
# markers_soupx = markers_soupx[order(markers_soupx$gene,-markers_soupx$tfidf),]
# markers_soupx = markers_soupx[!duplicated(markers_soupx$gene),]
# markers_soupx = markers_soupx[order(-markers_soupx$tfidf),]
# markers_soupx = markers_soupx[markers_soupx$tfidf > 1,]
# 
# sc  <- autoEstCont(sc, topMarkers = markers_soupx, soupQuantile = 0.8, maxMarkers = 100)

pdf(file=paste0(plot_dir_w_date, "/",sample_id,".", annotation, ".soupx_contamination_rate.pdf"), width=8, height = 8)
par(mfrow=c(1,1), mar=c(6, 4, 3, 3), oma=c(0,0,0,0))
sc  <- autoEstCont(sc, soupQuantile = 0.80)
dev.off()

print("SoupX complete")

estimated_contamination = unique(sc$metaData$rho)
#head(sc$soupProfile[order(sc$soupProfile$est, decreasing = T), ], n = 20)

# create a new SoupChannel object and use the estimated_contamination for count correction
# step needed to re-introduce P-cells in object
sc_full = SoupChannel(rnaseq_data_raw_filtered, rnaseq_data_nodub, calcSoupProfile=F)
sc_full = estimateSoup(sc_full,soupRange=c(as.numeric(barhop_rna) ,as.numeric(lower_rna))  )

meta    <- combined_data_seurat_nodoub@meta.data
umap    <- combined_data_seurat_nodoub@reductions$umap@cell.embeddings
sc_full  <- setClusters(sc_full, setNames(meta$seurat_clusters, rownames(meta)))
sc_full  <- setDR(sc_full, umap)

sc_full = setContaminationFraction(sc_full, estimated_contamination)

adj.matrix  <- adjustCounts(sc_full, roundToInt = T)

pdf(file=paste0(plot_dir_w_date, "/",sample_id,".", annotation, ".soupx_decontamination.", estimated_contamination, ".vet-6.binary.pdf"), width=16, height = 8)
plotChangeMap(sc, adj.matrix, geneSet = c("vet-6"), dataType = "binary")
dev.off()

# re-run clustering with Seurat
combined_data_seurat_soupx <- CreateSeuratObject(counts = adj.matrix, project = paste0(sample_id,"-soupX") )

combined_data_seurat_soupx <- SCTransform(combined_data_seurat_soupx, vst.flavor = "v2", method = "glmGamPoi")
combined_data_seurat_soupx <- RunPCA(combined_data_seurat_soupx, seed.use = 42, features = VariableFeatures(object = combined_data_seurat_soupx), npcs = 100)
ElbowPlot(combined_data_seurat_soupx, ndims = 100)

combined_data_seurat_soupx <- FindNeighbors(combined_data_seurat_soupx, dims = 1:50)
combined_data_seurat_soupx <- FindClusters(combined_data_seurat_soupx, resolution = 3, random.seed = 0)
combined_data_seurat_soupx <- RunUMAP(combined_data_seurat_soupx, seed.use = 42, dims = 1:50, return.model=TRUE, reduction.name = "umap_SCT")

# save seurat object
seurat_objects_dir = "seurat_objects/"
#saveRDS(combined_data_seurat_soupx, file = paste0(seurat_objects_dir, sample_id, "_postSoupX.", current_date, ".rds"))
write.table(combined_data_seurat_soupx[[]], file = paste0(seurat_objects_dir, sample_id, ".", annotation, ".postSoupX.", current_date, ".table.txt"), quote = F, sep = "\t", row.names = T, col.names = T)

pdf(file=paste0(plot_dir_w_date, "/",sample_id,".", annotation, ".umap_post_soupx.", estimated_contamination, "_soupX.50PCA.pdf"), width=8, height = 8)
DimPlot(combined_data_seurat_soupx, reduction = "umap_SCT", group.by = "seurat_clusters", label = TRUE, label.size = 4)
dev.off()

pdf(file=paste0(plot_dir_w_date, "/",sample_id,".", annotation, ".umap_post_soupx.", estimated_contamination, "_soupX.markers.pdf"), width=16, height = 16)
FeaturePlot(combined_data_seurat_soupx, order = T, features = c("med-1", "end-3", "tbx-35", "elt-7", "pes-10", "lsl-1")) & scale_colour_gradientn(colours = c("grey90", "cyan", "deepskyblue", "forestgreen", "darkorange", "darkred"))
FeaturePlot(combined_data_seurat_soupx, order = T, features = c("end-3", "end-1", "elt-7", "elt-2", "ceh-51", "lsl-1")) & scale_colour_gradientn(colours = c("grey90", "cyan", "deepskyblue", "forestgreen", "darkorange", "darkred"))
FeaturePlot(combined_data_seurat_soupx, order = T, features = c("tbx-33", "tbx-38", "tbx-40", "sdz-20", "pes-10", "vet-2")) & scale_colour_gradientn(colours = c("grey90", "cyan", "deepskyblue", "forestgreen", "darkorange", "darkred"))
dev.off()

lincRNA_genes <- gene_type$V1[gene_type$V2  == "lincRNA" & gene_type$V1 %nin% mt_genes$V1]

#combined_data_seurat_soupx = AddModuleScore(combined_data_seurat_soupx, features = list(c(lincRNA_genes)), assay = "RNA", name = "lincRNA_RNA", nbin = 10)
#combined_data_seurat_soupx = AddModuleScore(combined_data_seurat_soupx, features = list(c(lincRNA_genes)), assay = "SCT", name = "lincRNA_SCT")
combined_data_seurat_soupx$mito_pct <-  mt_genes_expression_pct[names(mt_genes_expression_pct) %in% colnames(combined_data_seurat_soupx)]
#combined_data_seurat_soupx$rrna_pct <-  rrna_genes_expression_pct[names(rrna_genes_expression_pct) %in% colnames(combined_data_seurat_soupx)]

pdf(file=paste0(plot_dir_w_date, "/",sample_id,".", annotation, ".umap_post_soupx.", estimated_contamination, "_soupX.qc_bad_clusters.pdf"), width=16, height = 16)
#FeaturePlot(combined_data_seurat_soupx, order = T, features = c("mito_pct", "rrna_pct", "lincRNA_SCT1", "nFeature_RNA")) & scale_colour_gradientn(colours = c("grey90", "cyan", "deepskyblue", "forestgreen", "darkorange", "darkred"))
FeaturePlot(combined_data_seurat_soupx, order = T, features = c("mito_pct", "nFeature_RNA")) & scale_colour_gradientn(colours = c("grey90", "cyan", "deepskyblue", "forestgreen", "darkorange", "darkred"))
dev.off()

# add ATAC data
counts <- Read10X_h5(paste0("data/sc/cellranger_arc/", sample_id, "/outs/raw_feature_bc_matrix.h5"))
colnames(counts$Peaks) <- sapply(strsplit(colnames(counts$Peaks), split='-', fixed=TRUE), `[`, 1)
merged_peaks = import.bed(merged_peaks_file)
fragpath <- paste0("data/sc/cellranger_arc/", sample_id, "/outs/atac_fragments_no_dash.tsv.gz")

fragments_exp031 <- CreateFragmentObject(fragpath, cells = colnames(combined_data_seurat_soupx))
merged_peaks_matrix_filtered <- FeatureMatrix(fragments_exp031, sep = c("-", "-"), features=merged_peaks, cells = colnames(combined_data_seurat_soupx))

combined_data_seurat_soupx[["ATAC"]] <- CreateChromatinAssay(
  counts = merged_peaks_matrix_filtered,
  sep = c(":", "-"),
  fragments = fragments_exp031)

DefaultAssay(combined_data_seurat_soupx) <- "SCT"
pdf(file=paste0(plot_dir_w_date, "/",sample_id,".", annotation, ".umap_post_soupx.", estimated_contamination, "_soupX.ATAC_features_counts.pdf"), width=24, height = 8)
p1 = DimPlot(combined_data_seurat_soupx, reduction = "umap_SCT", group.by = "seurat_clusters", label = TRUE, label.size = 4)
p2 = FeaturePlot(combined_data_seurat_soupx, features = "nFeature_ATAC", min.cutoff = 0, max.cutoff = 3000, order=T) & scale_colour_gradientn(colours = c("grey90", "cyan", "deepskyblue", "forestgreen", "darkorange", "darkred"))
p3 = FeaturePlot(combined_data_seurat_soupx, features = "nCount_ATAC", min.cutoff = 0, max.cutoff = 3000, order=T) & scale_colour_gradientn(colours = c("grey90", "cyan", "deepskyblue", "forestgreen", "darkorange", "darkred"))
p1 + p2 + p3
dev.off()

ATAC_counts <- CountFragments(fragments = fragpath, cells = colnames(combined_data_seurat_soupx))
DefaultAssay(combined_data_seurat_soupx) <- "ATAC"
combined_data_seurat_soupx$ATAC_count <- ATAC_counts$reads_count
combined_data_seurat_soupx <- FRiP(combined_data_seurat_soupx, assay = "ATAC", total.fragments = "ATAC_count", col.name = "FRiP", verbose = TRUE)

pdf(file=paste0(plot_dir_w_date, "/",sample_id,".", annotation, ".umap_post_soupx.", estimated_contamination, "_soupX.ATAC_features_FRiP.pdf"), width=24, height = 8)
p1 = DimPlot(combined_data_seurat_soupx, reduction = "umap_SCT", group.by = "seurat_clusters", label = TRUE, label.size = 4)
p2 = FeaturePlot(combined_data_seurat_soupx, features = "nFeature_ATAC", min.cutoff = 0, max.cutoff = 3000, order=T) & scale_colour_gradientn(colours = c("grey90", "cyan", "deepskyblue", "forestgreen", "darkorange", "darkred"))
p3 = FeaturePlot(combined_data_seurat_soupx, features = "FRiP", min.cutoff = 0, max.cutoff = 0.3, order=T) & scale_colour_gradientn(colours = c("grey90", "cyan", "deepskyblue", "forestgreen", "darkorange", "darkred"))
p1 + p2 + p3
dev.off()

# ATAC-seq clustering
DefaultAssay(combined_data_seurat_soupx) = "ATAC"
combined_data_seurat_soupx <- RunTFIDF(combined_data_seurat_soupx)
combined_data_seurat_soupx <- FindTopFeatures(combined_data_seurat_soupx, min.cutoff = 'q0')
combined_data_seurat_soupx <- RunSVD(combined_data_seurat_soupx)
DepthCor(combined_data_seurat_soupx, n = 50)
combined_data_seurat_soupx <- RunUMAP(object = combined_data_seurat_soupx, reduction = 'lsi', dims = 2:30, reduction.name = "umap_ATAC")
combined_data_seurat_soupx <- FindNeighbors(object = combined_data_seurat_soupx, reduction = 'lsi', dims = 2:30)
combined_data_seurat_soupx <- FindClusters(object = combined_data_seurat_soupx, verbose = T, algorithm = 3, resolution = 3)

pdf(file=paste0(plot_dir_w_date, "/",sample_id,".", annotation, ".umap_ATAC_MACS_peaks.pdf"), width=8, height = 8)
DimPlot(object = combined_data_seurat_soupx, label = TRUE)
dev.off()

saveRDS(combined_data_seurat_soupx, file = paste0(seurat_objects_dir, sample_id, ".", annotation, ".postSoupX.ATAC_MACS.rds"))
```
