## Supplementary material for "A lineage-resolved multimodal single-cell atlas reveals the genomic dynamics of early *C. elegans* development": Code and external data: seurat_soupx_launcher.20240207.sh

```
#!/bin/bash
#
#SBATCH --job-name=seurat_soupx
#SBATCH --output=slurm_out/seurat_soupx.%N.%j.out
#SBATCH --error=slurm_err/seurat_soupx.%N.%j.err
#SBATCH --ntasks=1
#SBATCH -N 1
#SBATCH -n 8
#SBATCH --mem-per-cpu=10000
#SBATCH --mail-type=END,FAIL
#SBATCH --mail-user=

sample=$1
barhop=$2
ambient=$3
annotation=$4
frag_min=$5
macs_peaks=$6

Rscript scripts/seurat_soupx.20240207.R $sample $barhop $ambient $annotation $frag_min $macs_peaks
```
