## Supplementary material for "A lineage-resolved multimodal single-cell atlas reveals the genomic dynamics of early *C. elegans* development": Code and external data: STAR_dedup.20231031.sh

```
#!/bin/bash
#
#SBATCH --job-name=STAR_dedup
#SBATCH --output=slurm_out/STAR_dedup.%N.%j.out
#SBATCH --error=slurm_err/STAR_dedup.%N.%j.err
#SBATCH --ntasks=1
#SBATCH -N 1
#SBATCH -n 1
#SBATCH --mem=32G
#SBATCH --mail-type=END,FAIL
#SBATCH --mail-user=

input_bam=$1

samtools index $input_bam

dir_name=$(dirname $input_bam)
file_name=$(basename $input_bam .bam)

umi_tools dedup --stdin=$input_bam --log=$dir_name/$file_name.dedup.log --error=$dir_name/$file_name.dedup.err --extract-umi-method=tag --umi-tag=UR --cell-tag=CR --per-cell > $dir_name/$file_name.dedup.bam
```
