## Supplementary material for "A lineage-resolved multimodal single-cell atlas reveals the genomic dynamics of early *C. elegans* development": Code and external data: STAR_launcher.sh

```
#!/bin/bash
#
#SBATCH --job-name=STARsolo
#SBATCH --output=slurm_out/STARsolo.%N.%j.out
#SBATCH --error=slurm_err/STARsolo.%N.%j.err
#SBATCH --ntasks=1
#SBATCH -N 1
#SBATCH -n 12
#SBATCH --mem=240G
#SBATCH --mail-type=END,FAIL
#SBATCH --mail-user=

genome_index=$1
fastq_files_R2=$2
fastq_files_R1=$3
output_dir_prefix=$4
sample_name=$5

STAR --genomeDir $genome_index --readFilesIn $fastq_files_R2 $fastq_files_R1 --soloType CB_UMI_Simple --soloCBwhitelist whitelist/737K-arc-v1.txt --soloUMIlen 12 --soloCellFilter None --soloFeatures GeneFull --soloMultiMappers EM --readFilesCommand zcat --runThreadN 12 --outFileNamePrefix $output_dir_prefix --outSAMtype BAM SortedByCoordinate --outWigType wiggle --twopassMode None --limitBAMsortRAM 10116470442 --outSAMattributes NH HI nM AS CR UR CB UB GX GN sS sQ sM RG XS --outSAMattrRGline "ID:"$sample_name --alignIntronMax 20000 
```
