## Supplementary material for "A lineage-resolved multimodal single-cell atlas reveals the genomic dynamics of early *C. elegans* development": Code and external data: STARsolo_indexing.sh

```
#!/BIN/BASH
#
#SBATCH --job-name=STARsolo
#SBATCH --output=slurm_out/STARsolo.%N.%j.out
#SBATCH --error=slurm_err/STARsolo.%N.%j.err
#SBATCH --ntasks=1
#SBATCH -N 1
#SBATCH -n 12
#SBATCH --mem-per-cpu=4000
#SBATCH --mail-type=END,FAIL
#SBATCH --mail-user=

input_gtf=$1
genome_index=$2

STAR --runMode genomeGenerate --genomeSAindexNbases 12 --sjdbGTFfile $input_gtf --runThreadN 10 --genomeDir star_idx/$genome_index --genomeFastaFiles species/elegans/genome/elegans.fa                                                                                                                                                                        
```
