## Supplementary material for "A lineage-resolved multimodal single-cell atlas reveals the genomic dynamics of early *C. elegans* development": Code and external data: STARsolo_output_parser.20231108.py

```
import sys
import os

smple_name=sys.argv[1]
gene_annotation=sys.argv[2]
wbid_gene_name=sys.argv[3]

raw_star_path = "data/sc/rnaseq/" + smple_name + "/star_" + gene_annotation + "/star_" + gene_annotation + "_Solo.out/GeneFull/raw"
wbid_gene_name_dict={}
for line in open(wbid_gene_name):
    wbid_gene_name_dict[line.split("\t")[0]] = line.rstrip().split("\t")[1]

cmd="mkdir " + raw_star_path + "/um_reads"
os.system(cmd)

new_feats = open(raw_star_path + "/um_reads/features.tsv", "w")
for line in open(raw_star_path + "/features.tsv"):
    G=line.split("\t")
    G[1] = wbid_gene_name_dict[G[1]]
    new_feats.write("\t".join(G))

new_feats.close()

cmd="gzip " + raw_star_path + "/um_reads/features.tsv"
os.system(cmd)
cmd="cp " + raw_star_path + "/barcodes.tsv " + raw_star_path + "/um_reads/barcodes.tsv; gzip " + raw_star_path + "/um_reads/barcodes.tsv"
os.system(cmd)
cmd="cp " + raw_star_path + "/UniqueAndMult-EM.mtx " + raw_star_path + "/um_reads/matrix.mtx; gzip " + raw_star_path + "/um_reads/matrix.mtx"
os.system(cmd)
```
