## Supplementary material for "A lineage-resolved multimodal single-cell atlas reveals the genomic dynamics of early *C. elegans* development": Code and external data: sum_TPM_per_gene.py

```
import sys

transcript_gene_table=sys.argv[1]
kallisto_output=sys.argv[2]
output_table=sys.argv[3]

new = open(output_table, "w")

transcript_gene_dict = {}
for line in open(transcript_gene_table):
    G=line.rstrip().split("\t")
    transcript_gene_dict[G[1]] = G[0]

gene_TPM = {}
for line in open(kallisto_output):
    G=line.rstrip().split("\t")
    if G[0] in transcript_gene_dict:
        if transcript_gene_dict[G[0]] not in gene_TPM:
            gene_TPM[transcript_gene_dict[G[0]]] = 0
        gene_TPM[transcript_gene_dict[G[0]]] += eval(G[4])

for i in gene_TPM:
    temp = i + "\t" + str(gene_TPM[i]) + "\n"
    new.write(temp)

new.close()
```
