## Supplementary material for "A lineage-resolved multimodal single-cell atlas reveals the genomic dynamics of early *C. elegans* development": Code and external data: transient_gene_promoters.motif_enrichment.R

```
# plot enrichment of motifs in transient promoters
library(Hmisc)
library(png)

current_date= paste(unlist(strsplit(as.character(Sys.Date()), "-")), collapse="")
plot_dir_w_date = paste0("paper_figures/", current_date)
dir.create(plot_dir_w_date, recursive=TRUE)

all_motifs_to_test = read.table("transient_genes/fimo/transient_genes.all.promoters/fimo.sites.0001.enrichment_in_transient_promoters.txt")


# plot enrichment
pdf(file=paste0(plot_dir_w_date, "/transient_promoters_motifs.enrichment.pdf"), width=12, height = 10, useDingbats = F)
par(mar=c(6, 3, 4, 4), xpd=TRUE, mfrow = c(1,2))
k = barplot(t(matrix(c(all_motifs_to_test$V4), ncol = 1, byrow = F)), beside = T, horiz = T, plot = F)
plot(1, type="n", xlab="", ylab="", xlim=c(0, 10), ylim=c(1, k[1, dim(k)[2] + 0.5]), xaxt = "n", yaxt = "n", bty = "n")
# read in sequentially all fwd/rev logos and add them to the left of the barplot
for (motif_n in c(1:dim(k)[2])){
  motif_name = all_motifs_to_test[motif_n,1]
  img_fwd<-readPNG(paste0("transient_genes/memechip/transient_genes.all.promoters.w_6_15/logos/", motif_name, ".fwd.png"))
  img_rev<-readPNG(paste0("transient_genes/memechip/transient_genes.all.promoters.w_6_15/logos/", motif_name, ".rev.png"))
  rasterImage(img_fwd,1, k[1,motif_n]-0.5, 4, k[1,motif_n]+0.5)
  rasterImage(img_rev,5, k[1,motif_n]-0.5, 9, k[1,motif_n]+0.5)
}
barplot(t(matrix(c(all_motifs_to_test$V4), ncol = 1, byrow = F)), beside = T, horiz = T, 
        main = "motif enrichment", col = c("grey"), xlab = "fraction of transient promoters with motif")
dev.off()
```
