## Supplementary material for "A lineage-resolved multimodal single-cell atlas reveals the genomic dynamics of early *C. elegans* development": Code and external data: transient_genes_DESeq.20241113.R

```
library(cluster)
library(Seurat)
library(Signac)
library(Hmisc)
library(RColorBrewer)
library(presto)
library(rtracklayer)
library(ComplexHeatmap)
library(DESeq2)
library(ggplot2)
#library(clusterProfiler)
#library(dplyr)

source("data/external_data/wormcat_function.R")

current_date= paste(unlist(strsplit(as.character(Sys.Date()), "-")), collapse="")
plot_dir_w_date = paste0("paper_figures/", current_date)
dir.create(plot_dir_w_date, recursive=TRUE)

# load RDS and WBid table
combined_seurat = readRDS("seurat_objects/wt_all.all_samples.all_cells.WS285_extended_no_ovlp.final_peakset_IDR.rds")
wb_id_common_names = read.table("species/elegans/gene_annotation/c_elegans.PRJNA13758.WS285.canonical_geneset.filtered.gene_name.txt")

# integrate cell_cycle info
cell_cycle_table = read.table("cell_cycle_prediction/cell_cycle_RF.txt")
combined_seurat$cell_cycle = cell_cycle_table$V2

# sample similar number of cells per each lineage
# do not include lineages present during/immediately after gastrulation
cell_lineage_levels <- c("ABx", "ABxx", "ABxxx", "ABxxxxx", "ABxxxxxx",
                         "EMS", "E", "Exx","Exxx", 
                         "MS", "MSx", "MSxxx", "MSxxxx", 
                         "C", "Cx", "Cxxx", 
                         "Dx", "DxxCxxx")

# sample up to 200 late cell cycle cells per lineage
cell_lineage_n_all=table(combined_seurat$cell_lineage[combined_seurat$cell_lineage %nin% c("unassigned", "P2", "P3", "P4", "Z2Z3",
                                                                                           "ABxxxx", "Ex", "MSxx", "Cxx") & combined_seurat$cell_cycle == "late"])
cell_lineage_barcodes_all = list()

sample_cell_n = 200
set.seed(100)
for (cell_l in cell_lineage_levels){
  if (as.numeric(cell_lineage_n_all[cell_l]) <= sample_cell_n){
    cell_l_barcodes = colnames(combined_seurat[,combined_seurat$cell_lineage == cell_l & combined_seurat$cell_cycle == "late"])
  }
  else {
    cell_l_barcodes = sample(colnames(combined_seurat[,combined_seurat$cell_lineage == cell_l & combined_seurat$cell_cycle == "late"]), size = sample_cell_n, replace = F)
  }
  cell_lineage_barcodes_all[[cell_l]] = cell_l_barcodes
}

combined_seurat_sample = subset(combined_seurat, cells = unlist(cell_lineage_barcodes_all))
#DimPlot(combined_seurat_sample, reduction = "umapWNN", label = T, group.by = "cell_lineage") & NoLegend()

# get aggregate UMI counts per each lineage
countData_lineage = as.data.frame(AggregateExpression(combined_seurat_sample, assays = "RNA", group.by = "cell_lineage", slot="counts"))
names(countData_lineage) = sapply(strsplit(names(countData_lineage), split='RNA.', fixed=TRUE), function(x) (x[2]))
countData_lineage = countData_lineage[order(factor(names(countData_lineage), levels = cell_lineage_levels))]

colData <- data.frame(row.names=colnames(countData_lineage),
                      cell_lineage = cell_lineage_levels,
                      pregastr_late = relevel(factor(c("early", "early", "early","late", "late",
                                                       "early", "early", "late", "late",
                                                       "early", "early", "late", "late",
                                                       "early", "early", "late",
                                                       "late", "late")), ref = "late"))

set.seed(42)

# run DESeq2
dds <- DESeqDataSetFromMatrix(countData=countData_lineage, colData=colData, design=~pregastr_late)
dds <- DESeq(dds)

# get results
res <- results(dds, alpha = 0.001, altHypothesis = "greater", lfcThreshold = 1)

# ditinguish transient genes with high/low expression in late cells
k = counts(dds, normalized=TRUE)
high_late = row.names(k[which(apply(k[,c(4,5,8,9,12,13,16,17,18)], 1, function(x){quantile(x, 0.25)}) > 25),])

DE_id_lfc_2_se7 = row.names(res[res$padj < 0.001 & !is.na(res$padj) & res$log2FoldChange > 2 & res$lfcSE < 0.7,])

DE_id_lfc_2_se7_hl = DE_id_lfc_2_se7[DE_id_lfc_2_se7 %in% high_late]
DE_id_lfc_2_se7_ll = DE_id_lfc_2_se7[DE_id_lfc_2_se7 %nin% high_late]

# output gene lists
dir.create("transient_genes")
write.table(wb_id_common_names[wb_id_common_names$V2 %in% DE_id_lfc_2_se7_hl,1], file = "transient_genes/transient_genes.high_postgastrulation_exp.txt", quote = F, row.names = F, col.names = "gene_name", append = F)
write.table(wb_id_common_names[wb_id_common_names$V2 %in% DE_id_lfc_2_se7_ll,1], file = "transient_genes/transient_genes.low_postgastrulation_exp.txt", quote = F, row.names = F, col.names = "gene_name", append = F)

# get average expression per cell lineage / cell type
# levels/colors for ordering
cell_type_levels <-  c("twocell", "ABx", "ABax", "ABpx", "earlyABxxx", "ABxxx", "ABala", "ABalp", "ABara", "ABarp", "ABpxa", "ABpxp", 
                       "earlyABxxxx", "ABxxxx", "ABalaa", "ABalap", "ABalpa", "ABalpp", "ABaraa", "ABarap", "ABarpa", "ABarpp", "ABpxaa", "ABpxap", "ABpxpa", "ABpxpp",
                       "ABxxxxx10", "ABxxxxx15", "ABxxxxx22", "ABxxxxx25", "ABxxxxx40", "ABxxxxx49", "ABxxxxx52", "ABxxxxx58", "ABxxxxx60", "ABxxxxx61", "ABxxxxx71", "ABxxxxx74", "ABxxxxx83",
                       "ABxxxxxx20", "ABxxxxxx23", "ABxxxxxx24", "ABxxxxxx3", "ABxxxxxx33", "ABxxxxxx36", "ABxxxxxx37", "ABxxxxxx47", "ABxxxxxx65", "ABxxxxxx69", "ABxxxxxx72", "ABxxxxxx73", "ABxxxxxx77", "ABxxxxxx82", "ABxxxxxx86", "ABxxxxxx9",
                       "earlyEMS", "EMS", "earlyEandMS", "E", "earlyEx", "Ex", "earlyExx", "Exx", "Exxx", 
                       "MS", "earlyMSx", "MSx", "MSa", "MSp", "earlyMSxx", "MSxx", "MSxa", "MSxp", "MSxxx28", "MSxxx29", "MSxxx42", "MSxxxx57",
                       "earlyC", "C", "earlyCx", "Ca", "Cp", "earlyCxx", "Cxx", "Cxa", "Cxp", "Cxxx59", "Cxxx78", "Cxxx79",
                       "D", "Dx", "DxxCxxx7",
                       "unassigned", "unassigned1", "unassigned14", "unassigned17", "unassigned5", "unassigned56", "unassigned6", "unassigned81", "unassigned85",
                       "P2", "P3", "P4", "Z2Z3")
cell_lineage_levels <- c("twocell", 
                         "ABx", "ABxx", "ABxxx", "ABxxxx", "ABxxxxx", "ABxxxxxx",
                         "EMS", "EandMS", "E", "Ex", "Exx","Exxx", 
                         "MS", "MSx", "MSxx", "MSxxx", "MSxxxx", 
                         "C", "Cx", "Cxx", "Cxxx", 
                         "D", "Dx", "DxxCxxx",
                         "unassigned", 
                         "P2", "P3", "P4", "Z2Z3")
#stage_levels_colnames <- c("twocell", "g4cell", "g8cell", "g15cell", "g26cell", "g46cell", "g87morecell", "Pcell", "Z2Z3", "unassigned")
stage_levels <- c("twocell", "4cell", "8cell", "15cell", "26cell", "46cell", "87morecell", "Pcell", "Z2Z3", "unassigned")

avgExp_lineage = as.data.frame(AverageExpression(combined_seurat, assays = "RNA", group.by = "cell_lineage", features = DE_id_lfc_2_se7))
names(avgExp_lineage) <- sapply(strsplit(names(avgExp_lineage), split='RNA.', fixed=TRUE), function(x) (x[2]))
avgExp_lineage = avgExp_lineage[order(factor(names(avgExp_lineage), levels = cell_lineage_levels))]

DE_id_lineage_heatmap = Heatmap(as.matrix(log10(avgExp_lineage+1)), name = "transient gene exp", cluster_columns = F, 
                             cluster_rows = T, use_raster = F, row_dend_reorder = TRUE, row_title_rot = 0, column_title_rot = 90, 
                             column_title_gp = gpar(cex = 0.8), row_title_gp = gpar(cex = 0.8), show_row_dend = F, show_row_names = F)
DE_id_lineage_heatmap = draw(DE_id_lineage_heatmap, padding = unit(c(8, 5, 10, 4), "mm"))

avgExp_type = as.data.frame(AverageExpression(combined_seurat, assays = "RNA", group.by = "cell_type", features = DE_id_lfc_2_se7))
names(avgExp_type) <- sapply(strsplit(names(avgExp_type), split='RNA.', fixed=TRUE), function(x) (x[2]))
avgExp_type = avgExp_type[order(factor(names(avgExp_type), levels = cell_type_levels))]


DE_id_type_heatmap = Heatmap(as.matrix(log10(avgExp_type+1)), name = "transient gene exp", cluster_columns = F, 
                             cluster_rows = T, use_raster = F, row_dend_reorder = TRUE, row_title_rot = 0, column_title_rot = 90, 
                             column_title_gp = gpar(cex = 0.8), row_title_gp = gpar(cex = 0.8), show_row_dend = F, show_row_names = F)
DE_id_type_heatmap = draw(DE_id_type_heatmap, padding = unit(c(8, 5, 10, 4), "mm"))

avgExp_stage = as.data.frame(AverageExpression(combined_seurat, assays = "RNA", group.by = "stage", features = DE_id_lfc_2_se7))
names(avgExp_stage) <- sapply(strsplit(names(avgExp_stage), split='RNA.', fixed=TRUE), function(x) (x[2]))
avgExp_stage = avgExp_stage[order(factor(names(avgExp_stage), levels = stage_levels))]
avgExp_stage = avgExp_stage[,c(1:9)]
names(avgExp_stage) = stage_levels[1:9]

DE_id_stage_heatmap = Heatmap(as.matrix(log10(avgExp_stage+1)), name = "transient gene exp", cluster_columns = F, 
                             cluster_rows = T, use_raster = F, row_dend_reorder = TRUE, row_title_rot = 0, column_title_rot = 90, 
                             column_title_gp = gpar(cex = 0.8), row_title_gp = gpar(cex = 0.8), show_row_dend = F, show_row_names = F)
DE_id_stage_heatmap = draw(DE_id_stage_heatmap, padding = unit(c(8, 5, 10, 4), "mm"))


# plot
pdf(file=paste0(plot_dir_w_date, "/transient_genes_heatmap.pdf"), width=16, height = 16)
print(DE_id_lineage_heatmap)
print(DE_id_type_heatmap)
print(DE_id_stage_heatmap)
dev.off()


# clusterProfiler
# transient_GO_BP <- compareCluster(geneCluster = list("transient"=wb_id_common_names[wb_id_common_names$V2 %in% DE_id_lfc_2_se7, 1]), fun = "enrichGO", ont = "BP", pvalueCutoff = 0.05, minGSSize = 3, pAdjustMethod = "BH", keyType = "WORMBASE", OrgDb='org.Ce.eg.db')
# transient_GO_MF <- compareCluster(geneCluster = list("transient"=wb_id_common_names[wb_id_common_names$V2 %in% DE_id_lfc_2_se7, 1]), fun = "enrichGO", ont = "MF", pvalueCutoff = 0.05, minGSSize = 3, pAdjustMethod = "BH", keyType = "WORMBASE", OrgDb='org.Ce.eg.db')
# transient_GO_CC <- compareCluster(geneCluster = list("transient"=wb_id_common_names[wb_id_common_names$V2 %in% DE_id_lfc_2_se7, 1]), fun = "enrichGO", ont = "CC", pvalueCutoff = 0.05, minGSSize = 3, pAdjustMethod = "BH", keyType = "WORMBASE", OrgDb='org.Ce.eg.db')
# 
# p1 = clusterProfiler::dotplot(transient_GO_BP, showCategory = 5)
# p2 = clusterProfiler::dotplot(transient_GO_MF, showCategory = 5)
# p3 = clusterProfiler::dotplot(transient_GO_CC, showCategory = 5)
# 
# pdf(file=paste0(plot_dir_w_date, "/transient_genes_ClusterProfiler.pdf"), width=12, height = 6)
# print(p1 | p2 | p3)
# dev.off()


# VlnPlots per gene
# Idents(combined_seurat) = "cell_lineage"
# Idents(combined_seurat) <- factor(x = Idents(combined_seurat), levels = cell_lineage_levels)
# color_code_lineage = c("grey",
#                        "dodgerblue4", "deepskyblue4", "deepskyblue3", "deepskyblue2","deepskyblue1","cyan",
#                        "darkorange", "darkorange3","gold4", "gold3", "gold2", "gold1",
#                        "darkred", "firebrick4", "firebrick3", "firebrick2", "firebrick1",
#                        "seagreen4", "seagreen3", "seagreen2", "seagreen1", 
#                        "mediumpurple1", "mediumpurple2", "mediumpurple4",
#                        "grey90", 
#                        rep("darkorchid1", 3), "darkorchid4")
# 
# pdf(file=paste0(plot_dir_w_date, "/transient_genes_vlnplots.pdf"), width=16, height = 12)
# for (i in c(1:(ceiling(length(DE_id_lfc_2_se7_ll)/4)))){
#   genes_to_plot = c(DE_id_lfc_2_se7_ll[4*(i-1) + 1],
#                     DE_id_lfc_2_se7_ll[4*(i-1) + 2],
#                     DE_id_lfc_2_se7_ll[4*(i-1) + 3],
#                     DE_id_lfc_2_se7_ll[4*(i-1) + 4])
#   p1 <- VlnPlot(combined_seurat, features = genes_to_plot[1], cols = color_code_lineage, log = T, assay = "RNA", slot = "data", raster = T) & NoLegend()
#   p2 <- VlnPlot(combined_seurat, features = genes_to_plot[2], cols = color_code_lineage, log = T, assay = "RNA", slot = "data", raster = T) & NoLegend()
#   p3 <- VlnPlot(combined_seurat, features = genes_to_plot[3], cols = color_code_lineage, log = T, assay = "RNA", slot = "data", raster = T) & NoLegend()
#   p4 <- VlnPlot(combined_seurat, features = genes_to_plot[4], cols = color_code_lineage, log = T, assay = "RNA", slot = "data", raster = T) & NoLegend()
#   print( (p1 | p2) / (p3 | p4) )
# }
# dev.off()
# 
# 
```
