## Supplementary material for "A lineage-resolved multimodal single-cell atlas reveals the genomic dynamics of early *C. elegans* development": Code and external data: ungapped_promoter_selection.py

```
import sys

promoter_file = sys.argv[1]
to_remove = sys.argv[2]
output_file = sys.argv[3]

prom_to_rm = {}
for line in open(to_remove):
    G=line.rstrip().split("\t")
    if G[0] not in prom_to_rm:
        prom_to_rm[G[0]] = []
    prom_to_rm[G[0]].append(G[1])

new = open(output_file, "w")
for line in open(promoter_file):
    G=line.rstrip().split("\t")
    if G[3] not in prom_to_rm:
        new.write(line)
    else:
        if G[5] not in prom_to_rm[G[3]]:
            new.write(line)

new.close()

        
```
