## Supplementary material for "A lineage-resolved multimodal single-cell atlas reveals the genomic dynamics of early *C. elegans* development": Code and external data: wormcat_launcher.R

```
source("data/external_data/wormcat_function.R")

args = commandArgs(trailingOnly=TRUE)

input_list <- args[1]
output_dir <- args[2]
wc_title <- args[3]

annotation_wormcat = "data/external_data/whole_genome_v2_nov-11-2021.csv"

worm_cat_fun(file_to_process = input_list,
             output_dir = output_dir, 
             title = wc_title, 
             annotation_file = "data/external_data/whole_genome_v2_nov-11-2021.csv", 
             input_type = "Wormbase.ID", zip_files = F)
```
