## Supplementary material for "A lineage-resolved multimodal single-cell atlas reveals the genomic dynamics of early *C. elegans* development": Code and external data: yapc_chipseq.20240325.sh

```
#!/bin/bash
#
#SBATCH --job-name=yapc
#SBATCH --output=slurm_out/yapc.%N.%j.err
#SBATCH --error=slurm_err/yapc.%N.%j.err
#SBATCH --ntasks=1
#SBATCH -N 1
#SBATCH -n 6
#SBATCH --mem-per-cpu=4000
#SBATCH --mail-type=END,FAIL
#SBATCH --mail-user=

rep1=$1
rep2=$2
sample_name=$3

input_dir=data/chipseq/macs

mkdir -p "data/chipseq/yapc/"$sample_name
yapc --smoothing-window-width 150 --fixed-peak-halfwidth 100 "data/chipseq/yapc/"$sample_name"/"$sample_name".model.smooth_100_yapc" $sample_name $input_dir"/"$rep1".model/"$rep1"_treat_pileup_sorted.bw" $input_dir"/"$rep2".model/"$rep2"_treat_pileup_sorted.bw"
```
