## Supplementary Figures for "A lineage-resolved multimodal single-cell atlas reveals the genomic dynamics of early *C. elegans* development"

### Supplementary Figure 1

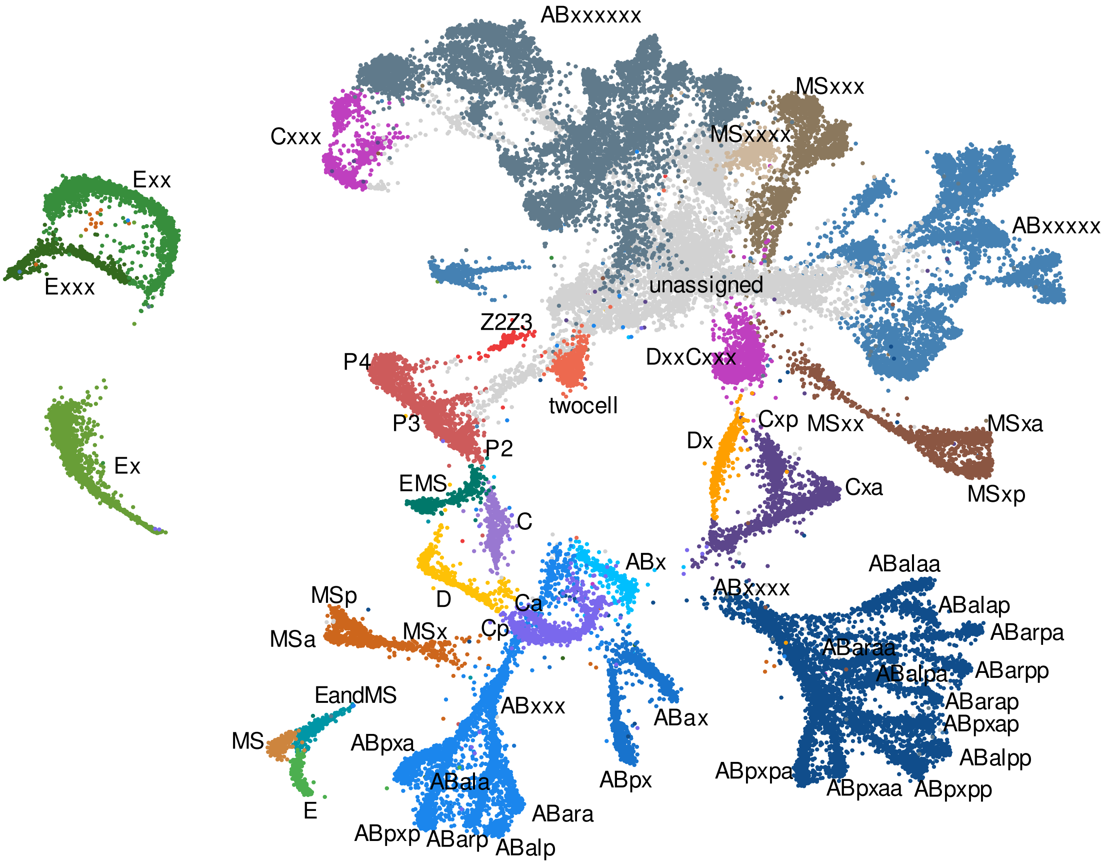

### Supplementary Figure 2

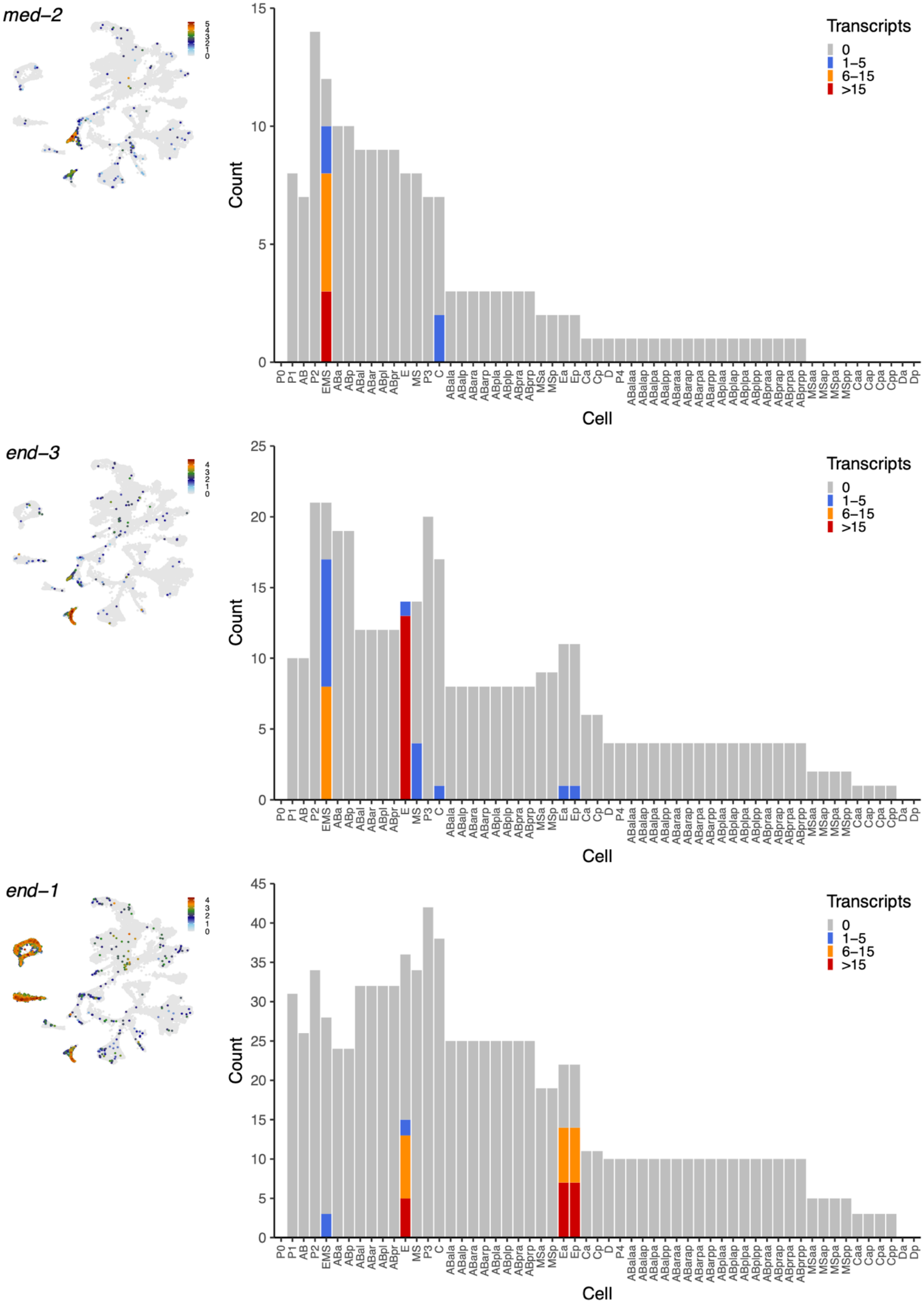

### Supplementary Figure 2 (continued)

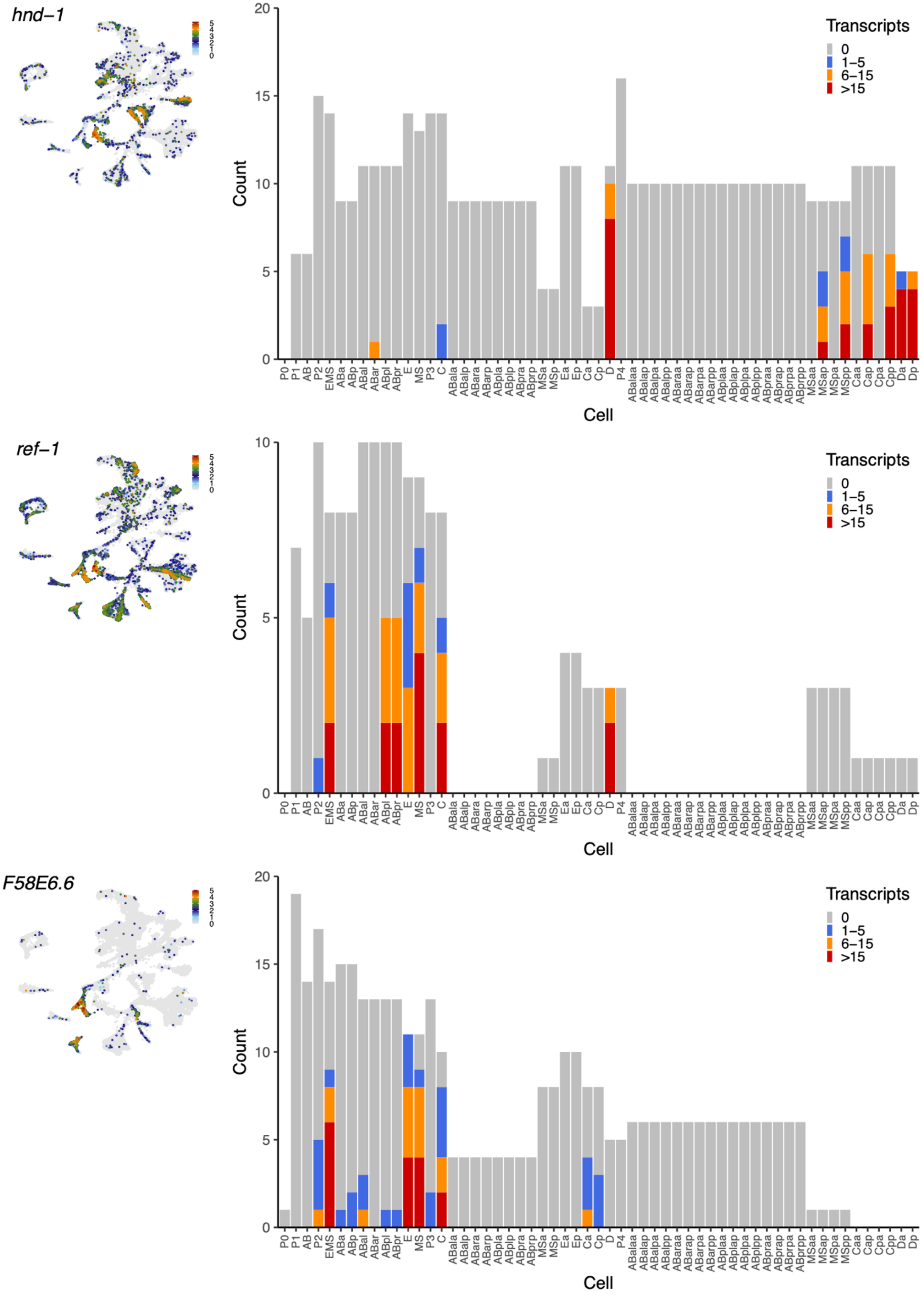

### Supplementary Figure 2 (continued)

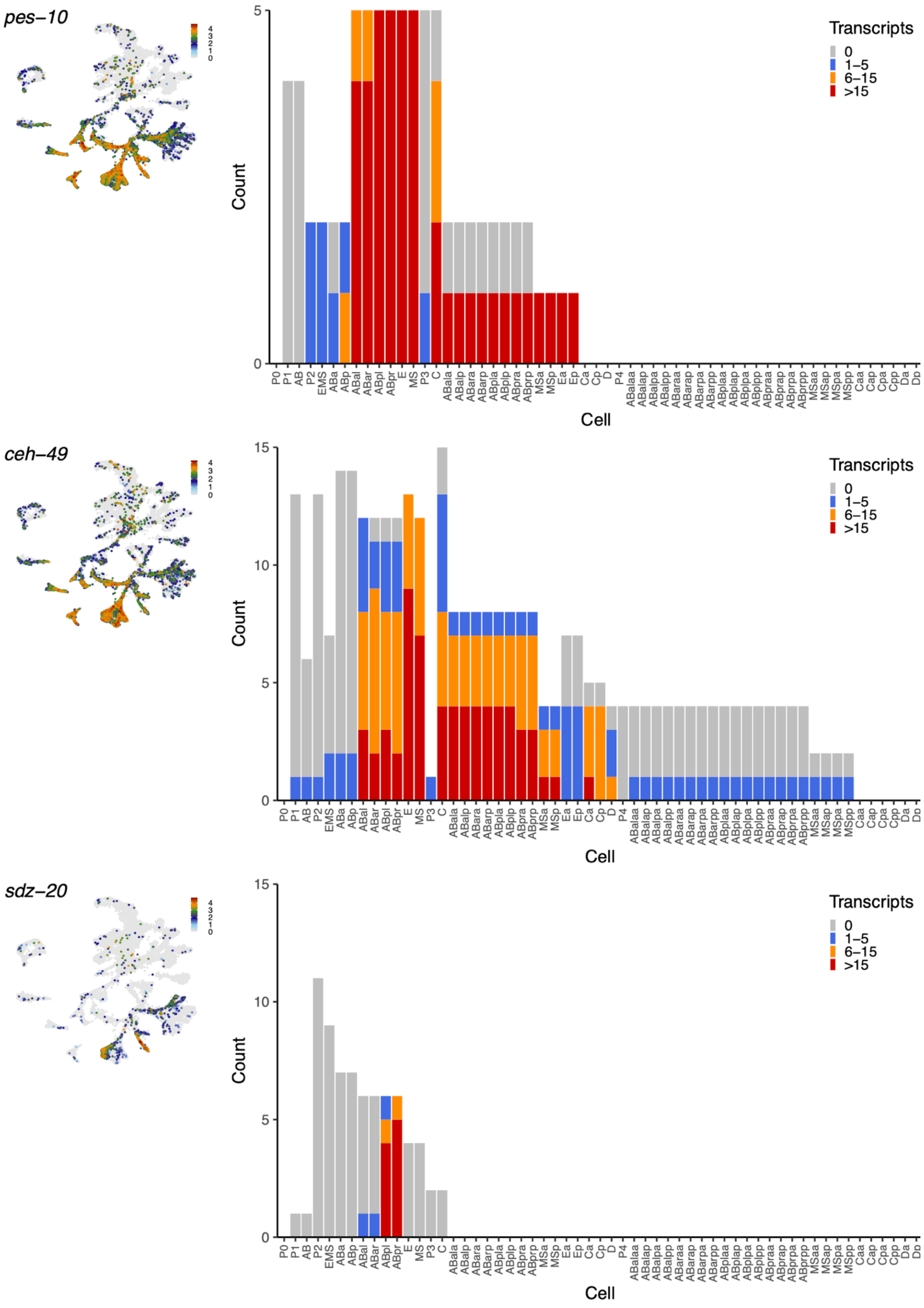

Supplementary Figure 2 (continued)

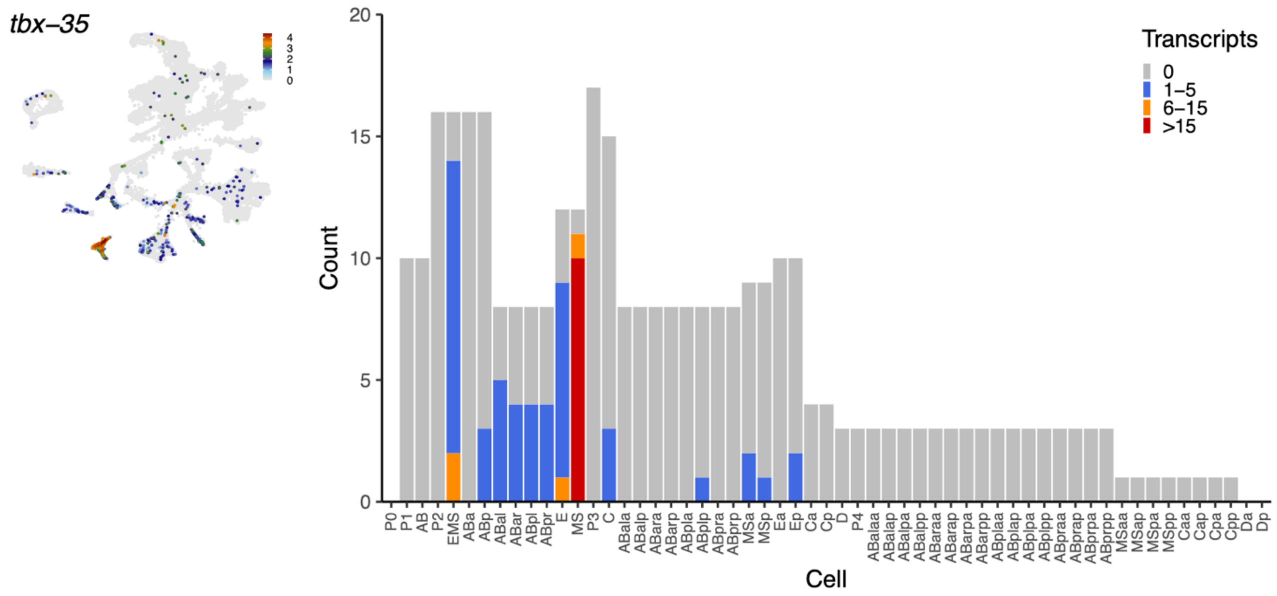


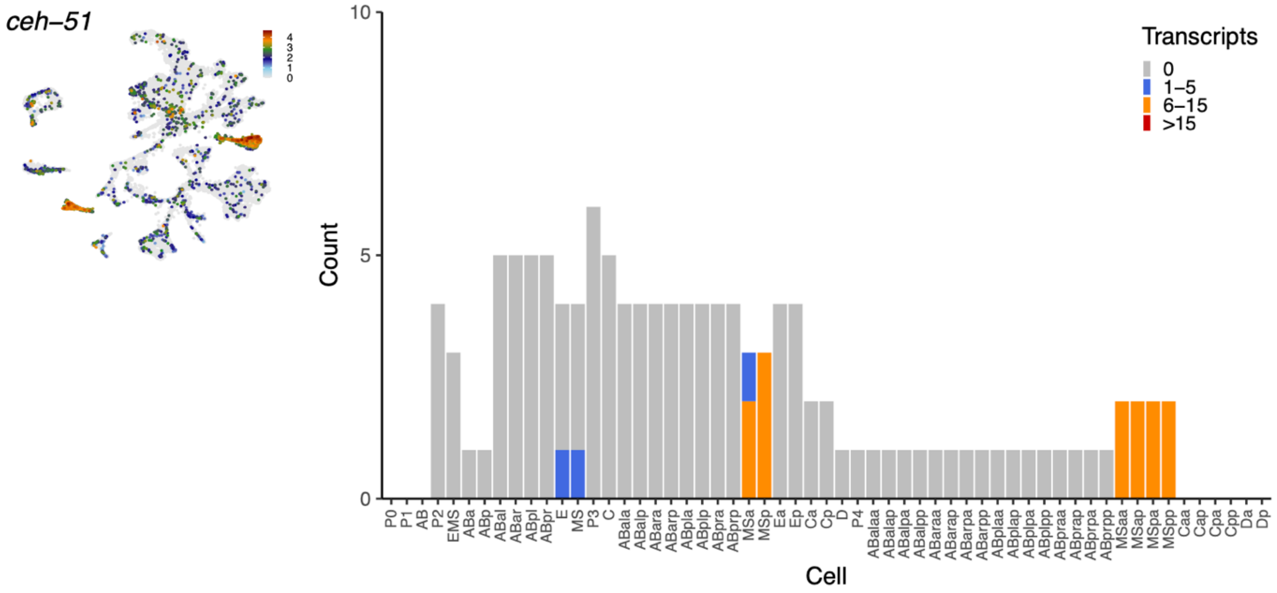


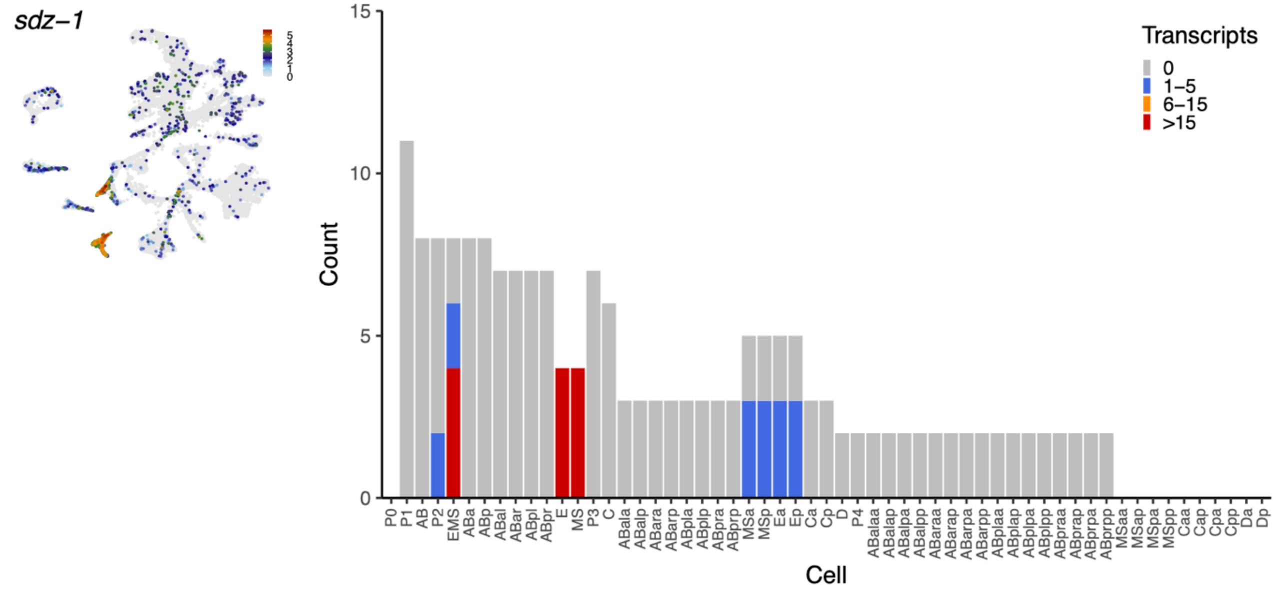

### Supplementary Figure 2 (continued)

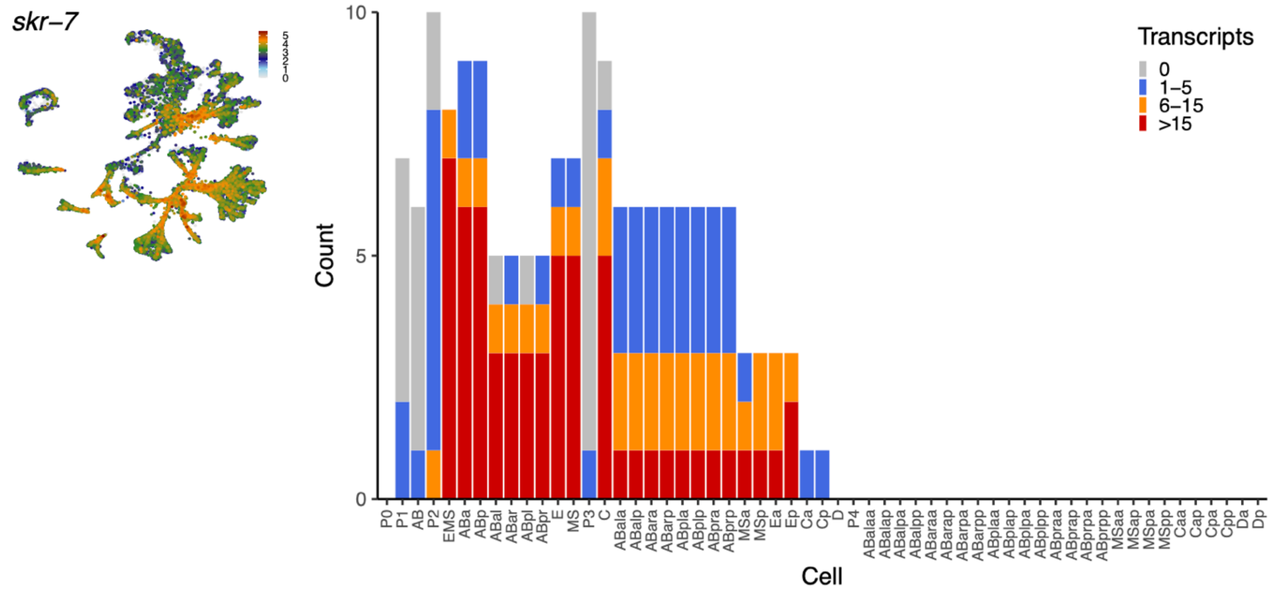


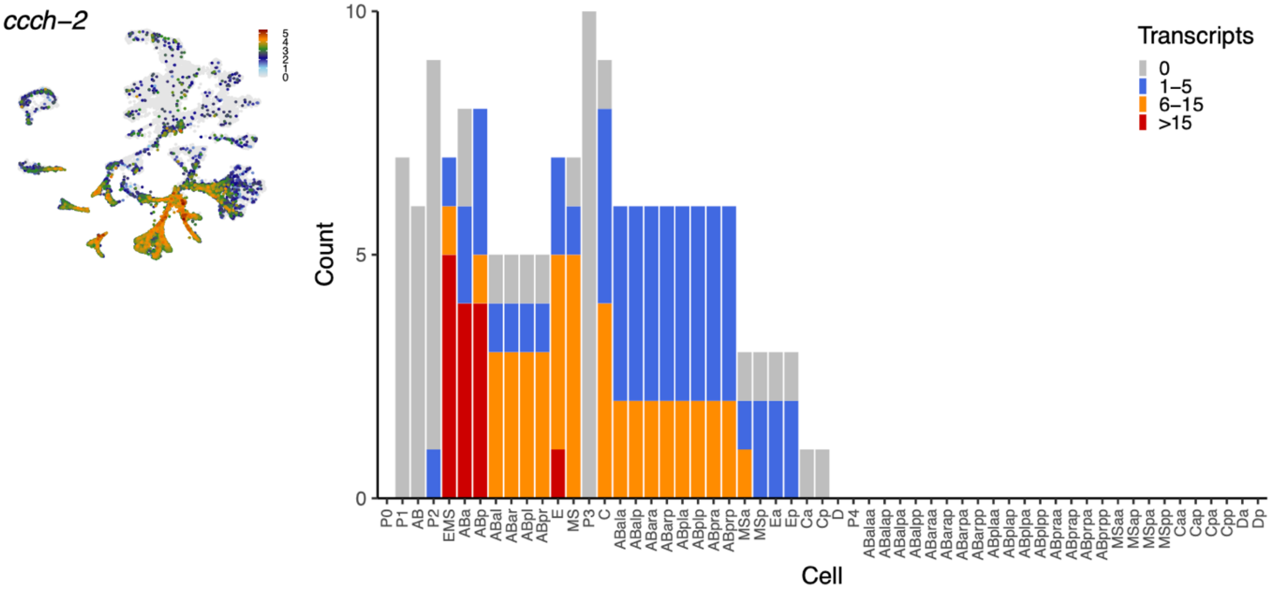


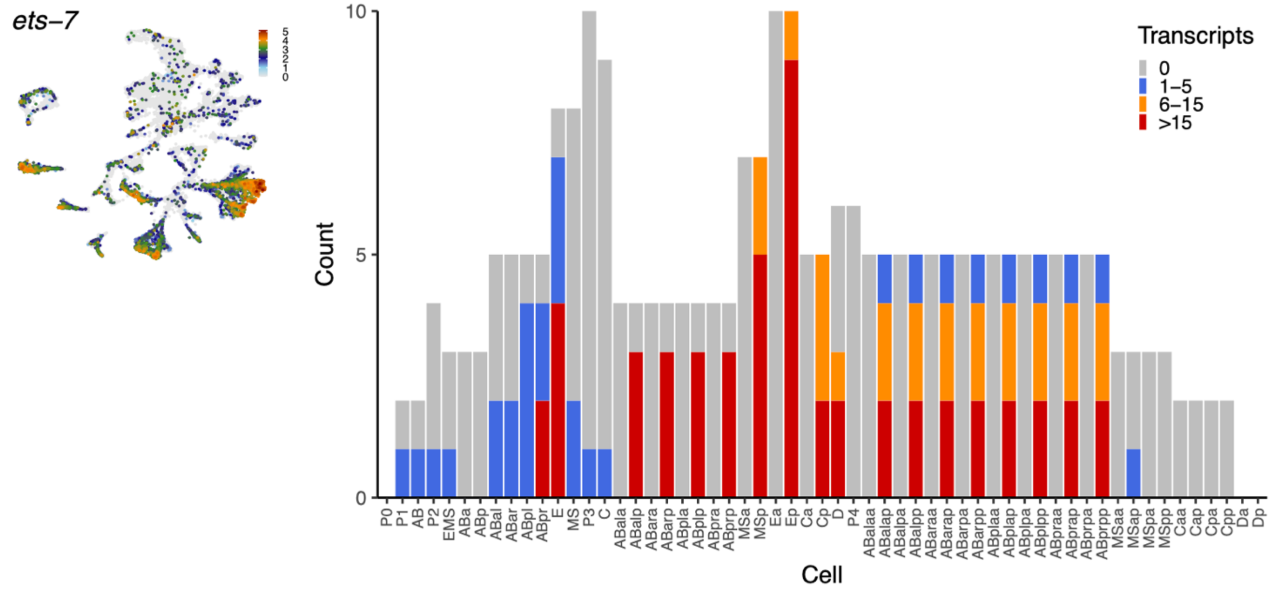

### Supplementary Figure 2 (continued)

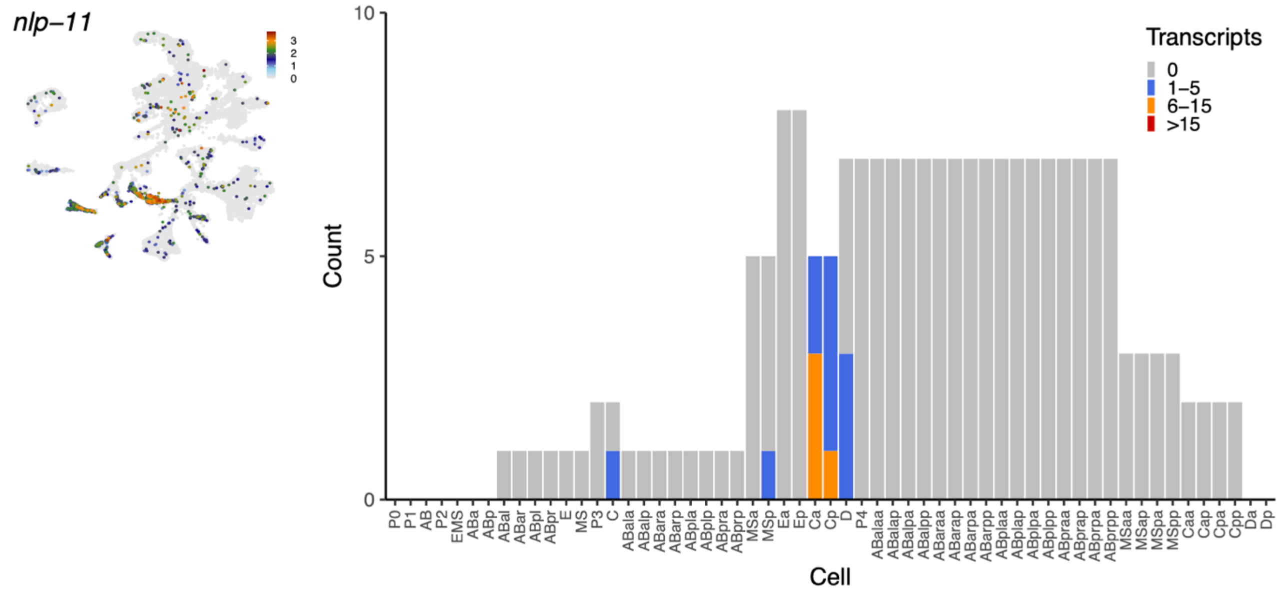


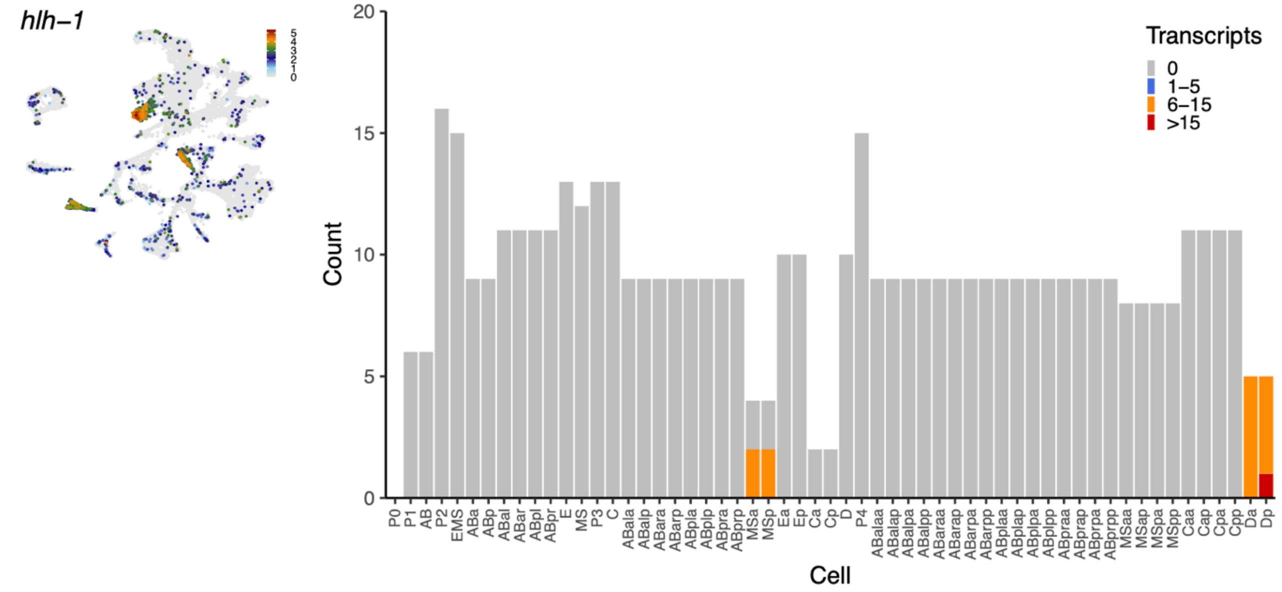


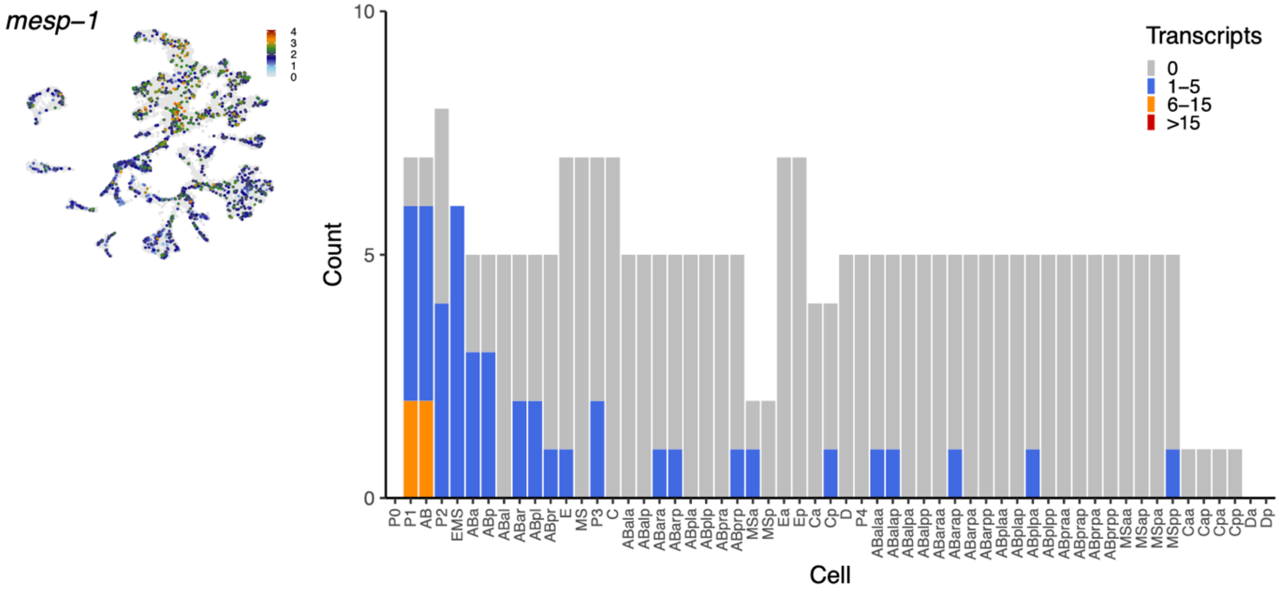

### Supplementary Figure 2 (continued)

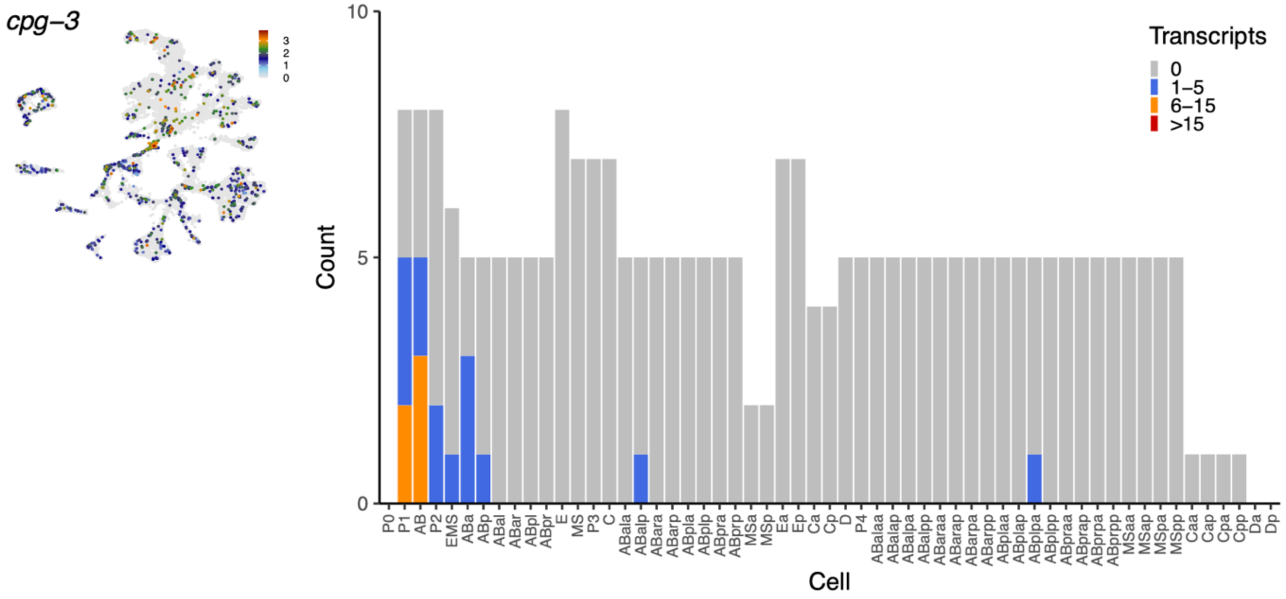

Supplementary Figure 3

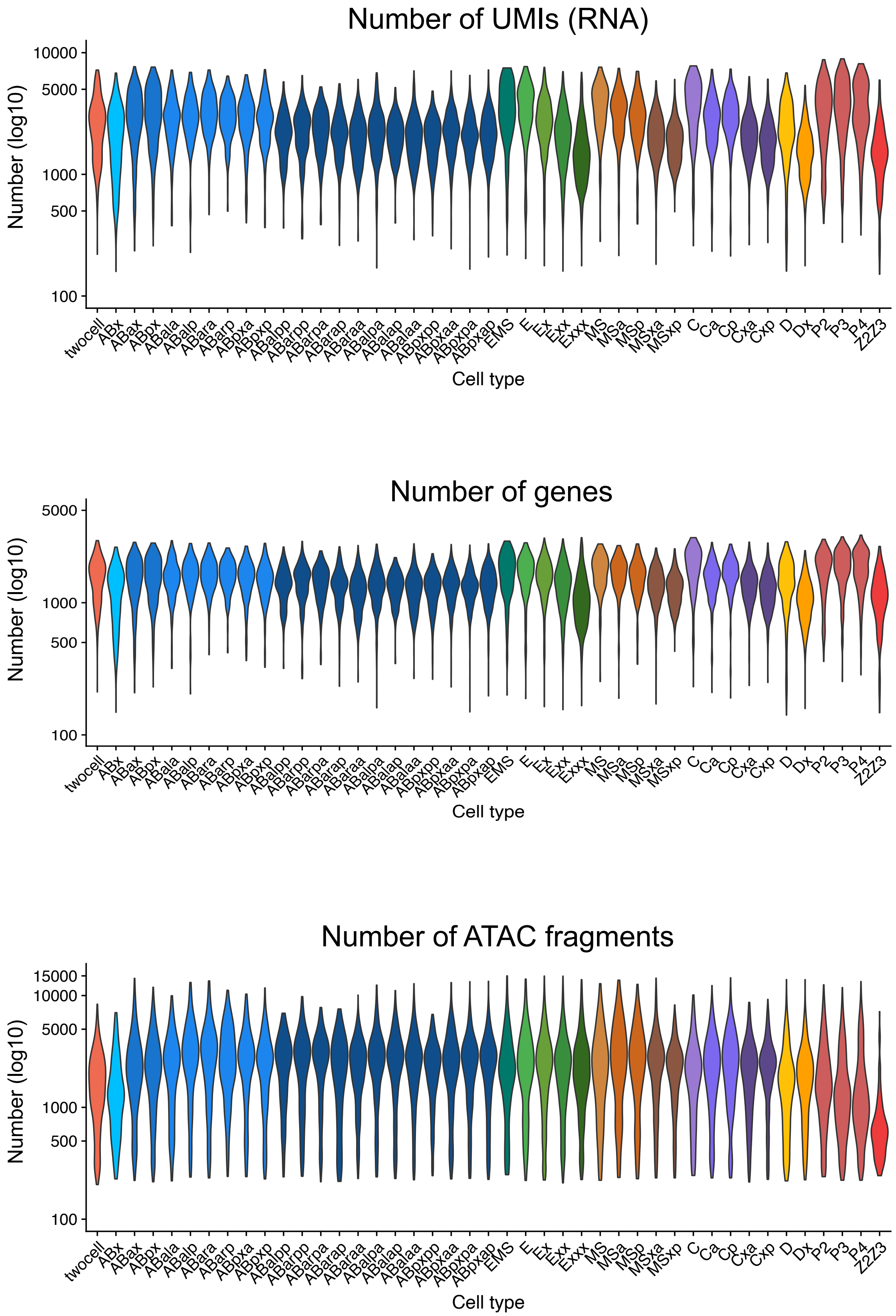

Supplementary Figure 4

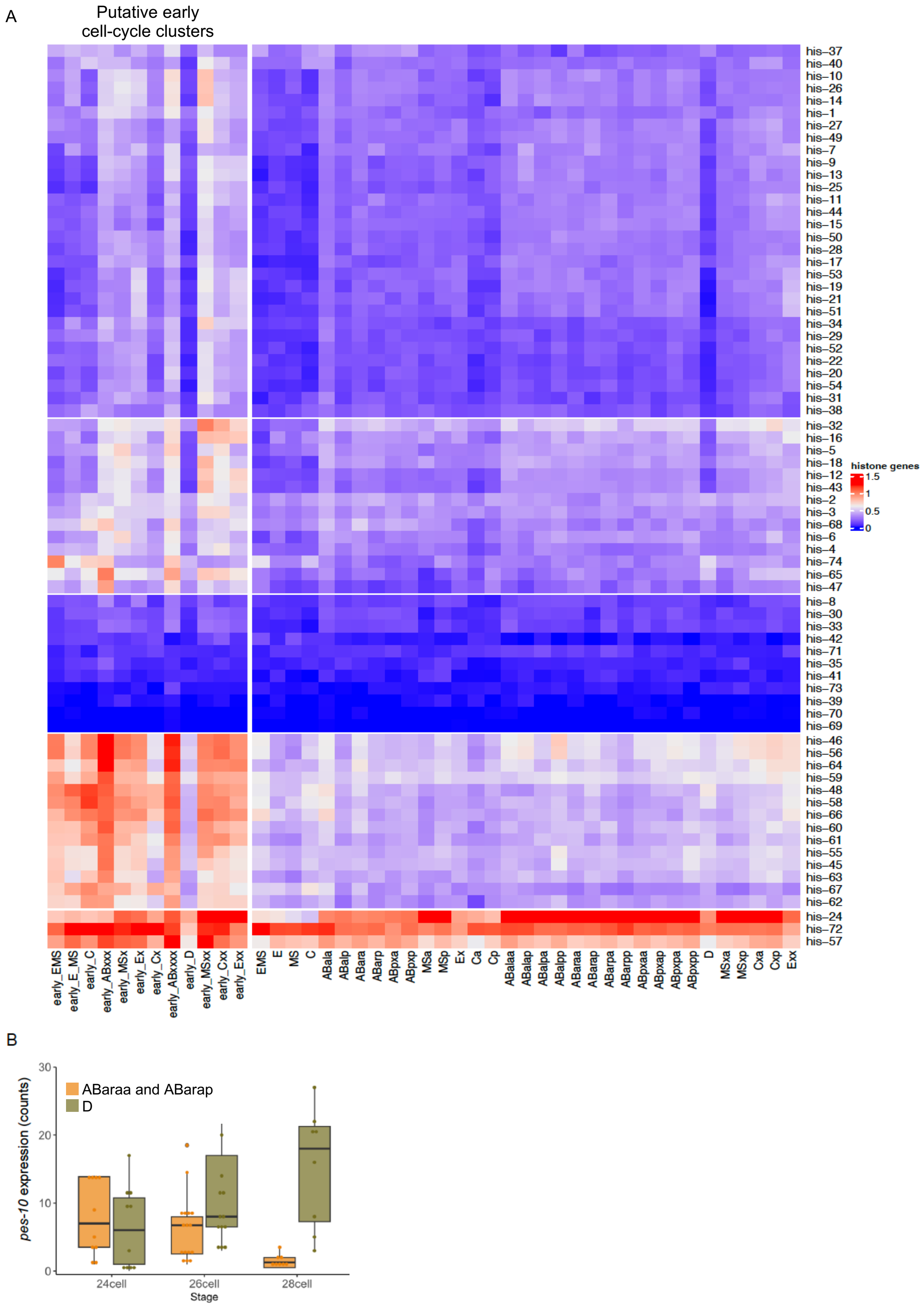

Supplementary Figure 5

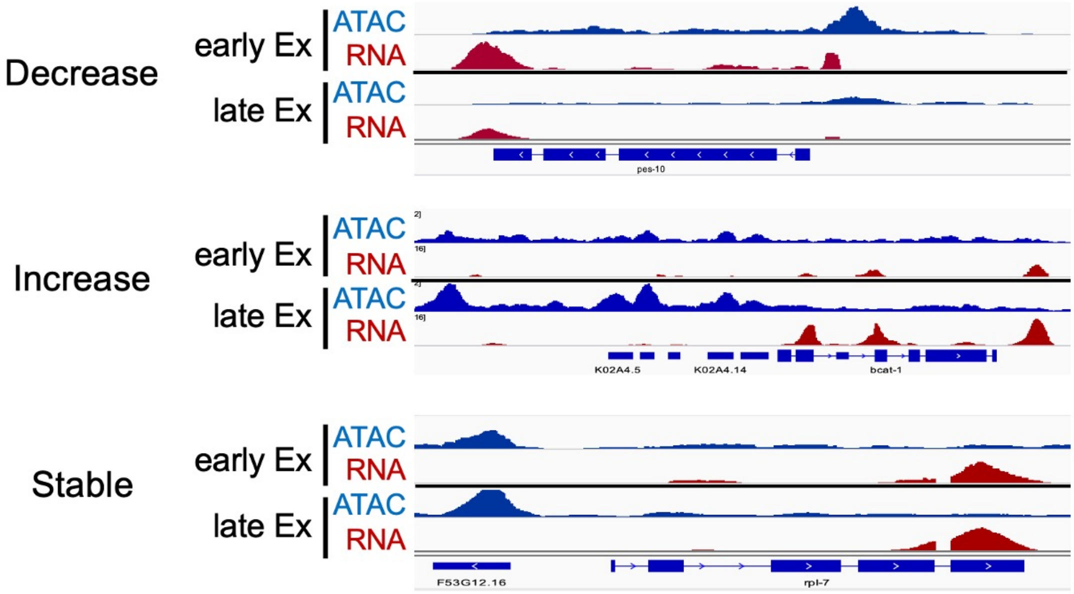

### Supplementary Figure 6

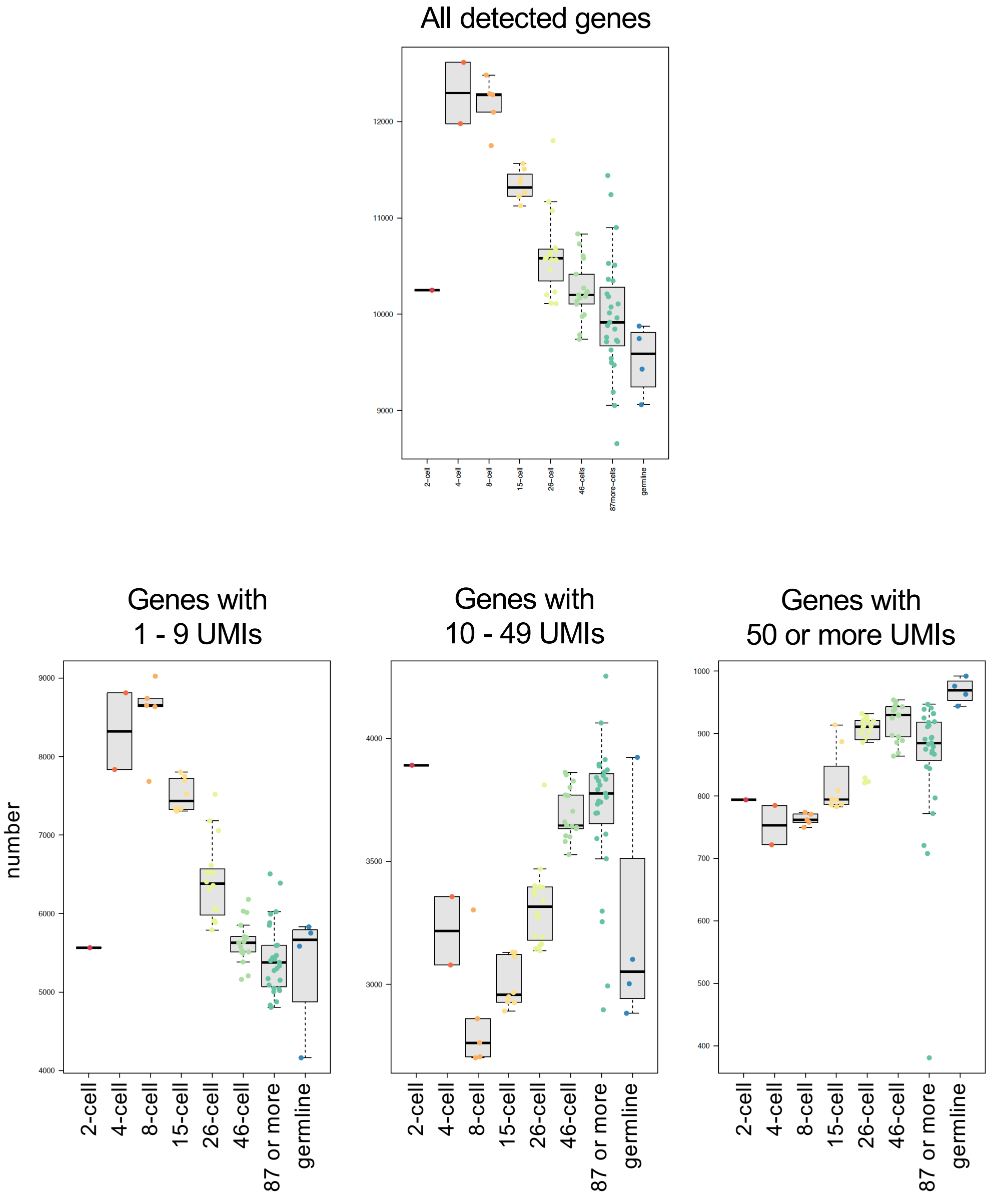

### Supplementary Figure 7

A

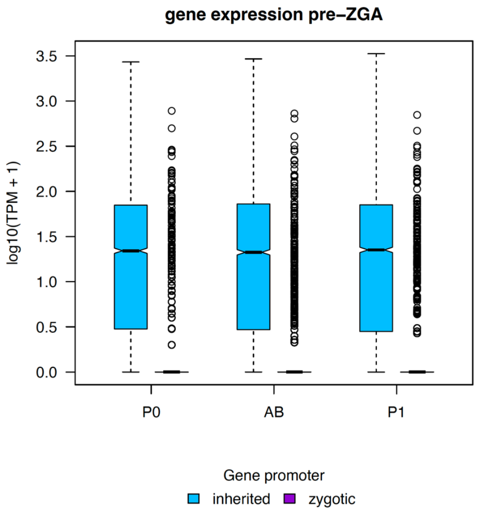


B

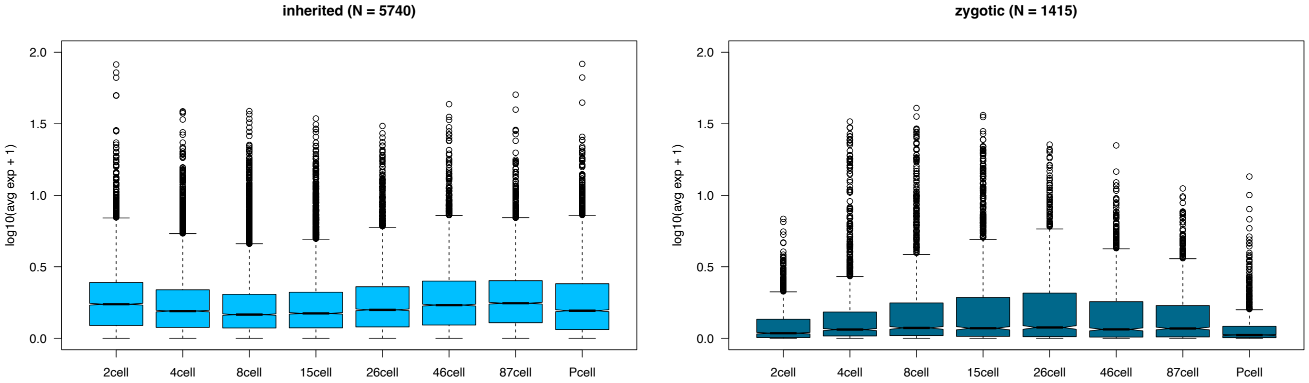

### Supplemental Figure 8

#### A Genes with inherited promoters

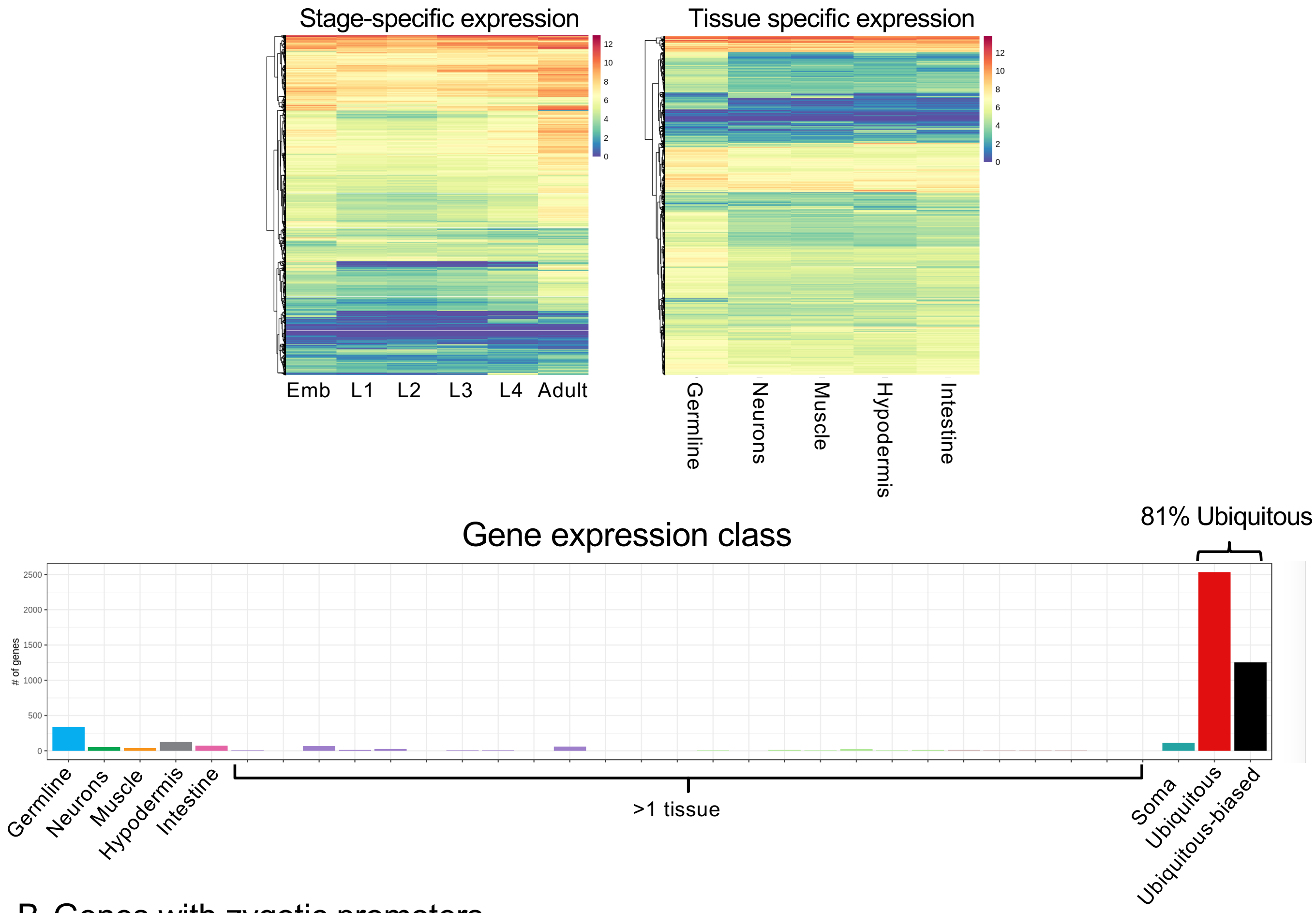


#### B Genes with zygotic promoters

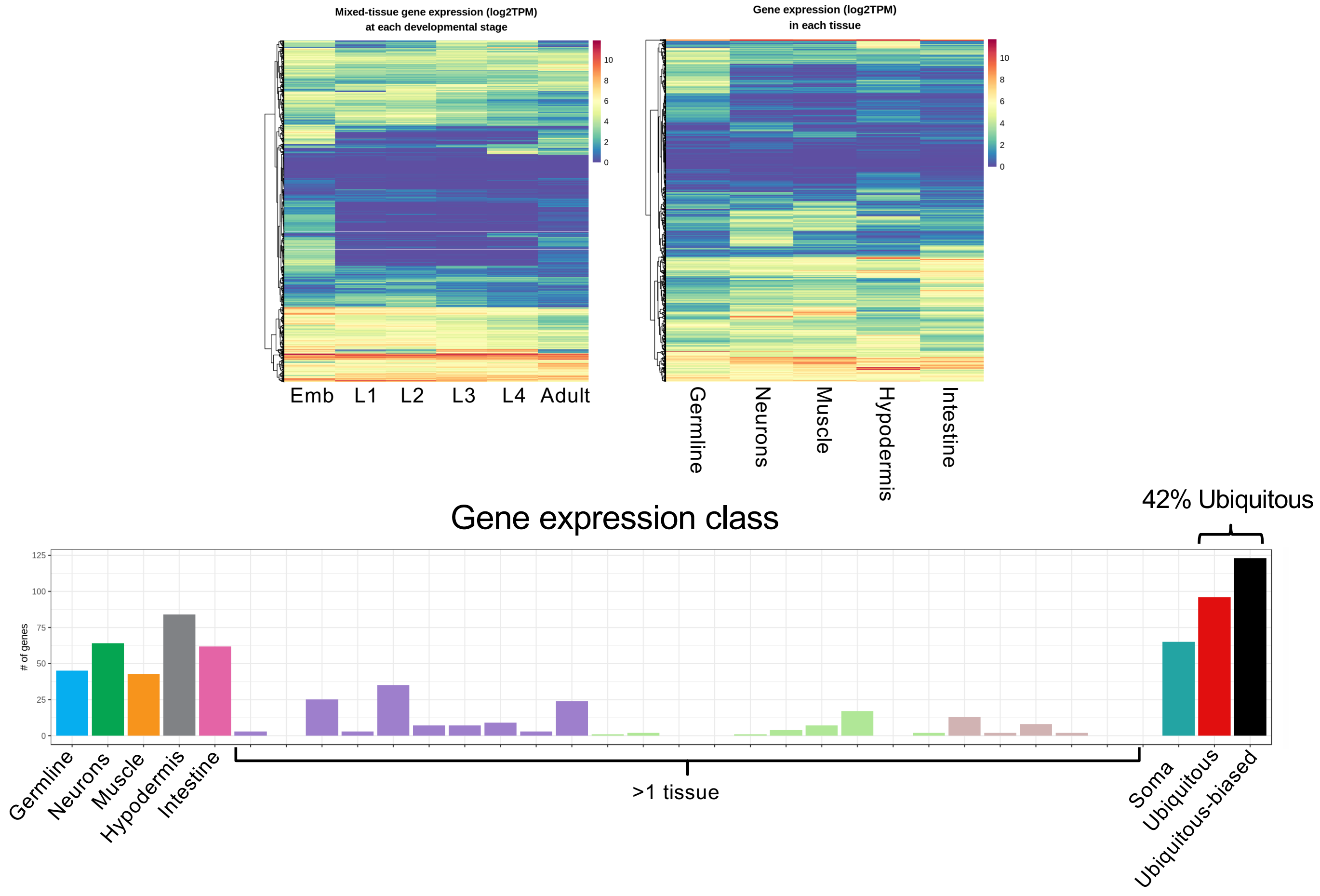

Supplementary Figure 9

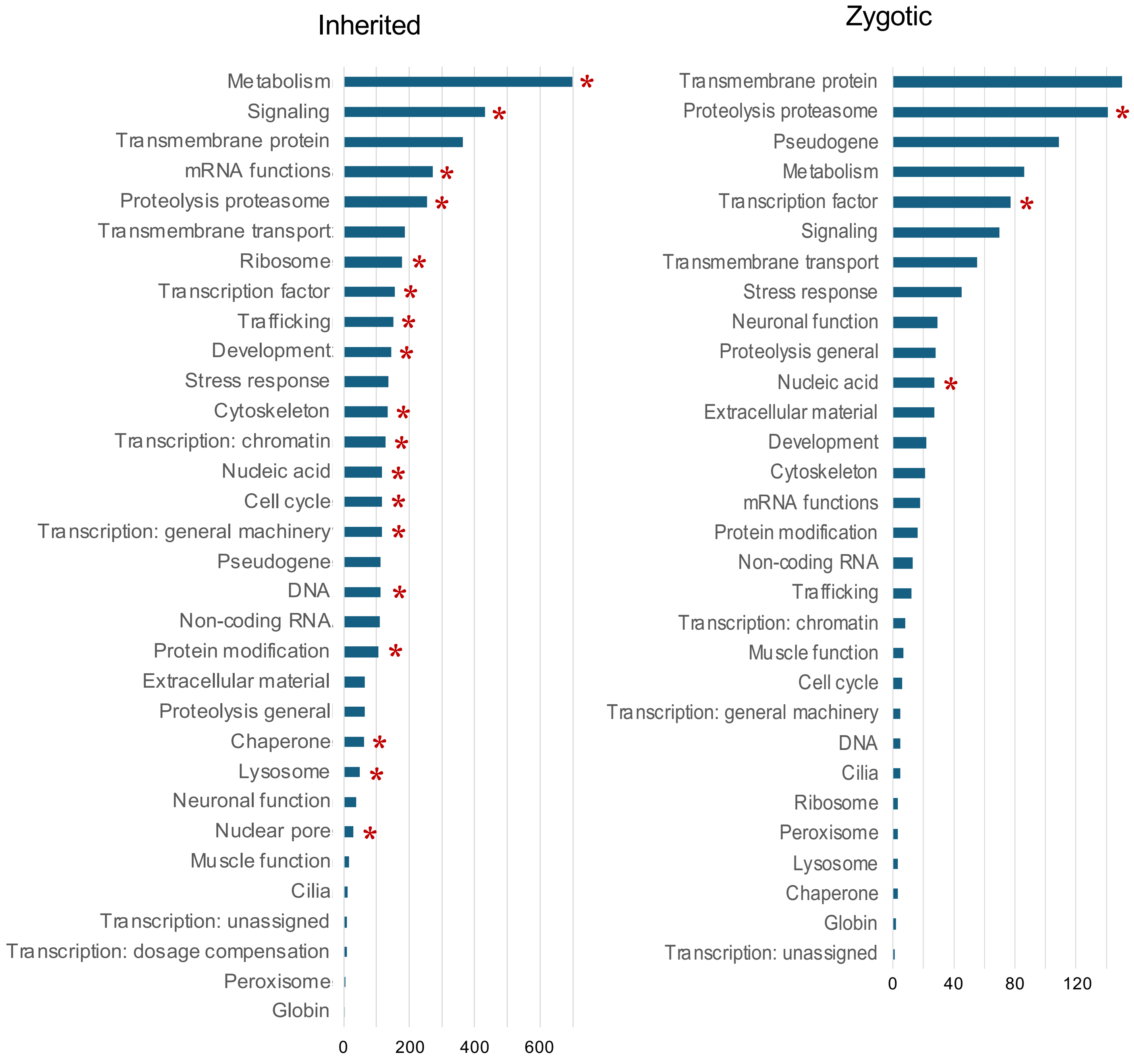


* significantly over-represented class

Supplementary Figure 10

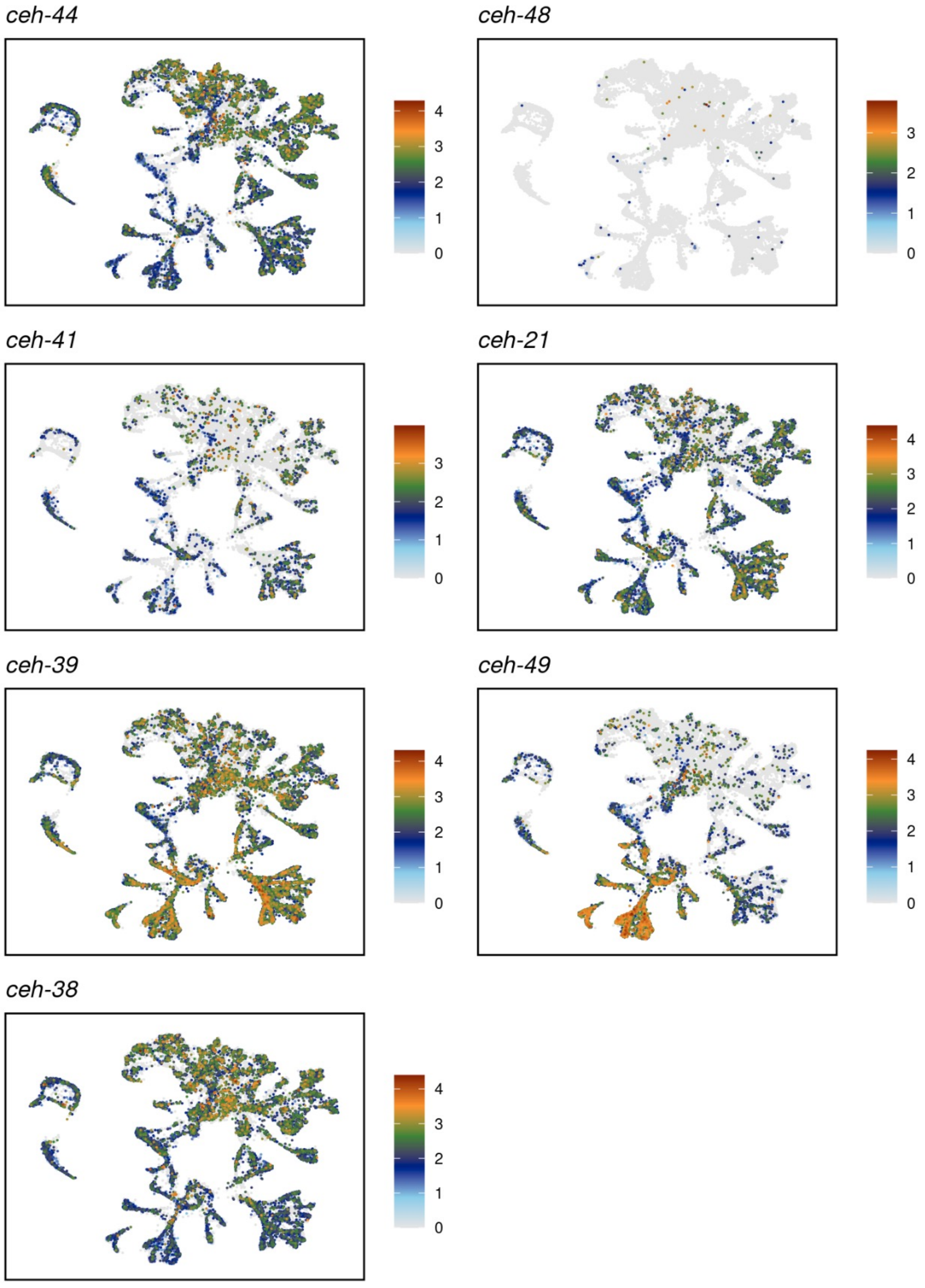

Supplementary Figure 11

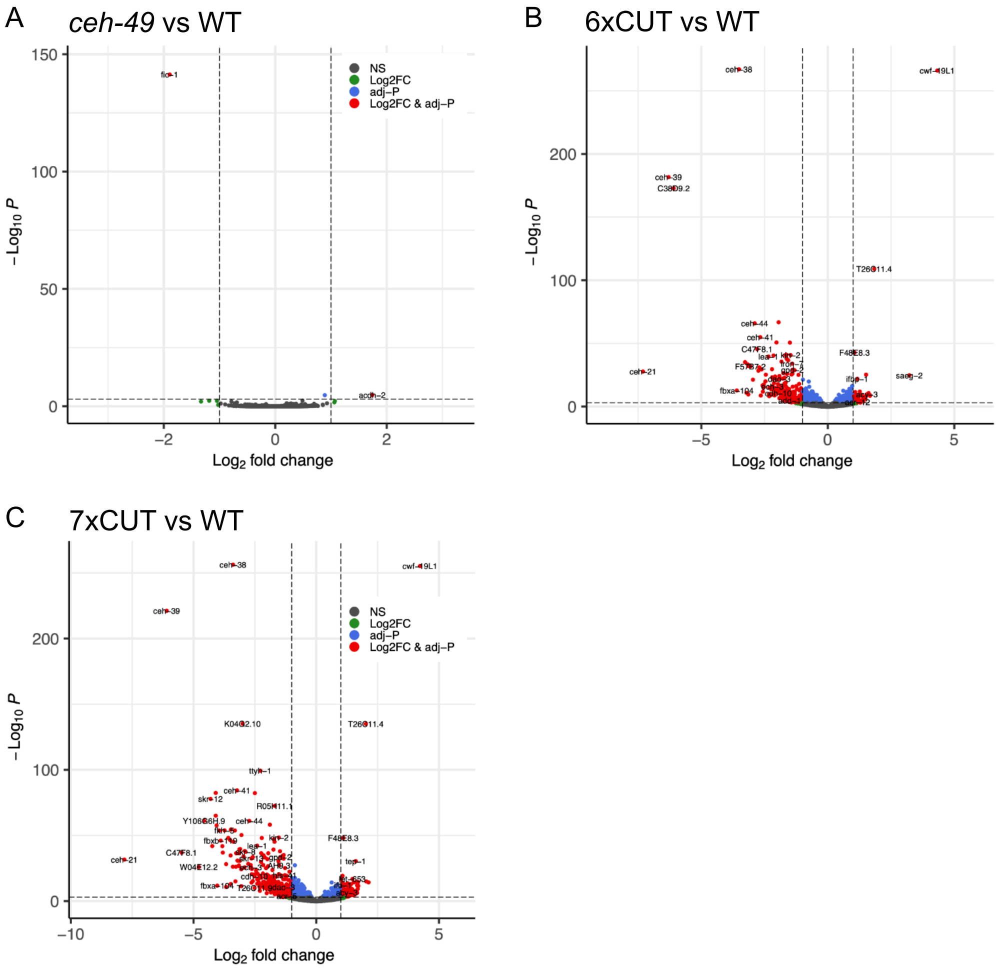

Supplementary Figure 12

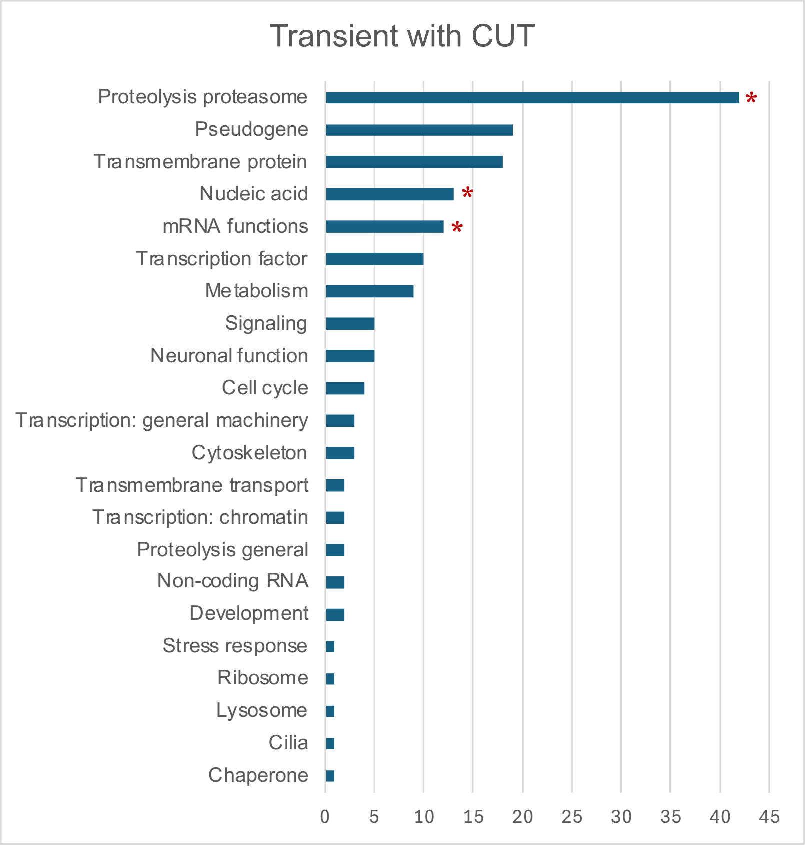


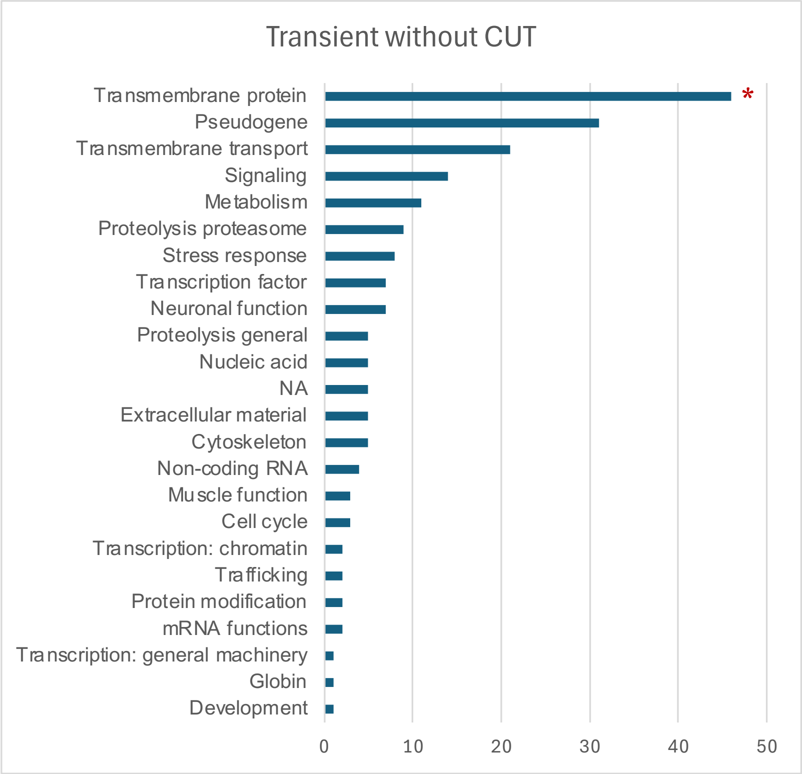


* significantly over-represented class

Supplementary Figure 13

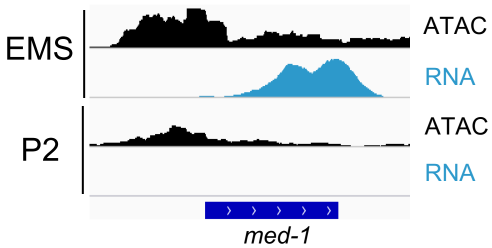
